# PubPlant – a continuously updated online resource for sequenced and published plant genomes

**DOI:** 10.1101/2025.03.12.642823

**Authors:** Rainer Schwacke, Marie Bolger, Björn Usadel

## Abstract

Advances in next-generation sequencing technologies over the last decade have substantially reduced the cost and effort required to sequence plant genomes. Whereas early efforts focused primarily on economically important crops and model species, attention has now turned to a broader range of plants, including those with larger and more complex genomes. In 2024, the genomes of 500 plant species were published, including 370 sequenced for the first time. Tracking and providing access to published plant genomes (now covering more than 1800 species) is an invaluable service for plant researchers. PubPlant is an online resource that serves this purpose by cataloging published plant genome sequences and offering multiple visualizations (https://www.plabipd.de/pubplant_main.html). It includes a chronology of genome publications, and cladograms to display the phylogenetic relationships among the sequenced plants. An overview diagram for seed plants highlights taxonomic orders and families with sequenced species, and reveals those that have been overlooked thus far. As a use case for PubPlant, we evaluated the status of sequenced food crops. We found that the five plant families featuring the most food crops were those containing the most sequenced plant species.

## Introduction

The first plant genome to be sequenced was that of the laboratory model *Arabidopsis thaliana*, published in the year 2000 (The Arabidopsis Genome Initiative, 2000). It took 10 further years to achieve the milestone of 20 sequenced plant genomes, but only another 4 years to pass 100 genomes, and by the year 2020 the milestone of 500 plant genomes had been achieved. Remarkably, 500 additional plant genomes were sequenced in the next 2 years (one tenth of the time needed for the first 500). This progress is still accelerating, mainly due to the advent of third-generation long-read sequencing technologies (Jiao and Schneeberger, 2017) and their continual refinement (Dumschott et al., 2020; Pucker et al., 2022). Advances in sequencing technologies have gone hand in hand with the development of more powerful bioinformatics algorithms for the assembly and annotation of genomic data. Genome assembly tools such as hifiasm (Cheng et al., 2021a) and verkko (Rautiainen et al., 2023) can integrate data from the two most popular long-read sequencing technologies, namely nanopore sequencing developed by Oxford Nanopore Technologies and single-molecule real-time sequencing commercialized as the PacBio platform by Pacific Biosciences (van Rengs et al., 2022).

The advances in third-generation sequencing have reduced the cost and effort needed for genome sequencing to such a degree that it is now within the means of even moderately-funded research groups. It is therefore unsurprising that recent years have seen a dramatic increase in the number of plant whole-genome sequencing projects (Fig. 1). By the end of 2024, more than 1800 plant species had been sequenced, more than 500 of which have been sequenced twice and more than 200 of which have been sequenced three or more times. The number of plant species that have been sequenced is increasing steadily, but the number of individual sequenced genomes is increasing at a much faster rate due to the re-sequencing of the same species multiple times. Re-sequencing is driven by the repetition of earlier sequencing efforts using more advanced technologies to obtain better and more complete genomes as well as intra-species pan-genome projects involving the sequencing of multiple individuals (such as different varieties, cultivars or ecotypes) within a species to explore the full genomic landscape (Golicz et al., 2016). Prominent examples of staple crop pan-genomes include maize (Hufford et al., 2021), barley (Jayakodi et al., 2020), wheat (Jiao et al., 2024), rice (Qin et al., 2021) and potato (Tang et al., 2022; Bozan et al., 2023), as well as fruit crops such as tomato (Gao et al., 2019; Zhou et al., 2022), and beverage crops such as tea (Chen et al., 2023; Tariq et al., 2024).

**Fig. 1.**
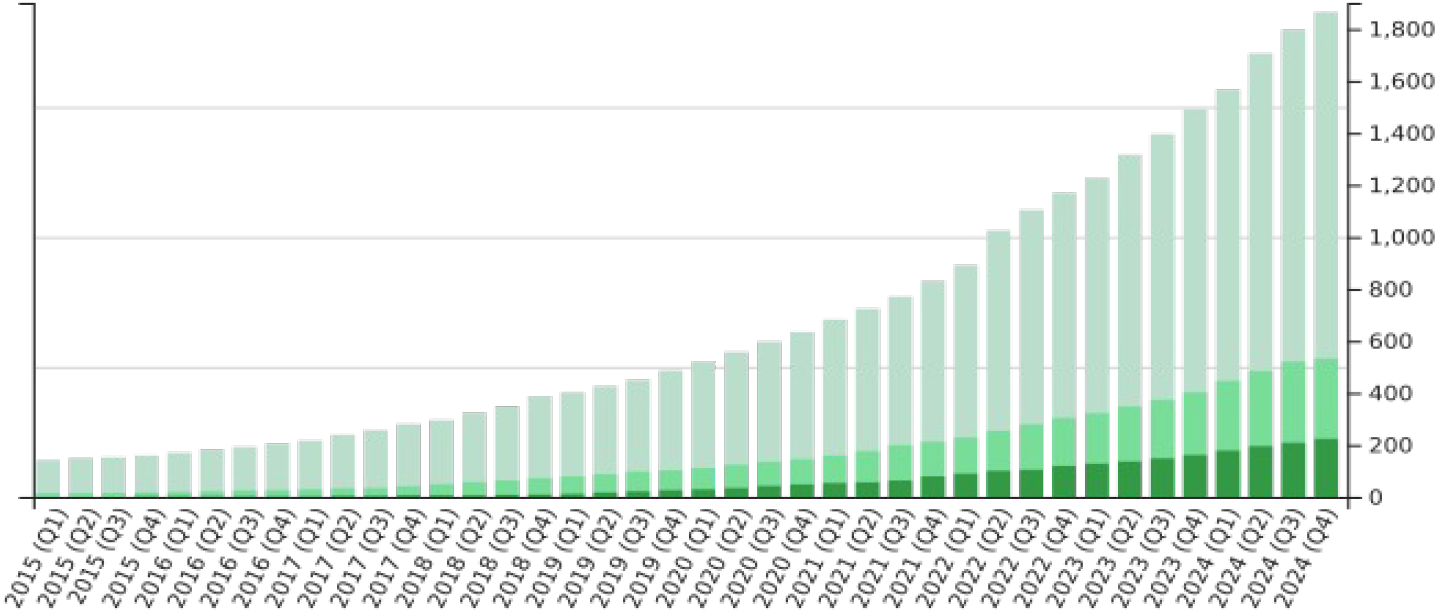
Number of plant species with sequenced and published genomes over time. The light green bars show the number of species that have been sequenced at least once, the medium green bars represent species that have been sequenced at least twice, and the dark green bars are those that have been sequenced three or more times. The bar chart shows quarterly data for the past decade.

The history of plant genome sequencing has been summarized at intervals to highlight the status of sequenced plant genomes at the time of publication, often focusing on particular technological advances (Michael and Jackson, 2013; Chen et al., 2019; Kersey, 2019; Shirasawa et al., 2021; Kress et al., 2022; Sun et al., 2022; Bernal-Gallardo and de Folter, 2024). But such is the pace of change that such review articles are often out of date by the time they are published. One attempt to present a more frequently updated resource is the Plants-Genomes-Technologies (N3) database (Xie et al., 2024). The current iteration is version 3.0 (accessed February 17, 2025), which was published on January 11, 2024. Here, we describe an additional online resource called PubPlant (https://www.plabipd.de/pubplant_main.html) that has tracked and continuously updated published plant genome sequences for almost a decade.

## Methods

### Tracking published plant genomes

PubPlant tracks published plant genomes by using the Google search engine, conducting a cited reference search in the Web of Science database (Clarivate, 2025) and the NCBI PubMed citation database (PubMed, 2025). To be included in PubPlant, the sequenced genome must belong to a plant (from the Archaeplastida group, which includes land plants, charophytes, green algae, glaucophytes, red algae and Rhodelphidophyta) and must be comprehensively described in a peer-reviewed journal. This means that the reads must be assembled into contigs, the contigs must have undergone scaffolding, and structural gene annotation must be completed. Accordingly, PubPlant excludes highly fragmented genomes or those with incomplete or missing structural annotations.

If a publication includes the genomes of multiple accessions representing a single species, such as an intra-species pan-genome, PubPlant counts the species as sequenced and published once. If a pan-genome is based on the genomes of different species – often described as a super pan-genome (Khan et al., 2020) – each species is counted individually. The accepted scientific names of vascular plant species are cross-checked against Plants of the World Online (POWO) 2025 (Govaerts et al., 2021). The lowest recognized taxonomic rank is the species. Accordingly, genomes of different subspecies or varieties are pooled. The phylogenetic positions of the plant species follow the classifications as listed in Table 1. PubPlant presents diagrams to illustrate the current status of sequenced plant genomes, and these are created using the Javascript libraries JQUERY (https://jquery.com), D3.JS (https://d3js.org) and VIS.JS (https://visjs.org).

**Table 1.** Resources used for phylogenetic classification in the diagrams presented by PubPlant.

| Plant group | Classification |
| --- | --- |
| Angiosperms | APG-IV system<br>(The Angiosperm Phylogeny Group, 2016;<br>Zuntini et al., 2024) |
| Gymnosperms | Yang et al. (2022) |
| Lycophytes and ferns | PPG-I system<br>(The Pteridophyte Phylogeny Group, 2016;<br>Hassler, 2025) |
| Bryophytes | Bechteler et al. (2023) |
| Primary algae | AlgaeBase (2025) |
| Others | NCBI Taxonomy (Schoch et al., 2020);<br>Catalogue of Life (2025) |

### Food crops

All statistical data concerning major food crop species were extracted from the Food and Agriculture Organization Corporate Statistical Database (FAOSTAT 2024). The data provided by FAOSTAT refer to crop categories with an annual production of more than 5000 tons in 2022. Each of these categories contains one or several food crops. The NCBI Taxonomy Database (Schoch et al., 2020) was used to assign correct scientific names to individual crops.

## Results and discussion

### Chronology of published plant genomes

PubPlant’s timeline view displays sequenced plant genomes according to the chronology of their first publication dates (Fig. 2, a full list of published plant genomes is presented in Supplementary Table 1). If there are two publications for the same plant species on the same date, the publication with the earliest acceptance date is listed. The timeline view groups the plant entries as belonging to dicotyledons, non-dicotyledons, non-angiosperms or algae (Fig. 2).

**Fig. 2.**
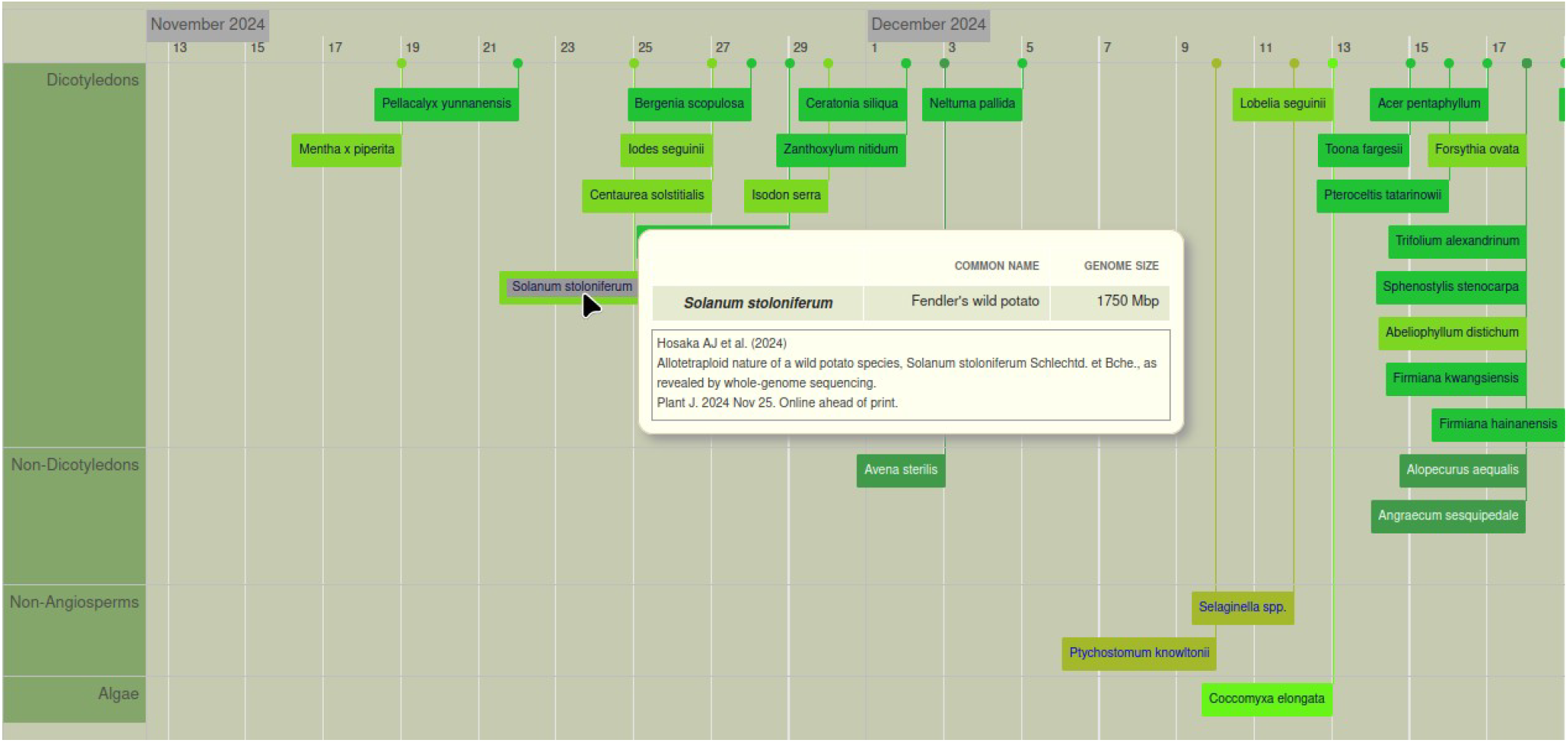
Timeline view of published plant genomes in PubPlant. The colored boxes refer to sequenced genomes of an individual species or a genus and multiple species (spp. for *species pluralis*) if the publication describes several species of the same genus. On the website, the mouse-over function displays the scientific name (or a list of scientific names if the box refers to several species of the same genus, as in the example shown for *Selaginella* spp.), the common plant name, the genome size, and the citation of the first publication. Clicking on the box links to the full-text publication in a new window.

### Phylogeny of sequenced plant genomes

PubPlant’s cladogram view arranges published plant genomes according to the phylogenetic position of each species (Fig. 3). The cladograms for flowering and non-flowering plants are displayed separately to improve legibility. Each entry in the cladogram provides a mouse-over function that displays a popup box containing the scientific name, common name and genome size of the plant. Publication details (one or more publications), including links to the corresponding full-text articles, are also provided in the popup box.

**Fig. 3.**
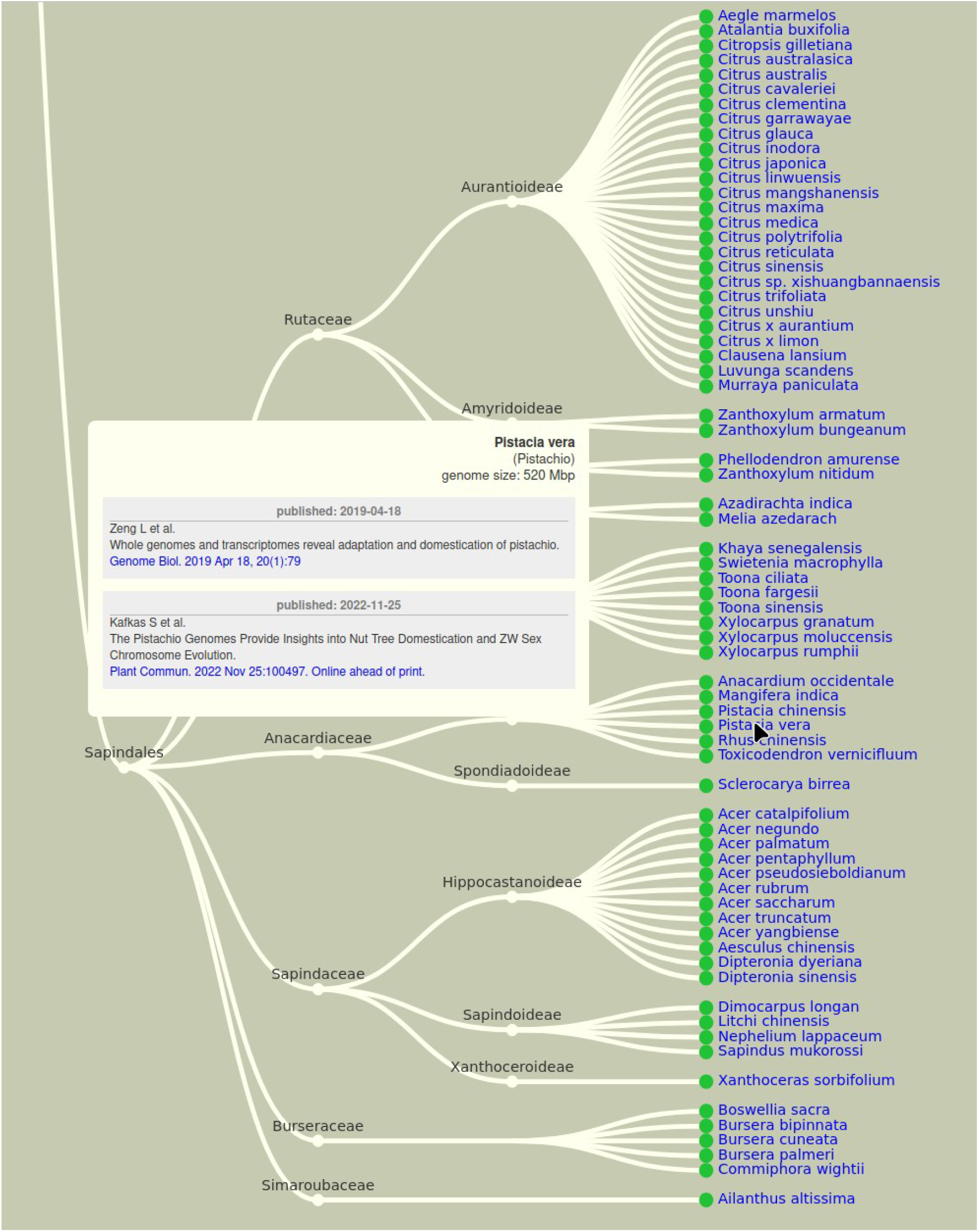
Cladogram view of plant species with sequenced genomes in PubPlant, showing the order Sapindales as an example. The cladogram view goes beyond the taxonomic rank of family, also showing subfamilies and species with sequenced genomes. On the website, the mouse-over function on the scientific name displays a popup box showing the genome size and listing publications (up to three for plants that have been sequenced multiple times) with links to the full-text article(s).

### Overview diagram of sequenced seed plants

The two major groups of seed plants are the angiosperms and gymnosperms. Angiosperms are by far the most diverse extant plant group, comprising more than 350,000 known species assigned to more than 400 families in 64 orders. The gymnosperms are a much smaller group, containing only 1100 living species, which is comparable to some of the larger angiosperm genera (e.g., *Solanum, Acacia* and *Rhododendron*) in terms of species numbers. By the end of 2024, 1700 angiosperm species and 26 gymnosperm species had been sequenced.

While the cladogram diagrams in PubPlant focus at the individual species level, the overview diagram provides a view at the taxonomic family level using an embedded progress bar. The cladogram displays the phylogenetic position of each plant family while the progress bar depicts the number of sequenced species in that family (Fig. 4). The overview diagram also includes families lacking any sequenced species thus far. This reveals that there are many plant families with no sequenced species (e.g., 25 of 38 families in the order Caryophyllales).

**Fig. 4.**
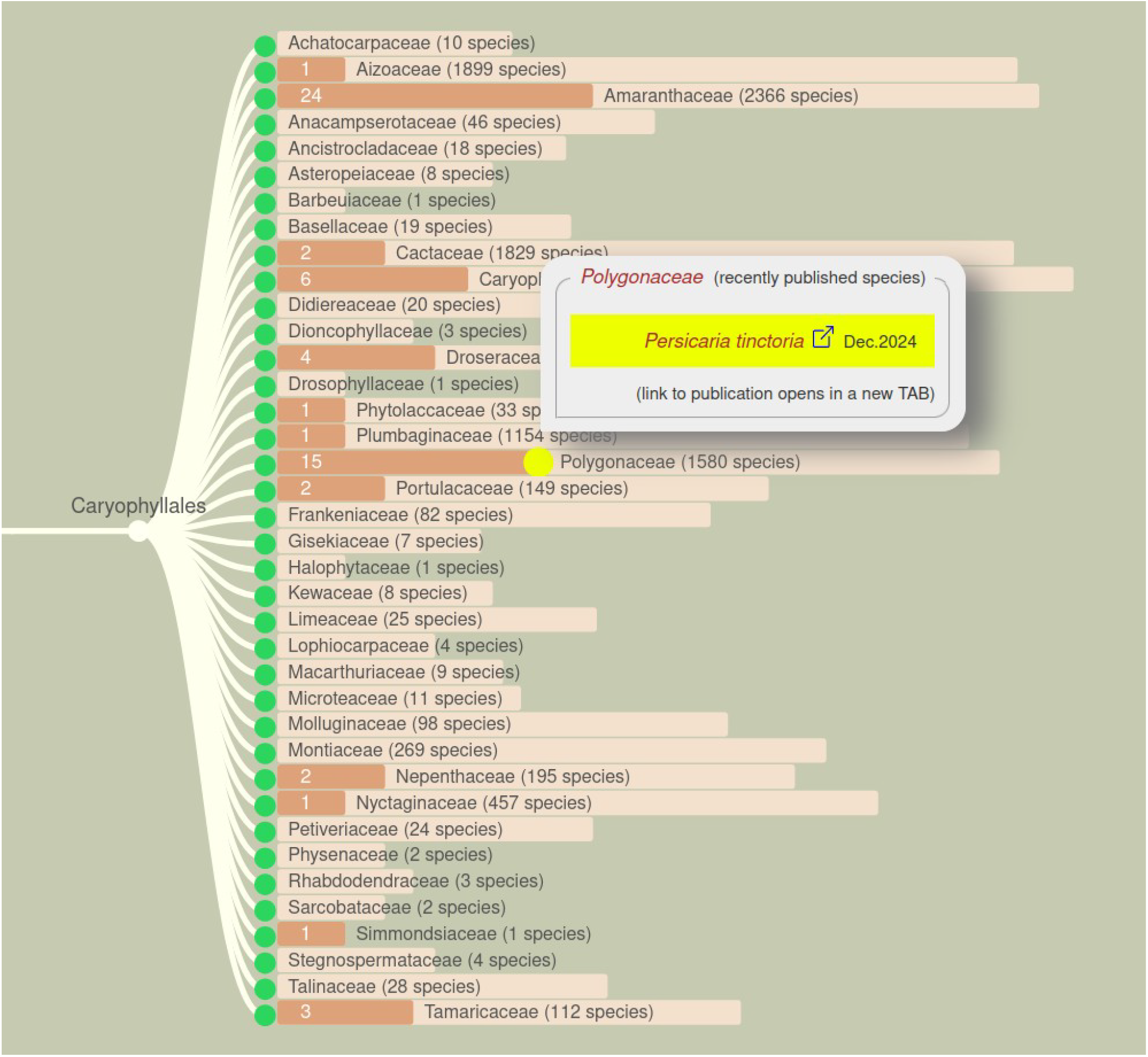
Overview diagram of sequenced angiosperms. The example shows the order Caryophyllales, which contains 38 families according to the APG-IV system. The family sizes are represented by light brown bars. The dark brown bars show how many of the species have been sequenced. All bars use a logarithmic scale. On the website, the mouse-over function on a dark bar shows the names of the genera and the number of species with sequenced genomes in a popup box. A yellow dot indicates recently published genomes. A mouse-over tooltip lists the individual species names and links to the full-text articles.

### Use case: sequenced food crops

As a use case, we evaluated the current status of sequenced food crops. Economically important food crop plants have always been a major target for sequencing efforts because this provides information about the genes responsible for agronomically favorable traits. The enormous genome sizes of some of these crops, mainly due to polyploidy, the presence of repetitive DNA (Jackson et al., 2011) and very long introns (Xu et al., 2024), hindered progress until the advent of long-read sequencing (Pellicer and Leitch, 2020). For example, the sizes of the onion and broad bean genomes are 16 Gbp (Hao et al., 2023) and 13 Gbp (Jayakodi et al., 2023), respectively. But thanks to the advances mentioned earlier, almost all of the food crop species listed in the FAOSTAT database (FAOSTAT 2024) have now been sequenced.

Further analysis of the most highly sequenced plant families (Poaceae, Fabaceae, Solanaceae, Rosaceae and Brassicaceae) indicated that these also contain the largest number of food crop species (Fig. 5, listed in Table 2). Other families such as Salicaceae, which do not contain any food crops, also may have sequenced species, but this reflects the numerous poplar and willow species in that family, which are important to the timber industry. Similarly, many species of Orchidaceae have been sequenced (Fig. 5) despite the presence of only one crop species (vanilla), due to the economic importance of many orchids as ornamental plants.

**Fig. 5.**
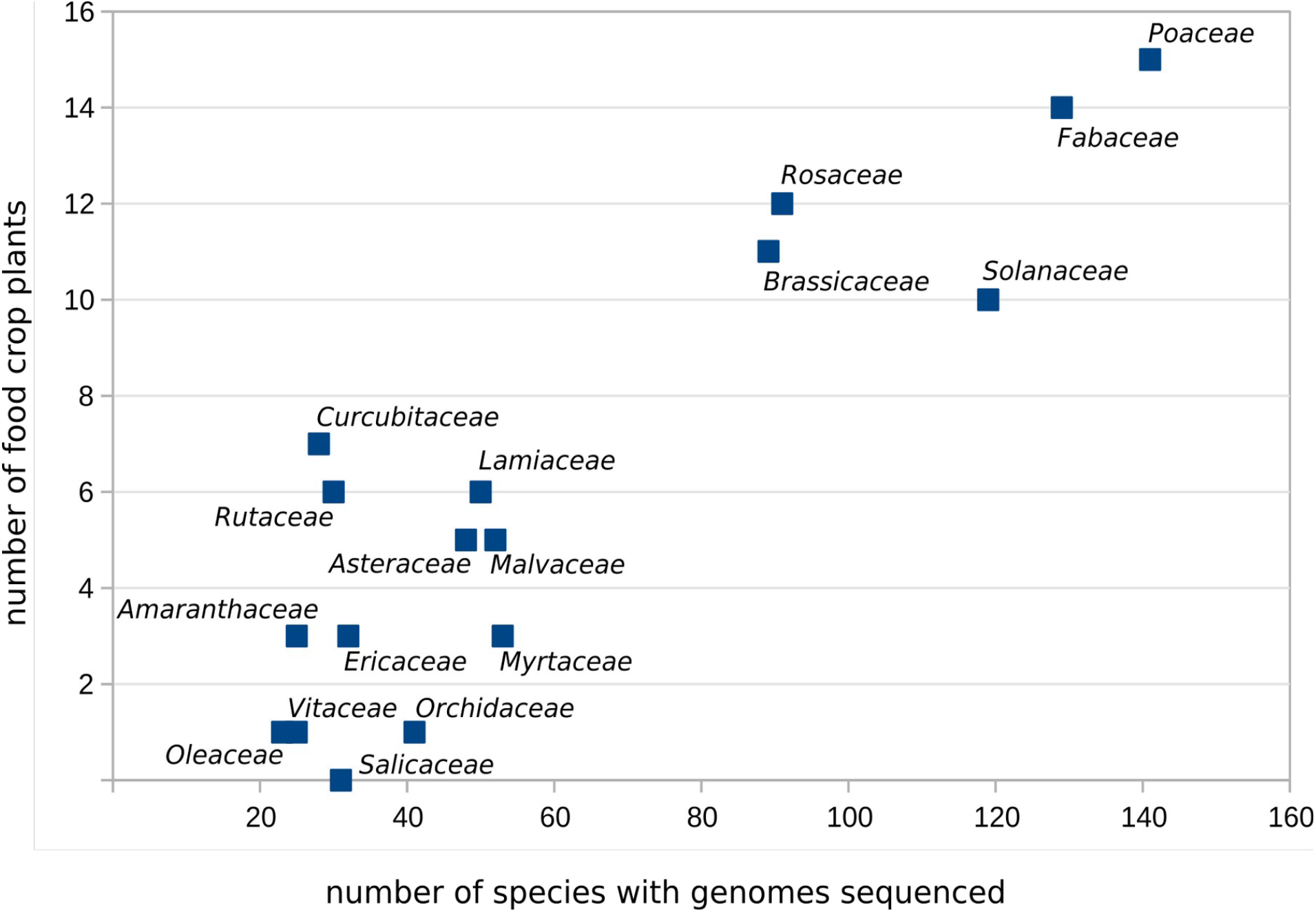
Scatter plot showing plant families in which more than 20 species have been sequenced. The *x*-axis indicates the number of species per family with sequenced genomes (evaluated February 2025) and the *y*-axis indicates the number of major food crops per family according to the FAOSTAT database (FAOSTAT 2024), including those that have yet to be sequenced.

**Table 2.**
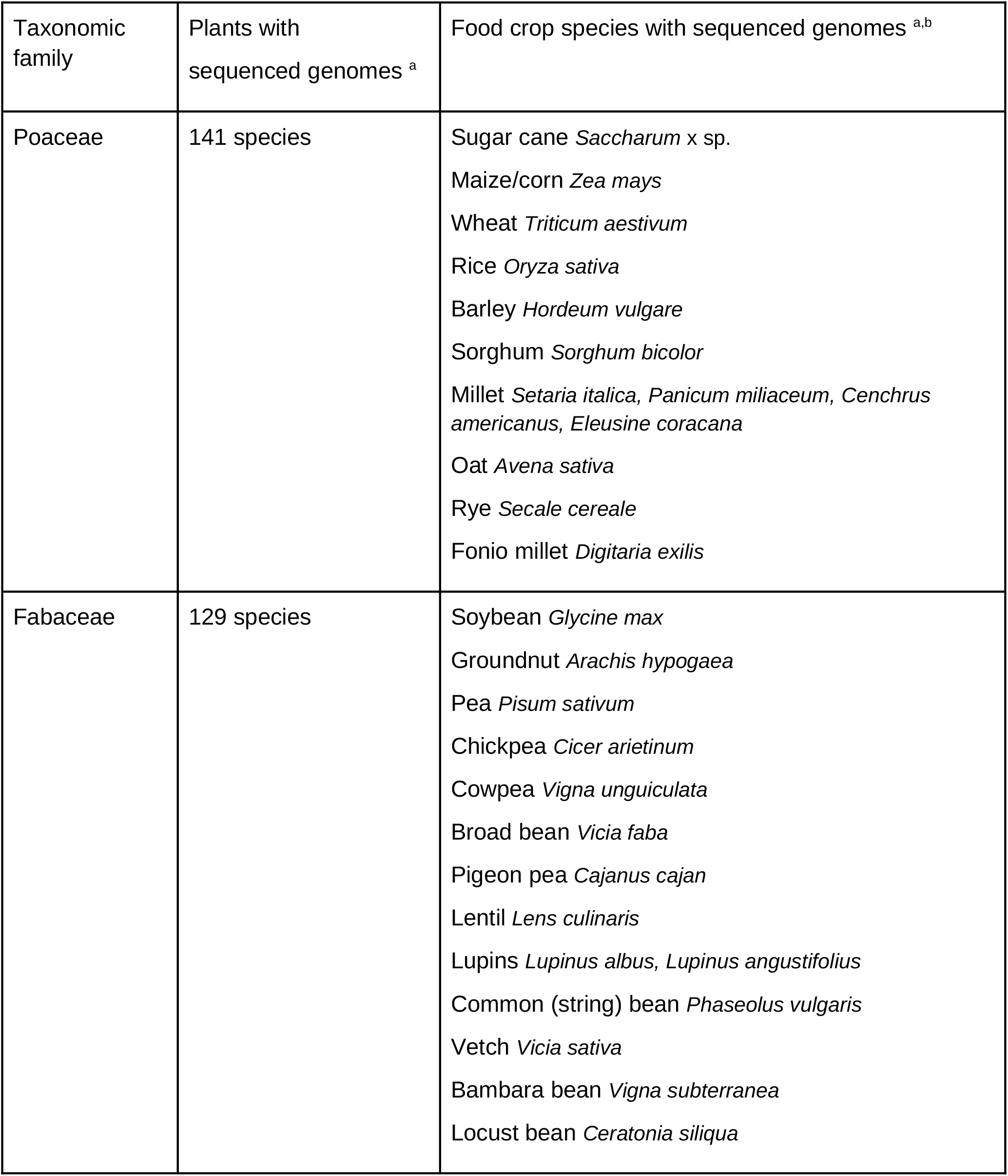
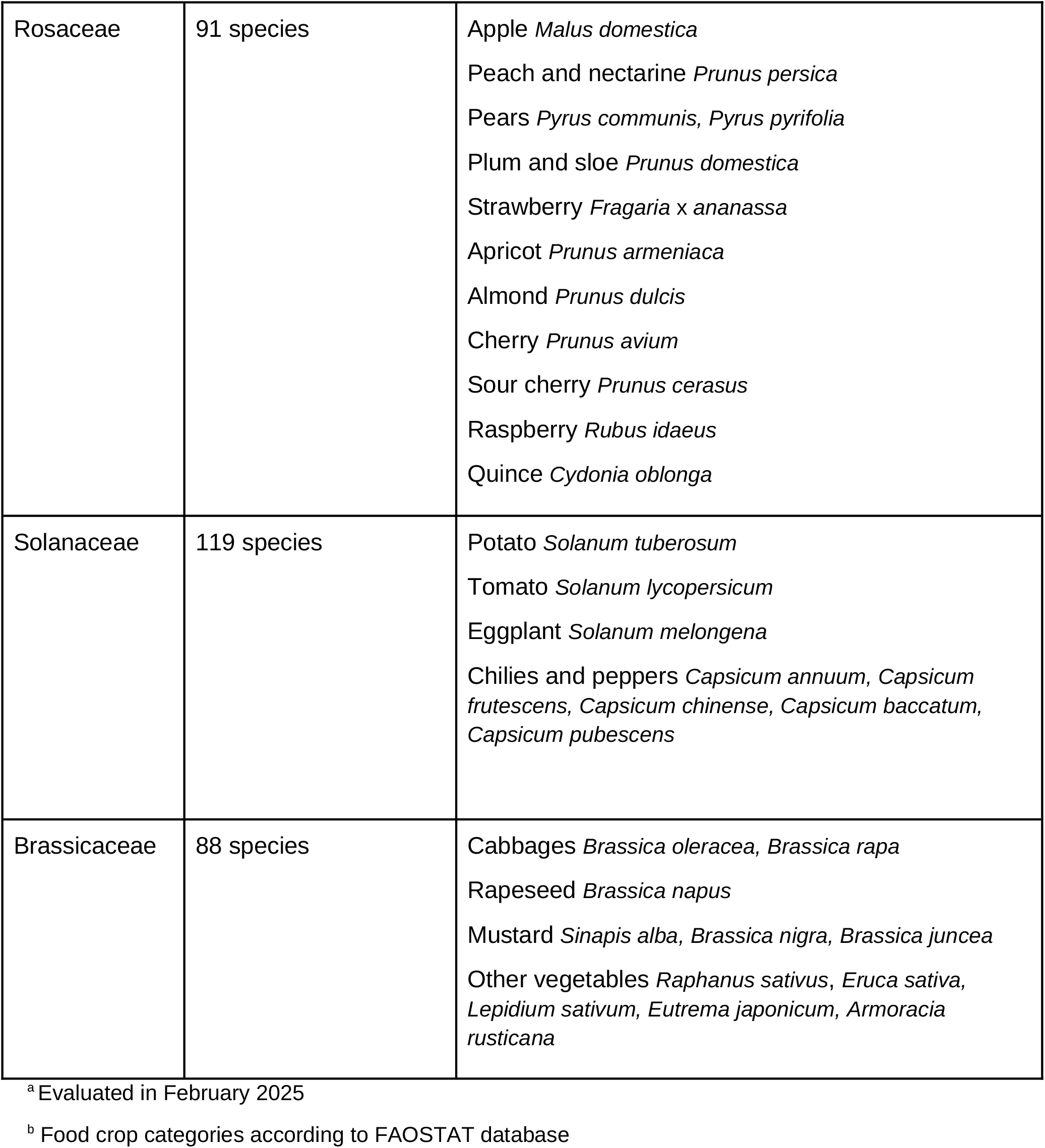
Plant families with the highest numbers of sequenced genomes.

| Taxonomic family | Plants with sequenced genomes <sup>a</sup> | Food crop species with sequenced genomes <sup>a,b</sup> |
| --- | --- | --- |
| Poaceae | 141 species | Sugar cane *Saccharum* x sp.<br>Maize/corn *Zea mays*<br>Wheat *Triticum aestivum*<br>Rice *Oryza sativa*<br>Barley *Hordeum vulgare*<br>Sorghum *Sorghum bicolor*<br>Millet *Setaria italica*, *Panicum miliaceum*, *Cenchrus americanus*, *Eleusine coracana*<br>Oat *Avena sativa*<br>Rye *Secale cereale*<br>Fonio millet *Digitaria exilis* |
| Fabaceae | 129 species | Soybean *Glycine max*<br>Groundnut *Arachis hypogaea*<br>Pea *Pisum sativum*<br>Chickpea *Cicer arietinum*<br>Cowpea *Vigna unguiculata*<br>Broad bean *Vicia faba*<br>Pigeon pea *Cajanus cajan*<br>Lentil *Lens culinaris*<br>Lupins *Lupinus albus*, *Lupinus angustifolius*<br>Common (string) bean *Phaseolus vulgaris*<br>Vetch *Vicia sativa*<br>Bambara bean *Vigna subterranea*<br>Locust bean *Ceratonia siliqua* |
| Rosaceae | 91 species | Apple *Malus domestica*<br>Peach and nectarine *Prunus persica*<br>Pears *Pyrus communis*, *Pyrus pyrifolia*<br>Plum and sloe *Prunus domestica*<br>Strawberry *Fragaria x ananassa*<br>Apricot *Prunus armeniaca*<br>Almond *Prunus dulcis*<br>Cherry *Prunus avium*<br>Sour cherry *Prunus cerasus*<br>Raspberry *Rubus idaeus*<br>Quince *Cydonia oblonga* |
| Solanaceae | 119 species | Potato *Solanum tuberosum*<br>Tomato *Solanum lycopersicum*<br>Eggplant *Solanum melongena*<br>Chilies and peppers *Capsicum annuum*, *Capsicum frutescens*, *Capsicum chinense*, *Capsicum baccatum*, *Capsicum pubescens* |
| Brassicaceae | 88 species | Cabbages *Brassica oleracea*, *Brassica rapa*<br>Rapeseed *Brassica napus*<br>Mustard *Sinapis alba*, *Brassica nigra*, *Brassica juncea*<br>Other vegetables *Raphanus sativus*, *Eruca sativa*, *Lepidium sativum*, *Eutrema japonicum*, *Armoracia rusticana* |

<sup>a</sup> Evaluated in February 2025
<sup>b</sup> Food crop categories according to FAOSTAT database

Only a few food crops remain to be sequenced (Table 3), including several culinary spices belonging to the Apiaceae (e.g. anise), leek (Amaryllidaceae), gooseberry and currants (Grossulariaceae). Notably, the blackcurrant (Grossulariaceae) is one of the most recently sequenced crop plants (Ziegler et al., 2024). Almost all of the food crops yet to be sequenced rank in the bottom half of the top 120 food crops ordered by production quantity (Table 3). Although all important food crop plants have now been sequenced, the remaining unsequenced species tend to be important in countries that contribute less to plant genome sequencing (Table 3). The countries that have contributed the most to the publication of crop plant genomes are China, the USA, Japan, Germany and Australia (Xie et al., 2024).

**Table 3.** Food crop plants that have not been sequenced, or have been sequenced very recently.

| Food crop category <sup>a</sup> | Ranking by production quantity <sup>b</sup> | Total production quantity (kiloton) | Main producers by production quantity (production share) <sup>c</sup> | Current status in genome sequencing <sup>d</sup> |
| --- | --- | --- | --- | --- |
| Anise, badian, coriander, cumin, caraway, fennel, juniper berries<br>*Pimpinella anisum*<br>*Illicium verum*<br>*Coriandrum sativum*<br>*Cuminum cyminum*<br>*Carum carvi*<br>*Foeniculum vulgare*<br>*Juniperus communis* | 76th | 2751 | India (68.6%)<br>Turkey (12.6%) | not sequenced (except *Coriandrum sativum*) |
| Leek<br>*Allium ampeloprasum* | 80th | 2109 | Indonesia (30.3%)<br>Turkey (8.0%) | not sequenced |
| Yerba Maté<br>*Ilex paraguariensis* | 82th | 1653 | Argentina (56.3%)<br>Brazil (37.4%) | sequenced (Vignale et al., 2025) |
| Currants<br>*Ribes nigrum*<br>*Ribes rubrum* | 97th | 764 | Russian Fed. (66.6%)<br>Poland (19.1%) | *Ribes nigrum* sequenced (Ziegler et al., 2024) |
| Yautía<br>*Xanthosoma sagittifolium* | 103th | 396 | Cuba (22.7%)<br>Venezuela (22.3%) | not sequenced |
| Kola nuts<br>*Cola acuminata*<br>*Cola nitida* | 104th | 315 | Nigeria (55.3%)<br>Côte d'Ivoire (18.6%) | not sequenced |
| Gooseberry<br>*Ribes uva-crispa* | 112th | 95 | Russian Fed. (89.0%)<br>Ukraine (8.0%) | not sequenced |
| Brazil nut<br>*Bertholletia excelsa* | 113th | 79 | Brazil (48.1%)<br>Bolivia (42.9%) | sequenced (Wang et al., 2025) |
| Locust bean<br>*Ceratonia siliqua* | 114th | 56 | Turkey (44.5%)<br>Morocco (39.1%) | sequenced (Akutsu et al., 2024) |
| Peppermint, spearmint<br>*Mentha x piperita*<br>*Mentha spicata* | 115th | 51 | Morocco (84.0%)<br>Argentina (13.7%) | *Mentha x piperita*<br>Sequenced (Talbot et al., 2024) |

- <sup>a</sup> Food crop categories according to the FAOSTAT database - <sup>b</sup> Ranking among the top 120 food crop categories (data for 2022) - <sup>c</sup> Top two countries with the largest share of world production (data for 2022) - <sup>d</sup> Evaluated in February 2025

## Conclusion

In recent years, the publication of newly sequenced plant genomes has increased to such an extent that it has become a weekly event (Fig. 1). Periodic review articles provide a snapshot of the situation at the time the manuscripts were written, but are quickly outdated. We therefore seek to highlight the online resource PubPlant, which provides up-to-date information on published plant genomes. PubPlant features a timeline view, where sequenced plant genomes are arranged in chronological order by the date of first publication (Fig. 2), and a cladogram view, showing all sequenced plant species arranged according to their phylogeny (Fig. 3). Separate cladograms are provided to display the phylogenetic positions of sequenced flowering and non-flowering plants. A summary diagram shows the phylogenetic position of all sequenced seed plants down to the taxonomic family rank, including those without any species sequenced thus far. It shows the total number of species for each family, the number of sequenced species, and the sequenced genera, while highlighting recently published genomes (Fig. 4).

As a use case for PubPlant, we evaluated the status of sequenced food crop plants, which tend to be prioritized for genome sequencing. Unsurprisingly, almost all major food crop species (FAOSTAT food crop categories ranked by production quantity, full list in Supplementary Table 2) have already been sequenced. In addition, the five plant families with the greatest number of sequenced plant genomes (Poaceae, Fabaceae, Rosaceae, Solanaceae and Brassicaceae) are also those containing the most major food crop species (Table 2).

PubPlant has been available online for more than 9 years and has been widely used, including as a resource to prepare review articles summarizing plant genome sequencing progress (Jiao and Schneeberger, 2017; Hao et al. 2022; Bernal-Gallardo and de Folter, 2024) and to evaluate the status of sequenced medicinal plants (Cheng et al., 2021b). Whereas other plant genome resources release irregular updates, PubPlant is updated on a monthly basis, providing a simple and intuitive source of current and historical information on sequenced plant genomes for the benefit of the entire plant research community.

## Supporting information

List of published sequenced plant genomes in chronological order

List of food crop categories (FAOSTAT) ranked by production quantity

## Supplementary material

Supplementary Table 1 | List of published sequenced plant genomes in chronological order.

Supplementary Table 2 | List of food crop categories (FAOSTAT) ranked by production quantity.

