## Supplementary material for "PubPlant – a continuously updated online resource for sequenced and published plant genomes": List of published sequenced plant genomes in chronological order

Plants with genomes sequenced and published ordered by the date of their first publication.

updated: Feb 11 2025, extracted from https://www.plabipd.de/pubplant_main.html

| | plant species | published | |
|---|---|---|---|
| 1924 | *Bracteacoccus bullatus* *(Bracteacoccaceae)* | 2025-02-10 | Liang H et al., Chromosome-level genomes of two Bracteacoccaceae highlight adaptations to biocrusts. Nat Commun. 2025 Feb 10, 16(1): 1492. |
| 1923 | *Bracteacoccus minor* *(Bracteacoccaceae)* | 2025-02-10 | Liang H et al., Chromosome-level genomes of two Bracteacoccaceae highlight adaptations to biocrusts. Nat Commun. 2025 Feb 10, 16(1): 1492. |
| 1922 | *Peganum multisectum* *(Nitrariaceae)* | 2025-02-10 | Xu H et al., Integration of De Novo Chromosome-Level Genome and Population Resequencing of Peganum (Nitrariaceae): A Case Study of Speciation and Evolutionary Trajectories in Arid Central Asia. Mol Ecol Resour. 2025 Feb 10: e14078. Online ahead of print. |
| 1921 | *Amorphophallus albus* *(Araceae)* | 2025-02-07 | Duan L et al., Genomic, transcriptomic and metabolomic analyses of Amorphophallus albus provides insights into the evolution and resistance to southern blight pathogen. Front Plant Sci. 2025 Feb 7: 15: 1518058. eCollection 2025. |
| 1920 | *Caragana korshinskii* *(Fabaceae)* | 2025-02-06 | Mei F et al., Genomic insights into drought adaptation of the forage shrub Caragana korshinskii (Fabaceae) widely planted in drylands. Plant J. 2025 Feb, 121(3): e17255. |
| 1919 | *Silene dioica* *(Caryophyllaceae)* | 2025-02-06 | Akagi T et al., Rapid and dynamic evolution of a giant Y chromosome in Silene latifolia. Science. 2025 Feb 7, 387(6734): 637-643. Epub 2025 Feb 6. |
| 1918 | *Silene vulgaris* *(Caryophyllaceae)* | 2025-02-06 | Akagi T et al., Rapid and dynamic evolution of a giant Y chromosome in Silene latifolia. Science. 2025 Feb 7, 387(6734): 637-643. Epub 2025 Feb 6. |
| 1917 | *Prunus speciosa* *(Rosaceae)* | 2025-02-04 | Fujiwara K et al., A Near Complete Genome Assembly of the Oshima Cherry Cerasus speciosa. Sci Data. 2025 Feb 4, 12(1): 162. |
| 1916 | *Micractinium tetrahymenae* *(Chlorellaceae)* | 2025-02-03 | Kelly JB et al., Genomic signatures of adaptation to stress reveal shared evolutionary trends between Tetrahymena utriculariae and its algal endosymbiont, Micractinium tetrahymenae. Mol Biol Evol. 2025 Feb 3: msaf030. Online ahead of print. |
| 1915 | *Capsella orientalis* *(Brassicaceae)* | 2025-01-31 | Kasianova AM et al., Chromosome-Scale Assembly of Capsella orientalis, Maternal Progenitor of Cosmopolitan Allotetraploid C. bursa-pastoris. Genome Biol Evol. 2025 Jan 6, 17(1): evaf009. |
| 1914 | *Pontederia cordata* *(Pontederiaceae)* | 2025-01-29 | Wang J et al., Chromosome-level genome assembly of Pontederia cordata L. provides insights into its rapid adaptation and variation of flower colors. DNA Res. 2025 Jan 29: dsaf002. Online ahead of print. |
| 1913 | *Capsicum chacoense* *(Solanaceae)* | 2025-01-28 | Zhang K et al., Transposon proliferation drives genome architecture and regulatory evolution in wild and domesticated peppers. Nat Plants. 2025 Jan 28. Online ahead of print. |
| 1912 | *Capsicum eximium* *(Solanaceae)* | 2025-01-28 | Zhang K et al., Transposon proliferation drives genome architecture and regulatory evolution in wild and domesticated peppers. Nat Plants. 2025 Jan 28. Online ahead of print. |
| 1911 | *Salsola tragus* *(Amaranthaceae)* | 2025-01-25 | Lemas JM et al., Assembly and Annotation of the Tetraploid Salsola tragus (Russian thistle) Genome. Genome Biol Evol. 2025 Jan 25: evaf014. Online ahead of print. |
| 1910 | *Stylosanthes angustifolia* *(Fabaceae)* | 2025-01-24 | Liu C et al., A chromosome-scale genome assembly of the pioneer plant Stylosanthes angustifolia: insights into genome evolution and drought adaptation. Gigascience. 2025 Jan 6: 14: giae118. |
| 1909 | *Urochloa eminii* *(Poaceae)* | 2025-01-24 | Ryan C et al., A haplotype-resolved chromosome-level genome assembly of Urochloa decumbens cv. Basilisk resolves its allopolyploid ancestry and composition. G3 (Bethesda). 2025 Jan 24:jkaf005. Online ahead of print. |
| 1908 | *Bertholletia excelsa* *(Lecythidaceae)* | 2025-01-22 | Wang H et al., Long-term human influence on the demography and genetic diversity of the hyperdominant Bertholletia excelsa in the Amazon Basin. Curr Biol. 2025 Jan 22:S0960-9822(24)01691-9. Online ahead of print. |
| 1907 | *Rosa laevigata* *(Rosaceae)* | 2025-01-22 | Wang Y et al., A high-quality chromosome-scale genome assembly of the Cherokee rose (Rosa laevigata). Sci Data. 12, 132 (2025). |
| 1906 | *Bergenia purpurascens* *(Saxifragaceae)* | 2025-01-21 | Liu XD et al., A Chromosome-level genome assembly of the alpine medicinal plant Bergenia purpurascens (Saxifragaceae). Sci Data. 12, 121 (2025). |
| 1905 | *Cinchona calisaya* *(Rubiaceae)* | 2025-01-21 | Du C et al., A near-complete genome assembly of Cinchona calisaya. Sci Data. 2025 Jan 21, 12(1): 122. |
| 1904 | *Cyperus iria* *(Cyperaceae)* | 2025-01-21 | Yang S et al., Chromosome-level genome assembly of Cyperus iria, an aggressive weed of rice. Sci Data. 2025 Jan 21, 12(1): 120. |
| 1903 | *Tylosema esculentum* *(Fabaceae)* | 2025-01-21 | Li J et al., Genome assembly and population analysis of tetraploid marama bean reveal two distinct genome types. Sci Rep. 2025 Jan 21, 15(1): 2665. |
| 1902 | *Calochortus tolmiei* *(Liliaceae)* | 2025-01-20 | Landis JB et al., Reference genome of Calochortus tolmiei Hook. & Arn. (Liliaceae), a cat's ear mariposa lily. G3 (Bethesda). 2025 Jan 20: jkaf008. Online ahead of print. |
| 1901 | *Bougainvillea glabra* *(Nyctaginaceae)* | 2025-01-18 | Lan L et al., Chromosome-level and haplotype-resolved genome assembly of Bougainvillea glabra. Sci Data. 12, 107 (2025). |
| 1900 | *Dovyalis afra* *(Salicaceae)* | 2025-01-15 | Kariba R et al., Draft genome sequence of Kei apple, an underutilized African tree crop Sci Data. 2025 Jan 15, 12(1): 70. |
| 1899 | *Riccia sorocarpa* *(Ricciaceae)* | 2025-01-15 | Krawczyk K et al., Chromosome-scale telomere to telomere genome assembly of common crystalwort (Riccia sorocarpa Bisch.). Sci Data. 12, 77 (2025). |
| 1898 | *Lagerstroemia excelsa* *(Lythraceae)* | 2025-01-11 | Qiao Z et al., Genome assembly and multiomic analyses reveal insights into flower and bark colors of Lagerstroemia excelsa. Plant Physiol Biochem. 2025 Jan 11, Online ahead of print. |
| 1897 | *Dendrobium aphyllum* *(Orchidaceae)* | 2025-01-10 | Li Y et al., Pangeneric genome analyses reveal the evolution and diversity of the orchid genus Dendrobium. Nat Plants. 2025 Jan 10. Online ahead of print. |
| 1896 | *Dendrobium crocatum* *(Orchidaceae)* | 2025-01-10 | Li Y et al., Pangeneric genome analyses reveal the evolution and diversity of the orchid genus Dendrobium. Nat Plants. 2025 Jan 10. Online ahead of print. |
| 1895 | *Dendrobium crumenatum* *(Orchidaceae)* | 2025-01-10 | Li Y et al., Pangeneric genome analyses reveal the evolution and diversity of the orchid genus Dendrobium. Nat Plants. 2025 Jan 10. Online ahead of print. |
| 1894 | *Dendrobium discolor* *(Orchidaceae)* | 2025-01-10 | Li Y et al., Pangeneric genome analyses reveal the evolution and diversity of the orchid genus Dendrobium. Nat Plants. 2025 Jan 10. Online ahead of print. |
| 1893 | *Dendrobium ellipsophyllum* *(Orchidaceae)* | 2025-01-10 | Li Y et al., Pangeneric genome analyses reveal the evolution and diversity of the orchid genus Dendrobium. Nat Plants. 2025 Jan 10. Online ahead of print. |
| 1892 | *Dendrobium formosum* *(Orchidaceae)* | 2025-01-10 | Li Y et al., Pangeneric genome analyses reveal the evolution and diversity of the orchid genus Dendrobium. Nat Plants. 2025 Jan 10. Online ahead of print. |
| 1891 | *Dendrobium hercoglossum* *(Orchidaceae)* | 2025-01-10 | Li Y et al., Pangeneric genome analyses reveal the evolution and diversity of the orchid genus Dendrobium. Nat Plants. 2025 Jan 10. Online ahead of print. |

| | plant species | published | |
|---|---|---|---|
| 1890 | *Dendrobium hybrid cv. CPS* *(Orchidaceae)* | 2025-01-10 | Li Y et al., Pangeneric genome analyses reveal the evolution and diversity of the orchid genus Dendrobium. Nat Plants. 2025 Jan 10. Online ahead of print. |
| 1889 | *Dendrobium jenkinsii* *(Orchidaceae)* | 2025-01-10 | Li Y et al., Pangeneric genome analyses reveal the evolution and diversity of the orchid genus Dendrobium. Nat Plants. 2025 Jan 10. Online ahead of print. |
| 1888 | *Dendrobium leonis* *(Orchidaceae)* | 2025-01-10 | Li Y et al., Pangeneric genome analyses reveal the evolution and diversity of the orchid genus Dendrobium. Nat Plants. 2025 Jan 10. Online ahead of print. |
| 1887 | *Dendrobium lindleyi* *(Orchidaceae)* | 2025-01-10 | Li Y et al., Pangeneric genome analyses reveal the evolution and diversity of the orchid genus Dendrobium. Nat Plants. 2025 Jan 10. Online ahead of print. |
| 1886 | *Dendrobium secundum* *(Orchidaceae)* | 2025-01-10 | Li Y et al., Pangeneric genome analyses reveal the evolution and diversity of the orchid genus Dendrobium. Nat Plants. 2025 Jan 10. Online ahead of print. |
| 1885 | *Dendrobium smillieae* *(Orchidaceae)* | 2025-01-10 | Li Y et al., Pangeneric genome analyses reveal the evolution and diversity of the orchid genus Dendrobium. Nat Plants. 2025 Jan 10. Online ahead of print. |
| 1884 | *Dendrobium tetragonum* *(Orchidaceae)* | 2025-01-10 | Li Y et al., Pangeneric genome analyses reveal the evolution and diversity of the orchid genus Dendrobium. Nat Plants. 2025 Jan 10. Online ahead of print. |
| 1883 | *Diospyros sp. deyangensis* *(Ebenaceae)* | 2025-01-08 | Guan C et al., The haplotype-resolved and chromosome-level reference genome assembly of Diospyros deyangensis provides insights into the evolution and juvenile growth of persimmon. Hortic Res. 2025 Jan: uhaf001. eCollection 2025 Jan. |
| 1882 | *Ilex paraguariensis* *(Aquifoliaceae)* | 2025-01-08 | Federico A Vignale FA et al., Yerba mate (Ilex paraguariensis) genome provides new insights into convergent evolution of caffeine biosynthesis. Elife. 2025 Jan 8: 14: e104759. |
| 1881 | *Citrus sp. xishuangbannaensis* *(Rutaceae)* | 2025-01-06 | Wang F et al., Haplotype-resolved genome of a papeda provides insights into the geographical origin and evolution of Citrus. J Integr Plant Biol. 2025 Jan 6. Online ahead of print. |
| 1880 | *Salvia sclarea* *(Lamiaceae)* | 2025-01-04 | Choi S et al., Chromosome-level genome assembly of Salvia sclarea. Sci Data. 2025 Jan 4, 12(1): 14. |
| 1879 | *Anthoceros fusiformis* *(Anthocerotaceae)* | 2025-01-03 | Schafran P et al., Pan-phylum genomes of hornworts reveal conserved autosomes but dynamic accessory and sex chromosomes. Nat Plants. 2025 Jan 3. Online ahead of print. |
| 1878 | *Leiosporoceros dussii* *(Leiosporocerotaceae)* | 2025-01-03 | Schafran P et al., Pan-phylum genomes of hornworts reveal conserved autosomes but dynamic accessory and sex chromosomes. Nat Plants. 2025 Jan 3. Online ahead of print. |
| 1877 | *Megaceros flagellaris* *(Dendrocerotaceae)* | 2025-01-03 | Schafran P et al., Pan-phylum genomes of hornworts reveal conserved autosomes but dynamic accessory and sex chromosomes. Nat Plants. 2025 Jan 3. Online ahead of print. |
| 1876 | *Notothylas orbicularis* *(Notothyladaceae)* | 2025-01-03 | Schafran P et al., Pan-phylum genomes of hornworts reveal conserved autosomes but dynamic accessory and sex chromosomes. Nat Plants. 2025 Jan 3. Online ahead of print. |
| 1875 | *Paraphymatoceros pearsonii* *(Notothyladaceae)* | 2025-01-03 | Schafran P et al., Pan-phylum genomes of hornworts reveal conserved autosomes but dynamic accessory and sex chromosomes. Nat Plants. 2025 Jan 3. Online ahead of print. |
| 1874 | *Phaeoceros carolinianus* *(Notothyladaceae)* | 2025-01-03 | Schafran P et al., Pan-phylum genomes of hornworts reveal conserved autosomes but dynamic accessory and sex chromosomes. Nat Plants. 2025 Jan 3. Online ahead of print. |
| 1873 | *Phaeoceros sp.* *(Notothyladaceae)* | 2025-01-03 | Schafran P et al., Pan-phylum genomes of hornworts reveal conserved autosomes but dynamic accessory and sex chromosomes. Nat Plants. 2025 Jan 3. Online ahead of print. |
| 1872 | *Phaeomegaceros chiloensis* *(Dendrocerotaceae)* | 2025-01-03 | Schafran P et al., Pan-phylum genomes of hornworts reveal conserved autosomes but dynamic accessory and sex chromosomes. Nat Plants. 2025 Jan 3. Online ahead of print. |
| 1871 | *Phymatoceros phymatodes* *(Phymatocerotaceae)* | 2025-01-03 | Schafran P et al., Pan-phylum genomes of hornworts reveal conserved autosomes but dynamic accessory and sex chromosomes. Nat Plants. 2025 Jan 3. Online ahead of print. |
| 1870 | *Urtica dioica* *(Urticaceae)* | 2025-01-03 | Hirabayashi K et al., A High-Quality Phased Genome Assembly of Stinging Nettle (Urtica dioica ssp. dioica). Plants. 2025, 14(1), 124 |
| 1869 | *Gloriosa superba* *(Colchicaceae)* | 2025-01-02 | Liang Y et al., The giant genome of lily provides insights into the hybridization of cultivated lilies. Nat Commun. 2025 Jan 2, 16(1): 45. |
| 1868 | *Lilium sargentiae* *(Liliaceae)* | 2025-01-02 | Liang Y et al., The giant genome of lily provides insights into the hybridization of cultivated lilies. Nat Commun. 2025 Jan 2, 16(1): 45. |
| 1867 | *Aegilops comosa* *(Poaceae)* | 2024-12-31 | Li H et al., Chromosome-scale assembly and annotation of the wild wheat relative Aegilops comosa. Sci Data. 2024 Dec 31, 11(1): 1454. |
| 1866 | *Palmaria palmata* *(Palmariaceae)* | 2024-12-30 | Rosignoli S et al., Draft genome of Palmaria palmata and intraspecific genetic variations in the North-East Atlantic. Algal Res. 2024 Dec:103887 |
| 1865 | *Isodon lophanthoides* *(Lamiaceae)* | 2024-12-28 | Liu J et al., Genome assembly and multi-omics analyses of Isodon lophanthoides provide insights into the distribution of medicinal metabolites induced by exogenous methyl jasmonate. BMC Plant Biol. 2024 Dec 28, 24(1): 1270. |
| 1864 | *Persicaria tinctoria* *(Polygonaceae)* | 2024-12-27 | Li Q et al., Chromosome-level genome assembly of the tetraploid medicinal and natural dye plant Persicaria tinctoria. Sci Data. 2024 Dec 27, 11(1): 1440. |
| 1863 | *Lonicera caerulea* *(Caprifoliaceae)* | 2024-12-26 | Wang J et al., Lonicera caerulea genome reveals molecular mechanisms of freezing tolerance and anthocyanin biosynthesis. J Adv Res. 2024 Dec 26: S2090-1232(24)00615-5. Online ahead of print. |
| 1862 | *Cardamine chenopodiifolia* *(Brassicaceae)* | 2024-12-23 | Emonet A et al., Polyploid genome assembly of Cardamine chenopodiifolia. Gigabyte 2024: 145 |
| 1861 | *Pandanus amaryllifolius* *(Pandanaceae)* | 2024-12-23 | Sidek N et al., Whole Genome Assembly, Annotation, and Comparative Genomic Analysis of Pandanus amaryllifolius. Plant Mol Biol Rep. 2024 Dec |
| 1860 | *Firmiana hainanensis* *(Malvaceae)* | 2024-12-19 | Dong Z et al., High-quality genome of Firmiana hainanensis provides insights into the evolution of Malvaceae subfamilies and the mechanism of their wood density formation. J Genet Genomics. 2024 Dec 19: S1673-8527(24)00362-X. Online ahead of print. |
| 1859 | *Abeliophyllum distichum* *(Oleaceae)* | 2024-12-18 | Jang H et al., Chromosome-level assemblies of the endemic Korean species Abeliophyllum distichum and Forsythia ovata. Sci Data. 2024 Dec 18, 11(1): 1372. |
| 1858 | *Alopecurus aequalis* *(Poaceae)* | 2024-12-18 | Wright J et al., Chromosome-scale genome assembly and de novo annotation of Alopecurus aequalis. Sci Data. 2024 Dec 18, 11(1): 1368. |
| 1857 | *Angraecum sesquipedale* *(Orchidaceae)* | 2024-12-18 | Jiang K et al., Genomic, transcriptomic, and metabolomic analyses reveal convergent evolution of oxime biosynthesis in Darwins orchid. Mol Plant. 2024 Dec 18, Online ahead of print. |
| 1856 | *Firmiana kwangsiensis* *(Malvaceae)* | 2024-12-18 | Wang B et al., A nearly telomere-to-telomere diploid genome assembly of Firmiana kwangsiensis, a threatened species in China. Sci Data. 2024 Dec 18, 11(1): 1394. |
| 1855 | *Forsythia ovata* *(Oleaceae)* | 2024-12-18 | Jang H et al., Chromosome-level assemblies of the endemic Korean species Abeliophyllum distichum and Forsythia ovata. Sci Data. 2024 Dec 18, 11(1): 1372. |
| 1854 | *Sphenostylis stenocarpa* *(Fabaceae)* | 2024-12-18 | Waweru B et al., Chromosome-scale assembly of the African yam bean genome. Sci Data. 2024 Dec 18, 11(1): 1384. |

| | plant species | published | |
|---|---|---|---|
| 1853 | *Trifolium alexandrinum* (*Fabaceae*) | 2024-12-18 | Sato MP et al., Near-complete telomere-to-telomere de novo genome assembly in Egyptian clover (Trifolium alexandrinum).<br>DNA Res. 2024 Dec 18: dsae036. Online ahead of print. |
| 1852 | *Acer pentaphyllum* (*Sapindaceae*) | 2024-12-17 | Li X et al., High-Resolution Genome Assembly and Population Genetic Study of the Endangered Maple Acer pentaphyllum (Sapindaceae): Implications for Conservation Strategies.<br>Hortic Res. 2024 Dec: uhae357. eCollection 2024 Dec. |
| 1851 | *Pteroceltis tatarinowii* (*Cannabaceae*) | 2024-12-16 | Qiao Q et al., Chromosome-level genome assembly of Pteroceltis tatarinowii provides new insights into evolution and fiber biosynthesis.<br>Ind Crops Prod. 2025 Feb, 224: 120316. Epub 2024 Dec 16. |
| 1850 | *Toona fargesii* (*Meliaceae*) | 2024-12-15 | Ma J et al., Chromosome-scale genomes of Toona fargesii provide insights into competency of root sprouting.<br>J Syst Evol. 2024, Online ahead of print. |
| 1849 | *Coccomyxa elongata* (*Coccomyxaceae*) | 2024-12-13 | Kraege A et al., High quality genome assembly and annotation (v1) of the eukaryotic freshwater microalga Coccomyxa elongata SAG 216-3b.<br>G3 (Bethesda). 2024 Dec 13: jkae294. Online ahead of print. |
| 1848 | *Lobelia seguinii* (*Campanulaceae*) | 2024-12-13 | Zhang N et al., A chromosome-level genome of Lobelia seguinii provides insights into the evolution of Campanulaceae and the lobeline biosynthesis.<br>Genomics. 2024 Dec 13: 110979. Online ahead of print. |
| 1847 | *Selaginella sellowii* (*Selaginellaceae*) | 2024-12-12 | Alejo-Jacuinde G et al., Gene family rearrangements and transcriptional priming drive the evolution of vegetative desiccation tolerance in Selaginella.<br>Plant J. 2024 Dec 12. Online ahead of print. |
| 1846 | *Selaginella silvestris* (*Selaginellaceae*) | 2024-12-12 | Alejo-Jacuinde G et al., Gene family rearrangements and transcriptional priming drive the evolution of vegetative desiccation tolerance in Selaginella.<br>Plant J. 2024 Dec 12. Online ahead of print. |
| 1845 | *Ptychostomum knowltonii* (*Bryaceae*) | 2024-12-10 | Ma C et al., Chromosome-level Genome Assembly and Annotation of the Arctic Moss Ptychostomum knowltonii.<br>Genome Biol Evol. 2024 Dec 10: evae268. Online ahead of print. |
| 1844 | *Neltuma pallida* (*Fabaceae*) | 2024-12-05 | La Torre Ramirez RD et al., A chromosome-level genome assembly of the Peruvian Algarrobo (Neltuma pallida) provides insights on its adaptation to its unique ecological niche.<br>G3 (Bethesda). 2024 Dec 05: jkae283. Online ahead of print. |
| 1843 | *Avena sterilis* (*Poaceae*) | 2024-12-03 | He Q et al., The near-complete genome assembly of hexaploid wild oat reveals its genome evolution and divergence with cultivated oats.<br>Nat Plants. 2024 Dec 3. Online ahead of print. |
| 1842 | *Berberis vulgaris* (*Berberidaceae*) | 2024-12-03 | Ruhsam M et al., The genome sequence of Berberis vulgaris L.<br>Wellcome Open Res. 2024 Dec 3: 9: 710. eCollection 2024. |
| 1841 | *Ceratonia siliqua* (*Fabaceae*) | 2024-12-02 | Akutsu M et al., De novo sequencing allows genome-wide identification of genes involved in galactomannan synthesis in locust bean (Ceratonia siliqua).<br>DNA Res. 2024 Dec 2: dsae033. |
| 1840 | *Zanthoxylum nitidum* (*Rutaceae*) | 2024-12-02 | Zhu Y et al., A high-quality chromosome-level genome assembly of the traditional Chinese medicinal herb Zanthoxylum nitidum.<br>Sci Data. 2024 Dec 2, 11(1): 1311. |
| 1839 | *Isodon serra* (*Lamiaceae*) | 2024-11-30 | Hou L et al., The Isodon serra genome sheds light on tanshinone biosynthesis and reveals the recursive karyotype evolutionary histories within Lamiales.<br>Plant J. 2024 Nov 30. Online ahead of print. |
| 1838 | *Phellodendron amurense* (*Rutaceae*) | 2024-11-29 | Xu Z et al., Convergent evolution of berberine biosynthesis.<br>Sci Adv. 2024 Nov 29, 10(48): eads3596. Epub 2024 Nov 29. |
| 1837 | *Bergenia scopulosa* (*Saxifragaceae*) | 2024-11-28 | Yang YX et al., The chromosome-level genome assembly of an endangered herb Bergenia scopulosa provides insights into local adaptation and genomic vulnerability under climate change.<br>Gigascience. 2024 Jan 2: 13: giae091. |
| 1836 | *Centaurea solstitialis* (*Asteraceae*) | 2024-11-27 | Reatini B et al., Chromosome-scale reference genome and RAD-based genetic map of yellow starthistle (Centaurea solstitialis) reveal putative structural variation and QTL associated with invader traits.<br>Genome Biol Evol. 2024 Nov 27: evae243. Online ahead of print. |
| 1835 | *Iodes seguinii* (*Icacinaceae*) | 2024-11-27 | Gong X et al., Chromosome-level genome assembly of Iodes seguinii and its metabonomic implications for rheumatoid arthritis treatment.<br>Plant Genome. 2024 Nov 27: e20534. Online ahead of print. |
| 1834 | *Solanum stoloniferum* (*Solanaceae*) | 2024-11-25 | Hosaka AJ et al., Allotetraploid nature of a wild potato species, Solanum stoloniferum Schlechtd. et Bche., as revealed by whole-genome sequencing.<br>Plant J. 2024 Nov 25. Online ahead of print. |
| 1833 | *Pellacalyx yunnanensis* (*Rhizophoraceae*) | 2024-11-22 | Shao S et al., Chromosomal-Level Genome Suggests Adaptive Constraints Leading to the Historical Population Decline in an Extremely Endangered Plant.<br>Mol Ecol Resour. 2024 Nov 22: e14045. Online ahead of print. |
| 1832 | *Mentha x piperita* (*Lamiaceae*) | 2024-11-19 | Talbot SC et al., A first look at the genome structure of hexaploid Mitcham peppermint (Mentha x piperita L.).<br>G3 (Bethesda). 2024 Nov 19: jkae195. Online ahead of print. |
| 1831 | *Clinopodium gracile* (*Lamiaceae*) | 2024-11-11 | Gao Y et al., Chromosome-level assembly of the Clinopodium gracile genome.<br>Front Plant Sci. 2024 Nov 11, 15: 1489102 eCollection 2024. |
| 1830 | *Ribes nigrum* (*Grossulariaceae*) | 2024-11-11 | Ziegler FMR et al., A full genome assembly reveals drought stress effects on gene expression and metabolite profiles in blackcurrant (Ribes nigrum L.).<br>Hortic Res. 2024 Nov: uhae313. eCollection 2024 Nov. |
| 1829 | *Euphrasia confusa* (*Orobanchaceae*) | 2024-11-07 | Twyford AD et al., The genome sequence of the confused eyebright, Euphrasia confusa Pugsley.<br>Wellcome Open Res. 2024 Nov 7: 9: 662. eCollection 2024. |
| 1828 | *Plinia cauliflora* (*Myrtaceae*) | 2024-11-07 | Zhao L et al., The telomere-to-telomere genome of Jaboticaba reveals the genetic basis of fruit color and citric acid content.<br>Int J Mol Sci. 2024 Nov 7, 25(22): 11951. |
| 1827 | *Uncaria rhynchophylla* (*Rubiaceae*) | 2024-11-07 | Hu T et al., Haploid-Phased Chromosomal Telomere-to-Telomere Genome Assembly of Medicinal Plant Uncaria rhynchophylla Dissects Genetic Controls on the Biosynthesis of Bioactive Alkaloids.<br>Plant Cell Environ. 2024 Nov 7. Online ahead of print. |
| 1826 | *Tillandsia duratii* (*Bromeliaceae*) | 2024-11-06 | Lyu X et al., Tracing the evolutionary and genetic footprints of atmospheric tillandsioids transition from land to air.<br>Nat Commun. 2024 Nov 6, 15(1): 9599. |
| 1825 | *Vitis davidii* (*Vitaceae*) | 2024-11-06 | Luo Y et al., Phased T2T genome assemblies facilitate the mining of disease-resistance genes in Vitis davidii.<br>Hortic Res. 2024 Nov: uhae306. eCollection 2024 Nov. |
| 1824 | *Vriesea erythrodactylon* (*Bromeliaceae*) | 2024-11-06 | Lyu X et al., Tracing the evolutionary and genetic footprints of atmospheric tillandsioids transition from land to air.<br>Nat Commun. 2024 Nov 6, 15(1): 9599. |
| 1823 | *Cryptomeria japonica* (*Cupressaceae*) | 2024-11-05 | Fujino T et al., A chromosome-level genome assembly of a model conifer plant, the Japanese cedar, Cryptomeria japonica D. Don.<br>BMC Genomics. 2024 Nov 5, 25(1): 1039. |
| 1822 | *Codonopsis pilosula* (*Campanulaceae*) | 2024-11-04 | Chen BZ et al., Chromosome-scale genome assembly of Codonopsis pilosula and comparative genomic analyses shed light on its genome evolution.<br>Front Plant Sci. 2024 Nov 05, 15: 1469375 eCollection 2024. |
| 1821 | *Medicago arabica* (*Fabaceae*) | 2024-11-04 | Christenhusz MJM et al., The genome sequence of spotted medick, Medicago arabica (L.) Huds. (Fabaceae)<br>Wellcome Open Res. 2024 Nov 4: 9: 116. eCollection 2024. |
| 1820 | *Salix cheilophila* (*Salicaceae*) | 2024-11-04 | Gao B et al., A telomere-to-telomere genome assembly of Salix cheilophila reveals its evolutionary signatures for environmental adaptation.<br>Plant Commun. 2024 Nov 4: 101182. Online ahead of print. |
| 1819 | *Vitis retordii* (*Vitaceae*) | 2024-11-04 | Liu Z et al., Grapevine pangenome facilitates trait genetics and genomic breeding.<br>Nat Genet (2024). |
| 1818 | *Rhodamnia argentea* (*Myrtaceae*) | 2024-11-02 | Chen SH et al., Chromosome-level genome assembly of the Australian rainforest tree Rhodamnia argentea (malletwood)<br>Genome Biol Evol. 2024 Nov 2: evae238. Online ahead of print. |

| | plant species | published | |
|---|---|---|---|
| 1817 | *Urtica urens* (*Urticaceae*) | 2024-11-01 | Christenhusz MJM et al., The genome sequence of the small nettle, Urtica urens L. (Urticaceae). Wellcome Open Res. 2024 Nov 1: 9: 639. eCollection 2024. |
| 1816 | *Dalbergia candenatensis* (*Fabaceae*) | 2024-10-31 | Shi M et al., Chromosome-scale genome assembly of the mangrove climber species Dalbergia candenatensis. Sci Data. 11, 1187 (2024). |
| 1815 | *Magnolia sieboldii* (*Magnoliaceae*) | 2024-10-30 | Lu X et al., The Chromosome-Scale Genome of Magnolia sieboldii K. Koch Provides Insight Into the Evolutionary Position of Magnoliids and Seed Germination. Mol Ecol Resour. 2024 Oct 30:e14030. Online ahead of print. |
| 1814 | *Curcuma petiolata* (*Zingiberaceae*) | 2024-10-29 | Liao X et al., Inversions encounter relaxed genetic constraints and balance birth and death of TPS genes in Curcuma. Nat Commun. 2024 Oct 29, 15(1): 9349. |
| 1813 | *Draba incana* (*Brassicaceae*) | 2024-10-29 | Ruhsam M et al., The genome sequence of Hoary Whitlowgrass, Draba incana L. Wellcome Open Res. 2024 Oct 29: 9: 630. eCollection 2024. |
| 1812 | *Gaultheria prostrata* (*Ericaceae*) | 2024-10-29 | Lin YJ et al., First De Novo genome assembly and characterization of Gaultheria prostrata. Front Plant Sci. 2024 Oct 29: 15: 1456102. eCollection 2024. |
| 1811 | *Lilium davidii* (*Liliaceae*) | 2024-10-24 | Xu S et al., The evolutionary tale of lilies: Giant genomes derived from transposon insertions and polyploidization. Innovation (Camb). 2024 Oct 24, 5(6): 100726. eCollection 2024 Nov 4. |
| 1810 | *Penstemon eatonii* (*Plantaginaceae*) | 2024-10-23 | Jarvis DE et al., Whole-genome assembly and annotation of the firecracker penstemon (Penstemon eatonii). J Hered. 2024 Oct 23: esae058. Online ahead of print. |
| 1809 | *Pinus densiflora* (*Pinaceae*) | 2024-10-20 | Jang MJ et al., Haplotype-resolved genome assembly and resequencing analysis provide insights into genome evolution and allelic imbalance in Pinus densiflora. Nat Genet. (2024). |
| 1808 | *Inga laevigata* (*Fabaceae*) | 2024-10-17 | Schley RJ et al., The genome sequence of Inga leiocalycina Benth. Wellcome Open Res. 2024 Oct 17: 9: 606. eCollection 2024. |
| 1807 | *Inga oerstediana* (*Fabaceae*) | 2024-10-17 | Schley RJ et al., The genome sequence of Inga oerstediana Benth. Wellcome Open Res. 2024 Oct 17: 9: 607. eCollection 2024. |
| 1806 | *Sambucus nigra* (*Viburnaceae*) | 2024-10-17 | Christenhusz MJM et al., The genome sequence of black elder, Sambucus nigra Linnaeus, 1753 (Adoxaceae). Wellcome Open Res. 2024 Oct 17: 9: 609. eCollection 2024. |
| 1805 | *Beckmannia syzigachne* (*Poaceae*) | 2024-10-16 | Han Y et al., Assembling a reference-quality genome and resequencing diverse accessions of Beckmannia syzigachne provide insights into population structure and gene family evolution. Plant Commun. 2024 Oct 15: 101174. Online ahead of print. |
| 1804 | *Arctium minus* (*Asteraceae*) | 2024-10-15 | Christenhusz MJM et al., The genome sequence of lesser burdock, Arctium minus (Hill) Bernh. (Asteraceae). Wellcome Open Res. 2024 Oct 15: 9: 589. eCollection 2024. |
| 1803 | *Adansonia digitata* (*Malvaceae*) | 2024-10-12 | Kitony JK et al., Chromosome-level baobab genome illuminates its evolutionary trajectory and environmental adaptation. Nat Commun. 15, 8833 (2024). |
| 1802 | *Rosa hybrid cv. Samantha* (*Rosaceae*) | 2024-10-11 | Zhang Z et al., Haplotype-resolved genome assembly and resequencing provide insights into the origin and breeding of modern rose. Nat Plants. (2024). |
| 1801 | *Digitaria radicosa* (*Poaceae*) | 2024-10-10 | Minoji K et al., A chromosome-scale genome assembly of Timorese crabgrass (Digitaria radicosa): a useful genomic resource for the Poaceae. G3 (Bethesda). 2024 Oct 10: jkae242. Online ahead of print. |
| 1800 | *Medicago lupulina* (*Fabaceae*) | 2024-10-09 | Ruhsam M et al., The genome sequence of the Black Medic, Medicago lupulina L. Wellcome Open Res. 2024 Oct 9: 9: 574. eCollection 2024. |
| 1799 | *Malpighia emarginata* (*Malpighiaceae*) | 2024-10-07 | Shirasawa K et al., Chromosome-scale genome assembly of acerola (Malpighia emarginata DC.). DNA Res. 2024 Oct 5: dsae029. |
| 1798 | *Liquidambar styraciflua* (*Altingiaceae*) | 2024-10-03 | Ma Y et al., Chromosome-level genome assembly of American sweetgum (Liquidambar styraciflua, Altingiaceae). Sci Data. 2024 Oct 3, 11(1): 1078. |
| 1797 | *Salvia divinorum* (*Lamiaceae*) | 2024-10-01 | Ford SA et al., A chromosome level reference genome of Diviner's sage (Salvia divinorum) provides insight into salvinorin A biosynthesis. BMC Plant Biol. 2024 Oct 1, 24(1): 914. |
| 1796 | *Rosa gigantea* (*Rosaceae*) | 2024-09-30 | Zhou L et al., Multi-omics analyzes of Rosa gigantea illuminate tea scent biosynthesis and release mechanisms. Nat Commun. 2024 Sep 30, 15(1): 8469. |
| 1795 | *Rhododendron bailiense* (*Ericaceae*) | 2024-09-29 | Wen S et al., Chromosome-level genome assembly of a rare karst-growing Rhododendron species provides insights into its evolution and environmental adaptation. J Syst Evol. 2024, Online ahead of print. |
| 1794 | *Cirsium nipponicum* (*Asteraceae*) | 2024-09-27 | Choi BY et al., De Novo Genome Assembly and Phylogenetic Analysis of Cirsium nipponicum. Genes. 2024, 15, 1269. |
| 1793 | *Ilex hylonoma* (*Aquifoliaceae*) | 2024-09-27 | Feng L et al., Integrated genomic, transcriptomic, and metabolomic analyses of Ilex hylonoma provide insights into the triterpenoid saponin biosynthesis. Plant J. 2024 Sep 27. Online ahead of print. |
| 1792 | *Ziziphus mauritiana* (*Rhamnaceae*) | 2024-09-26 | Guo M et al., Genomes of autotetraploid wild and cultivated Ziziphus mauritiana reveal polyploid evolution and crop domestication. Plant Physiol. 2024 Sep 26: kiae512. Online ahead of print. |
| 1791 | *Citrus garrawayae* (*Rutaceae*) | 2024-09-24 | Nakandala U et al., The genomes of Australian wild limes. Plant Mol Biol. 2024 Sep 24, 114(5): 102. |
| 1790 | *Fraxinus angustifolia* (*Oleaceae*) | 2024-09-21 | Liu JN et al., Pan-genome analyses of eleven Fraxinus species provide insights into salt adaptation in ash trees. Plant Commun. 2024 Sep 21: 101137. Online ahead of print. |
| 1789 | *Fraxinus baroniana* (*Oleaceae*) | 2024-09-21 | Liu JN et al., Pan-genome analyses of eleven Fraxinus species provide insights into salt adaptation in ash trees. Plant Commun. 2024 Sep 21: 101137. Online ahead of print. |
| 1788 | *Fraxinus chinensis* (*Oleaceae*) | 2024-09-21 | Liu JN et al., Pan-genome analyses of eleven Fraxinus species provide insights into salt adaptation in ash trees. Plant Commun. 2024 Sep 21: 101137. Online ahead of print. |
| 1787 | *Fraxinus hubeiensis* (*Oleaceae*) | 2024-09-21 | Liu JN et al., Pan-genome analyses of eleven Fraxinus species provide insights into salt adaptation in ash trees. Plant Commun. 2024 Sep 21: 101137. Online ahead of print. |
| 1786 | *Fraxinus mandshurica* (*Oleaceae*) | 2024-09-21 | Liu JN et al., Pan-genome analyses of eleven Fraxinus species provide insights into salt adaptation in ash trees. Plant Commun. 2024 Sep 21: 101137. Online ahead of print. |
| 1785 | *Fraxinus ornus* (*Oleaceae*) | 2024-09-21 | Liu JN et al., Pan-genome analyses of eleven Fraxinus species provide insights into salt adaptation in ash trees. Plant Commun. 2024 Sep 21: 101137. Online ahead of print. |
| 1784 | *Fraxinus sogdiana* (*Oleaceae*) | 2024-09-21 | Liu JN et al., Pan-genome analyses of eleven Fraxinus species provide insights into salt adaptation in ash trees. Plant Commun. 2024 Sep 21: 101137. Online ahead of print. |
| 1783 | *Fraxinus velutina* (*Oleaceae*) | 2024-09-21 | Liu JN et al., Pan-genome analyses of eleven Fraxinus species provide insights into salt adaptation in ash trees. Plant Commun. 2024 Sep 21: 101137. Online ahead of print. |
| 1782 | *Amphiroa fragilissima* (*Lithophyllaceae*) | 2024-09-20 | Zhang H et al., Genome of Halimeda opuntia reveals differentiation of subgenomes and molecular bases of multinucleation and calcification in algae. Proc Natl Acad Sci U S A. 2024 Sep 24, 121(39): e2403222121. Epub 2024 Sep 20. |
| 1781 | *Halimeda opuntia* (*Halimedaceae*) | 2024-09-20 | Zhang H et al., Genome of Halimeda opuntia reveals differentiation of subgenomes and molecular bases of multinucleation and calcification in algae. Proc Natl Acad Sci U S A. 2024 Sep 24, 121(39): e2403222121. Epub 2024 Sep 20. |

| | plant species | published | |
|---|---|---|---|
| 1780 | *Antiaris toxicaria* (*Moraceae*) | 2024-09-18 | Miao K et al., Haplotype-resolved genome assembly of the upas tree (Antiaris toxicaria).<br>Sci Data. 2024 Sep 18, 11(1): 1011. |
| 1779 | *Bromus sterilis* (*Poaceae*) | 2024-09-18 | Christenhusz MJM et al., The genome sequence of barren brome, Bromus sterilis L. (Poaceae).<br>Wellcome Open Res. 2024 Sep 18: 9: 534. eCollection 2024. |
| 1778 | *Coptis teeta* (*Ranunculaceae*) | 2024-09-18 | Wang Y et al., A haplotype-resolved genome assembly of Coptis teeta, an endangered plant of significant medicinal value.<br>Sci Data. 2024 Sep 18, 11(1): 1012. |
| 1777 | *Madhuca longifolia* (*Sapotaceae*) | 2024-09-18 | Wang S et al., High-quality chromosome-level genomic insights into molecular adaptation to low-temperature stress in Madhuca longifolia in southern subtropical China.<br>BMC Genomics. 2024 Sep 18, 25(1): 877. |
| 1776 | *Rhododendron championiae* (*Ericaceae*) | 2024-09-18 | Xia XM et al., Genomic insights into adaptive evolution of the species-rich cosmopolitan plant genus Rhododendron.<br>Cell Rep. 2024 Sep 18, 43(10): 114745. Online ahead of print. |
| 1775 | *Rhododendron micranthum* (*Ericaceae*) | 2024-09-18 | Xia XM et al., Genomic insights into adaptive evolution of the species-rich cosmopolitan plant genus Rhododendron.<br>Cell Rep. 2024 Sep 18, 43(10): 114745. Online ahead of print. |
| 1774 | *Rhododendron mucronulatum* (*Ericaceae*) | 2024-09-18 | Xia XM et al., Genomic insights into adaptive evolution of the species-rich cosmopolitan plant genus Rhododendron.<br>Cell Rep. 2024 Sep 18, 43(10): 114745. Online ahead of print. |
| 1773 | *Rhododendron neriiflorum* (*Ericaceae*) | 2024-09-18 | Xia XM et al., Genomic insights into adaptive evolution of the species-rich cosmopolitan plant genus Rhododendron.<br>Cell Rep. 2024 Sep 18, 43(10): 114745. Online ahead of print. |
| 1772 | *Rhododendron redowskianum* (*Ericaceae*) | 2024-09-18 | Xia XM et al., Genomic insights into adaptive evolution of the species-rich cosmopolitan plant genus Rhododendron.<br>Cell Rep. 2024 Sep 18, 43(10): 114745. Online ahead of print. |
| 1771 | *Lupinus mutabilis* (*Fabaceae*) | 2024-09-12 | Pancaldi F et al., The genome of Lupinus mutabilis: Evolution and genetics of an emerging bio-based crop.<br>Plant J. 2024 Sep 12. Online ahead of print. |
| 1770 | *Comarum palustre* (*Rosaceae*) | 2024-09-05 | Christenhusz MJM et al., The genome sequence of the marsh cinquefoil, Comarum palustre L., also known as Potentilla palustris (L.) Scop. (Rosaceae).<br>Wellcome Open Res. 2024 Sep 5, 9,518. eCollection 2024. |
| 1769 | *Stephania longa* (*Menispermaceae*) | 2024-09-05 | Shang H et al., Genome assembly of Stephania longa provides insight into cepharanthine biosynthesis.<br>Front Plant Sci. 2024 Sep 05, 15: 1414636 eCollection 2024. |
| 1768 | *Bambusa odashimae* (*Poaceae*) | 2024-09-04 | Wang YJ et al., Haplotype-resolved nonaploid genome provides insights into in vitro flowering in bamboo.<br>Hortic Res. 2024 Sep: uhae250. eCollection 2024 Sep. |
| 1767 | *Chenopodium album* (*Amaranthaceae*) | 2024-09-04 | Mian S et al., The genome sequence of fat-hen, Chenopodium album L.<br>Wellcome Open Res. 2024 Sep 4, 9: 508. eCollection 2024. |
| 1766 | *Gongora gibba* (*Orchidaceae*) | 2024-09-04 | Guizar Amador MF et al., The Gongora gibba genome assembly provides new insights into the evolution of floral scent in male euglossine bee-pollinated orchids.<br>G3 (Bethesda). 2024 Sep 4: jkae211. Online ahead of print. |
| 1765 | *Sageretia thea* (*Rhamnaceae*) | 2024-09-04 | Jo J et al., The first chromosomal-level genome assembly of Sageretia thea using Nanopore long reads and Pore-C technology.<br>Sci Data. 2024 Sep 4, 11(1): 959. |
| 1764 | *Chionanthus retusus* (*Oleaceae*) | 2024-09-03 | Wang J et al., A telomere-to-telomere gap-free reference genome of Chionanthus retusus provides insights into the molecular mechanism underlying petal shape changes.<br>Hortic Res. 2024 Sep: uhae249. eCollection 2024 Sep. |
| 1763 | *Canavalia gladiata* (*Fabaceae*) | 2024-08-27 | Yuan L et al., The genomes of 5 underutilized Papilionoideae crops provide insights into root nodulation and disease resistance.<br>Gigascience. 2024: 13: giae063. |
| 1762 | *Clitoria ternatea* (*Fabaceae*) | 2024-08-27 | Yuan L et al., The genomes of 5 underutilized Papilionoideae crops provide insights into root nodulation and disease resistance.<br>Gigascience. 2024: 13: giae063. |
| 1761 | *Crotalaria linifolia* (*Fabaceae*) | 2024-08-27 | Yuan L et al., The genomes of 5 underutilized Papilionoideae crops provide insights into root nodulation and disease resistance.<br>Gigascience. 2024: 13: giae063. |
| 1760 | *Petroselinum crispum* (*Apiaceae*) | 2024-08-27 | Uncu AT et al., Whole-genome sequencing and identification of antimicrobial peptide coding genes in parsley (Petroselinum crispum), an important culinary and medicinal Apiaceae species.<br>Funct Integr Genomics. 2024 Aug 27, 24(5): 142. |
| 1759 | *Phaseolus coccineus* (*Fabaceae*) | 2024-08-27 | Yuan L et al., The genomes of 5 underutilized Papilionoideae crops provide insights into root nodulation and disease resistance.<br>Gigascience. 2024: 13: giae063. |
| 1758 | *Roscoea debilis* (*Zingiberaceae*) | 2024-08-27 | Peng XC et al., Draft genome of Roscoea Debilis, the first genome in the alpine ginger Roscoea (Zingiberaceae).<br>BMC Genom Data. 2024 Aug 27, 25(1): 77. |
| 1757 | *Malus coronaria* (*Rosaceae*) | 2024-08-26 | Svara A et al., Advancing apple genetics research: Malus coronaria and Malus ioensis genomes and a gene family-based pangenome of native North American apples.<br>DNA Res. 2024 Aug 26: dsae026. Online ahead of print. |
| 1756 | *Malus ioensis* (*Rosaceae*) | 2024-08-26 | Svara A et al., Advancing apple genetics research: Malus coronaria and Malus ioensis genomes and a gene family-based pangenome of native North American apples.<br>DNA Res. 2024 Aug 26: dsae026. Online ahead of print. |
| 1755 | *Eleocharis vivipara* (*Cyperaceae*) | 2024-08-23 | Liu H et al., The genome of Eleocharis vivipara elucidates the genetics of C3-C4 photosynthetic plasticity and karyotype evolution in the Cyperaceae.<br>J Integr Plant Biol. 2024 Aug 23. Online ahead of print. |
| 1754 | *Hevea benthamiana* (*Euphorbiaceae*) | 2024-08-22 | Fang Y et al., Pan-genome and phylogenomic analyses highlight Hevea species delineation and rubber trait evolution.<br>Nat Commun. 2024 Aug 22, 15(1): 7232. |
| 1753 | *Hevea nitida* (*Euphorbiaceae*) | 2024-08-22 | Fang Y et al., Pan-genome and phylogenomic analyses highlight Hevea species delineation and rubber trait evolution.<br>Nat Commun. 2024 Aug 22, 15(1): 7232. |
| 1752 | *Hevea pauciflora* (*Euphorbiaceae*) | 2024-08-22 | Fang Y et al., Pan-genome and phylogenomic analyses highlight Hevea species delineation and rubber trait evolution.<br>Nat Commun. 2024 Aug 22, 15(1): 7232. |
| 1751 | *Chrysosplenium sinicum* (*Saxifragaceae*) | 2024-08-16 | Liu S et al., The Chrysosplenium sinicum genome provides insights into adaptive evolution of shade plants.<br>Commun Biol. 2024 Aug 16, 7(1): 1004. |
| 1750 | *Malus delavayi* (*Rosaceae*) | 2024-08-16 | Tian J et al., Chromosome-scale genome assembly of Docynia delavayi provides new insights into the alpha-Farnesene biosynthesis.<br>Int J Biol Macromol. 2024 Aug 16: 134820. Online ahead of print. |
| 1749 | *Flueggea virosa* (*Phyllanthaceae*) | 2024-08-13 | Chen BZ et al., Chromosome-level genome assembly and annotation of Flueggea virosa (Phyllanthaceae).<br>Sci Data. 2024 Aug 13, 11(1): 875. |
| 1748 | *Hakea prostrata* (*Proteaceae*) | 2024-08-13 | Nestor BJ et al., Identification and expression analysis of Phosphate Transporter 1 (PHT1) genes in the highly phosphorus-use-efficient Hakea prostrata (Proteaceae).<br>Plant Cell Environ. 2024 Aug 13. Online ahead of print. |
| 1747 | *Salix babylonica* (*Salicaceae*) | 2024-08-12 | He L et al., Allopolyploidization from two dioecious ancestors leads to recurrent evolution of sex chromosomes.<br>Nat Commun. 15, 6893 (2024). |
| 1746 | *Trichosanthes truncata* (*Cucurbitaceae*) | 2024-08-12 | Hu Y et al., Chromosome-level assembly and evolution analysis of the Trichosanthes truncata genome.<br>Sci Data. 2024 Aug 12, 11(1): 872. |
| 1745 | *Eleocharis dulcis* (*Cyperaceae*) | 2024-08-10 | Chen Y et al., Telomere-to-telomere genome assembly of Eleocharis dulcis and expression profiles during corm development.<br>Sci Data. 11, 869 (2024). |
| 1744 | *Cochlearia pyrenaica* (*Brassicaceae*) | 2024-08-07 | Bray SM et al., Kinetochore and ionomic adaptation to whole-genome duplication in Cochlearia shows evolutionary convergence in three autopolyploids.<br>Cell Rep. 2024 Aug 7, 43(8): 114576. Online ahead of print. |

| | plant species | published | |
|---|---|---|---|
| 1743 | *Rhododendron nivale* *(Ericaceae)* | 2024-08-07 | Lyu ZY et al., The first high-altitude autotetraploid haplotype-resolved genome assembled (Rhododendron nivale subsp. boreale) provides new insights into mountaintop adaptation. Gigascience. 2024 Jan 2: 13: giae052. |
| 1742 | *Kokia cookei* *(Malvaceae)* | 2024-08-06 | Kayal E et al., Genomic diversity and evolution in the Hawaiian Islands endemic Kokia (Malvaceae). G3 (Bethesda). 2024 Aug 6: jkae180. Online ahead of print. |
| 1741 | *Kokia drynarioides* *(Malvaceae)* | 2024-08-06 | Kayal E et al., Genomic diversity and evolution in the Hawaiian Islands endemic Kokia (Malvaceae). G3 (Bethesda). 2024 Aug 6: jkae180. Online ahead of print. |
| 1740 | *Kokia kauaiensis* *(Malvaceae)* | 2024-08-06 | Kayal E et al., Genomic diversity and evolution in the Hawaiian Islands endemic Kokia (Malvaceae). G3 (Bethesda). 2024 Aug 6: jkae180. Online ahead of print. |
| 1739 | *Syzygium maire* *(Myrtaceae)* | 2024-08-06 | Balkwill CG et al., De-novo assembly of a reference genome for a critically threatened Aotearoa New Zealand tree species, Syzygium maire (Myrtaceae). Tree Genet Genomes. 2024, 20(5) |
| 1738 | *Miscanthus x giganteus* *(Poaceae)* | 2024-08-05 | Zeng X et al., Chromosome-level scaffolding of haplotype-resolved assemblies using Hi-C data without reference genomes. Nat Plants. 2024 Aug 5. Online ahead of print. |
| 1737 | *Rubus leucanthus* *(Rosaceae)* | 2024-08-05 | Wu W et al., A high-quality genome assembly reveals adaptations underlying glossy, wax-coated leaves in the heat-tolerant wild raspberry Rubus leucanthus. DNA Res. 2024 Aug 5: dsae024. Online ahead of print. |
| 1736 | *Lindenbergia philippensis* *(Orobanchaceae)* | 2024-08-02 | Chen BZ et al., Chromosome-level assembly of Lindenbergia philippensis and comparative genomic analyses shed light on genome evolution in Lamiales. Front Plant Sci. 2024 Aug 02, 15: 1444234 eCollection 2024. |
| 1735 | *Sanguina aurantia* *(Chlamydomonadaceae)* | 2024-08-02 | Raymond BB et al., Snow alga Sanguina aurantia as revealed through de novo genome assembly and annotation. G3 (Bethesda). 2024 Aug 02: jkae181. Online ahead of print. |
| 1734 | *Solanum cleistogamum* *(Solanaceae)* | 2024-08-02 | Satterlee JW et al., Convergent evolution of plant prickles by repeated gene co-option over deep time. Science. 2024 Aug 2, 385(6708): eado1663. Epub 2024 Aug 2. |
| 1733 | *Solanum prinophyllum* *(Solanaceae)* | 2024-08-02 | Satterlee JW et al., Convergent evolution of plant prickles by repeated gene co-option over deep time. Science. 2024 Aug 2, 385(6708): eado1663. Epub 2024 Aug 2. |
| 1732 | *Hygrophila difformis* *(Acanthaceae)* | 2024-07-30 | Li G et al., Water wisteria genome reveals environmental adaptation and heterophylly regulation in amphibious plants. Plant Cell Environ. 2024 Jul 30. Online ahead of print. |
| 1731 | *Biscutella laevigata* *(Brassicaceae)* | 2024-07-29 | Beringer M et al., Biased retention of environment-responsive genes following genome fractionation. Mol Biol Evol. 2024 Jul 29: msae155. Online ahead of print. |
| 1730 | *Elymus sibiricus* *(Poaceae)* | 2024-07-26 | Shen W et al., Chromosome-scale assembly of the wild cereal relative Elymus sibiricus. Sci Data. 2024 Jul 26, 11(1): 823. |
| 1729 | *Ophrys sphegodes* *(Orchidaceae)* | 2024-07-26 | Russo A et al., Genome of the early spider-orchid Ophrys sphegodes provides insights into sexual deception and pollinator adaptation. Nat Commun. 15, 6308 (2024). |
| 1728 | *Drepanocaryum sewerzowi* *(Lamiaceae)* | 2024-07-24 | Smit SJ et al., Genome Report: Pseudomolecule-scale genome assemblies of Drepanocaryum sewerzowii and Marmoritis complanata. G3 (Bethesda). 2024 Jul 24: jkae172. Online ahead of print. |
| 1727 | *Houttuynia cordata* *(Saururaceae)* | 2024-07-24 | Huang P et al., A Genome Assembly of decaploid Houttuynia cordata Provides Insights into the evolution of Houttuynia and the biosynthesis of alkaloids. Hortic Res. 2024 Jul: uhae203. eCollection 2024 Jul. |
| 1726 | *Nepeta complanata* *(Lamiaceae)* | 2024-07-24 | Smit SJ et al., Genome Report: Pseudomolecule-scale genome assemblies of Drepanocaryum sewerzowii and Marmoritis complanata. G3 (Bethesda). 2024 Jul 24: jkae172. Online ahead of print. |
| 1725 | *Saponaria officinalis* *(Caryophyllaceae)* | 2024-07-23 | Jo S et al., Unlocking saponin biosynthesis in soapwort. Nat Chem Biol. (2024). |
| 1724 | *Reaumuria soongarica* *(Tamaricaceae)* | 2024-07-22 | Song M et al., Chromosome-level genome assembly and annotation of xerophyte secretohalophyte Reaumuria soongarica. Sci Data. 11, 812 (2024). |
| 1723 | *Cananga odorata* *(Annonaceae)* | 2024-07-19 | Zheng Y et al., The chromosome-level genome assembly of Cananga odorata provides insights into its evolution and terpenoid biosynthesis. New Phytol. 2024 Jul 19. Online ahead of print. |
| 1722 | *Cochlearia groenlandica* *(Brassicaceae)* | 2024-07-18 | Kim J et al., Whole-genome sequencing of 13 Arctic plants and draft genomes of Oxyria digyna and Cochlearia groenlandica. Sci Data. 2024 Jul 18, 11(1): 793. |
| 1721 | *Oxyria digyna* *(Polygonaceae)* | 2024-07-18 | Kim J et al., Whole-genome sequencing of 13 Arctic plants and draft genomes of Oxyria digyna and Cochlearia groenlandica. Sci Data. 2024 Jul 18, 11(1): 793. |
| 1720 | *Pinellia ternata* *(Araceae)* | 2024-07-18 | Xue T et al., A chromosome-level Pinellia ternata genome assembly provides insight into the evolutionary origin of ephedrine and acrid raphide formation. Med Plant Biol. 3: e013 |
| 1719 | *Aquilaria yunnanensis* *(Thymelaeaceae)* | 2024-07-17 | Li M et al., Chromosome-level genome assembly of Aquilaria yunnanensis. Sci Data. 2024 Jul 17, 11(1): 790. |
| 1718 | *Gentiana straminea* *(Gentianaceae)* | 2024-07-17 | Ala KG et al., Insights from the first chromosome-level genome assembly of the alpine gentian Gentiana straminea Maxim. (Insight from genome assembly of Gentiana straminea). DNA Res. 2024 Jul 17: dsae022. Online ahead of print. |
| 1717 | *Geum urbanum* *(Rosaceae)* | 2024-07-16 | Christenhusz MJM et al., The genome sequence of wood avens, Geum urbanum L., 1753. Wellcome Open Res. 2024 Jul 16:8:371. eCollection 2023. |
| 1716 | *Magnolia kwangsiensis* *(Magnoliaceae)* | 2024-07-13 | Shi CY et al., A high-quality chromosome-level genome assembly of the endangered tree Kmeria septentrionalis. Sci Data. 2024 Jul 13, 11(1): 775. |
| 1715 | *Hibiscus sabdariffa* *(Malvaceae)* | 2024-07-12 | Kim T et al., Genome assembly of Hibiscus sabdariffa L. provides insights into metabolisms of medicinal natural products. G3 (Bethesda). 2024 Jul 12: jkae134. Online ahead of print. |
| 1714 | *Fragaria chinensis* *(Rosaceae)* | 2024-07-11 | Lin H et al., Genomic data provides insights into the evolutionary history and adaptive differentiation of two tetraploid strawberries. Hortic Res. 2024 Jul: uhae194. eCollection 2024 Jul. |
| 1713 | *Fragaria corymbosa* *(Rosaceae)* | 2024-07-11 | Lin H et al., Genomic data provides insights into the evolutionary history and adaptive differentiation of two tetraploid strawberries. Hortic Res. 2024 Jul: uhae194. eCollection 2024 Jul. |
| 1712 | *Citrullus rehmii* *(Cucurbitaceae)* | 2024-07-08 | Zhang Y et al., Telomere-to-telomere Citrullus super-pangenome provides direction for watermelon breeding. Nat Genet. 2024 Jul 08. Online ahead of print. |
| 1711 | *Citrullus ecirrhosus* *(Cucurbitaceae)* | 2024-07-08 | Zhang Y et al., Telomere-to-telomere Citrullus super-pangenome provides direction for watermelon breeding. Nat Genet. 2024 Jul 08. Online ahead of print. |
| 1710 | *Citrullus naudinianus* *(Cucurbitaceae)* | 2024-07-08 | Zhang Y et al., Telomere-to-telomere Citrullus super-pangenome provides direction for watermelon breeding. Nat Genet. 2024 Jul 08. Online ahead of print. |
| 1709 | *Helwingia omeiensis* *(Helwingiaceae)* | 2024-07-02 | Chen Y et al., Chromosome-level genome assembly of Helwingia omeiensis: the first genome in the family Helwingiaceae. Sci Data. 11, 719 (2024). |
| 1708 | *Orobanche coerulescens* *(Orobanchaceae)* | 2024-07-02 | Kim B et al., Chromosome-level genome assembly of Korean holoparasitic plants, Orobanche coerulescens. Sci Data. 11, 714 (2024) |
| 1707 | *Schoenoplectus tabernaemontani* *(Cyperaceae)* | 2024-07-01 | Li Y et al., Chromosome-scale genome assembly for soft-stem bulrush (Schoenoplectus tabernaemontani) confirms a clade-specific whole genome duplication in Cyperaceae. Genome Biol Evol. 2024 Jul 01: evae141. Online ahead of print. |

| | plant species | published | |
|---|---|---|---|
| 1706 | *Tamarix austromongolica* *(Tamaricaceae)* | 2024-07-01 | Gong S et al., A Chromosome-level Genome Assembly Provides Insights into the Local Adaptation of Tamarix austromongolica in the Yellow River Basin, China. DNA Res. 2024 Jul 01: dsae021. Online ahead of print. |
| 1705 | *cf. Sphaerocystis sp.* *(Sphaerocystidaceae)* | 2024-06-28 | Ciftci O et al., Long-read based hybrid genome assembly and annotation of snow algal strain CCCryo 101-99 (cf. Sphaerocystis sp., Chlamydomonadales). Genome Biol Evol. 2024 Jun 28: evae140. Online ahead of print. |
| 1704 | *Corymbia calophylla* *(Myrtaceae)* | 2024-06-27 | Ahrens CW et al., Genomic determinants, architecture, and constraints in drought-related traits in Corymbia calophylla. BMC Genomics. 2024 Jun 27, 25(1): 640. |
| 1703 | *Dendrobium loddigesii* *(Orchidaceae)* | 2024-06-27 | Du Z et al., Multi-omics analyses and botanical perfumer hypothesis provide insights into the formation and maintenance of aromatic characteristics of Dendrobium loddigesii flowers. Plant Physiol Biochem. 2024 Jun 27: 214: 108891. Online ahead of print. |
| 1702 | *Camphora longepaniculata* *(Lauraceae)* | 2024-06-26 | Yan K et al., Chromosome genome assembly of the Camphora longepaniculata (Gamble) with PacBio and Hi-C sequencing data. Front Plant Sci. 2024 Jun 26, 15: 1372127. eCollection 2024. |
| 1701 | *Elaeis oleifera* *(Arecaceae)* | 2024-06-26 | Low ETL et al., Chromosome-scale Elaeis guineensis and E. oleifera assemblies: Comparative genomics of oil palm and other Arecaceae. G3 (Bethesda). 2024 Jun 26: jkae135. Online ahead of print. |
| 1700 | *Geum rupestre* *(Rosaceae)* | 2024-06-25 | Li WG et al., Chromosome-level genome assembly of a cliff plant Taihangia rupestris var. ciliata provides insights into its adaptation and demographic history. BMC Plant Biol. 2024 Jun 25, 24(1): 596. |
| 1699 | *Castanea sativa* *(Fagaceae)* | 2024-06-22 | Bianco L et al., The de novo, chromosome-level genome assembly of the sweet chestnut (Castanea sativa Mill.) Cv. Marrone Di Chiusa Pesio. BMC Genom Data. 2024 Jun 22, 25(1): 64. |
| 1698 | *Cnidium monnieri* *(Apiaceae)* | 2024-06-22 | Wang Z et al., Chromosome-level genome assembly of Cnidium monnieri, a highly demanded traditional Chinese medicine. Sci Data. 2024 Jun 22, 11(1): 667. |
| 1697 | *Atractylodes lancea* *(Asteraceae)* | 2024-06-21 | Zhang C et al., Genome resequencing reveals the genetic basis of Population Evolution, local adaptation, and rewiring of the rhizomes metabolome in Atractylodes lancea. Hortic Res. 2024 Jun: uhae167. eCollection 2024 Jun. |
| 1696 | *Euphorbia tirucalli* *(Euphorbiaceae)* | 2024-06-21 | Wei Z et al., Chromosome-level genome assembly of Euphorbia tirucalli (Euphorbiaceae), a highly stress-tolerant oil plant Sci Data. 2024 Jun 21, 11(1): 658. |
| 1695 | *Prunus pseudocerasus* *(Rosaceae)* | 2024-06-21 | Jiu S et al., Haplotype-resolved genome assembly for tetraploid Chinese cherry (Prunus pseudocerasus) offers insights into fruit firmness. Hortic Res. 2024 Jun: uhae142. eCollection 2024 Jun. |
| 1694 | *Ajuga decumbens* *(Lamiaceae)* | 2024-06-19 | Gao Y et al., Chromosome-level genome assembly of Ajuga decumbens. Front Plant Sci. 2024 Jun 19: 15: 1413468. eCollection 2024. |
| 1693 | *Prunus conradinae* *(Rosaceae)* | 2024-06-19 | Jiu S et al., Chromosome-level genome assembly provides insights into the genetic diversity, evolution, and flower development of Prunus conradinae. Mol Hortic. 2024 Jun 19, 4(1): 25. |
| 1692 | *Sinojackia xylocarpa* *(Styracaceae)* | 2024-06-18 | Zhu S et al., The jacktree genome and population genomics provides insights for the mechanisms of the germination obstacle and the conservation of endangered ornamental plants. Hortic Res. 2024 Jun: uhae166. eCollection 2024 Jun. |
| 1691 | *Vaccinium floribundum* *(Ericaceae)* | 2024-06-18 | Albuja-Quintana M et al., Genome Report: First reference genome of Vaccinium floribundum Kunth, an emblematic Andean species. G3 (Bethesda). 2024 Jun 18: jkae136. Online ahead of print. |
| 1690 | *Ischnosiphon puberulus* *(Marantaceae)* | 2024-06-17 | Palma-Silva C et al., Ecological transcriptomics reveals stress response pathways of a ground-herb species in a waterlogging gradient of Amazonian riparian forests. Mol Ecol. 2024 Jun 17: e17437. Online ahead of print. |
| 1689 | *Lepidium apetalum* *(Brassicaceae)* | 2024-06-17 | Yan H et al., A chromosome-level genome assembly and annotation of the medicinal plant Lepidium apetalum. BMC Genom Data. 2024 Jun 17, 25(1): 61. |
| 1688 | *Bidens alba* *(Asteraceae)* | 2024-06-13 | Wang H et al., The genomes of Dahlia pinnata, Cosmos bipinnatus, and Bidens alba in tribe Coreopsideae provide insights into polyploid evolution and inulin biosynthesis. Gigascience. 2024 Jan 2: 13: giae032. |
| 1687 | *Climacium dendroides* *(Climaciaceae)* | 2024-06-13 | Bell D et al., The genome sequence of the tree-moss, Climacium dendroides (Hedw.) F.Weber & D.Mohr (Climaciaceae). Wellcome Open Res. 2024 Jun 13: 9: 311. eCollection 2024. |
| 1686 | *Cosmos bipinnatus* *(Asteraceae)* | 2024-06-13 | Wang H et al., The genomes of Dahlia pinnata, Cosmos bipinnatus, and Bidens alba in tribe Coreopsideae provide insights into polyploid evolution and inulin biosynthesis. Gigascience. 2024 Jan 2: 13: giae032. |
| 1685 | *Dahlia pinnata* *(Asteraceae)* | 2024-06-13 | Wang H et al., The genomes of Dahlia pinnata, Cosmos bipinnatus, and Bidens alba in tribe Coreopsideae provide insights into polyploid evolution and inulin biosynthesis. Gigascience. 2024 Jan 2: 13: giae032. |
| 1684 | *Paratinospora sagittata* *(Menispermaceae)* | 2024-06-12 | Alami MM et al., Chromosome-scale genome assembly of medicinal plant Tinospora sagittata (Oliv.) Gagnep. from the Menispermaceae family. Sci Data. 11, 610 (2024). |
| 1683 | *Prasiola crispa* *(Prasiolaceae)* | 2024-06-10 | Kosugi M et al., Characterization of the far-red light absorbing light-harvesting chlorophyll a/b binding complex, a derivative of the distinctive Lhca gene family in green algae. Front Plant Sci. 2024 Jun 10, 15: 1409116. eCollection 2024. |
| 1682 | *Pyrus x sinkiangensis* *(Rosaceae)* | 2024-06-10 | Li Q et al., Haplotype-resolved T2T genome assemblies and pangenome graph of pear reveal diverse patterns of allele-specific expression and genomic basis of fruit quality traits. Plant Commun. 2024 Jun 10: 101000. Online ahead of print. |
| 1681 | *Camellia crapnelliana* *(Theaceae)* | 2024-06-07 | Zhang F et al., Chromosome-scale genome assembly of oil-tea tree Camellia crapnelliana. Sci Data. 11, 599 (2024). |
| 1680 | *Deinandra increscens* *(Asteraceae)* | 2024-06-07 | McEvoy SL et al., The reference genome of an endangered Asteraceae, Deinandra increscens subsp. villosa, endemic to the Central Coast of California. G3 (Bethesda). 2024 Jun 07: jkae117. Online ahead of print. |
| 1679 | *Ancylonema nordenskioeldii* *(Mesotaeniaceae)* | 2024-06-06 | Bowles AMC et al., Metagenome-assembled genome of the glacier alga Ancylonema yields insights into the evolution of streptophyte life on ice and land. New Phytol. 2024 Jun 6. Online ahead of print. |
| 1678 | *Ehretia macrophylla* *(Ehretiaceae)* | 2024-06-05 | Cheng S et al., Haplotype-resolved chromosome-level genome assembly of Ehretia macrophylla. Sci Data. 11, 589 (2024) |
| 1677 | *Theobroma grandiflorum* *(Malvaceae)* | 2024-06-05 | Alves RM et al., Genomic decoding of Theobroma grandiflorum (cupuassu) at chromosomal scale: evolutionary insights for horticultural innovation. Gigascience. 2024 Jan 2: 13: giae027. |
| 1676 | *Platanus x hispanica* *(Platanaceae)* | 2024-06-03 | Yan X et al., Genome evolution of the ancient hexaploid Platanus x acerifolia (London planetree). Proc Natl Acad Sci U S A. 2024 Jun 11, 121(24): e2319679121. Epub 2024 Jun 3. |
| 1675 | *Syringa pinnatifolia* *(Oleaceae)* | 2024-06-02 | Gao J et al., A Syringa pinnatifolia genome assembly reveals the zerumbone biosynthesis machinery with a cytochrome P450-catalysed epoxidation reaction in eudicots. Plant Biotechnol J. 2024 Jun 2. Online ahead of print. |
| 1674 | *Carex breviculmis* *(Cyperaceae)* | 2024-06-01 | Yuan T et al., The genome assembly of Carex breviculmis provides evidence for its phylogenetic localization and environmental adaptation. Ann Bot. 2024 Jun 1: mcae085. Online ahead of print. |
| 1673 | *Pseudocydonia sinensis* *(Rosaceae)* | 2024-05-31 | Yang Y et al., A chromosome-level genome assembly of Chinese quince (Pseudocydonia sinensis). Front Plant Sci. 2024 May 31: 15: 1368861. eCollection 2024. |
| 1672 | *Warburgia ugandensis* *(Canellaceae)* | 2024-05-30 | Luo L et al., The chromosome-level genomes of the herbal magnoliids Warburgia ugandensis and Saururus chinensis. Sci Data. 2024 May 30, 11(1): 554. |
| 1671 | *Magnolia x alba* *(Magnoliaceae)* | 2024-05-29 | Jiang S et al., A high-quality haplotype genome of Michelia alba DC reveals differences in methylation patterns and flower characteristics. Mol Hortic. 2024 May 29, 4(1): 23. |
| 1670 | *Malus honanensis x domestica* *(Rosaceae)* | 2024-05-29 | Li J et al., The chromosome-level genome assembly of the dwarfing apple interstock Malus hybrid SH6. Sci Data. 2024 May 29, 11(1): 552. |

| | plant species | published | |
|---|---|---|---|
| 1669 | *Physaria fendleri* (Brassicaceae) | 2024-05-28 | Johnston CR et al., First draft reference genome and annotation of the alternative oil species Physaria fendleri. G3 (Bethesda). 2024 May 28: jkae114. Online ahead of print. |
| 1668 | *Apatococcus fuscideae* (Chlorellaceae) | 2024-05-24 | Puginier C et al., Phylogenomics reveals the evolutionary origins of lichenization in chlorophyte algae. Nat Commun. 2024 May 24, 15(1): 4452. |
| 1667 | *Apatococcus lobatus* (Chlorellaceae) | 2024-05-24 | Puginier C et al., Phylogenomics reveals the evolutionary origins of lichenization in chlorophyte algae. Nat Commun. 2024 May 24, 15(1): 4452. |
| 1666 | *Citrus glauca* (Rutaceae) | 2024-05-24 | Singh K et al., Chromosome-Scale, De Novo, Phased Genome Assemblies of Three Australian Limes: Citrus australasica, C. inodora, and C. glauca. Plants (Basel). 2024 May 24, 13(11): 1460. |
| 1665 | *Citrus inodora* (Rutaceae) | 2024-05-24 | Singh K et al., Chromosome-Scale, De Novo, Phased Genome Assemblies of Three Australian Limes: Citrus australasica, C. inodora, and C. glauca. Plants (Basel). 2024 May 24, 13(11): 1460. |
| 1664 | *Coccomyxa pringsheimii* (Coccomyxaceae) | 2024-05-24 | Puginier C et al., Phylogenomics reveals the evolutionary origins of lichenization in chlorophyte algae. Nat Commun. 2024 May 24, 15(1): 4452. |
| 1663 | *Elliptochloris bilobata* (P.incertae sedis) | 2024-05-24 | Puginier C et al., Phylogenomics reveals the evolutionary origins of lichenization in chlorophyte algae. Nat Commun. 2024 May 24, 15(1): 4452. |
| 1662 | *Myrmecia bisecta* (Trebouxiaceae) | 2024-05-24 | Puginier C et al., Phylogenomics reveals the evolutionary origins of lichenization in chlorophyte algae. Nat Commun. 2024 May 24, 15(1): 4452. |
| 1661 | *Symbiochloris irregularis* (Trebouxiaceae) | 2024-05-24 | Puginier C et al., Phylogenomics reveals the evolutionary origins of lichenization in chlorophyte algae. Nat Commun. 2024 May 24, 15(1): 4452. |
| 1660 | *Trebouxia sp.* (Trebouxiaceae) | 2024-05-24 | Puginier C et al., Phylogenomics reveals the evolutionary origins of lichenization in chlorophyte algae. Nat Commun. 2024 May 24, 15(1): 4452. |
| 1659 | *Cicer bijugum* (Fabaceae) | 2024-05-23 | Khan AW et al., Cicer super-pangenome provides insights into species evolution and agronomic trait loci for crop improvement in chickpea. Nat Genet. 2024 May 23. Online ahead of print. |
| 1658 | *Cicer chorassanicum* (Fabaceae) | 2024-05-23 | Khan AW et al., Cicer super-pangenome provides insights into species evolution and agronomic trait loci for crop improvement in chickpea. Nat Genet. 2024 May 23. Online ahead of print. |
| 1657 | *Cicer cuneatum* (Fabaceae) | 2024-05-23 | Khan AW et al., Cicer super-pangenome provides insights into species evolution and agronomic trait loci for crop improvement in chickpea. Nat Genet. 2024 May 23. Online ahead of print. |
| 1656 | *Cicer echinospermum* (Fabaceae) | 2024-05-23 | Khan AW et al., Cicer super-pangenome provides insights into species evolution and agronomic trait loci for crop improvement in chickpea. Nat Genet. 2024 May 23. Online ahead of print. |
| 1655 | *Cicer judaicum* (Fabaceae) | 2024-05-23 | Khan AW et al., Cicer super-pangenome provides insights into species evolution and agronomic trait loci for crop improvement in chickpea. Nat Genet. 2024 May 23. Online ahead of print. |
| 1654 | *Cicer pinnatifidum* (Fabaceae) | 2024-05-23 | Khan AW et al., Cicer super-pangenome provides insights into species evolution and agronomic trait loci for crop improvement in chickpea. Nat Genet. 2024 May 23. Online ahead of print. |
| 1653 | *Cicer yamashitae* (Fabaceae) | 2024-05-23 | Khan AW et al., Cicer super-pangenome provides insights into species evolution and agronomic trait loci for crop improvement in chickpea. Nat Genet. 2024 May 23. Online ahead of print. |
| 1652 | *Ottelia alismoides* (Hydrocharitaceae) | 2024-05-23 | Wang ZF et al., Genome assembly of Ottelia alismoides, a multiple-carbon utilisation aquatic plant. BMC Genom Data. 2024 May 23, 25(1): 48. |
| 1651 | *Physcomitrellopsis africana* (Funariaceae) | 2024-05-23 | Vuruputoor VS et al., Crossroads of assembling a moss genome: navigating contaminants and horizontal gene transfer in the moss Physcomitrellopsis africana. G3 (Bethesda). 2024 May 23: jkae104. Online ahead of print. |
| 1650 | *Ficus simplicissima* (Moraceae) | 2024-05-22 | Huang W et al., A high-quality chromosome-level genome assembly of Ficus hirta. Sci Data. 2024 May 22, 11(1): 526. |
| 1649 | *Camellia granthamiana* (Theaceae) | 2024-05-20 | Hong Kong Biodiversity Genomics Consortium et al., Genome assembly of the rare and endangered Grantham's camellia, Camellia granthamiana. GigaByte. 2024 May 20, 2024: gigabyte124. eCollection 2024. |
| 1648 | *Capsicum rhomboideum* (Solanaceae) | 2024-05-20 | Chen W et al., Two telomere-to-telomere gapless genomes reveal insights into Capsicum evolution and capsaicinoid biosynthesis. Nat Commun. 15, 4295 (2024). |
| 1647 | *Dasiphora fruticosa* (Rosaceae) | 2024-05-20 | Yang FS et al., Signatures of adaptation and purifying selection in highland populations of Dasiphora fruticosa. Mol Biol Evol. 2024 May 20: msae099. Online ahead of print. |
| 1646 | *Erythranthe nasuta* (Phrymaceae) | 2024-05-20 | Whitener MR et al., Patterns of genomic variation reveal a single evolutionary origin of the wild allotetraploid Mimulus sookensis. Evolution. 2024 May 20: qpae079. Online ahead of print. |
| 1645 | *Salicornia bigelovii* (Amaranthaceae) | 2024-05-20 | Salazar OR et al., SOS1 tonoplast neo-localization and the RGG protein SALTY are important in the extreme salinity tolerance of Salicornia bigelovii. Nat Commun. 2024 May 20, 15(1): 4279. |
| 1644 | *Vernicia montana* (Euphorbiaceae) | 2024-05-18 | Li W et al., Genome assembly and resequencing shed light on evolution, population selection, and sex identification in Vernicia montana. Hortic Res. 2024 May: uhae141. eCollection 2024 May. |
| 1643 | *Lespedeza daurica* (Fabaceae) | 2024-05-17 | Yan Q et al., The genome of Lespedeza potaninii reveals biased subgenome evolution and drought adaptation. Plant Physiol. 2024 May 17: kiae283. Online ahead of print. |
| 1642 | *Salicornia europaea* (Amaranthaceae) | 2024-05-15 | Mian S et al., The genome sequence of purple glasswort, Salicornia ramosissima Woods (Amaranthaceae). Wellcome Open Res. 2024 May 15: 9: 257. eCollection 2024. |
| 1641 | *Acacia confusa* (Fabaceae) | 2024-05-09 | Chen R et al., The genomes of seven economic Caesalpinioideae trees provide insights into polyploidization history and secondary metabolite biosynthesis. Plant Commun. 2024 May 9: 100944. Online ahead of print. |
| 1640 | *Albizia julibrissin* (Fabaceae) | 2024-05-09 | Chen R et al., The genomes of seven economic Caesalpinioideae trees provide insights into polyploidization history and secondary metabolite biosynthesis. Plant Commun. 2024 May 9: 100944. Online ahead of print. |
| 1639 | *Biancaea sappan* (Fabaceae) | 2024-05-09 | Chen R et al., The genomes of seven economic Caesalpinioideae trees provide insights into polyploidization history and secondary metabolite biosynthesis. Plant Commun. 2024 May 9: 100944. Online ahead of print. |
| 1638 | *Delonix regia* (Fabaceae) | 2024-05-09 | Chen R et al., The genomes of seven economic Caesalpinioideae trees provide insights into polyploidization history and secondary metabolite biosynthesis. Plant Commun. 2024 May 9: 100944. Online ahead of print. |
| 1637 | *Gleditsia sinensis* (Fabaceae) | 2024-05-09 | Chen R et al., The genomes of seven economic Caesalpinioideae trees provide insights into polyploidization history and secondary metabolite biosynthesis. Plant Commun. 2024 May 9: 100944. Online ahead of print. |
| 1636 | *Leucaena leucocephala* (Fabaceae) | 2024-05-09 | Chen R et al., The genomes of seven economic Caesalpinioideae trees provide insights into polyploidization history and secondary metabolite biosynthesis. Plant Commun. 2024 May 9: 100944. Online ahead of print. |
| 1635 | *Spinacia tetrandra* (Amaranthaceae) | 2024-05-07 | She H et al., Insights into spinach domestication from genome sequences of two wild spinach progenitors, Spinacia turkestanica and Spinacia tetrandra. New Phytol. 2024 Jul, 243(1): 477-494. Epub 2024 May 7. |
| 1634 | *Linum trigynum* (Linaceae) | 2024-05-06 | Gutierrez-Valencia J et al., Genetic causes and genomic consequences of breakdown of distyly in Linum trigynum. Mol Biol Evol. 2024 May 6: msae087. Online ahead of print. |
| 1633 | *Solanum berthaultii* (Solanaceae) | 2024-05-04 | Achakkagari SR et al., The phased Solanum okadae genome and Petota pangenome analysis of 23 other potato wild relatives and hybrids. Sci Data. 2024 May 4, 11(1): 454. |

| | plant species | published | |
|---|---|---|---|
| 1632 | *Solanum chiquidenum* *(Solanaceae)* | 2024-05-04 | Achakkagari SR et al., The phased Solanum okadae genome and Petota pangenome analysis of 23 other potato wild relatives and hybrids. Sci Data. 2024 May 4, 11(1): 454. |
| 1631 | *Solanum gandarillasii* *(Solanaceae)* | 2024-05-04 | Achakkagari SR et al., The phased Solanum okadae genome and Petota pangenome analysis of 23 other potato wild relatives and hybrids. Sci Data. 2024 May 4, 11(1): 454. |
| 1630 | *Solanum gracilifrons* *(Solanaceae)* | 2024-05-04 | Achakkagari SR et al., The phased Solanum okadae genome and Petota pangenome analysis of 23 other potato wild relatives and hybrids. Sci Data. 2024 May 4, 11(1): 454. |
| 1629 | *Solanum medians* *(Solanaceae)* | 2024-05-04 | Achakkagari SR et al., The phased Solanum okadae genome and Petota pangenome analysis of 23 other potato wild relatives and hybrids. Sci Data. 2024 May 4, 11(1): 454. |
| 1628 | *Solanum okadae* *(Solanaceae)* | 2024-05-04 | Achakkagari SR et al., The phased Solanum okadae genome and Petota pangenome analysis of 23 other potato wild relatives and hybrids. Sci Data. 2024 May 4, 11(1): 454. |
| 1627 | *Solanum tarnii x tuberosum* *(Solanaceae)* | 2024-05-04 | Achakkagari SR et al., The phased Solanum okadae genome and Petota pangenome analysis of 23 other potato wild relatives and hybrids. Sci Data. 2024 May 4, 11(1): 454. |
| 1626 | *Cinnamomum chago* *(Lauraceae)* | 2024-05-03 | Tao L et al., Chromosome-level genome assembly of the threatened resource plant Cinnamomum chago. Sci Data. 2024 May 3, 11(1): 447. |
| 1625 | *Pedicularis kansuensis* *(Orobanchaceae)* | 2024-05-02 | Fang L et al., Chromosome-level genome assembly of Pedicularis kansuensis illuminates genome evolution of facultative parasitic plant. Mol Ecol Resour. 2024 May 2: e13966. Online ahead of print. |
| 1624 | *Barthea barthei* *(Melastomataceae)* | 2024-05-01 | Huang W et al., De novo genome assembly and population genomics of a shrub tree Barthea barthei (Hance) krass provide insights into the adaptive color variations. Front Plant Sci. 2024 May 1: 15: 1365686. eCollection 2024. |
| 1623 | *Triplostegia glandulifera* *(Caprifoliaceae)* | 2024-05-01 | Zhang J et al., Coping with alpine habitats: genomic insights into the adaptation strategies of Triplostegia glandulifera (Caprifoliaceae). Hortic Res. 2024 May 1, 11(5): uhae077. eCollection 2024 May. |
| 1622 | *Zygnema cf. cylindricum* *(Zygnemataceae)* | 2024-05-01 | Feng X et al., Genomes of multicellular algal sisters to land plants illuminate signaling network evolution. Nat Genet. 2024 May 1. Online ahead of print. |
| 1621 | *Zygnema circumcarinatum* *(Zygnemataceae)* | 2024-05-01 | Feng X et al., Genomes of multicellular algal sisters to land plants illuminate signaling network evolution. Nat Genet. 2024 May 1. Online ahead of print. |
| 1620 | *Tillandsia fasciculata* *(Bromeliaceae)* | 2024-04-30 | Crego CG et al., CAM evolution is associated with gene family expansion in an explosive bromeliad radiation. Plant Cell. 2024 Apr 30: koae130. Online ahead of print. |
| 1619 | *Tillandsia leiboldiana* *(Bromeliaceae)* | 2024-04-30 | Crego CG et al., CAM evolution is associated with gene family expansion in an explosive bromeliad radiation. Plant Cell. 2024 Apr 30: koae130. Online ahead of print. |
| 1618 | *Excoecaria agallocha* *(Euphorbiaceae)* | 2024-04-25 | Hong Kong Biodiversity Genomics Consortium et al., Genome assembly of the milky mangrove Excoecaria agallocha. GigaByte. 2024 Apr 25, 2024: gigabyte119. eCollection 2024. |
| 1617 | *Limnomonas spitsbergensis* *(C.incertae sedis)* | 2024-04-25 | Hulatt CJ et al., The genome of the Arctic snow-alga Limnomonas spitsbergensis (Chlamydomonadales). G3 (Bethesda). 2024 Apr 25: jkae086. Online ahead of print. |
| 1616 | *Populus nigra* *(Salicaceae)* | 2024-04-24 | Christenhusz MJM et al., The genome sequence of black poplar, Populus nigra subsp. betulifolia L., 1753 (Salicaceae). Wellcome Open Res. 2024 Apr 24: 9: 228. eCollection 2024. |
| 1615 | *Populus deltoides x canadensis* *(Salicaceae)* | 2024-04-23 | Luo J et al., Haplotype-resolved genome assembly of poplar line NL895 provides a valuable tree genomic resource. For Res (Fayettev). 2024 Apr 23, 4: e015. eCollection 2024. |
| 1614 | *Triticum timopheevii* *(Poaceae)* | 2024-04-23 | Grewal S et al., Chromosome-scale genome assembly of bread wheat wild relative Triticum timopheevii. Sci Data 11, 420 (2024). |
| 1613 | *Cotoneaster glaucophyllus* *(Rosaceae)* | 2024-04-22 | Meng K et al., Chromosome-scale genome assembly and annotation of Cotoneaster glaucophyllus. Sci Data 11, 406 (2024). |
| 1612 | *Bryopsis sp.* *(Bryopsidaceae)* | 2024-04-20 | Ochiai KK et al., Genome sequence and cell biological toolbox of the highly regenerative, coenocytic green feather alga Bryopsis. Plant J. 2024 Apr 20. Online ahead of print. |
| 1611 | *Rorippa aquatica* *(Brassicaceae)* | 2024-04-18 | Sakamoto T et al., A chromosome-level genome assembly for the amphibious plant Rorippa aquatica reveals its allotetraploid origin and mechanisms of heterophylly upon submergence. Commun Biol. 7, 431 (2024). |
| 1610 | *Aristolochia californica* *(Aristolochiaceae)* | 2024-04-15 | Chaturvedi S et al., A draft reference genome assembly of California Pipevine, Aristolochia californica Torr. J Hered. 2024 Apr 15: esae023. Online ahead of print. |
| 1609 | *Coffea eugenioides* *(Rubiaceae)* | 2024-04-15 | Salojarvi J et al., The genome and population genomics of allopolyploid Coffea arabica reveal the diversification history of modern coffee cultivars. Nat Genet. 2024 Apr, 56(4): 721-731. Epub 2024 Apr 15. |
| 1608 | *Breynia spatulifolia* *(Phyllanthaceae)* | 2024-04-12 | Li F et al., Haplotype-resolved genomes of octoploid species in Phyllanthaceae family reveal a critical role for polyploidization and hybridization in speciation. Plant J. 2024 Apr 12. Online ahead of print. |
| 1607 | *Hippophae tibetana* *(Elaeagnaceae)* | 2024-04-12 | Zhang G et al., Chromosome-level genome assembly of Hippophae tibetana provides insights into high-altitude adaptation and flavonoid biosynthesis. BMC Biol. 2024 Apr 12, 22(1): 82. |
| 1606 | *Phillyrea angustifolia* *(Oleaceae)* | 2024-04-11 | Castric V et al., The homomorphic self-incompatibility system in Oleaceae is controlled by a hemizygous genomic region expressing a gibberellin pathway gene. Curr Biol. 2024 Apr 11: S0960-9822(24)00387-7. Online ahead of print. |
| 1605 | *Scleromitrion diffusum* *(Rubiaceae)* | 2024-04-11 | Gao Y et al., Telomere-to-telomere genome assembly of Oldenlandia diffusa. DNA Res. 2024 Apr 11: dsae012. Online ahead of print. |
| 1604 | *Rheum palmatum* *(Polygonaceae)* | 2024-04-10 | Zhang T et al., A chromosome-level genome reveals genome evolution and molecular basis of anthraquinone biosynthesis in Rheum palmatum. BMC Plant Biol. 2024 Apr 10, 24(1): 261. |
| 1603 | *Stenogyne calaminthoides* *(Lamiaceae)* | 2024-04-10 | Tomlin CM et al., Allopolyploid origin and diversification of the Hawaiian endemic mints. Nat Commun. 2024 Apr 10, 15(1): 3109. |
| 1602 | *Caragana lidou* *(Fabaceae)* | 2024-04-09 | Cao J et al., Chromosome-level genome assembly and characterization of the Calophaca sinica genome. DNA Res. 2024 Apr 9: dsae011. Online ahead of print. |
| 1601 | *Saposhnikovia divaricata* *(Apiaceae)* | 2024-04-09 | Wang ZH et al., Genomic, transcriptomic, and metabolomic analyses provide insights into the evolution and development of a medicinal plant Saposhnikovia divaricata (Apiaceae). Hortic Res. 2024 Apr: uhae105. eCollection 2024 Apr. |
| 1600 | *Angophora floribunda* *(Myrtaceae)* | 2024-04-08 | Ferguson S et al., Plant genome evolution in the genus Eucalyptus driven by structural rearrangements that promote sequence divergence. Genome Res. 2024 Apr 8: gr.277999.123. Online ahead of print. |
| 1599 | *Corymbia maculata* *(Myrtaceae)* | 2024-04-08 | Ferguson S et al., Plant genome evolution in the genus Eucalyptus driven by structural rearrangements that promote sequence divergence. Genome Res. 2024 Apr 8: gr.277999.123. Online ahead of print. |
| 1598 | *Eucalyptus brandiana* *(Myrtaceae)* | 2024-04-08 | Ferguson S et al., Plant genome evolution in the genus Eucalyptus driven by structural rearrangements that promote sequence divergence. Genome Res. 2024 Apr 8: gr.277999.123. Online ahead of print. |
| 1597 | *Eucalyptus caleyi* *(Myrtaceae)* | 2024-04-08 | Ferguson S et al., Plant genome evolution in the genus Eucalyptus driven by structural rearrangements that promote sequence divergence. Genome Res. 2024 Apr 8: gr.277999.123. Online ahead of print. |
| 1596 | *Eucalyptus cladocalyx* *(Myrtaceae)* | 2024-04-08 | Ferguson S et al., Plant genome evolution in the genus Eucalyptus driven by structural rearrangements that promote sequence divergence. Genome Res. 2024 Apr 8: gr.277999.123. Online ahead of print. |

| | plant species | published | |
|---|---|---|---|
| 1595 | *Eucalyptus cloeziana* (Myrtaceae) | 2024-04-08 | Ferguson S et al., Plant genome evolution in the genus Eucalyptus driven by structural rearrangements that promote sequence divergence. Genome Res. 2024 Apr 8: gr.277999.123. Online ahead of print. |
| 1594 | *Eucalyptus coolabah* (Myrtaceae) | 2024-04-08 | Ferguson S et al., Plant genome evolution in the genus Eucalyptus driven by structural rearrangements that promote sequence divergence. Genome Res. 2024 Apr 8: gr.277999.123. Online ahead of print. |
| 1593 | *Eucalyptus curtisii* (Myrtaceae) | 2024-04-08 | Ferguson S et al., Plant genome evolution in the genus Eucalyptus driven by structural rearrangements that promote sequence divergence. Genome Res. 2024 Apr 8: gr.277999.123. Online ahead of print. |
| 1592 | *Eucalyptus dawsonii* (Myrtaceae) | 2024-04-08 | Ferguson S et al., Plant genome evolution in the genus Eucalyptus driven by structural rearrangements that promote sequence divergence. Genome Res. 2024 Apr 8: gr.277999.123. Online ahead of print. |
| 1591 | *Eucalyptus decipiens* (Myrtaceae) | 2024-04-08 | Ferguson S et al., Plant genome evolution in the genus Eucalyptus driven by structural rearrangements that promote sequence divergence. Genome Res. 2024 Apr 8: gr.277999.123. Online ahead of print. |
| 1590 | *Eucalyptus erythrocorys* (Myrtaceae) | 2024-04-08 | Ferguson S et al., Plant genome evolution in the genus Eucalyptus driven by structural rearrangements that promote sequence divergence. Genome Res. 2024 Apr 8: gr.277999.123. Online ahead of print. |
| 1589 | *Eucalyptus fibrosa* (Myrtaceae) | 2024-04-08 | Ferguson S et al., Plant genome evolution in the genus Eucalyptus driven by structural rearrangements that promote sequence divergence. Genome Res. 2024 Apr 8: gr.277999.123. Online ahead of print. |
| 1588 | *Eucalyptus globulus* (Myrtaceae) | 2024-04-08 | Ferguson S et al., Plant genome evolution in the genus Eucalyptus driven by structural rearrangements that promote sequence divergence. Genome Res. 2024 Apr 8: gr.277999.123. Online ahead of print. |
| 1587 | *Eucalyptus guilfoylei* (Myrtaceae) | 2024-04-08 | Ferguson S et al., Plant genome evolution in the genus Eucalyptus driven by structural rearrangements that promote sequence divergence. Genome Res. 2024 Apr 8: gr.277999.123. Online ahead of print. |
| 1586 | *Eucalyptus lansdowneana* (Myrtaceae) | 2024-04-08 | Ferguson S et al., Plant genome evolution in the genus Eucalyptus driven by structural rearrangements that promote sequence divergence. Genome Res. 2024 Apr 8: gr.277999.123. Online ahead of print. |
| 1585 | *Eucalyptus leucophloia* (Myrtaceae) | 2024-04-08 | Ferguson S et al., Plant genome evolution in the genus Eucalyptus driven by structural rearrangements that promote sequence divergence. Genome Res. 2024 Apr 8: gr.277999.123. Online ahead of print. |
| 1584 | *Eucalyptus marginata* (Myrtaceae) | 2024-04-08 | Ferguson S et al., Plant genome evolution in the genus Eucalyptus driven by structural rearrangements that promote sequence divergence. Genome Res. 2024 Apr 8: gr.277999.123. Online ahead of print. |
| 1583 | *Eucalyptus melliodora x sideroxylon* (Myrtaceae) | 2024-04-08 | Ferguson S et al., Plant genome evolution in the genus Eucalyptus driven by structural rearrangements that promote sequence divergence. Genome Res. 2024 Apr 8: gr.277999.123. Online ahead of print. |
| 1582 | *Eucalyptus microcorys* (Myrtaceae) | 2024-04-08 | Ferguson S et al., Plant genome evolution in the genus Eucalyptus driven by structural rearrangements that promote sequence divergence. Genome Res. 2024 Apr 8: gr.277999.123. Online ahead of print. |
| 1581 | *Eucalyptus paniculata* (Myrtaceae) | 2024-04-08 | Ferguson S et al., Plant genome evolution in the genus Eucalyptus driven by structural rearrangements that promote sequence divergence. Genome Res. 2024 Apr 8: gr.277999.123. Online ahead of print. |
| 1580 | *Eucalyptus polyanthemos* (Myrtaceae) | 2024-04-08 | Ferguson S et al., Plant genome evolution in the genus Eucalyptus driven by structural rearrangements that promote sequence divergence. Genome Res. 2024 Apr 8: gr.277999.123. Online ahead of print. |
| 1579 | *Eucalyptus pumila* (Myrtaceae) | 2024-04-08 | Ferguson S et al., Plant genome evolution in the genus Eucalyptus driven by structural rearrangements that promote sequence divergence. Genome Res. 2024 Apr 8: gr.277999.123. Online ahead of print. |
| 1578 | *Eucalyptus regnans* (Myrtaceae) | 2024-04-08 | Ferguson S et al., Plant genome evolution in the genus Eucalyptus driven by structural rearrangements that promote sequence divergence. Genome Res. 2024 Apr 8: gr.277999.123. Online ahead of print. |
| 1577 | *Eucalyptus shirleyi* (Myrtaceae) | 2024-04-08 | Ferguson S et al., Plant genome evolution in the genus Eucalyptus driven by structural rearrangements that promote sequence divergence. Genome Res. 2024 Apr 8: gr.277999.123. Online ahead of print. |
| 1576 | *Eucalyptus tenuipes* (Myrtaceae) | 2024-04-08 | Ferguson S et al., Plant genome evolution in the genus Eucalyptus driven by structural rearrangements that promote sequence divergence. Genome Res. 2024 Apr 8: gr.277999.123. Online ahead of print. |
| 1575 | *Eucalyptus victrix* (Myrtaceae) | 2024-04-08 | Ferguson S et al., Plant genome evolution in the genus Eucalyptus driven by structural rearrangements that promote sequence divergence. Genome Res. 2024 Apr 8: gr.277999.123. Online ahead of print. |
| 1574 | *Eucalyptus viminalis* (Myrtaceae) | 2024-04-08 | Ferguson S et al., Plant genome evolution in the genus Eucalyptus driven by structural rearrangements that promote sequence divergence. Genome Res. 2024 Apr 8: gr.277999.123. Online ahead of print. |
| 1573 | *Eucalyptus virginea* (Myrtaceae) | 2024-04-08 | Ferguson S et al., Plant genome evolution in the genus Eucalyptus driven by structural rearrangements that promote sequence divergence. Genome Res. 2024 Apr 8: gr.277999.123. Online ahead of print. |
| 1572 | *Silybum marianum* (Asteraceae) | 2024-04-05 | Kim KD et al., Chromosome-level genome assembly of milk thistle (Silybum marianum (L.) Gaertn.). Sci Data 11, 342 (2024). |
| 1571 | *Mimosa bimucronata* (Fabaceae) | 2024-04-03 | Jia H et al., Haplotype-resolved genome of Mimosa bimucronata revealed insights into leaf movement and nitrogen fixation. BMC Genomics. 2024 Apr 3, 25(1): 334. |
| 1570 | *Corylus chinensis* (Betulaceae) | 2024-04-02 | Yang Z et al., Conservation genomics provides insights into genetic resilience and adaptation of the endangered Chinese hazelnut, Corylus chinensis. Plant Divers. 2024 Apr 2, 46(3): 294-308. eCollection 2024 May. |
| 1569 | *Quercus rex* (Fagaceae) | 2024-04-02 | Liu X et al., Chromosome-scale genomes of Quercus sichourensis and Quercus rex provide insights into the evolution and adaptation of Fagaceae. J Genet Genomics. 2024 Apr 2: S1673-8527(24)00065-1. Online ahead of print. |
| 1568 | *Quercus sichourensis* (Fagaceae) | 2024-04-02 | Liu X et al., Chromosome-scale genomes of Quercus sichourensis and Quercus rex provide insights into the evolution and adaptation of Fagaceae. J Genet Genomics. 2024 Apr 2: S1673-8527(24)00065-1. Online ahead of print. |
| 1567 | *Ammopiptanthus mongolicus* (Fabaceae) | 2024-04-01 | Feng L et al., A reference-grade genome of the xerophyte Ammopiptanthus mongolicus sheds light on its evolution history in legumes and drought tolerance mechanisms. Plant Commun. 2024 Apr 1:100891. Online ahead of print. |
| 1566 | *Grona styracifolia* (Fabaceae) | 2024-03-30 | Zeng S et al., The high-quality genome of Grona styracifolia uncover the genomic mechanism of high level of schaftoside, a promising drug candidate for treatment of COVID-19. Hortic Res. 2024 Mar: uhae089. eCollection 2024 Mar. |
| 1565 | *Pachypodium lamerei* (Apocynaceae) | 2024-03-30 | Cuello C et al., The Madagascar palm genome provides new insights on the evolution of Apocynaceae specialized metabolism. Heliyon. 2024 Mar 14, 10(6): e28078. eCollection 2024 Mar 30. |
| 1564 | *Alstonia scholaris* (Apocynaceae) | 2024-03-27 | Chen H et al., Chromosome-level Alstonia scholaris genome unveils evolutionary insights into biosynthesis of monoterpenoid indole alkaloids. iScience. 2024 Mar 27, 27(5): 109599. eCollection 2024 May 17. |
| 1563 | *Pinus albicaulis* (Pinaceae) | 2024-03-25 | Neale DB et al., A Genome Sequence for the Threatened Whitebark Pine. G3 (Bethesda). 2024 Mar 25: jkae061. Online ahead of print. |
| 1562 | *Epilobium angustifolium* (Onagraceae) | 2024-03-20 | Christenhusz MJM et al., The genome sequence of rosebay willowherb Chamaenerion angustifolium (L.) Scop., 1771 (syn. Epilobium angustifolium L., 1753) (Onagraceae). Wellcome Open Res. 2024 Mar 20: 9: 163. eCollection 2024. |
| 1561 | *Scutellaria minor* (Lamiaceae) | 2024-03-20 | Mian S et al., The genome sequence of the Lesser Skullcap, Scutellaria minor Huds., 1762 (Lamiaceae). Wellcome Open Res. 2024 Mar 20: 9: 165. eCollection 2024. |
| 1560 | *Ampelocalamus luodianensis* (Poaceae) | 2024-03-15 | Ma PF et al., Genome assemblies of 11 bamboo species highlight diversification induced by dynamic subgenome dominance. Nat Genet. 2024 Mar 15. Online ahead of print. |
| 1559 | *Dendrocalamus sinicus* (Poaceae) | 2024-03-15 | Ma PF et al., Genome assemblies of 11 bamboo species highlight diversification induced by dynamic subgenome dominance. Nat Genet. 2024 Mar 15. Online ahead of print. |

| | plant species | published | |
|---|---|---|---|
| 1558 | *Hsuehochloa calcarea* *(Poaceae)* | 2024-03-15 | Ma PF et al., Genome assemblies of 11 bamboo species highlight diversification induced by dynamic subgenome dominance. Nat Genet. 2024 Mar 15. Online ahead of print. |
| 1557 | *Melocanna baccifera* *(Poaceae)* | 2024-03-15 | Ma PF et al., Genome assemblies of 11 bamboo species highlight diversification induced by dynamic subgenome dominance. Nat Genet. 2024 Mar 15. Online ahead of print. |
| 1556 | *Otatea glauca* *(Poaceae)* | 2024-03-15 | Ma PF et al., Genome assemblies of 11 bamboo species highlight diversification induced by dynamic subgenome dominance. Nat Genet. 2024 Mar 15. Online ahead of print. |
| 1555 | *Rhipidocladum racemiflorum* *(Poaceae)* | 2024-03-15 | Ma PF et al., Genome assemblies of 11 bamboo species highlight diversification induced by dynamic subgenome dominance. Nat Genet. 2024 Mar 15. Online ahead of print. |
| 1554 | *Teucrium marum* *(Lamiaceae)* | 2024-03-15 | Smit SJ et al., The genomic and enzymatic basis for iridoid biosynthesis in cat thyme (Teucrium marum). Plant J. 2024 Mar 15. Online ahead of print. |
| 1553 | *Musa ornata* *(Musaceae)* | 2024-03-14 | Xiao TW et al., Chromosome-level genome assemblies of Musa ornata and M. velutina provide insights into pericarp dehiscence and anthocyanin biosynthesis in banana. Hortic Res. 2024 Mar 14: uhae079. |
| 1552 | *Musa velutina* *(Musaceae)* | 2024-03-14 | Xiao TW et al., Chromosome-level genome assemblies of Musa ornata and M. velutina provide insights into pericarp dehiscence and anthocyanin biosynthesis in banana. Hortic Res. 2024 Mar 14: uhae079. |
| 1551 | *Pontederia crassipes* *(Pontederiaceae)* | 2024-03-14 | Huang Y et al., A reference genome of Commelinales provides insights into the commelinids evolution and global spread of water hyacinth (Pontederia crassipes). Gigascience. 2024 Jan 2: 13: giae006. |
| 1550 | *Torenia fournieri* *(Linderniaceae)* | 2024-03-14 | Song J et al., Genome assembly and multi-omic analyses reveal the mechanisms underlying flower color formation in Torenia fournieri . Plant Genome. 2024 Mar 14: e20439. Online ahead of print. |
| 1549 | *Menispermum dauricum* *(Menispermaceae)* | 2024-03-13 | An Z et al., Lineage-Specific CYP80 Expansion and Benzylisoquinoline Alkaloid Diversity in Early-Diverging Eudicots. Adv Sci (Weinh). 2024 Mar 13: e2309990. Online ahead of print. |
| 1548 | *Poa trivialis* *(Poaceae)* | 2024-03-13 | Brunharo CACG et al., Chromosome-scale genome assembly of Poa trivialis and population genomics reveal widespread gene flow in a cool-season grass seed production system. Plant Direct. 2024 Mar 13, 8(3):e575. eCollection 2024 Mar. |
| 1547 | *Thymus mandschuricus* *(Lamiaceae)* | 2024-03-13 | Jia L et al., Chromosome-level genome of Thymus mandschuricus reveals molecular mechanism of aroma compounds biosynthesis. Front Plant Sci. 2024 Mar 13: 15: 1368869 . eCollection 2024. |
| 1546 | *Clarkia xantiana* *(Onagraceae)* | 2024-03-11 | Sianta SA et al., The extent of introgression between incipient Clarkia species is determined by temporal environmental variation and mating system. Proc Natl Acad Sci U S A. 2024 Mar 19, 121(12): e2316008121. Epub 2024 Mar 11. |
| 1545 | *Musella lasiocarpa* *(Musaceae)* | 2024-03-08 | Zhao W et al., Characterization of O-Methyltransferases in the Biosynthesis of Phenylphenalenone Phytoalexins Based on the Telomere-to-Telomere Gap-less Genome of Musella lasiocarpa. Hortic Res. 2024 Mar 08: uhae042. |
| 1544 | *Bolboschoenus planiculmis* *(Cyperaceae)* | 2024-03-06 | Ning Y et al., Chromosome-scale genome assembly for clubrush (Bolboschoenus planiculmis) indicates a karyotype with high chromosome number and heterogeneous centromere distribution. Genome Biol Evol. 2024 Mar 6: evae039. Online ahead of print. |
| 1543 | *Idesia polycarpa* *(Salicaceae)* | 2024-03-06 | Zuo Y et al., The Idesia polycarpa genome provides insights into its evolution and oil biosynthesis. Cell Rep. 2024 Mar 6, 43(3): 113909. Online ahead of print. |
| 1542 | *Pseudoroegneria libanotica* *(Poaceae)* | 2024-03-06 | Zhai X et al., A chromosome level genome assembly of Pseudoroegneria libanotica reveals a key kcs gene involves in the cuticular wax elongation for drought resistance. BMC Genomics. 2024 Mar 06, 25(1): 253. |
| 1541 | *Calotropis procera* *(Apocynaceae)* | 2024-03-05 | Gaafar ARZ et al., First draft of the nuclear genome assembly and annotation of the multi-stress tolerant desert giant milkweed Calotropis procera. S Afr J Bot. 2024 March, 166: 442-454. Epub 2024 Mar. |
| 1540 | *Haberlea rhodopensis* *(Gesneriaceae)* | 2024-03-05 | Gupta S et al., The genome of Haberlea rhodopensis provides insights into the mechanisms for tolerance to multiple extreme environments. Cell Mol Life Sci. 2024 Mar 5, 81(1): 117. |
| 1539 | *Rubus rosifolius* *(Rosaceae)* | 2024-03-02 | Wang Y et al., Chromosome-scale genome, together with transcriptome and metabolome, provided insights into the evolution and anthocyanin biosynthesis of Rubus rosaefolius Sm. (Rosaceae). Hortic Res. 2024 Mar 02: uhae064. |
| 1538 | *Luzula sylvatica* *(Juncaceae)* | 2024-03-01 | Goodwin ZA et al., The genome sequence of great wood-rush, Luzula sylvatica (Huds) Gaudin. Wellcome Open Res. 2024 Mar 1:9:124. eCollection 2024. |
| 1537 | *Notholithocarpus densiflorus* *(Fagaceae)* | 2024-03-01 | Cai Y et al., Assembly and analysis of the genome of Notholithocarpus densiflorus. G3 (Bethesda). 2024 Mar 1: jkae043. Online ahead of print. |
| 1536 | *Polygonum aviculare* *(Polygonaceae)* | 2024-03-01 | Christenhusz MJM et al., The genome sequence of common knotgrass, Polygonum aviculare L. (Polygonaceae). Wellcome Open Res. 2024 Mar 1: 9: 112. eCollection 2024. |
| 1535 | *Psophocarpus tetragonolobus* *(Fabaceae)* | 2024-03-01 | Ho WK et al., A genomic toolkit for winged bean Psophocarpus tetragonolobus. Nat Commun. 2024 Mar 1, 15(1): 1901. |
| 1534 | *Sherardia arvensis* *(Rubiaceae)* | 2024-03-01 | Christenhusz MJM et al., The genome sequence of field madder, Sherardia arvensis L., 1753 (Rubiaceae). Wellcome Open Res. 2024 Mar 1: 9: 126. eCollection 2024. |
| 1533 | *Baccaurea ramiflora* *(Phyllanthaceae)* | 2024-02-28 | Huang J et al., Genome assembly provides insights into the genome evolution of Baccaurea ramiflora Lour. Sci Rep. 14, 4867 (2024). |
| 1532 | *Syntrichia ruralis* *(Pottiaceae)* | 2024-02-28 | Zhang X et al., Syntrichia ruralis: emerging model moss genome reveals a conserved and previously unknown regulator of desiccation in flowering plants. New Phytol. 2024 Feb 28. Online ahead of print. |
| 1531 | *Camellia japonica* *(Theaceae)* | 2024-02-27 | Hu Z et al., Genomics insights into flowering and floral pattern formation: regional duplication and seasonal pattern of gene expression in Camellia. BMC Biol. 2024 Feb 27, 22(1): 50. |
| 1530 | *Dicorynia guianensis* *(Fabaceae)* | 2024-02-27 | Schmitt S et al., Low-frequency somatic mutations are heritable in tropical trees Dicorynia guianensis and Sextonia rubra. Proc Natl Acad Sci U S A. 2024 Mar 5, 121(10): e2313312121. Epub 2024 Feb 27. |
| 1529 | *Lolium multiflorum* *(Poaceae)* | 2024-02-27 | Goettelmann F et al., High-resolution bulked segregant analysis enables candidate gene identification for bacterial wilt resistance in Italian ryegrass (Lolium multiflorum Lam.) Plant J. 2024 Feb 27. Online ahead of publication |
| 1528 | *Sextonia rubra* *(Lauraceae)* | 2024-02-27 | Schmitt S et al., Low-frequency somatic mutations are heritable in tropical trees Dicorynia guianensis and Sextonia rubra. Proc Natl Acad Sci U S A. 2024 Mar 5, 121(10): e2313312121. Epub 2024 Feb 27. |
| 1527 | *Hopea chinensis* *(Dipterocarpaceae)* | 2024-02-23 | Wang R et al., Dipterocarpoidae genomics reveal their demography and adaptations to Asian rainforests. Nat Commun. 15, 1683 (2024). |
| 1526 | *Hopea reticulata* *(Dipterocarpaceae)* | 2024-02-23 | Wang R et al., Dipterocarpoidae genomics reveal their demography and adaptations to Asian rainforests. Nat Commun. 15, 1683 (2024). |
| 1525 | *Shorea robusta* *(Dipterocarpaceae)* | 2024-02-23 | Wang R et al., Dipterocarpoidae genomics reveal their demography and adaptations to Asian rainforests. Nat Commun. 15, 1683 (2024). |
| 1524 | *Stephania cephalantha* *(Menispermaceae)* | 2024-02-20 | Leng L et al., Cepharanthine analogs mining and genomes of Stephania accelerate anti-coronavirus drug discovery. Nat Commun. 2024 Feb 20, 15(1): 1537. |
| 1523 | *Stephania japonica* *(Menispermaceae)* | 2024-02-20 | Liu Z et al., The genome of Stephania japonica provides insights into the biosynthesis of cepharanthine. Cell Rep. 2024 Feb 20, 43(3): 113832. Online ahead of print. |
| 1522 | *Stephania yunnanensis* *(Menispermaceae)* | 2024-02-20 | Leng L et al., Cepharanthine analogs mining and genomes of Stephania accelerate anti-coronavirus drug discovery. Nat Commun. 2024 Feb 20, 15(1): 1537. |

| | plant species | published | |
|---|---|---|---|
| 1521 | *Spiraea crenata* *(Rosaceae)* | 2024-02-17 | Laczko L et al., The draft genome of Spiraea crenata L. (Rosaceae) - the first complete genome in tribe Spiraeeae. Sci Data. 2024 Feb 17, 11(1): 219. |
| 1520 | *Ceiba pentandra* *(Malvaceae)* | 2024-02-05 | Shao L et al., High-quality genomes of Bombax ceiba and Ceiba pentandra provide insights into evolution of Malvaceae species and differentiation of natural fiber. Plant Commun. 2024 Feb 5:100832. Online ahead of print. |
| 1519 | *Cissus quadrangularis* *(Vitaceae)* | 2024-02-02 | Li Q et al., The Cissus quadrangularis genome reveals its adaptive features in arid habitat. Hortic Res. 2024 Feb: uhae038. eCollection 2024 Feb. |
| 1518 | *Quercus tomentella* *(Fagaceae)* | 2024-02-02 | Mead A et al., The genome assembly of Island Oak (Quercus tomentella), a relictual island tree species. J Hered. 2024 Feb 2: esae002. Online ahead of print. |
| 1517 | *Chrysanthemum indicum* *(Asteraceae)* | 2024-01-31 | Deng Y et al., Genomic insights into the evolution of flavonoid biosynthesis and O-methyltransferase and glucosyltransferase in Chrysanthemum indicum. Cell Rep. 2024 Jan 31, 43(2): 113725. Online ahead of print. |
| 1516 | *Zygophyllum xanthoxylum* *(Zygophyllaceae)* | 2024-01-29 | Ma Q et al., Genomic analysis reveals phylogeny of Zygophyllales and mechanism for water retention of a succulent xerophyte. Plant Physiol. 2024 Jan 29 Online ahead of print. |
| 1515 | *Cymodocea nodosa* *(Cymodoceaceae)* | 2024-01-26 | Ma X et al., Seagrass genomes reveal ancient polyploidy and adaptations to the marine environment. Nat Plants. 2024 Jan 26. Online ahead of print. |
| 1514 | *Posidonia oceanica* *(Posidoniaceae)* | 2024-01-26 | Ma X et al., Seagrass genomes reveal ancient polyploidy and adaptations to the marine environment. Nat Plants. 2024 Jan 26. Online ahead of print. |
| 1513 | *Potamogeton acutifolius* *(Potamogetonaceae)* | 2024-01-26 | Ma X et al., Seagrass genomes reveal ancient polyploidy and adaptations to the marine environment. Nat Plants. 2024 Jan 26. Online ahead of print. |
| 1512 | *Thalassia testudinum* *(Hydrocharitaceae)* | 2024-01-26 | Ma X et al., Seagrass genomes reveal ancient polyploidy and adaptations to the marine environment. Nat Plants. 2024 Jan 26. Online ahead of print. |
| 1511 | *Hippophae gyantsensis* *(Elaeagnaceae)* | 2024-01-25 | Chen M et al., Chromosome-level genome assembly of Hippophae gyantsensis. Sci Data. 2024 Jan 25, 11(1): 126. |
| 1510 | *Rhus chinensis* *(Anacardiaceae)* | 2024-01-25 | Ni BB et al., A chromosome-scale genome of Rhus chinensis Mill. provides new insights into plant-insect interaction and gallotannins biosynthesis. Plant J. 2024 Jan 25. Online ahead of print. |
| 1509 | *Momordica balsamina* *(Cucurbitaceae)* | 2024-01-24 | Vinay ND et al., High-quality Momordica balsamina genome elucidates its potential use in improving stress resilience and therapeutic properties of bitter gourd. Front Plant Sci. 2024 Jan 24: 14: 1258042. eCollection 2023. |
| 1508 | *Cryptocoryne crispatula* *(Araceae)* | 2024-01-20 | Qian Z et al., The chromosome-level genome of the submerged plant Cryptocoryne crispatula provides insights into the terrestrial-freshwater transition in Araceae. DNA Res. 2024 Jan 20: dsae003. Online ahead of print. |
| 1507 | *Persicaria maackiana* *(Polygonaceae)* | 2024-01-19 | Kim KR et al., Genome assembly and microsatellite marker development using Illumina and PacBio sequencing in Persicaria maackiana (Polygonaceae) from Korea. Genes Genomics. 2024 Jan 19. Online ahead of print. |
| 1506 | *Huperzia asiatica* *(Lycopodiaceae)* | 2024-01-18 | Li C et al., Extraordinary preservation of gene collinearity over three hundred million years revealed in homosporous lycophytes. Proc Natl Acad Sci U S A. 2024 Jan 23, 121(4): e2312607121. Epub 2024 Jan 18. |
| 1505 | *Lycopodium complanatum* *(Lycopodiaceae)* | 2024-01-18 | Li C et al., Extraordinary preservation of gene collinearity over three hundred million years revealed in homosporous lycophytes. Proc Natl Acad Sci U S A. 2024 Jan 23, 121(4): e2312607121. Epub 2024 Jan 18. |
| 1504 | *Mentha suaveolens* *(Lamiaceae)* | 2024-01-17 | Yang H et al., A haplotype-resolved gap-free genome assembly provides novel insight into monoterpenoid diversification in Mentha suaveolens Variegata. Hortic Res. 2024 Jan: uhae022. eCollection 2024 Jan. |
| 1503 | *Cryptotaenia japonica* *(Apiaceae)* | 2024-01-12 | Liu H et al., The high-quality genome of Cryptotaenia japonica and comparative genomics analysis reveals anthocyanin biosynthesis in Apiaceae. Plant J. 2024 Jan 12. Online ahead of print. |
| 1502 | *Magnolia sinica* *(Magnoliaceae)* | 2024-01-11 | Cai L et al., The chromosome-scale genome of Magnolia sinica (Magnoliaceae) provides insights into the conservation of plant species with extremely small populations (PSESP). Gigascience. 2024 Jan 2, 13: giad110. |
| 1501 | *Ambrosia trifida* *(Asteraceae)* | 2024-01-06 | Yin L et al., Chromosome-level genome of Ambrosia trifida provides insights into adaptation and the evolution of pollen allergens. Int J Biol Macromol. 2024 Jan 6: 129232. Online ahead of print. |
| 1500 | *Fragaria moupinensis* *(Rosaceae)* | 2024-01-06 | Qiao Q et al., Genomic analyses provide insights into sex differentiation of tetraploid strawberry (Fragaria moupinensis). Plant Biotechnol J. 2024 Jan 6. Online ahead of print. |
| 1499 | *Fraxinus nigra* *(Oleaceae)* | 2024-01-06 | Fleck SJ et al., High quality genomes produced from single MinION flow cells clarify polyploid and demographic histories of critically endangered Fraxinus (ash) species. Commun Biol. 2024 Jan 6, 7(1): 54. |
| 1498 | *Onobrychis viciifolia* *(Fabaceae)* | 2024-01-05 | He J et al., A chromosome-level genome assembly for Onobrychis viciifolia reveals gene copy number gain underlying enhanced proanthocyanidin biosynthesis. Commun Biol. 2024 Jan 5, 7(1): 19. |
| 1497 | *Ilex aquifolium* *(Aquifoliaceae)* | 2024-01-03 | Christenhusz MJM et al., The genome sequence of the English holly, Ilex aquifolium L. (Aquifoliaceae). Wellcome Open Res. 2024 Jan 3: 9: 1. eCollection 2024. |
| 1496 | *Phanera championii* *(Fabaceae)* | 2024-01-03 | Lu Y et al., Haplotype-resolved genome assembly of Phanera championii reveals molecular mechanisms of flavonoid synthesis and adaptive evolution. Plant J. 2024 Jan 3. Online ahead of print. |
| 1495 | *Corydalis yanhusuo* *(Papaveraceae)* | 2024-01-02 | Xu D et al., Haplotype-resolved genome assembly of Corydalis yanhusuo, a traditional Chinese medicine with unusual telomere motif. Hortic Res. 2024 Jan 02: uhad296. |
| 1494 | *Datura wrightii* *(Solanaceae)* | 2024-01-02 | Goldberg JK et al., A de novo long-read genome assembly of the sacred datura plant (Datura wrightii) reveals a role of tandem gene duplications in the evolution of herbivore-defense response. BMC Genomics. 2024 Jan 2, 25(1): 15. |
| 1493 | *Escallonia herrerae* *(Escalloniaceae)* | 2024-01-02 | Chanderbali AS et al., Draft genome assemblies for two species of Escallonia (Escalloniales). BMC Genom Data. 2024 Jan 2, 25(1): 1. |
| 1492 | *Escallonia rubra* *(Escalloniaceae)* | 2024-01-02 | Chanderbali AS et al., Draft genome assemblies for two species of Escallonia (Escalloniales). BMC Genom Data. 2024 Jan 2, 25(1): 1. |
| 1491 | *Adenophora triphylla* *(Campanulaceae)* | 2023-12-30 | Kang JN et al., First Contiguous Genome Assembly of Japanese Lady Bell (Adenophora triphylla) and Insights into Development of Different Leaf Types. Genes (Basel). 2023 Dec 30, 15(1): 58. |
| 1490 | *Penstemon davidsonii* *(Plantaginaceae)* | 2023-12-29 | Ostevik KL et al., Genome report: Whole genome sequence and annotation of Penstemon davidsonii. G3 (Bethesda). 2023 Dec 29: jkad296. Online ahead of print. |
| 1489 | *Crocus sativus* *(Iridaceae)* | 2023-12-27 | Xu Z et al., Crocus genome reveals the evolutionary origin of crocin biosynthesis. Acta Pharm Sin B. 2024 Apr, 14(4): 1878-1891. Epub 2023 Dec 27. |
| 1488 | *Euryodendron excelsum* *(Pentaphylacaceae)* | 2023-12-26 | Wang ZF et al., Chromosome-level genome assembly and demographic history of Euryodendron excelsum in monotypic genus endemic to China. DNA Res. 2024 Feb 1, 31(1): dsad028. |
| 1487 | *Breynia androgyna* *(Phyllanthaceae)* | 2023-12-25 | Xia F et al., Polyploid Genome Assembly Provides Insights into Morphological Development and Ascorbic Acid Accumulation of Sauropus androgynus. Int J Mol Sci. 2023 Dec 25, 25(1): 300. |
| 1486 | *Vaccinium vitis-idaea* *(Ericaceae)* | 2023-12-24 | Hirabayashi K et al., Unveiling the evolutionary history of lingonberry (Vaccinium vitis-idaea L.) through genome sequencing and assembly of European and North American subspecies. G3 (Bethesda). 2023 Dec 24: jkad294. Online ahead of print. |
| 1485 | *Anisodus tanguticus* *(Solanaceae)* | 2023-12-20 | Yang J et al., Multiple independent losses of the biosynthetic pathway for two tropane alkaloids in the Solanaceae family. Nat Commun. 2023 Dec 20, 14(1): 8457. |

| | plant species | published | |
|---|---|---|---|
| 1484 | *Brugmansia arborea* *(Solanaceae)* | 2023-12-20 | Yang J et al., Multiple independent losses of the biosynthetic pathway for two tropane alkaloids in the Solanaceae family. Nat Commun. 2023 Dec 20, 14(1): 8457. |
| 1483 | *Lycium chinense* *(Solanaceae)* | 2023-12-20 | Yang J et al., Multiple independent losses of the biosynthetic pathway for two tropane alkaloids in the Solanaceae family. Nat Commun. 2023 Dec 20, 14(1): 8457. |
| 1482 | *Mandragora caulescens* *(Solanaceae)* | 2023-12-20 | Yang J et al., Multiple independent losses of the biosynthetic pathway for two tropane alkaloids in the Solanaceae family. Nat Commun. 2023 Dec 20, 14(1): 8457. |
| 1481 | *Agave hybrid cv. No.11648* *(Asparagaceae)* | 2023-12-19 | Yang Z et al., A chromosome-level genome assembly of Agave hybrid NO.11648 provides insights into the CAM photosynthesis. Hortic Res. 2023 Dec: uhad269. eCollection 2023 Dec. |
| 1480 | *Penthorum chinense* *(Penthoraceae)* | 2023-12-19 | Wang Z et al., Insights into the Superrosids Phylogeny and Flavonoid Synthesis from the Telomere-to-Telomere Gap-Free Genome Assembly of Penthorum chinense Pursh. Hortic Res. 2023 Dec: uhad274. eCollection 2023 Dec. |
| 1479 | *Vitis acerifolia* *(Vitaceae)* | 2023-12-19 | Cochetel N et al., A super-pangenome of the North American wild grape species. Genome Biol. 2023 Dec 19, 24(1): 290. |
| 1478 | *Vitis aestivalis* *(Vitaceae)* | 2023-12-19 | Cochetel N et al., A super-pangenome of the North American wild grape species. Genome Biol. 2023 Dec 19, 24(1): 290. |
| 1477 | *Vitis berlandieri* *(Vitaceae)* | 2023-12-19 | Cochetel N et al., A super-pangenome of the North American wild grape species. Genome Biol. 2023 Dec 19, 24(1): 290. |
| 1476 | *Vitis girdiana* *(Vitaceae)* | 2023-12-19 | Cochetel N et al., A super-pangenome of the North American wild grape species. Genome Biol. 2023 Dec 19, 24(1): 290. |
| 1475 | *Vitis monticola* *(Vitaceae)* | 2023-12-19 | Cochetel N et al., A super-pangenome of the North American wild grape species. Genome Biol. 2023 Dec 19, 24(1): 290. |
| 1474 | *Vitis mustangensis* *(Vitaceae)* | 2023-12-19 | Cochetel N et al., A super-pangenome of the North American wild grape species. Genome Biol. 2023 Dec 19, 24(1): 290. |
| 1473 | *Vitis rupestris* *(Vitaceae)* | 2023-12-19 | Cochetel N et al., A super-pangenome of the North American wild grape species. Genome Biol. 2023 Dec 19, 24(1): 290. |
| 1472 | *Passiflora foetida* *(Passifloraceae)* | 2023-12-18 | Zou Y et al., Multi-integrated genomic data for Passiflora foetida provides insights into genome size evolution and floral development in Passiflora. Mol Hortic. 2023 Dec 18, 3(1): 27. |
| 1471 | *Quercus glauca* *(Fagaceae)* | 2023-12-18 | Luo CS et al., High-quality haplotype-resolved genome assembly for ring-cup oak (Quercus glauca) provides insight into oaks demographic dynamics. Mol Ecol Resour. 2023 Dec 18: e13914. Online ahead of print. |
| 1470 | *Microcos paniculata* *(Malvaceae)* | 2023-12-15 | Liu D et al., Haplotype-resolved chromosomal-level genome assembly of Buzhaye (Microcos paniculata). Sci Data. 10, 901 (2023). |
| 1469 | *Nyctanthes arbor-tristis* *(Oleaceae)* | 2023-12-15 | Patil AB et al., Genomic and Transcriptomic Analyses Illuminates Unique Traits of Elusive Night Flowering Jasmine Parijat (Nyctanthes arbor-tristis). Physiol Plant. 2023 Nov-Dec, 175(6): e14119. |
| 1468 | *Acacia crassicarpa* *(Fabaceae)* | 2023-12-14 | Massaro I et al., Genome Report: Chromosome-level genome of the transformable northern wattle, Acacia crassicarpa. G3 (Bethesda). 2023 Dec 14: jkad284. Online ahead of print. |
| 1467 | *Castanopsis chinensis* *(Fagaceae)* | 2023-12-14 | Chen P et al., Draft genome of Castanopsis chinensis, a dominant species safeguarding biodiversity in subtropical broadleaved evergreen forests. BMC Genom Data. 2023 Dec 14, 24(1): 78. |
| 1466 | *Melaleuca quinquenervia* *(Myrtaceae)* | 2023-12-14 | Chen SH et al., A high-quality pseudo-phased genome for Melaleuca quinquenervia shows allelic diversity of NLR-type resistance genes. Gigascience. 2022 Dec 28, 12: giad102. |
| 1465 | *Dicranostigma leptopodum* *(Papaveraceae)* | 2023-12-12 | Lei W et al., From genomics to metabolomics: Deciphering sanguinarine biosynthesis in Dicranostigma leptopodum. Int J Biol Macromol. 2023 Dec 11: 128727. Online ahead of print. |
| 1464 | *Euptelea pleiosperma* *(Eupteleaceae)* | 2023-12-11 | Xiang KL et al., Genomic data and ecological niche modeling reveal an unusually slow rate of molecular evolution in the Cretaceous Eupteleaceae. Sci China Life Sci. 2023 Dec 11. Online ahead of print. |
| 1463 | *Populus cathayana* *(Salicaceae)* | 2023-12-11 | Xiang X et al., Populus cathayana genome and population resequencing provide insights into its evolution and adaptation. Hortic Res. 2023 Dec: uhad255. eCollection 2023 Dec. |
| 1462 | *Lithocarpus polystachyus* *(Fagaceae)* | 2023-12-06 | Liu H et al., Chromosome-scale genome assembly of sweet tea (Lithocarpus polystachyus Rehder). Sci Data. 2023 Dec 6, 10(1): 873. |
| 1461 | *Dendrocalamus brandisii* *(Poaceae)* | 2023-12-05 | Jiang J et al., Chromosomal-level genome and metabolome analyses of highly heterozygous allohexaploid Dendrocalamus brandisii elucidate shoot quality and developmental characteristics. J Integr Plant Biol. 2023 Dec 5. Online ahead of print. |
| 1460 | *Ipomoea pes-caprae* *(Convolvulaceae)* | 2023-12-04 | Cheng Y et al., Unveiling the genomic blueprint of salt stress: insights from Ipomoea pes-caprae L. Seed Biol. 2023, 2: 21 |
| 1459 | *Coccomyxa viridis* *(Coccomyxaceae)* | 2023-12-02 | Tagirdzhanova G et al., Genomic analysis of Coccomyxa viridis, a common low-abundance alga associated with lichen symbioses. Sci Rep. 2023 Dec 2, 13(1): 21285. |
| 1458 | *Orinus kokonorica* *(Poaceae)* | 2023-12-02 | Qu K et al., A genome assembly for Orinus kokonorica provides insights into the origin, adaptive evolution and further diversification of two closely related grass genera. Commun Biol. 2023 Dec 2, 6(1): 1223. |
| 1457 | *Khaya senegalensis* *(Meliaceae)* | 2023-11-25 | Sahu SK et al., Chromosome-scale genomes of commercially important mahoganies, Swietenia macrophylla and Khaya senegalensis. Sci Data. 2023 Nov 25, 10(1): 832. |
| 1456 | *Aeonium arboreum* *(Crassulaceae)* | 2023-11-24 | Han H et al., A chromosome-scale genome sequence of Aeonium(Aeonium arboreum Velour) provides novel insights into the evolution of anthocyanin synthesis . Gene. 2023 Nov 24:148031. Online ahead of print. |
| 1455 | *Rauvolfia tetraphylla* *(Apocynaceae)* | 2023-11-24 | Stander EA et al., The Rauvolfia tetraphylla genome suggests multiple distinct biosynthetic routes for yohimbane monoterpene indole alkaloids. Commun Biol. 2023 Nov 24, 6(1): 1197. |
| 1454 | *Nepenthes gracilis* *(Nepenthaceae)* | 2023-11-23 | Saul F et al., Subgenome dominance shapes novel gene evolution in the decaploid pitcher plant Nepenthes gracilis. Nat Plants. 2023 Nov 23. Online ahead of print. |
| 1453 | *Kitagawia praeruptora* *(Apiaceae)* | 2023-11-20 | Song C et al., A chromosome-scale genome of Peucedanum praeruptorum provide insights into Apioideae evolution and medicinal ingredient biosynthesis. Int J Biol Macromol. 2023 Nov 20: 128218. Online ahead of print. |
| 1452 | *Syzygium cumini* *(Myrtaceae)* | 2023-11-17 | Chakraborty A et al., Genome sequencing of Syzygium cumini (jamun) reveals adaptive evolution in secondary metabolism pathways associated with its medicinal properties. Front Plant Sci. 2023 Nov 17: 14: 1260414. eCollection 2023. |
| 1451 | *Helianthus tuberosus* *(Asteraceae)* | 2023-11-15 | Wang S et al., A haplotype-resolved chromosome-level genome assembly for the hexaploid Jerusalem artichoke provides insights into its origin, evolution and inulin metabolism. Plant Commun. 2023 Nov 15: 100767. Online ahead of print. |
| 1450 | *Knoxia roxburghii* *(Rubiaceae)* | 2023-11-15 | Zhang Y et al., A chromosome-level genome assembly of the Knoxia roxburghii (Rubiaceae). Sci Data. 2023 Nov 15, 10(1): 803. |
| 1449 | *Malus asiatica* *(Rosaceae)* | 2023-11-15 | Wang T et al., Pan-genome analysis of 13 Malus accessions reveals structural and sequence variations associated with fruit traits. Nat Commun. 2023 Nov 15, 14(1): 7377. |
| 1448 | *Malus orientalis* *(Rosaceae)* | 2023-11-15 | Wang T et al., Pan-genome analysis of 13 Malus accessions reveals structural and sequence variations associated with fruit traits. Nat Commun. 2023 Nov 15, 14(1): 7377. |

| | plant species | published | |
|---|---|---|---|
| 1447 | *Musa x paradisiaca* (*Musaceae*) | 2023-11-15 | Xie WZ et al., Two haplotype-resolved genome assemblies for AAB allotriploid bananas provide insights into banana subgenome asymmetric evolution and Fusarium wilt control. Plant Commun. 2023 Nov 15: 100766. Online ahead of print. |
| 1446 | *Paeonia ludlowii* (*Paeoniaceae*) | 2023-11-14 | Xiao PX et al., High-quality assembly and methylome of a Tibetan wild tree peony genome (Paeonia ludlowii) reveal the evolution of giant genome architecture. Hortic Res. 2023 Nov: uhad241. eCollection 2023 Nov. |
| 1445 | *Vicia villosa* (*Fabaceae*) | 2023-11-13 | Fuller T et al., A reference assembly for the legume cover crop hairy vetch (Vicia villosa). GigaByte. 2023 Nov 13, 2023: gigabyte98. eCollection 2023. |
| 1444 | *Agastache rugosa* (*Lamiaceae*) | 2023-11-10 | Park HS et al., A chromosome-level genome assembly of Korean mint (Agastache rugosa). Sci Data. 2023 Nov 10, 10(1): 792. |
| 1443 | *Carex pumila* (*Cyperaceae*) | 2023-11-10 | Kim KR et al., Genome Assembly and Microsatellite Marker Development Using Illumina and PacBio Sequencing in the Carex pumila (Cyperaceae) from Korea. Genes (Basel). 2023 Nov 10, 14(11): 2063. |
| 1442 | *Leonurus japonicus* (*Lamiaceae*) | 2023-11-09 | Li P et al., Multi-Omics Analyses of Two Leonurus Species Illuminate Leonurine Biosynthesis and Its Evolution. Mol Plant. 2023 Nov 9: S1674-2052(23)00358-1. Online ahead of print. |
| 1441 | *Leonurus sibiricus* (*Lamiaceae*) | 2023-11-09 | Li P et al., Multi-Omics Analyses of Two Leonurus Species Illuminate Leonurine Biosynthesis and Its Evolution. Mol Plant. 2023 Nov 9: S1674-2052(23)00358-1. Online ahead of print. |
| 1440 | *Tinospora cordifolia* (*Menispermaceae*) | 2023-11-08 | R Namitha et al., De novo genome assembly and annotation of the medicinal plant Tinospora cordifolia (Willd.) Miers ex Hook. f. & Thom Funct Integr Genomics. 2023 Nov 8, 23(4): 330. |
| 1439 | *Agapanthus africanus* (*Amaryllidaceae*) | 2023-11-06 | Hao F et al., Chromosome-level genomes of three key Allium crops and their trait evolution. Nat Genet. 2023 Nov, 55(11): 1976-1986. Epub 2023 Nov 6. |
| 1438 | *Prinsepia uniflora* (*Rosaceae*) | 2023-11-02 | Zhang L et al., A High-Quality Reference Genome Assembly of Prinsepia uniflora (Rosaceae). Genes 2023, 14(11), 2035 |
| 1437 | *Vitis zhejiang-adstricta* (*Vitaceae*) | 2023-11-01 | Li H et al., The genome of Vitis zhejiang-adstricta strengthens the protection and utilization of the endangered ancient grape endemic to China. Plant Cell Physiol. 2023 Nov 1: pcad140. Online ahead of print. |
| 1436 | *Prunus hongpingensis* (*Rosaceae*) | 2023-10-27 | Dai X et al., Genomic evidence for evolutionary history and local adaptation of two endemic apricots: Prunus hongpingensis and P. zhengheensis. Hortic Res. 2023 Oct: uhad215. eCollection 2023 Oct. |
| 1435 | *Prunus zhengheensis* (*Rosaceae*) | 2023-10-27 | Dai X et al., Genomic evidence for evolutionary history and local adaptation of two endemic apricots: Prunus hongpingensis and P. zhengheensis. Hortic Res. 2023 Oct: uhad215. eCollection 2023 Oct. |
| 1434 | *Brachypodium sylvaticum* (*Poaceae*) | 2023-10-26 | Lei L et al., The reference genome and abiotic stress responses of the model perennial grass Brachypodium sylvaticum. G3 (Bethesda). 2023 Oct 26: jkad245. Online ahead of print. |
| 1433 | *Aegilops umbellulata* (*Poaceae*) | 2023-10-25 | Abrouk M et al., Chromosome-scale assembly of the wild wheat relative Aegilops umbellulata. Sci Data. 2023 Oct 25, 10(1): 739. |
| 1432 | *Sesbania cannabina* (*Fabaceae*) | 2023-10-25 | Luo H et al., Telomere-to-telomere genome of the allotetraploid legume Sesbania cannabina reveals transposon-driven subgenome divergence and mechanisms of alkaline stress tolerance. Sci China Life Sci. 2023 Oct 25. Online ahead of print. |
| 1431 | *Syzygium samarangense* (*Myrtaceae*) | 2023-10-25 | Wei X et al., The Haplotype-resolved Autotetraploid Genome Assembly Provides Insights into the genomic evolution and fruit divergence in Wax apple (Syzygium samarangense (Bl.) Merr.et Perry). Hortic Res. 2023 Oct: uhad214. eCollection 2023 Oct. |
| 1430 | *Picea mariana* (*Pinaceae*) | 2023-10-24 | Lo T et al., Assembly and annotation of the black spruce genome provide insights on spruce phylogeny and evolution of stress response. G3 (Bethesda). 2023 Oct 24: jkad247. Online ahead of print. |
| 1429 | *Corylus americana* (*Betulaceae*) | 2023-10-23 | Brainard SH et al., The first two chromosome-scale genome assemblies of American hazelnut enable comparative genomic analysis of the genus Corylus. Plant Biotechnol J. 2023 Oct 23. Online ahead of print. |
| 1428 | *Impatiens capensis* (*Balsaminaceae*) | 2023-10-23 | Gadagkar SR et al., De-novo whole genome assembly of the orange jewelweed, Impatiens capensis Meerb. (Balsaminaceae) using nanopore long-read sequencing. PeerJ. 2023 Oct 23, 11: e16328. eCollection 2023. |
| 1427 | *Rhodiola chrysanthemifolia* (*Crassulaceae*) | 2023-10-23 | Zhang DQ et al., Two chromosome-level genome assemblies of Rhodiola shed new light on genome evolution in rapid radiation and evolution of the biosynthetic pathway of salidroside. Plant J. 2023 Oct 23. Online ahead of print. |
| 1426 | *Rhodiola kirilowii* (*Crassulaceae*) | 2023-10-23 | Zhang DQ et al., Two chromosome-level genome assemblies of Rhodiola shed new light on genome evolution in rapid radiation and evolution of the biosynthetic pathway of salidroside. Plant J. 2023 Oct 23. Online ahead of print. |
| 1425 | *Silene conica* (*Caryophyllaceae*) | 2023-10-20 | Fields PD et al., Chromosome-level genome assembly for the angiosperm Silene conica. Genome Biol Evol. 2023 Oct 20: evad192. Online ahead of print. |
| 1424 | *Vaccinium duclouxii* (*Ericaceae*) | 2023-10-19 | Zeng T et al., The Telomere-to-telomere gap-free reference genome of wild blueberry (Vaccinium duclouxii) provides its high soluble sugar and anthocyanin accumulation. Hortic Res. 2023 Oct: uhad209. eCollection 2023 Oct. |
| 1423 | *Heritiera fomes* (*Malvaceae*) | 2023-10-18 | Wang J et al., Evolution of sea-surfing plant propagule as revealed by the genomes of Heritiera mangroves. Plant J. 2023 Oct 18. Online ahead of print. |
| 1422 | *Niphotrichum japonicum* (*Grimmiaceae*) | 2023-10-18 | Zhou X et al., Chromosome-level genome assembly of Niphotrichum japonicum provides new insights into heat stress responses in mosses. Front Plant Sci. 2023 Oct 18: 14: 1271357. eCollection 2023. |
| 1421 | *Heracleum sosnowskyi* (*Apiaceae*) | 2023-10-17 | Schelkunov MI et al., The genome of the toxic invasive species Heracleum sosnowskyi carries an increased number of genes despite absence of recent whole-genome duplications. Plant J. 2023 Oct 17. Online ahead of print. |
| 1420 | *Pterocarpus macrocarpus* (*Fabaceae*) | 2023-10-17 | Yang J et al., Chromosome-scale genomes of five Hongmu species in Leguminosae. Sci Data. 2023 Oct 17, 10(1): 710. |
| 1419 | *Senna siamea* (*Fabaceae*) | 2023-10-17 | Yang J et al., Chromosome-scale genomes of five Hongmu species in Leguminosae. Sci Data. 2023 Oct 17, 10(1): 710. |
| 1418 | *Ballota nigra* (*Lamiaceae*) | 2023-10-12 | Christenhusz MJM et al., The genome sequence of black horehound, Ballota nigra L. subsp. foetida (Lam.) Hayek (Lamiaceae). Wellcome Open Res. 2023 Oct 12: 8: 439. eCollection 2023. |
| 1417 | *Pulicaria dysenterica* (*Asteraceae*) | 2023-10-12 | Christenhusz MJM et al., The genome sequence of common fleabane, Pulicaria dysenterica (L.) Bernh. (Asteraceae). Wellcome Open Res. 2023 Oct 12: 8: 447. eCollection 2023 |
| 1416 | *Quercus ilex* (*Fagaceae*) | 2023-10-12 | Rey MD et al., A first draft genome of holm oak ( Quercus ilex subsp. ballota), the most representative species of the Mediterranean forest and the Spanish agrosylvopastoral ecosystem DEHESA. Front Mol Biosci. 2023 Oct 12: 10: 1242943. eCollection 2023. |
| 1415 | *Syzygium aqueum* (*Myrtaceae*) | 2023-10-06 | Ouadi S et al., Chromosome-scale assemblies of S. malaccense, S. aqueum, S. jambos, and S. syzygioides provide insights into the evolution of Syzygium genomes. Front Plant Sci. 2023 Oct 6: 14: 1248780. eCollection 2023. |
| 1414 | *Syzygium jambos* (*Myrtaceae*) | 2023-10-06 | Ouadi S et al., Chromosome-scale assemblies of S. malaccense, S. aqueum, S. jambos, and S. syzygioides provide insights into the evolution of Syzygium genomes. Front Plant Sci. 2023 Oct 6: 14: 1248780. eCollection 2023. |
| 1413 | *Syzygium malaccense* (*Myrtaceae*) | 2023-10-06 | Ouadi S et al., Chromosome-scale assemblies of S. malaccense, S. aqueum, S. jambos, and S. syzygioides provide insights into the evolution of Syzygium genomes. Front Plant Sci. 2023 Oct 6: 14: 1248780. eCollection 2023. |
| 1412 | *Syzygium syzygioides* (*Myrtaceae*) | 2023-10-06 | Ouadi S et al., Chromosome-scale assemblies of S. malaccense, S. aqueum, S. jambos, and S. syzygioides provide insights into the evolution of Syzygium genomes. Front Plant Sci. 2023 Oct 6: 14: 1248780. eCollection 2023. |

| | plant species | published | |
|---|---|---|---|
| 1411 | *Dipteronia dyeriana* *(Sapindaceae)* | 2023-10-05 | Feng Y et al., Genome sequences and population genomics provide insights into the demographic history, inbreeding, and mutation load of two living fossil tree species of Dipteronia. Plant J. 2023 Oct 5. Online ahead of print. |
| 1410 | *Dipteronia sinensis* *(Sapindaceae)* | 2023-10-05 | Feng Y et al., Genome sequences and population genomics provide insights into the demographic history, inbreeding, and mutation load of two living fossil tree species of Dipteronia. Plant J. 2023 Oct 5. Online ahead of print. |
| 1409 | *Iris pallida* *(Iridaceae)* | 2023-10-05 | Bruccoleri RE et al., Genome assembly of the bearded iris, Iris pallida. GigaByte. 2023 Oct 5: 2023: gigabyte94. eCollection 2023. |
| 1408 | *Sesamum alatum* *(Pedaliaceae)* | 2023-10-05 | Miao H et al., Genomic evolution and insights into agronomic trait innovations of Sesamum Species. Plant Commun. 2023 Oct 5: 100729. Online ahead of print. |
| 1407 | *Sesamum angolense* *(Pedaliaceae)* | 2023-10-05 | Miao H et al., Genomic evolution and insights into agronomic trait innovations of Sesamum Species. Plant Commun. 2023 Oct 5: 100729. Online ahead of print. |
| 1406 | *Sesamum angustifolium* *(Pedaliaceae)* | 2023-10-05 | Miao H et al., Genomic evolution and insights into agronomic trait innovations of Sesamum Species. Plant Commun. 2023 Oct 5: 100729. Online ahead of print. |
| 1405 | *Sesamum calycinum* *(Pedaliaceae)* | 2023-10-05 | Miao H et al., Genomic evolution and insights into agronomic trait innovations of Sesamum Species. Plant Commun. 2023 Oct 5: 100729. Online ahead of print. |
| 1404 | *Sesamum latifolium* *(Pedaliaceae)* | 2023-10-05 | Miao H et al., Genomic evolution and insights into agronomic trait innovations of Sesamum Species. Plant Commun. 2023 Oct 5: 100729. Online ahead of print. |
| 1403 | *Sesamum radiatum* *(Pedaliaceae)* | 2023-10-05 | Miao H et al., Genomic evolution and insights into agronomic trait innovations of Sesamum Species. Plant Commun. 2023 Oct 5: 100729. Online ahead of print. |
| 1402 | *Aegle marmelos* *(Rutaceae)* | 2023-10-02 | Huang Y et al., Pangenome analysis provides insight into the evolution of the orange subfamily and a key gene for citric acid accumulation in citrus fruits. Nat Genet. 2023 Oct 2. Online ahead of print. |
| 1401 | *Citropsis gilletiana* *(Rutaceae)* | 2023-10-02 | Huang Y et al., Pangenome analysis provides insight into the evolution of the orange subfamily and a key gene for citric acid accumulation in citrus fruits. Nat Genet. 2023 Oct 2. Online ahead of print. |
| 1400 | *Citrus australasica* *(Rutaceae)* | 2023-10-02 | Huang Y et al., Pangenome analysis provides insight into the evolution of the orange subfamily and a key gene for citric acid accumulation in citrus fruits. Nat Genet. 2023 Oct 2. Online ahead of print. |
| 1399 | *Citrus linwuensis* *(Rutaceae)* | 2023-10-02 | Huang Y et al., Pangenome analysis provides insight into the evolution of the orange subfamily and a key gene for citric acid accumulation in citrus fruits. Nat Genet. 2023 Oct 2. Online ahead of print. |
| 1398 | *Citrus mangshanensis* *(Rutaceae)* | 2023-10-02 | Huang Y et al., Pangenome analysis provides insight into the evolution of the orange subfamily and a key gene for citric acid accumulation in citrus fruits. Nat Genet. 2023 Oct 2. Online ahead of print. |
| 1397 | *Luvunga scandens* *(Rutaceae)* | 2023-10-02 | Huang Y et al., Pangenome analysis provides insight into the evolution of the orange subfamily and a key gene for citric acid accumulation in citrus fruits. Nat Genet. 2023 Oct 2. Online ahead of print. |
| 1396 | *Isoetes sinensis* *(Isoetaceae)* | 2023-09-30 | Cui J et al., Chromosome-level reference genome of tetraploid Isoetes sinensis provides insights into evolution and adaption of lycophytes. Gigascience. 2022 Dec 28: 12: giad079. |
| 1395 | *Cornus wilsoniana* *(Cornaceae)* | 2023-09-29 | He Z et al., A chromosome-level genome assembly provides insights into Cornus wilsoniana evolution, oil biosynthesis and floral bud development. Hortic Res. 2023 Sep 29: uhad196. eCollection 2023 Sep. |
| 1394 | *Fraxinus americana* *(Oleaceae)* | 2023-09-29 | Zhang H et al., The chromosome-level genome assembly of Fraxinus americana provides insights into the evolution of Oleaceae plants. Int J Biol Macromol. 2023 Sep 29: 127132. Online ahead of print. |
| 1393 | *Adenosma indiana* *(Plantaginaceae)* | 2023-09-28 | Huang H et al., A chromosome-level reference genome of an aromatic medicinal plant Adenosma buchneroides. Sci Data. 2023 Sep28, 10(1): 660. |
| 1392 | *Lactuca virosa* *(Asteraceae)* | 2023-09-23 | Xiong W et al., Genome assembly and analysis of Lactuca virosa: implications for lettuce breeding. G3 (Bethesda). 2023 Sep 23: jkad204. Online ahead of print. |
| 1391 | *Balanophora fungosa* *(Balanophoraceae)* | 2023-09-21 | Chen X et al., Balanophora genomes display massively convergent evolution with other extreme holoparasites and provide novel insights into parasite-host interactions. Nat Plants. 2023 Sep 21. Online ahead of print. |
| 1390 | *Balanophora subcupularis* *(Balanophoraceae)* | 2023-09-21 | Chen X et al., Balanophora genomes display massively convergent evolution with other extreme holoparasites and provide novel insights into parasite-host interactions. Nat Plants. 2023 Sep 21. Online ahead of print. |
| 1389 | *Dalbergia cultrata* *(Fabaceae)* | 2023-09-21 | Huang P et al., Chromosome-level genome assembly and population genetic analysis of a near-threatened rosewood species (Dalbergia cultrata Pierre Graham ex Benth) provide insights into its evolutionary and cold stress responses. Front Plant Sci. 2023 Sep 21, 14: 1212967. eCollection 2023. |
| 1388 | *Scurrula parasitica* *(Loranthaceae)* | 2023-09-21 | Chen X et al., Balanophora genomes display massively convergent evolution with other extreme holoparasites and provide novel insights into parasite-host interactions. Nat Plants. 2023 Sep 21. Online ahead of print. |
| 1387 | *Selaginella kraussiana* *(Selaginellaceae)* | 2023-09-15 | Liu W et al., Genome and transcriptome of Selaginella kraussiana reveal evolution of root apical meristems in vascular plants. Curr Biol. 2023 Sep 6, S0960-9822(23)01141-7. Online ahead of print. |
| 1386 | *Chaenomeles speciosa* *(Rosaceae)* | 2023-09-14 | He S et al., A Telomere-to-telomere Reference Genome Provides Genetic Insight into the Pentacyclic Triterpenoid Biosynthesis in Chaenomeles speciosa. Hortic Res. 2023 Sep 14: uhad183. eCollection 2023 Sep. |
| 1385 | *Quercus rubra* *(Fagaceae)* | 2023-09-14 | Kapoor B et al., A haplotype-resolved chromosome-scale genome for Quercus rubra L. provides insights into the genetics of adaptive traits for red oak species. G3 (Bethesda). 2023 Sep 14, jkad209. Online ahead of print. |
| 1384 | *Vitis x doaniana* *(Vitaceae)* | 2023-09-14 | Zou C et al., A Multi-Tiered Haplotype Strategy to Enhance Phased Assembly and Fine-Mapping of a Disease Resistance Locus. Plant Physiol. 2023 Sep 14, kiad494. Online ahead of print. |
| 1383 | *Juglans cinerea* *(Juglandaceae)* | 2023-09-13 | Guzman-Torres CR et al., Conserving a threatened North American walnut: a chromosome-scale reference genome for butternut (Juglans cinerea). G3 (Bethesda). 2023 Sep 13, jkad189. Online ahead of print. |
| 1382 | *Hibiscus trionum* *(Malvaceae)* | 2023-09-11 | Koshimizu S et al., Genome and transcriptome analyses reveal genes involved in the formation of fine ridges on petal epidermal cells in Hibiscus trionum. DNA Res. 2023 Oct 1, 30(5): dsad019. |
| 1381 | *Zantedeschia elliottiana* *(Araceae)* | 2023-09-09 | Wang Y et al., Chromosome level genome assembly of colored calla lily (Zantedeschia elliottiana) Sci Data. 2023 Sep 9, 10(1): 605. |
| 1380 | *Aconitum vilmorinianum* *(Ranunculaceae)* | 2023-09-08 | Zhao D et al., Multi-omics analysis reveals the evolutionary origin of diterpenoid alkaloid biosynthesis pathways in Aconitum. J Integr Plant Biol. 2023 Sep 8. Online ahead of print. |
| 1379 | *Capsicum frutescens* *(Solanaceae)* | 2023-09-07 | Liu F et al., Genomes of cultivated and wild Capsicum species provide insights into pepper domestication and population differentiation. Nat Commun. 2023 Sep 7, 14(1): 5487. |
| 1378 | *Capsicum pubescens* *(Solanaceae)* | 2023-09-07 | Liu F et al., Genomes of cultivated and wild Capsicum species provide insights into pepper domestication and population differentiation. Nat Commun. 2023 Sep 7, 14(1): 5487. |
| 1377 | *Saussurea involucrata* *(Asteraceae)* | 2023-09-05 | Sun Y et al., Genome assembly of the snow lotus species Saussurea involucrata provides insights into acacetin and rutin biosynthesis and tolerance to an alpine environment. Hortic Res. 2023 Sep 05: uhad180. eCollection 2023 Sep. |
| 1376 | *Phyllanthus emblica* *(Phyllanthaceae)* | 2023-09-01 | Mahajan S et al., Genome of Phyllanthus emblica: the medicinal plant Amla with super antioxidant properties. Front Plant Sci. 2023 Sep 1, 14: 1210078. eCollection 2023. |

| | plant species | published | |
|---|---|---|---|
| 1375 | *Vitis labrusca* (*Vitaceae*) | 2023-08-31 | Li B et al., Vitis labrusca genome assembly reveals diversification between wild and cultivated grapevine genomes. Front Plant Sci. 2023 Aug 31, 14: 1234130. eCollection 2023. |
| 1374 | *Malus fusca* (*Rosaceae*) | 2023-08-28 | Mansfeld BN et al., A haplotype resolved chromosome-scale assembly of North American wild apple Malus fusca and comparative genomics of the fire blight Mfu10 locus. Plant J. 2023 Nov, 116(4): 989-1002. Epub 2023 Aug 28. |
| 1373 | *Rheum officinale* (*Polygonaceae*) | 2023-08-26 | Zhang H et al., The haplotype-resolved genome assembly of autotetraploid rhubarb Rheum officinale provides insights into the genome evolution and massive accumulation of anthraquinones. Plant Commun. 2023 Aug 26: 100677. Online ahead of print. |
| 1372 | *Cenchrus alopecuroides* (*Poaceae*) | 2023-08-23 | Teng K et al., Chromosome-level reference genome assembly provides insights into the evolution of Pennisetum alopecuroides. Front Plant Sci. 2023 Aug 23, 14: 1195479. eCollection 2023. |
| 1371 | *Rheum tanguticum* (*Polygonaceae*) | 2023-08-23 | Li Y et al., A chromosome-scale Rhubarb (Rheum tanguticum) genome assembly provides insights into the evolution of anthraquinone biosynthesis. Commun Biol. 2023 Aug 23, 6(1): 867. |
| 1370 | *Sophora flavescens* (*Fabaceae*) | 2023-08-22 | Kang YJ et al., Sophora genomes provide insight into the evolution of alkaloid metabolites along with small-scale gene duplication. BMC Genomics. 2023 Aug 22, 24(1): 475. |
| 1369 | *Sophora koreensis* (*Fabaceae*) | 2023-08-22 | Kang YJ et al., Sophora genomes provide insight into the evolution of alkaloid metabolites along with small-scale gene duplication. BMC Genomics. 2023 Aug 22, 24(1): 475. |
| 1368 | *Chloris gayana* (*Poaceae*) | 2023-08-18 | Maybery-Reupert K et al., Development of genomic resources for Rhodes Grass (Chloris gayana), draft genome and annotated variant discovery. Front Plant Sci. 2023 Aug 18, 14: 1239290. eCollection 2023. |
| 1367 | *Chloris virgata* (*Poaceae*) | 2023-08-17 | Ferreira LAI et al., Evolution of ACCase-inhibitor resistance in Chloris virgata is conferred by a Trp2027-Cys mutation in the herbicide target site. Pest Manag Sci. 2023 Aug 17. Online ahead of print. |
| 1366 | *Dalbergia sissoo* (*Fabaceae*) | 2023-08-17 | Sahu SK et al., Chromosome-scale genome of Indian rosewood (Dalbergia sissoo). Front Plant Sci. 2023 Aug 17: 14: 1218515. eCollection 2023 |
| 1365 | *Portulaca oleracea* (*Portulacaceae*) | 2023-08-17 | Wang X et al., Gene duplications facilitate C4-CAM compatibility in common purslane. Plant Physiol. 2023 Aug 17: kiad451. Online ahead of print. |
| 1364 | *Fagopyrum homotropicum* (*Polygonaceae*) | 2023-08-10 | Fawcett JA et al., Genome sequencing reveals the genetic architecture of heterostyly and domestication history of common buckwheat Nat Plants. 2023 Aug 10. Online ahead of print. |
| 1363 | *Takakia lepidozioides* (*Takakiaceae*) | 2023-08-09 | Hu R et al., Adaptive evolution of the enigmatic Takakia now facing climate change in Tibet. Cell. 2023 Aug 3, S0092-8674(23)00736-5. Online ahead of print. |
| 1362 | *Dalbergia cochinchinensis* (*Fabaceae*) | 2023-08-07 | Hung TH et al., Range-wide differential adaptation and genomic offset in critically endangered Asian rosewoods. Proc Natl Acad Sci U S A. 2023 Aug 15, 120(33): e2301603120. Epub 2023 Aug 7 |
| 1361 | *Dalbergia oliveri* (*Fabaceae*) | 2023-08-07 | Hung TH et al., Range-wide differential adaptation and genomic offset in critically endangered Asian rosewoods. Proc Natl Acad Sci U S A. 2023 Aug 15, 120(33): e2301603120. Epub 2023 Aug 7 |
| 1360 | *Fragaria nipponica* (*Rosaceae*) | 2023-08-03 | Jin X et al., Haplotype-resolved genomes of wild octoploid progenitors illuminate genomic diversifications from wild relatives to cultivated strawberry. Nat Plants. 2023 Aug 3. Online ahead of print. |
| 1359 | *Fragaria virginiana* (*Rosaceae*) | 2023-08-03 | Jin X et al., Haplotype-resolved genomes of wild octoploid progenitors illuminate genomic diversifications from wild relatives to cultivated strawberry. Nat Plants. 2023 Aug 3. Online ahead of print. |
| 1358 | *Ochroma pyramidale* (*Malvaceae*) | 2023-08-03 | Sahu SK et al., Chromosome-scale genomes of commercial timber trees (Ochroma pyramidale, Mesua ferrea, and Tectona grandis). Sci Data 10, 512 (2023). |
| 1357 | *Triticum monococcum* (*Poaceae*) | 2023-08-02 | Ahmed HI et al., Einkorn genomics sheds light on history of the oldest domesticated wheat. Nature. 2023 Aug 2. Online ahead of print. |
| 1356 | *Prosopis alba* (*Fabaceae*) | 2023-07-31 | Kong W et al., Genome and evolution of Prosopis alba Griseb. a drought and salinity tolerant tree legume crop for arid climates. Plants People Planet. 2023 July 31, Epub 2023 Jul 31. |
| 1355 | *Robinia pseudoacacia* (*Fabaceae*) | 2023-07-31 | Wang Z et al., Chromosome-level genome assembly and population genomics of Robinia pseudoacacia reveal the genetic basis for its wide cultivation. Commun Biol. 2023 Jul 31, 6(1): 797. |
| 1354 | *Hedera helix* (*Araliaceae*) | 2023-07-26 | Christenhusz MJM et al., The genome sequence of common ivy, Hedera helix L., 1753. Wellcome Open Res. 2023 Jul 26: 8: 325. eCollection 2023. |
| 1353 | *Tamarix chinensis* (*Tamaricaceae*) | 2023-07-26 | Liu JN et al., Genomic analyses provide insights into the evolution and salinity adaptation of halophyte Tamarix chinensis. Gigascience. 2022 Dec 28, 12: giad053. |
| 1352 | *Ailanthus altissima* (*Simaroubaceae*) | 2023-07-25 | Schley RJ et al., The genome sequence of the tree of heaven, Ailanthus altissima (Mill.) Swingle, 1916. Wellcome Open Res. 2023 Jul 25: 8: 321. eCollection 2023. |
| 1351 | *Armoracia rusticana* (*Brassicaceae*) | 2023-07-25 | Shen F et al., The allotetraploid horseradish genome provides insights into subgenome diversification and formation of critical traits. Nat Commun. 2023 Jul 25, 14(1): 4102. |
| 1350 | *Citrullus amarus* (*Cucurbitaceae*) | 2023-07-25 | Wu S et al., A Citrullus genus super-pangenome reveals extensive variations in wild and cultivated watermelons and sheds light on watermelon evolution and domestication. Plant Biotechnol J. 2023 Oct, 21(10): 1926-1928. Epub 2023 Jul 25. |
| 1349 | *Citrullus colocynthis* (*Cucurbitaceae*) | 2023-07-25 | Wu S et al., A Citrullus genus super-pangenome reveals extensive variations in wild and cultivated watermelons and sheds light on watermelon evolution and domestication. Plant Biotechnol J. 2023 Oct, 21(10): 1926-1928. Epub 2023 Jul 25. |
| 1348 | *Citrullus mucosospermus* (*Cucurbitaceae*) | 2023-07-25 | Wu S et al., A Citrullus genus super-pangenome reveals extensive variations in wild and cultivated watermelons and sheds light on watermelon evolution and domestication. Plant Biotechnol J. 2023 Oct, 21(10): 1926-1928. Epub 2023 Jul 25. |
| 1347 | *Sparganium stoloniferum* (*Typhaceae*) | 2023-07-25 | Zou Y et al., Genomic analysis of the emergent aquatic plant Sparganium stoloniferum provides insights into its clonality, local adaptation and demographic history. Mol Ecol Resour. 2023 Jul 25. Online ahead of print. |
| 1346 | *Lagerstroemia indica* (*Lythraceae*) | 2023-07-21 | Zhou Y et al., Genome assembly and resequencing analyses provide new insights into the evolution and ornamental traits of crape myrtle. Hortic Res. 2023 July 21: uhad146 eCollection 2023 July. |
| 1345 | *Prunus tenella* (*Rosaceae*) | 2023-07-21 | Qin Y et al., Chromosome-Level Genome Assembly and Population Genomic Analyses Reveal Geographic Variation and Population Genetic Structure of Prunus tenella. Int J Mol Sci. 2023 Jul 21, 24(14): 11735. |
| 1344 | *Vigna hirtella* (*Fabaceae*) | 2023-07-20 | Pootakham W et al., Genome assemblies of Vigna reflexo-pilosa (creole bean) and its progenitors, Vigna hirtella and Vigna trinervia, revealed homoeolog expression bias and expression-level dominance in the allotetraploid. Gigascience. 2022 Dec 28, 12: giad050. |
| 1343 | *Vigna reflexopilosa* (*Fabaceae*) | 2023-07-20 | Pootakham W et al., Genome assemblies of Vigna reflexo-pilosa (creole bean) and its progenitors, Vigna hirtella and Vigna trinervia, revealed homoeolog expression bias and expression-level dominance in the allotetraploid. Gigascience. 2022 Dec 28, 12: giad050. |
| 1342 | *Brasenia schreberi* (*Cabombaceae*) | 2023-07-19 | Lu B et al., Chromosome-level genome assembly of watershield (Brasenia schreberi). Sci Data. 2023 Jul 19, 10(1): 467. |
| 1341 | *Lonicera macranthoides* (*Caprifoliaceae*) | 2023-07-14 | Yin X et al., Comparative genomics of the medicinal plants Lonicera macranthoides and L. japonica provides insight into genus genome evolution and hederagenin-based saponin biosynthesis. Plant Biotechnol J. 2023 Jul 14. Online ahead of print. |
| 1340 | *Tiarella polyphylla* (*Saxifragaceae*) | 2023-07-14 | Liu L et al., Phylogenomic and syntenic data demonstrate complex evolutionary processes in early radiation of the rosids. Mol Ecol Resour. 2023 Jul 14. Online ahead of print. |

| | plant species | published | |
|---|---|---|---|
| 1339 | *Rhododendron vialii* *(Ericaceae)* | 2023-07-12 | Chang Y et al., A haplotype-resolved genome assembly of Rhododendron vialii based on PacBio HiFi reads and Hi-C data. Sci Data. 2023 Jul 12, 10(1): 451. |
| 1338 | *Brassica gravinae* *(Brassicaceae)* | 2023-07-11 | Guerreiro R et al., A genomic panel for studying C3-C4 intermediate photosynthesis in the Brassiceae tribe. Plant Cell Environ. 2023 Jul 11. Online ahead of print. |
| 1337 | *Brassica repanda* *(Brassicaceae)* | 2023-07-11 | Guerreiro R et al., A genomic panel for studying C3-C4 intermediate photosynthesis in the Brassiceae tribe. Plant Cell Environ. 2023 Jul 11. Online ahead of print. |
| 1336 | *Brassica tournefortii* *(Brassicaceae)* | 2023-07-11 | Guerreiro R et al., A genomic panel for studying C3-C4 intermediate photosynthesis in the Brassiceae tribe. Plant Cell Environ. 2023 Jul 11. Online ahead of print. |
| 1335 | *Carrichtera annua* *(Brassicaceae)* | 2023-07-11 | Guerreiro R et al., A genomic panel for studying C3-C4 intermediate photosynthesis in the Brassiceae tribe. Plant Cell Environ. 2023 Jul 11. Online ahead of print. |
| 1334 | *Diplotaxis acris* *(Brassicaceae)* | 2023-07-11 | Guerreiro R et al., A genomic panel for studying C3-C4 intermediate photosynthesis in the Brassiceae tribe. Plant Cell Environ. 2023 Jul 11. Online ahead of print. |
| 1333 | *Diplotaxis erucoides* *(Brassicaceae)* | 2023-07-11 | Guerreiro R et al., A genomic panel for studying C3-C4 intermediate photosynthesis in the Brassiceae tribe. Plant Cell Environ. 2023 Jul 11. Online ahead of print. |
| 1332 | *Diplotaxis harra* *(Brassicaceae)* | 2023-07-11 | Guerreiro R et al., A genomic panel for studying C3-C4 intermediate photosynthesis in the Brassiceae tribe. Plant Cell Environ. 2023 Jul 11. Online ahead of print. |
| 1331 | *Diplotaxis muralis* *(Brassicaceae)* | 2023-07-11 | Guerreiro R et al., A genomic panel for studying C3-C4 intermediate photosynthesis in the Brassiceae tribe. Plant Cell Environ. 2023 Jul 11. Online ahead of print. |
| 1330 | *Diplotaxis tenuifolia* *(Brassicaceae)* | 2023-07-11 | Guerreiro R et al., A genomic panel for studying C3-C4 intermediate photosynthesis in the Brassiceae tribe. Plant Cell Environ. 2023 Jul 11. Online ahead of print. |
| 1329 | *Diplotaxis tenuisiliqua* *(Brassicaceae)* | 2023-07-11 | Guerreiro R et al., A genomic panel for studying C3-C4 intermediate photosynthesis in the Brassiceae tribe. Plant Cell Environ. 2023 Jul 11. Online ahead of print. |
| 1328 | *Diplotaxis viminea* *(Brassicaceae)* | 2023-07-11 | Guerreiro R et al., A genomic panel for studying C3-C4 intermediate photosynthesis in the Brassiceae tribe. Plant Cell Environ. 2023 Jul 11. Online ahead of print. |
| 1327 | *Eutrema japonicum* *(Brassicaceae)* | 2023-07-11 | Tanaka H et al., Haplotype-resolved chromosomal-level assembly of wasabi (Eutrema japonicum) genome. Sci Data. 2023 Jul 11, 10(1): 441. |
| 1326 | *Moricandia nitens* *(Brassicaceae)* | 2023-07-11 | Guerreiro R et al., A genomic panel for studying C3-C4 intermediate photosynthesis in the Brassiceae tribe. Plant Cell Environ. 2023 Jul 11. Online ahead of print. |
| 1325 | *Moricandia sinaica* *(Brassicaceae)* | 2023-07-11 | Guerreiro R et al., A genomic panel for studying C3-C4 intermediate photosynthesis in the Brassiceae tribe. Plant Cell Environ. 2023 Jul 11. Online ahead of print. |
| 1324 | *Moricandia spinosa* *(Brassicaceae)* | 2023-07-11 | Guerreiro R et al., A genomic panel for studying C3-C4 intermediate photosynthesis in the Brassiceae tribe. Plant Cell Environ. 2023 Jul 11. Online ahead of print. |
| 1323 | *Moricandia suffruticosa* *(Brassicaceae)* | 2023-07-11 | Guerreiro R et al., A genomic panel for studying C3-C4 intermediate photosynthesis in the Brassiceae tribe. Plant Cell Environ. 2023 Jul 11. Online ahead of print. |
| 1322 | *Polygala tenuifolia* *(Polygalaceae)* | 2023-07-11 | Meng F et al., Genome Assembly of Polygala tenuifolia Provides Insights into Its Karyotype Evolution and Triterpenoid Saponin Biosynthesis. Hortic Res. 2023 Jul 11: uhad139. |
| 1321 | *Heliophila variabilis* *(Brassicaceae)* | 2023-07-10 | Huang Y et al., The meso-octoploid Heliophila variabilis genome sheds a new light on the impact of polyploidization and diploidization on the diversity of the Cape flora. Plant J. 2023 Jul 10. Online ahead of print. |
| 1320 | *Rheum nobile* *(Polygonaceae)* | 2023-07-10 | Feng T et al., The genome of the glasshouse plant noble rhubarb (Rheum nobile) provides a window into alpine adaptation. Commun Biol. 2023 Jul 10, 6(1): 706. |
| 1319 | *Wurfbainia longiligularis* *(Zingiberaceae)* | 2023-07-10 | Yang P et al., Comparing genomes of Fructus Amomi-producing species reveals genetic basis of volatile terpenoid divergence. Plant Physiol. 2023 Jul 10: kiad400. Online ahead of print. |
| 1318 | *Lavandula x intermedia* *(Lamiaceae)* | 2023-07-03 | Li J et al., Decoupling subgenomes within hybrid lavandin provide new insights into speciation and monoterpenoid diversification of Lavandula. Plant Biotechnol J. 2023 Jul 3. Online ahead of print. |
| 1317 | *Ormosia emarginata* *(Fabaceae)* | 2023-06-29 | Liu PP et al., Genome Assemblies of Two Ormosia Species: Gene Duplication Related to Their Evolutionary Adaptation. Agronomy. 2023, 13(7), 1757 |
| 1316 | *Ormosia semicastrata* *(Fabaceae)* | 2023-06-29 | Liu PP et al., Genome Assemblies of Two Ormosia Species: Gene Duplication Related to Their Evolutionary Adaptation. Agronomy. 2023, 13(7), 1757 |
| 1315 | *Lycopodium clavatum* *(Lycopodiaceae)* | 2023-06-28 | Yu JG et al., The first homosporous lycophyte genome revealed the association between the recent dynamic accumulation of LTR-RTs and genome size variation. Plant Mol Biol. 2023 Jun 28. Online ahead of print. |
| 1314 | *Metasequoia glyptostroboides* *(Cupressaceae)* | 2023-06-28 | Fu F et al., The Metasequoia genome and evolutionary relationship among redwoods. Plant Commun. 2023 Jun 28, 100643. Online ahead of print. |
| 1313 | *Scilla litardierei* *(Asparagaceae)* | 2023-06-28 | Radosavljevic I et al., Towards the Investigation of the Adaptive Divergence in a Species of Exceptional Ecological Plasticity: Chromosome-Scale Genome Assembly of Chouardia litardierei (Hyacinthaceae). Int J Mol Sci. 2023 Jun 28, 24(13): 10755. |
| 1312 | *Poa infirma* *(Poaceae)* | 2023-06-26 | Benson CW et al., Homoeologous evolution of the allotetraploid genome of Poa annua L. BMC Genomics. 2023 Jun 26, 24(1): 350. |
| 1311 | *Poa supina* *(Poaceae)* | 2023-06-26 | Benson CW et al., Homoeologous evolution of the allotetraploid genome of Poa annua L. BMC Genomics. 2023 Jun 26, 24(1): 350. |
| 1310 | *Pterocarpus santalinus* *(Fabaceae)* | 2023-06-26 | Ghosh Dasgupta M et al., The draft genome reveals early duplication event in Pterocarpus santalinus: an endemic timber species. Planta. 2023 Jun 26, 258(2): 27. |
| 1309 | *Trapa incisa* *(Lythraceae)* | 2023-06-24 | Qu M et al., Chromosome-level assemblies of cultivated water chestnut Trapa bicornis and its wild relative Trapa incisa. Sci Data. 2023 Jun 24, 10(1): 407 |
| 1308 | *Acorus gramineus* *(Acoraceae)* | 2023-06-20 | Ma L et al., Diploid and tetraploid genomes of Acorus and the evolution of monocots. Nat Commun. 2023 Jun 20, 14(1): 3661. |
| 1307 | *Murraya paniculata* *(Rutaceae)* | 2023-06-20 | Yang T et al., Chromosome-level genome assembly of Murraya paniculata sheds light on biosynthesis of floral volatiles. BMC Biol. 2023 Jun 20, 21(1): 142. |
| 1306 | *Cyamopsis tetragonoloba* *(Fabaceae)* | 2023-06-19 | Gaikwad K et al., The chromosome-scale genome assembly of cluster bean provides molecular insight into edible gum (galactomannan) biosynthesis family genes. Sci Rep. 2023 Jun 19, 13(1): 9941. |
| 1305 | *Juglans californica* *(Juglandaceae)* | 2023-06-19 | Fitz-Gibbon S et al., Reference genome of California walnut, Juglans californica, and resemblance with other genomes in the order Fagales. J Hered. 2023 Jun 19, esad036. Online ahead of print. |
| 1304 | *Bougainvillea x buttiana* *(Nyctaginaceae)* | 2023-06-13 | Lan L et al., A high-quality Bougainvillea genome provides new insights into the evolutionary history and pigment biosynthetic pathways in the Caryophyllales. Hortic Res. 2023 Jun 13: uhad124. |

| | plant species | published | |
|---|---|---|---|
| 1303 | *Chionographis japonica* (Melanthiaceae) | 2023-06-13 | Kuo YT et al., Holocentromeres can consist of merely a few megabase-sized satellite arrays. Nat Commun. 2023 Jun 13, 14(1): 3502. |
| 1302 | *Thalia dealbata* (Marantaceae) | 2023-06-13 | Tang M et al., A near-complete genome assembly of Thalia dealbata Fraser (Marantaceae). Front Plant Sci. 2023 Jun 13, 14: 1183361. eCollection 2023. |
| 1301 | *Alpinia oxyphylla* (Zingiberaceae) | 2023-06-08 | Pan K et al., Chromosome-level genome and multi-omics analyses provide insights into the geo-herbalism properties of Alpinia oxyphylla. Front Plant Sci. 2023 Jun 8, 14: 1161257. eCollection 2023. |
| 1300 | *Silene latifolia* (Caryophyllaceae) | 2023-06-07 | Yue J et al., The origin and evolution of sex chromosomes, revealed by sequencing of the Silene latifolia female genome. Curr Biol. 2023 Jun 5: S0960-9822(23)00678-4. Online ahead of print. |
| 1299 | *Cenchrus fungigraminus* (Poaceae) | 2023-06-03 | Zheng H et al., A near-complete genome assembly of the allotetrapolyploid Cenchrus fungigraminus (JUJUNCAO) provides insights into its evolution and C4 photosynthesis. Plant Commun. 2023 Jun 3: 100633. Online ahead of print. |
| 1298 | *Abelmoschus esculentus* (Malvaceae) | 2023-06-02 | Wang R et al., The genome of okra (Abelmoschus esculentus) provides insights into its genome evolution and high nutrient content. Hortic Res. 2023 Jun 02: uhad120. |
| 1297 | *Dittrichia graveolens* (Asteraceae) | 2023-06-01 | McEvoy SL et al., Chromosome-level reference genome of stinkwort, Dittrichia graveolens (L.) Greuter: a resource for studies on invasion, range expansion, and evolutionary adaptation under global change. J Hered. 2023 Jun 1, esad033. Online ahead of print. |
| 1296 | *Santalum yasi* (Santalaceae) | 2023-06-01 | Hong Z et al., Chromosome-level genome assemblies from two sandalwood species provide insights into the evolution of the Santalales. Commun Biol. 2023 Jun 1, 6(1): 587. |
| 1295 | *Solanum rostratum* (Solanaceae) | 2023-06-01 | Zhang Y et al., Chromosome-level genome assembly and annotation of the prickly nightshade Solanum rostratum Dunal. Sci Data. 2023 Jun 1, 10(1): 341. |
| 1294 | *Actinidia hemsleyana* (Actinidiaceae) | 2023-05-31 | Yu X et al., Genomic analyses reveal dead-end hybridization between two deeply divergent kiwifruit species rather than homoploid hybrid speciation. Plant J. 2023 May 31. Online ahead of print. |
| 1293 | *Actinidia zhejiangensis* (Actinidiaceae) | 2023-05-31 | Yu X et al., Genomic analyses reveal dead-end hybridization between two deeply divergent kiwifruit species rather than homoploid hybrid speciation. Plant J. 2023 May 31. Online ahead of print. |
| 1292 | *Melilotus officinalis* (Fabaceae) | 2023-05-31 | He Q et al., Chromosome-scale assembly and analysis of Melilotus officinalis genome for SSR development and nodulation genes analysis. Plant Genome. 2023 May 31: e20345. Online ahead of print. |
| 1291 | *Anisodus stramoniifolius* (Solanaceae) | 2023-05-25 | Wu Y et al., Phylogenomic discovery of deleterious mutations facilitates hybrid potato breeding. Cell. 2023 May 25, 186(11): 2313-2328.e15. Epub 2023 May 4. |
| 1290 | *Datura innoxia* (Solanaceae) | 2023-05-25 | Wu Y et al., Phylogenomic discovery of deleterious mutations facilitates hybrid potato breeding. Cell. 2023 May 25, 186(11): 2313-2328.e15. Epub 2023 May 4. |
| 1289 | *Lycianthes biflora* (Solanaceae) | 2023-05-25 | Wu Y et al., Phylogenomic discovery of deleterious mutations facilitates hybrid potato breeding. Cell. 2023 May 25, 186(11): 2313-2328.e15. Epub 2023 May 4. |
| 1288 | *Nicandra physalodes* (Solanaceae) | 2023-05-25 | Wu Y et al., Phylogenomic discovery of deleterious mutations facilitates hybrid potato breeding. Cell. 2023 May 25, 186(11): 2313-2328.e15. Epub 2023 May 4. |
| 1287 | *Nicotiana longiflora* (Solanaceae) | 2023-05-25 | Wu Y et al., Phylogenomic discovery of deleterious mutations facilitates hybrid potato breeding. Cell. 2023 May 25, 186(11): 2313-2328.e15. Epub 2023 May 4. |
| 1286 | *Physalis angulata* (Solanaceae) | 2023-05-25 | Wu Y et al., Phylogenomic discovery of deleterious mutations facilitates hybrid potato breeding. Cell. 2023 May 25, 186(11): 2313-2328.e15. Epub 2023 May 4. |
| 1285 | *Plantago major* (Plantaginaceae) | 2023-05-25 | Lyu S et al., Genome assembly of the pioneer species Plantago major L. (Plantaginaceae) provides insight into its global distribution and adaptation to metal-contaminated soil. DNA Res. 2023 May 25: dsad013. Online ahead of print. |
| 1284 | *Solanum americanum* (Solanaceae) | 2023-05-25 | Wu Y et al., Phylogenomic discovery of deleterious mutations facilitates hybrid potato breeding. Cell. 2023 May 25, 186(11): 2313-2328.e15. Epub 2023 May 4. |
| 1283 | *Solanum atropurpureum* (Solanaceae) | 2023-05-25 | Wu Y et al., Phylogenomic discovery of deleterious mutations facilitates hybrid potato breeding. Cell. 2023 May 25, 186(11): 2313-2328.e15. Epub 2023 May 4. |
| 1282 | *Solanum capsicoides* (Solanaceae) | 2023-05-25 | Wu Y et al., Phylogenomic discovery of deleterious mutations facilitates hybrid potato breeding. Cell. 2023 May 25, 186(11): 2313-2328.e15. Epub 2023 May 4. |
| 1281 | *Solanum dulcamara* (Solanaceae) | 2023-05-25 | Wu Y et al., Phylogenomic discovery of deleterious mutations facilitates hybrid potato breeding. Cell. 2023 May 25, 186(11): 2313-2328.e15. Epub 2023 May 4. |
| 1280 | *Solanum erianthum* (Solanaceae) | 2023-05-25 | Wu Y et al., Phylogenomic discovery of deleterious mutations facilitates hybrid potato breeding. Cell. 2023 May 25, 186(11): 2313-2328.e15. Epub 2023 May 4. |
| 1279 | *Solanum laciniatum* (Solanaceae) | 2023-05-25 | Wu Y et al., Phylogenomic discovery of deleterious mutations facilitates hybrid potato breeding. Cell. 2023 May 25, 186(11): 2313-2328.e15. Epub 2023 May 4. |
| 1278 | *Solanum lyratum* (Solanaceae) | 2023-05-25 | Wu Y et al., Phylogenomic discovery of deleterious mutations facilitates hybrid potato breeding. Cell. 2023 May 25, 186(11): 2313-2328.e15. Epub 2023 May 4. |
| 1277 | *Solanum macrocarpon* (Solanaceae) | 2023-05-25 | Wu Y et al., Phylogenomic discovery of deleterious mutations facilitates hybrid potato breeding. Cell. 2023 May 25, 186(11): 2313-2328.e15. Epub 2023 May 4. |
| 1276 | *Solanum mammosum* (Solanaceae) | 2023-05-25 | Wu Y et al., Phylogenomic discovery of deleterious mutations facilitates hybrid potato breeding. Cell. 2023 May 25, 186(11): 2313-2328.e15. Epub 2023 May 4. |
| 1275 | *Solanum ochranthum* (Solanaceae) | 2023-05-25 | Wu Y et al., Phylogenomic discovery of deleterious mutations facilitates hybrid potato breeding. Cell. 2023 May 25, 186(11): 2313-2328.e15. Epub 2023 May 4. |
| 1274 | *Solanum pseudocapsicum* (Solanaceae) | 2023-05-25 | Wu Y et al., Phylogenomic discovery of deleterious mutations facilitates hybrid potato breeding. Cell. 2023 May 25, 186(11): 2313-2328.e15. Epub 2023 May 4. |
| 1273 | *Solanum retroflexum* (Solanaceae) | 2023-05-25 | Wu Y et al., Phylogenomic discovery of deleterious mutations facilitates hybrid potato breeding. Cell. 2023 May 25, 186(11): 2313-2328.e15. Epub 2023 May 4. |
| 1272 | *Solanum scabrum* (Solanaceae) | 2023-05-25 | Wu Y et al., Phylogenomic discovery of deleterious mutations facilitates hybrid potato breeding. Cell. 2023 May 25, 186(11): 2313-2328.e15. Epub 2023 May 4. |
| 1271 | *Solanum seaforthianum* (Solanaceae) | 2023-05-25 | Wu Y et al., Phylogenomic discovery of deleterious mutations facilitates hybrid potato breeding. Cell. 2023 May 25, 186(11): 2313-2328.e15. Epub 2023 May 4. |
| 1270 | *Solanum septemlobum* (Solanaceae) | 2023-05-25 | Wu Y et al., Phylogenomic discovery of deleterious mutations facilitates hybrid potato breeding. Cell. 2023 May 25, 186(11): 2313-2328.e15. Epub 2023 May 4. |
| 1269 | *Solanum sisymbriifolium* (Solanaceae) | 2023-05-25 | Wu Y et al., Phylogenomic discovery of deleterious mutations facilitates hybrid potato breeding. Cell. 2023 May 25, 186(11): 2313-2328.e15. Epub 2023 May 4. |
| 1268 | *Solanum spirale* (Solanaceae) | 2023-05-25 | Wu Y et al., Phylogenomic discovery of deleterious mutations facilitates hybrid potato breeding. Cell. 2023 May 25, 186(11): 2313-2328.e15. Epub 2023 May 4. |

| | plant species | published | |
|---|---|---|---|
| 1267 | *Solanum stramoniifolium* (*Solanaceae*) | 2023-05-25 | Wu Y et al., Phylogenomic discovery of deleterious mutations facilitates hybrid potato breeding. Cell. 2023 May 25, 186(11): 2313-2328.e15. Epub 2023 May 4. |
| 1266 | *Solanum torvum* (*Solanaceae*) | 2023-05-25 | Wu Y et al., Phylogenomic discovery of deleterious mutations facilitates hybrid potato breeding. Cell. 2023 May 25, 186(11): 2313-2328.e15. Epub 2023 May 4. |
| 1265 | *Solanum viarum* (*Solanaceae*) | 2023-05-25 | Wu Y et al., Phylogenomic discovery of deleterious mutations facilitates hybrid potato breeding. Cell. 2023 May 25, 186(11): 2313-2328.e15. Epub 2023 May 4. |
| 1264 | *Solanum wrightii* (*Solanaceae*) | 2023-05-25 | Wu Y et al., Phylogenomic discovery of deleterious mutations facilitates hybrid potato breeding. Cell. 2023 May 25, 186(11): 2313-2328.e15. Epub 2023 May 4. |
| 1263 | *Tubocapsicum anomalum* (*Solanaceae*) | 2023-05-25 | Wu Y et al., Phylogenomic discovery of deleterious mutations facilitates hybrid potato breeding. Cell. 2023 May 25, 186(11): 2313-2328.e15. Epub 2023 May 4. |
| 1262 | *Prunus mongolica* (*Rosaceae*) | 2023-05-23 | Zhu Q et al., Chromosome-level genome assembly of an endangered plant Prunus mongolica using PacBio and Hi-C technologies. DNA Res. 2023 May 23, dsad012. Online ahead of print. |
| 1261 | *Platycarya strobilacea* (*Juglandaceae*) | 2023-05-22 | Cao Y et al., Genomic Insights into Adaptation to Karst Limestone and Incipient Speciation in East Asian Platycarya spp. (Juglandaceae). Mol Biol Evol. 2023 May 22: msad121. Online ahead of print. |
| 1260 | *Apocynum pictum* (*Apocynaceae*) | 2023-05-19 | Gao G et al., Comparative genome and metabolome analyses uncover the evolution and flavonoid biosynthesis between Apocynum venetum and Apocynum hendersonii. iScience. 2023 Apr 28, 26(5): 106772 eCollection 2023 May 19. |
| 1259 | *Apocynum venetum* (*Apocynaceae*) | 2023-05-19 | Gao G et al., Comparative genome and metabolome analyses uncover the evolution and flavonoid biosynthesis between Apocynum venetum and Apocynum hendersonii. iScience. 2023 Apr 28, 26(5): 106772 eCollection 2023 May 19. |
| 1258 | *Chenopodium watsonii* (*Amaranthaceae*) | 2023-05-17 | Young LA et al., A chromosome-scale reference of Chenopodium watsonii helps elucidate relationships within the North American A-genome Chenopodium species and with quinoa. Plant Genome. 2023 May 17: e20349. Online ahead of print. |
| 1257 | *Typha angustifolia* (*Typhaceae*) | 2023-05-17 | Liao Y et al., Chromosome-level genome and high nitrogen stress response of the widespread and ecologically important wetland plant Typha angustifolia. Front Plant Sci. 2023 May 17, 14: 1138498. eCollection 2023. |
| 1256 | *Cuscuta epithymum* (*Convolvulaceae*) | 2023-05-16 | Neumann P et al., Disruption of the standard kinetochore in holocentric Cuscuta species. Proc Natl Acad Sci U S A. 2023 May 23, 120(21): e2300877120. doi: Epub 2023 May 16. |
| 1255 | *Cuscuta europaea* (*Convolvulaceae*) | 2023-05-16 | Neumann P et al., Disruption of the standard kinetochore in holocentric Cuscuta species. Proc Natl Acad Sci U S A. 2023 May 23, 120(21): e2300877120. doi: Epub 2023 May 16. |
| 1254 | *Bothriochloa decipiens* (*Poaceae*) | 2023-05-12 | De Silva NP et al., Genome assembly of an Australian native grass species reveals a recent whole-genome duplication and biased gene retention of genes involved in stress response. Gigascience. 2022 Dec 28, 12: giad034. |
| 1253 | *Diospyros kaki* (*Ebenaceae*) | 2023-05-11 | Li H et al., Allele-aware chromosome-level genome assembly of the autohexaploid Diospyros kaki Thunb. Sci Data. 2023 May 11, 10(1): 270. |
| 1252 | *Elettaria cardamomum* (*Zingiberaceae*) | 2023-05-10 | Gaikwad AB et al., Small cardamom genome: development and utilization of microsatellite markers from a draft genome sequence of Elettaria cardamomum Maton. Front Plant Sci. 2023 May 10, 14: 1161499. eCollection 2023. |
| 1251 | *Prunus cerasus* (*Rosaceae*) | 2023-05-10 | Goeckeritz CZ et al., Genome of tetraploid sour cherry (Prunus cerasus L.) Montmorency identifies three distinct ancestral Prunus genomes. Hortic Res. 2023 May 10: uhad097. eCollection 2023 Mai. |
| 1250 | *Coriaria nepalensis* (*Coriariaceae*) | 2023-05-09 | Zhao SW et al., Haplotype-resolved genome assembly of Coriaria nepalensis a non-legume nitrogen-fixing shrub. Sci Data. 2023 May 9, 10(1): 259. |
| 1249 | *Helichrysum umbraculigerum* (*Asteraceae*) | 2023-05-01 | Berman P et al., Parallel evolution of cannabinoid biosynthesis. Nat Plants. 2023 May 1. Online ahead of print. |
| 1248 | *Eucalyptus urophylla x grandis* (*Myrtaceae*) | 2023-04-28 | Shen C et al., E. urophylla x E. grandis high-quality genome and comparative genomics provide insights on evolution and diversification of eucalyptus. BMC Genomics. 2023 Apr 28, 24(1): 223. |
| 1247 | *Castanopsis purpurella* (*Fagaceae*) | 2023-04-25 | Huang WC et al., A chromosome-scale genome assembly of Castanopsis hystrix provides new insights into the evolution and adaptation of Fagaceae species. Front Plant Sci. 2023 Apr 25, 14: 1174972. eCollection 2023. |
| 1246 | *Solanum arcanum* (*Solanaceae*) | 2023-04-19 | Jiang L et al., Chromosome-scale genome assembly-assisted identification of Mi-9 gene in Solanum arcanum accession LA2157, conferring heat-stable resistance to Meloidogyne incognita. Plant Biotechnol J. 2023 Apr 19. Online ahead of print. |
| 1245 | *Anisodus acutangulus* (*Solanaceae*) | 2023-04-17 | Wang YJ et al., Genomic and structural basis for evolution of tropane alkaloid biosynthesis. Proc Natl Acad Sci U S A. 2023 Apr 25, 120(17): e2302448120. Epub 2023 Apr 17. |
| 1244 | *Erythroxylum novogranatense* (*Erythroxylaceae*) | 2023-04-17 | Wang YJ et al., Genomic and structural basis for evolution of tropane alkaloid biosynthesis. Proc Natl Acad Sci U S A. 2023 Apr 25, 120(17): e2302448120. Epub 2023 Apr 17. |
| 1243 | *Vigna stipulacea* (*Fabaceae*) | 2023-04-17 | Takahashi Y et al., Domesticating Vigna stipulacea: Chromosome-Level genome assembly reveals VsPSAT1 as a candidate gene decreasing hard-seededness. Front Plant Sci. 2023 Apr 17, 14: 1119625. eCollection 2023. |
| 1242 | *Scaevola taccada* (*Goodeniaceae*) | 2023-04-16 | Li S et al., Chromosomal-Scale Genome Assemblies ofTwo Coastal Plant Species, Scaevola taccada and S. hainanensis - Insight into Adaptation Outside of the Common Range. Int. J. Mol. Sci. 2023, 24(8), 7355 |
| 1241 | *Nepenthes mirabilis* (*Nepenthaceae*) | 2023-04-14 | Gao Y et al., Draft genome and transcriptome of Nepenthes mirabilis, a carnivorous plant in China. BMC Genom Data. 2023 Apr 14, 24(1): 21. |
| 1240 | *Chrysanthemum x morifolium* (*Asteraceae*) | 2023-04-11 | Song A et al., Analyses of a chromosome-scale genome assembly reveal the origin and evolution of cultivated chrysanthemum. Nat Commun. 2023 Apr 11, 14(1): 2021. |
| 1239 | *Prunus pusilliflora* (*Rosaceae*) | 2023-04-10 | Jiu S et al., Chromosome-scale genome assembly of Prunus pusilliflora provides novel insights into genome evolution, disease resistance, and dormancy release in Cerasus L. Hortic Res. 2023 Apr 10: uhad062. eCollection 2023 Apr. |
| 1238 | *Solanum chmielewskii* (*Solanaceae*) | 2023-04-06 | Li N et al., Super-pangenome analyses highlight genomic diversity and structural variation across wild and cultivated tomato species. Nat Genet. 2023 May, 55(5): 852-860. Epub 2023 Apr 6. |
| 1237 | *Solanum corneliomulleri* (*Solanaceae*) | 2023-04-06 | Li N et al., Super-pangenome analyses highlight genomic diversity and structural variation across wild and cultivated tomato species. Nat Genet. 2023 May, 55(5): 852-860. Epub 2023 Apr 6. |
| 1236 | *Solanum neorickii* (*Solanaceae*) | 2023-04-06 | Li N et al., Super-pangenome analyses highlight genomic diversity and structural variation across wild and cultivated tomato species. Nat Genet. 2023 May, 55(5): 852-860. Epub 2023 Apr 6. |
| 1235 | *Solanum peruvianum* (*Solanaceae*) | 2023-04-06 | Li N et al., Super-pangenome analyses highlight genomic diversity and structural variation across wild and cultivated tomato species. Nat Genet. 2023 May, 55(5): 852-860. Epub 2023 Apr 6. |
| 1234 | *Tagetes erecta* (*Asteraceae*) | 2023-04-05 | Qi X et al., Chromosome-scale genome assembly of marigold (Tagetes erecta L.): an ornamental plant and feedstock for industrial lutein production. Hortic Plant J. 2023 Apr, 9(6): 1119-1130 |
| 1233 | *Antirrhinum hispanicum* (*Plantaginaceae*) | 2023-04-04 | Zhu S et al., The snapdragon genomes reveal the evolutionary dynamics of the S locus supergene. Mol Biol Evol. 2023 Apr 4: msad080. Online ahead of print. |
| 1232 | *Antirrhinum linkianum* (*Plantaginaceae*) | 2023-04-04 | Zhu S et al., The snapdragon genomes reveal the evolutionary dynamics of the S locus supergene. Mol Biol Evol. 2023 Apr 4: msad080. Online ahead of print. |
| 1231 | *Misopates orontium* (*Plantaginaceae*) | 2023-04-04 | Zhu S et al., The snapdragon genomes reveal the evolutionary dynamics of the S locus supergene. Mol Biol Evol. 2023 Apr 4: msad080. Online ahead of print. |

| | plant species | published | |
|---|---|---|---|
| 1230 | *Przewalskia tangutica* *(Solanaceae)* | 2023-04-04 | Wu Y et al., The genome sequence and demographic history of Przewalskia tangutica (Solanaceae), an endangered alpine plant on the Qinghai-Tibet Plateau. DNA Res. 2023 Apr 4: dsad005. Online ahead of print. |
| 1229 | *Citrus australis* *(Rutaceae)* | 2023-04-03 | Nakandala U et al., Haplotype resolved chromosome level genome assembly of Citrus australis reveals disease resistance and other citrus specific genes. Hortic Res. 2023 Apr 03: uhad058. eCollection 2023 Apr. |
| 1228 | *Poa pratensis* *(Poaceae)* | 2023-04-01 | Phillips AR et al., A happy accident: a novel turfgrass reference genome. G3 (Bethesda). 2023 Apr 1: jkad073. Online ahead of print. |
| 1227 | *Zygophyllum mongolicum* *(Zygophyllaceae)* | 2023-03-31 | Liu B et al., Chromosome-level genome assembly of the endangered plant Tetraena mongolica. DNA Res. 2023 Mar 31: dsad004. Online ahead of print. |
| 1226 | *Tripidium rufipilum* *(Poaceae)* | 2023-03-30 | Wang T et al., A complete gap-free diploid genome in Saccharum complex and the genomic footprints of evolution in the highly polyploid Saccharum genus. Nat Plants. 2023 Mar 30. Online ahead of print. |
| 1225 | *Citrus polytrifolia* *(Rutaceae)* | 2023-03-29 | Zhang S et al., Insights into identifying resistance genes for cold and disease stresses through chromosome-level reference genome analyses of Poncirus polyandra. Genomics. 2023 Mar 29:110617. Online ahead of print. |
| 1224 | *Lactuca saligna* *(Asteraceae)* | 2023-03-29 | Xiong W et al., The genome of Lactuca saligna, a wild relative of lettuce, provides insight into non-host resistance to the downy mildew Bremia lactucae. Plant J. 2023 Mar 29. Online ahead of print. |
| 1223 | *Ambrosia artemisiifolia* *(Asteraceae)* | 2023-03-27 | Battlay P et al., Large haploblocks underlie rapid adaptation in the invasive weed Ambrosia artemisiifolia. Nat Commun. 2023 Mar 27, 14(1): 1717. |
| 1222 | *Bromus tectorum* *(Poaceae)* | 2023-03-27 | Revolinski SR et al., Preadapted to adapt: underpinnings of adaptive plasticity revealed by the downy brome genome. Commun Biol. 2023 Mar 27, 6(1): 326. |
| 1221 | *Casuarina cunninghamiana* *(Casuarinaceae)* | 2023-03-27 | Zhang Y et al., Chromosome-scale de novo genome assembly and annotation of three representative Casuarina species: C. equisetifolia, C. glauca, and C. cunninghamiana. Plant J. 2023 Mar 27. Online ahead of print. |
| 1220 | *Alnus rubra* *(Betulaceae)* | 2023-03-26 | Hixson KK et al., Annotated genome sequence of a fast-growing diploid clone of red alder (Alnus rubra Bong.). G3 (Bethesda). 2023 Mar 26: jkad060. Online ahead of print. |
| 1219 | *Quillaja saponaria* *(Quillajaceae)* | 2023-03-24 | Reed J et al., Elucidation of the pathway for biosynthesis of saponin adjuvants from the soapbark tree. Science. 2023 Mar 24, 379(6638): 1252-1264. Epub 2023 Mar 23. |
| 1218 | *Rhododendron irroratum* *(Ericaceae)* | 2023-03-21 | Wu X et al., Evolutionary history of two evergreen Rhododendron species as revealed by chromosome-level genome assembly. Front Plant Sci. 2023 Mar 21, 14: 1123707. eCollection 2023. |
| 1217 | *Costus lasius* *(Costaceae)* | 2023-03-16 | Harencar J et al., Genome assemblies and comparison of two neotropical spiral gingers: Costus pulverulentus and C. lasius. J Hered. 2023 Mar 16: esad018. Online ahead of print. |
| 1216 | *Costus pulverulentus* *(Costaceae)* | 2023-03-16 | Harencar J et al., Genome assemblies and comparison of two neotropical spiral gingers: Costus pulverulentus and C. lasius. J Hered. 2023 Mar 16: esad018. Online ahead of print. |
| 1215 | *Atropa belladonna* *(Solanaceae)* | 2023-03-15 | Zhang F et al., Revealing evolution of tropane alkaloid biosynthesis by analyzing two genomes in the Solanaceae family. Nat Commun. 14, 1446 (2023). |
| 1214 | *Reynoutria multiflora* *(Polygonaceae)* | 2023-03-15 | Zhao Y et al., The first chromosome-level Fallopia multiflora genome assembly provides insights into stilbene biosynthesis. Hortic Res. 2023 Mar 15, 10(3): uhad047. eCollection 2023 |
| 1213 | *Torreya grandis* *(Taxaceae)* | 2023-03-10 | Lou H et al., The Torreya grandis genome illuminates the origin and evolution of gymnosperm-specific sciadonic acid biosynthesis. Nat Commun. 2023 Mar 10, 14(1): 1315. |
| 1212 | *Lindera megaphylla* *(Lauraceae)* | 2023-03-08 | Tian XC et al., Unique gene duplications and conserved microsynteny potentially associated with resistance to wood decay in the Lauraceae. Front Plant Sci. 2023 Mar 8: 14: 1122549. eCollection 2023. |
| 1211 | *Vicia faba* *(Fabaceae)* | 2023-03-08 | Jayakodi M et al., The giant diploid faba genome unlocks variation in a global protein crop. Nature. 2023 Mar 8. Online ahead of print. |
| 1210 | *Actinidia arguta* *(Actinidiaceae)* | 2023-03-06 | Akagi T et al., Recurrent neo-sex chromosome evolution in kiwifruit. Nat Plants. 2023 Mar 6. Online ahead of print. |
| 1209 | *Actinidia polygama* *(Actinidiaceae)* | 2023-03-06 | Akagi T et al., Recurrent neo-sex chromosome evolution in kiwifruit. Nat Plants. 2023 Mar 6. Online ahead of print. |
| 1208 | *Actinidia rufa* *(Actinidiaceae)* | 2023-03-06 | Akagi T et al., Recurrent neo-sex chromosome evolution in kiwifruit. Nat Plants. 2023 Mar 6. Online ahead of print. |
| 1207 | *Petrea volubilis* *(Verbenaceae)* | 2023-03-03 | Hamilton JP et al., Chromosome-scale assembly of the Verbenaceae species Queen's Wreath (Petrea volubilis L.). BMC Genom Data. 2023 Mar 3, 24(1): 14. |
| 1206 | *Tetrastigma hemsleyanum* *(Vitaceae)* | 2023-03-02 | Zhu S et al., Chromosome-level reference genome of Tetrastigma hemsleyanum (Vitaceae) provides insights into genomic evolution and the biosynthesis of phenylpropanoids and flavonoids. Plant J. 2023 Mar 2. Online ahead of print. |
| 1205 | *Commiphora wightii* *(Burseraceae)* | 2023-02-28 | Banerjee RP et al., De Novo Hybrid Assembled Draft Genome of Commiphora wightii (Arnott) Bhandari Reveals Key Enzymes Involved in Phytosterol Biosynthesis. Life 2023, 13(3), 662 |
| 1204 | *Annona cherimola* *(Annonaceae)* | 2023-02-21 | Talavera A et al., Genomics in neglected and underutilized fruit crops: A chromosome-scale genome sequence of cherimoya (Annona cherimola). Plants People Planet. 2023: 1-16 |
| 1203 | *Cymbidium crassifolium* *(Orchidaceae)* | 2023-02-21 | Fan W et al., High-quality Cymbidium mannii genome and multifaceted regulation of crassulacean acid metabolism in epiphytes. Plant Commun. 2023 Feb 21, 100564. Online ahead of print. |
| 1202 | *Cyperus esculentus* *(Cyperaceae)* | 2023-02-21 | Zhao X et al., Chromosome-scale genome assembly of the yellow nutsedge (Cyperus esculentus). Genome Biol Evol. 2023 Feb 21, evad027. Online ahead of print. |
| 1201 | *Musa beccarii* *(Musaceae)* | 2023-02-21 | Wang ZF et al., Genome assembly of Musa beccarii shows extensive chromosomal rearrangements and genome expansion during evolution of Musaceae genomes. Gigascience. 2023 Feb 21, 12: giad005. |
| 1200 | *Oldenlandia corymbosa* *(Rubiaceae)* | 2023-02-21 | Julca I et al., Genomic, transcriptomic, and metabolomic analysis of Oldenlandia corymbosa reveals the biosynthesis and mode of action of anti-cancer metabolites. J Integr Plant Biol. 2023 Feb 21. Online ahead of print. |
| 1199 | *Acacia pachyceras* *(Fabaceae)* | 2023-02-16 | Habibi N et al., Genome survey and genetic characterization of Acacia pachyceras O. Schwartz. Front Plant Sci. 2023 Feb 16, 14: 1062401. eCollection 2023. |
| 1198 | *Quercus dentata* *(Fagaceae)* | 2023-02-15 | Wang WB et al., Chromosome-scale genome assembly and insights into the metabolome and gene regulation of leaf color transition in an important oak species, Quercus dentata. New Phytol. 2023 Feb 15. Online ahead of print. |
| 1197 | *Citrus x limon* *(Rutaceae)* | 2023-02-14 | Bao Y et al., A gap-free and haplotype-resolved lemon genome provides insights into flavor synthesis and Huanglongbing (HLB) tolerance. Hortic Res. 2023 Feb 14: uhad020. |
| 1196 | *Prunus campanulata* *(Rosaceae)* | 2023-02-14 | Nie C et al., Genome assembly, resequencing and genome-wide association analyses provide novel insights into the origin, evolution and flower colour variations of flowering cherry. Plant J. 2023 Feb 14. Online ahead of print. |
| 1195 | *Magnolia obovata* *(Magnoliaceae)* | 2023-02-10 | Zhou L et al., The genome of Magnolia hypoleuca provides a new insight into cold tolerance and the evolutionary position of magnoliids. Front Plant Sci. 2023 Feb 10, 14: 1108701. eCollection 2023. |
| 1194 | *Euphorbia peplus* *(Euphorbiaceae)* | 2023-02-09 | Johnson AR et al., Chromosome-level genome assembly of Euphorbia peplus, a model system for plant latex, reveals that relative lack of Ty3 transposons contributed to its small genome size. Genome Biol Evol. 2023 Feb 9, evad018. Online ahead of print. |

| | plant species | published | |
|---|---|---|---|
| 1193 | *Sphagnum angustifolium* *(Sphagnaceae)* | 2023-02-06 | Healey AL et al., Newly identified sex chromosomes in the Sphagnum (peat moss) genome alter carbon sequestration and ecosystem dynamics. Nat Plants. (2023). |
| 1192 | *Sphagnum divinum* *(Sphagnaceae)* | 2023-02-06 | Healey AL et al., Newly identified sex chromosomes in the Sphagnum (peat moss) genome alter carbon sequestration and ecosystem dynamics. Nat Plants. (2023). |
| 1191 | *Salix integra* *(Salicaceae)* | 2023-02-02 | Hyden B et al., De Novo Assembly and Annotation of 11 Diverse Shrub Willow ( Salix) Genomes Reveals Novel Gene Organization in Sex-Linked Regions. Int J Mol Sci. 2023 Feb 2, 24(3): 2904. |
| 1190 | *Salix koriyanagi* *(Salicaceae)* | 2023-02-02 | Hyden B et al., De Novo Assembly and Annotation of 11 Diverse Shrub Willow ( Salix) Genomes Reveals Novel Gene Organization in Sex-Linked Regions. Int J Mol Sci. 2023 Feb 2, 24(3): 2904. |
| 1189 | *Salix purpurea* *(Salicaceae)* | 2023-02-02 | Hyden B et al., De Novo Assembly and Annotation of 11 Diverse Shrub Willow ( Salix) Genomes Reveals Novel Gene Organization in Sex-Linked Regions. Int J Mol Sci. 2023 Feb 2, 24(3): 2904. |
| 1188 | *Salix udensis* *(Salicaceae)* | 2023-02-02 | Hyden B et al., De Novo Assembly and Annotation of 11 Diverse Shrub Willow ( Salix) Genomes Reveals Novel Gene Organization in Sex-Linked Regions. Int J Mol Sci. 2023 Feb 2, 24(3): 2904. |
| 1187 | *Codonopsis lanceolata* *(Campanulaceae)* | 2023-02-01 | Jang W et al., The chromosome-level genome assembly of lance asiabell (Codonopsis lanceolata), a medicinal and vegetable plant of the Campanulaceae family. Front Genet. 2023 Feb 1, 14: 1100819. eCollection 2023. |
| 1186 | *Melastoma malabathricum* *(Melastomataceae)* | 2023-01-27 | Zhong Y et al., Chromosomal-level genome assembly of Melastoma candidum provides insights into trichome evolution. Front Plant Sci. 2023 Jan 27, 14: 1126319. eCollection 2023. |
| 1185 | *Plantago ovata* *(Plantaginaceae)* | 2023-01-27 | Herliana L et al., A chromosome-level genome assembly of Plantago ovata. Sci Rep. 2023 Jan 27, 13(1): 1528. |
| 1184 | *Vitis adenoclada* *(Vitaceae)* | 2023-01-25 | Cheng G et al., Chromosome-scale genomics, metabolomics, and transcriptomics provide insight into the synthesis and regulation of phenols in Vitis adenoclada grapes. Front Plant Sci. 2023 Jan 25, 14: 1124046. eCollection 2023. |
| 1183 | *Cleome gynandra* *(Cleomaceae)* | 2023-01-24 | Hoang NV et al., The Gynandropsis gynandra genome provides insights into whole-genome duplications and the evolution of C4 photosynthesis in Cleomaceae. Plant Cell. 2023 Jan 24: koad018. Online ahead of print. |
| 1182 | *Medakamo hakoo* *(Coccomyxaceae)* | 2023-01-23 | Kato S et al., Genomic analysis of an ultrasmall freshwater green alga, Medakamo hakoo. Commun Biol. 2023 Jan 23, 6(1): 89. |
| 1181 | *Coleus barbatus* *(Lamiaceae)* | 2023-01-20 | Bryson AE et al., Uncovering a miltiradiene biosynthetic gene cluster in the Lamiaceae reveals a dynamic evolutionary trajectory. Nat Commun. 2023 Jan 20, 14(1): 343. |
| 1180 | *Leonotis leonurus* *(Lamiaceae)* | 2023-01-20 | Bryson AE et al., Uncovering a miltiradiene biosynthetic gene cluster in the Lamiaceae reveals a dynamic evolutionary trajectory. Nat Commun. 2023 Jan 20, 14(1): 343. |
| 1179 | *Prunella vulgaris* *(Lamiaceae)* | 2023-01-20 | Bryson AE et al., Uncovering a miltiradiene biosynthetic gene cluster in the Lamiaceae reveals a dynamic evolutionary trajectory. Nat Commun. 2023 Jan 20, 14(1): 343. |
| 1178 | *Chenopodium berlandieri* *(Amaranthaceae)* | 2023-01-19 | Samuels ME et al., Genomic Sequence of Canadian Chenopodium berlandieri: A North American Wild Relative of Quinoa. Plants (Basel). 2023 Jan 19, 12(3): 467. |
| 1177 | *Rhodomyrtus tomentosa* *(Myrtaceae)* | 2023-01-19 | Li F et al., Gap-free genome assembly and comparative analysis reveal the evolution and anthocyanin accumulation mechanism of Rhodomyrtus tomentosa. Hortic Res. 2023 Jan 19, 10(3): uhad005. eCollection 2023 Mar. |
| 1176 | *Cardamine occulta* *(Brassicaceae)* | 2023-01-18 | Li LZ et al., Common evolutionary trajectory of short life-cycle in Brassicaceae ruderal weeds. Nat Commun. 14, 290 (2023). |
| 1175 | *Herpetospermum pedunculosum* *(Cucurbitaceae)* | 2023-01-18 | Yang Y et al., Chromosome-Level Genome Assembly of Herpetospermum pedunculosum (Cucurbitaceae). Genome Biol Evol. 2023 Jan 18: evad005. Online ahead of print. |
| 1174 | *Lathyrus sativus* *(Fabaceae)* | 2023-01-17 | Rajarammohan S et al., Genome sequencing and assembly of Lathyrus sativus - a nutrient-rich hardy legume crop. Sci Data. 2023 Jan 17, 10(1): 32. |
| 1173 | *Turnera subulata* *(Passifloraceae)* | 2023-01-07 | Henning PM et al., Annotation of the Turnera subulata (Passifloraceae) Draft Genome Reveals the S-Locus Evolved after the Divergence of Turneroideae from Passifloroideae in a Stepwise Manner. Plants (Basel). 2023 Jan 7, 12(2): 286. |
| 1172 | *Nepeta tenuifolia* *(Lamiaceae)* | 2023-01-06 | Liu C et al., A chromosome-level genome assembly reveals that a bipartite gene cluster formed via an inverted duplication controls monoterpenoid biosynthesis in Japanese catnip. Mol Plant. 2023 Jan 6: S1674-2052(23)00004-7. Online ahead of print. |
| 1171 | *Saururus chinensis* *(Saururaceae)* | 2023-01-05 | Xue JY et al., The Saururus chinensis genome provides insights into the evolution of pollination strategies and herbaceousness in magnoliids. Plant J. 2023 Mar, 113(5): 1021-1034. |
| 1170 | *Cyanidium caldarium* *(Cyanidiaceae)* | 2023-01-04 | Cho CH et al., Genome-wide signatures of adaptation to extreme environments in red algae. Nat Commun. 14, 10 (2023). |
| 1169 | *Actinidia latifolia* *(Actinidiaceae)* | 2022-12-31 | Han X et al., Two haplotype-resolved, gap-free genome assemblies of Actinidia latifolia and Actinidia chinensis shed light on regulation mechanisms of vitamin C and sucrose metabolism in kiwifruit. Mol Plant. 2022 Dec 31, S1674-2052(22)00476-2. Online ahead of print. |
| 1168 | *Pinellia pedatisecta* *(Araceae)* | 2022-12-30 | Qian Z et al., The high-quality Pinellia pedatisecta genome reveals a key role of tandem duplication in the expansion of its agglutinin genes. Hortic Res. 2022 Dec 30: uhac289. eCollection 2022 |
| 1167 | *Oenanthe linearis* *(Apiaceae)* | 2022-12-28 | Liu JX et al., High-quality genome sequence reveals a young polyploidization and provides insights into cellulose and lignin biosynthesis in water dropwort (Oenanthe sinensis). Ind Crops Prod. 2023 Mar, 193, 116203 Epub 2022 Dec 28. |
| 1166 | *Poa annua* *(Poaceae)* | 2022-12-28 | Robbins MD et al., Chromosome-scale Genome Assembly and Annotation of Allotetraploid Annual Bluegrass (Poa annua L.) Genome Biol Evol. 2022 Dec 28: evac180. Online ahead of print. |
| 1165 | *Marsdenia tenacissima* *(Apocynaceae)* | 2022-12-27 | Zhou Y et al., The genome of Marsdenia tenacissima provides insights into calcium adaptation and tenacissoside biosynthesis. Plant J. 2022 Dec 27. Online ahead of print. |
| 1164 | *Melia azedarach* *(Meliaceae)* | 2022-12-26 | Cui G et al., Meliaceae genomes provide insights into wood development and limonoids biosynthesis. Plant Biotechnol J. 2023 Mar, 21(3): 574-590. Epub 2022 Dec 26. |
| 1163 | *Cornus controversa* *(Cornaceae)* | 2022-12-15 | Dong C et al., Karyotype evolution of the Asterids insights from the first genome sequences of the family Cornaceae. DNA Res. 2023 Feb 1, 30(1): dsac051. |
| 1162 | *Isodon rubescens* *(Lamiaceae)* | 2022-12-13 | Sun Y et al., A chromosome-level genome of Isodon rubescens reveals that tandem-duplicated CYP706V oxidase genes control oridonin biosynthesis in shoot apex. Mol Plant. 2022 Dec 13: S1674-2052(22)00442-7. Online ahead of print. |
| 1161 | *Paspalum vaginatum* *(Poaceae)* | 2022-12-13 | Sun G et al., Genome of Paspalum vaginatum and the role of trehalose mediated autophagy in increasing maize biomass. Nat Commun. 2022 Dec 13, 13: 7731. |
| 1160 | *Artocarpus hirsutus* *(Moraceae)* | 2022-12-12 | Patil AB et al., Jack of all trades: Genome assembly of Wild Jack and comparative genomics of Artocarpus. Front Plant Sci. 2022 Dec 12, 13: 1029540. eCollection 2022. |
| 1159 | *Carex myosuroides* *(Cyperaceae)* | 2022-12-12 | Ning Y et al., The chromosome-scale genome of Kobresia myosuroides sheds light on karyotype evolution and recent diversification of a dominant herb group on the Qinghai-Tibet Plateau. DNA Res. 2022 Dec 12: dsac049. Online ahead of print. |
| 1158 | *Amaranthus tricolor* *(Amaranthaceae)* | 2022-12-06 | Wang H et al., Chromosome-scale Amaranthus tricolor genome provides insights into the evolution of the genus Amaranthus and the mechanism of betalain biosynthesis. DNA Res. 2022 Dec 6: dsac050. Online ahead of print. |

| | plant species | published | |
|---|---|---|---|
| 1157 | *Gossypium sturtianum* *(Malvaceae)* | 2022-12-06 | Wang M et al., Genomic innovation and regulatory rewiring during evolution of the cotton genus Gossypium. Nat Genet. 2022 Dec 6. Online ahead of print. |
| 1156 | *Sophora moorcroftiana* *(Fabaceae)* | 2022-12-06 | Yin X et al., Sophora moorcroftiana genome analysis suggests association between sucrose metabolism and drought adaptation. Plant Physiol. 2022 Dec 6: kiac558. Online ahead of print. |
| 1155 | *Acer palmatum* *(Sapindaceae)* | 2022-12-02 | Chen Z et al., Chromosomal-level genome and multi-omics dataset provides new insights into leaf pigmentation in Acer palmatum. Int J Biol Macromol. 2022 Dec 2: S0141-8130(22)02868-9. Online ahead of print. |
| 1154 | *Rhododendron prattii* *(Ericaceae)* | 2022-12-02 | Ma Y et al., Pervasive hybridization during evolutionary radiation of Rhododendron subgenus Hymenanthes in mountains of southwest China. Natl Sci Rev. 2022 Dec 2, 9(12): nwac276. eCollection 2022 Dec. |
| 1153 | *Alopecurus myosuroides* *(Poaceae)* | 2022-12-01 | Cai L et al., The blackgrass genome reveals patterns of non-parallel evolution of polygenic herbicide resistance. New Phytol. 2022 Dec 1. Online ahead of print. |
| 1152 | *Paeonia ostii* *(Paeoniaceae)* | 2022-11-28 | Yuan J et al., Genomic basis of the giga-chromosomes and giga-genome of tree peony Paeonia ostii. Nat Commun. 2022 Nov 28, 13(1): 7328. |
| 1151 | *Lonicera maackii* *(Caprifoliaceae)* | 2022-11-26 | Kesel E et al., Whole-Genome Sequence, Assembly and Annotation of an Invasive Plant, Lonicera maackii (Amur Honeysuckle). Plants (Basel). 2022 Nov 26, 11(23): 3253 |
| 1150 | *Cremastra appendiculata* *(Orchidaceae)* | 2022-11-25 | Wang J et al., A draft genome of the medicinal plant Cremastra appendiculata (D. Don) provides insights into the colchicine biosynthetic pathway. Commun Biol. 5, 1294 (2022). |
| 1149 | *Protea cynaroides* *(Proteaceae)* | 2022-11-24 | Chang J et al., The Genome of the King Protea, Protea cynaroides. Plant J. 2022 Nov 24. Online ahead of print. |
| 1148 | *Rhamphospermum arvense* *(Brassicaceae)* | 2022-11-24 | Yang T et al., Sinapis genomes provide insights into whole-genome triplication (WGT) and divergence patterns within tribe Brassiceae. Plant J. 2022 Nov 24. Online ahead of print. |
| 1147 | *Epimedium pubescens* *(Berberidaceae)* | 2022-11-14 | Shen G et al., The discovery of a key prenyltransferase gene assisted by a chromosome-level Epimedium pubescens genome. Front Plant Sci. 2022 Nov 14, 13: 1034943. eCollection 2022. |
| 1146 | *Phyllanthus cochinchinensis* *(Phyllanthaceae)* | 2022-11-09 | Zhang W et al., The first high-quality chromosome-level genome assembly of Phyllanthaceae (Phyllanthus cochinchinensis) provides insights into flavonoid biosynthesis. Planta. 2022 Nov 9, 256(6): 109. |
| 1145 | *Vitis x labruscana* *(Vitaceae)* | 2022-11-07 | Shirasawa K et al., De novo whole-genome assembly in an interspecific hybrid table grape, Shine Muscat. DNA Res. 2022 Nov 7: dsac040. Online ahead of print. |
| 1144 | *Allium fistulosum* *(Amaryllidaceae)* | 2022-11-05 | Liao N et al., Chromosome-level genome assembly of bunching onion illuminates genome evolution and flavor formation in Allium crops. Nat Commun. 2022 Nov 5, 13(1): 6690. |
| 1143 | *Rubus argutus* *(Rosaceae)* | 2022-11-04 | Bruna T et al., A chromosome-length genome assembly and annotation of blackberry (Rubus argutus, cv. Hillquist). G3 (Bethesda). 2022 Nov 4: jkac289. Online ahead of print. |
| 1142 | *Pistacia chinensis* *(Anacardiaceae)* | 2022-11-02 | Hegde SN et al., De novo genome assembly and annotation of gall-forming medicinal plant Pistacia chinensis subsp. integerrima (J. L. Stewart ex Brandis) Rech. f. J Genet. 2022, 101: 51 |
| 1141 | *Lolium rigidum* *(Poaceae)* | 2022-11-01 | Paril J et al., Rounding up the annual ryegrass genome: High-quality reference genome of Lolium rigidum. Front Genet. 2022 Nov 1, 13: 1012694. eCollection 2022. |
| 1140 | *Populus suaveolens* *(Salicaceae)* | 2022-11-01 | Sang Y et al., Genomic insights into local adaptation and future climate-induced vulnerability of a keystone forest tree in East Asia. Nat Commun. 2022 Nov 1, 13(1): 6541. |
| 1139 | *Capparis spinosa* *(Capparaceae)* | 2022-10-30 | Wang L et al., The Capparis spinosa var. herbacea genome provides the first genomic instrument for a diversity and evolution study of the Capparaceae family. Gigascience. 2022 Oct 30, 11: giac106. |
| 1138 | *Anacardium occidentale* *(Anacardiaceae)* | 2022-10-28 | Savadi S et al., De novo assembly and characterization of the draft genome of the cashew (Anacardium occidentale L.). Sci Rep. 2022 Oct 28, 12(1): 18187. |
| 1137 | *Vitis berlandieri x riparia* *(Vitaceae)* | 2022-10-28 | Minio A et al., HiFi chromosome-scale diploid assemblies of the grape rootstocks 110R, Kober 5BB, and 101-14 Mgt. Sci Data. 2022 Oct 28, 9(1): 660. |
| 1136 | *Vitis berlandieri x rupestris* *(Vitaceae)* | 2022-10-28 | Minio A et al., HiFi chromosome-scale diploid assemblies of the grape rootstocks 110R, Kober 5BB, and 101-14 Mgt. Sci Data. 2022 Oct 28, 9(1): 660. |
| 1135 | *Vitis rupestris x riparia* *(Vitaceae)* | 2022-10-28 | Minio A et al., HiFi chromosome-scale diploid assemblies of the grape rootstocks 110R, Kober 5BB, and 101-14 Mgt. Sci Data. 2022 Oct 28, 9(1): 660. |
| 1134 | *Astragalus mongholicus* *(Fabaceae)* | 2022-10-27 | Chen Y et al., A reference-grade genome assembly for Astragalus mongholicus and insights into the biosynthesis and high accumulation of triterpenoids and flavonoids in its roots. Plant Commun. 2022 Oct 27, 100469. Online ahead of print. |
| 1133 | *Dipterocarpus alatus* *(Dipterocarpaceae)* | 2022-10-27 | Tian Z et al., Thirteen Dipterocarpoideae genomes provide insights into their evolution and borneol biosynthesis. Plant Commun. 2022 Oct 27, 100464. Online ahead of print. |
| 1132 | *Dipterocarpus gracilis* *(Dipterocarpaceae)* | 2022-10-27 | Tian Z et al., Thirteen Dipterocarpoideae genomes provide insights into their evolution and borneol biosynthesis. Plant Commun. 2022 Oct 27, 100464. Online ahead of print. |
| 1131 | *Dipterocarpus intricatus* *(Dipterocarpaceae)* | 2022-10-27 | Tian Z et al., Thirteen Dipterocarpoideae genomes provide insights into their evolution and borneol biosynthesis. Plant Commun. 2022 Oct 27, 100464. Online ahead of print. |
| 1130 | *Dipterocarpus zeylanicus* *(Dipterocarpaceae)* | 2022-10-27 | Tian Z et al., Thirteen Dipterocarpoideae genomes provide insights into their evolution and borneol biosynthesis. Plant Commun. 2022 Oct 27, 100464. Online ahead of print. |
| 1129 | *Hopea mollissima* *(Dipterocarpaceae)* | 2022-10-27 | Tian Z et al., Thirteen Dipterocarpoideae genomes provide insights into their evolution and borneol biosynthesis. Plant Commun. 2022 Oct 27, 100464. Online ahead of print. |
| 1128 | *Hopea odorata* *(Dipterocarpaceae)* | 2022-10-27 | Tian Z et al., Thirteen Dipterocarpoideae genomes provide insights into their evolution and borneol biosynthesis. Plant Commun. 2022 Oct 27, 100464. Online ahead of print. |
| 1127 | *Shorea henryana* *(Dipterocarpaceae)* | 2022-10-27 | Tian Z et al., Thirteen Dipterocarpoideae genomes provide insights into their evolution and borneol biosynthesis. Plant Commun. 2022 Oct 27, 100464. Online ahead of print. |
| 1126 | *Shorea roxburghii* *(Dipterocarpaceae)* | 2022-10-27 | Tian Z et al., Thirteen Dipterocarpoideae genomes provide insights into their evolution and borneol biosynthesis. Plant Commun. 2022 Oct 27, 100464. Online ahead of print. |
| 1125 | *Shorea wangtianshuea* *(Dipterocarpaceae)* | 2022-10-27 | Tian Z et al., Thirteen Dipterocarpoideae genomes provide insights into their evolution and borneol biosynthesis. Plant Commun. 2022 Oct 27, 100464. Online ahead of print. |
| 1124 | *Vatica guangxiensis* *(Dipterocarpaceae)* | 2022-10-27 | Tian Z et al., Thirteen Dipterocarpoideae genomes provide insights into their evolution and borneol biosynthesis. Plant Commun. 2022 Oct 27, 100464. Online ahead of print. |
| 1123 | *Vatica odorata* *(Dipterocarpaceae)* | 2022-10-27 | Tian Z et al., Thirteen Dipterocarpoideae genomes provide insights into their evolution and borneol biosynthesis. Plant Commun. 2022 Oct 27, 100464. Online ahead of print. |
| 1122 | *Vatica rassak* *(Dipterocarpaceae)* | 2022-10-27 | Tian Z et al., Thirteen Dipterocarpoideae genomes provide insights into their evolution and borneol biosynthesis. Plant Commun. 2022 Oct 27, 100464. Online ahead of print. |
| 1121 | *Voacanga thouarsii* *(Apocynaceae)* | 2022-10-27 | Cuello C et al., Genome assembly of the medicinal plant Voacanga thouarsii. Genome Biol Evol. 2022 Oct 27: evac158. Online ahead of print. |

| | plant species | published | |
|---|---|---|---|
| 1120 | *Populus wilsonii* (*Salicaceae*) | 2022-10-25 | Li C et al., Chromosome-scale genome assembly provides insights into the molecular mechanisms of tissue development of Populus wilsonii. Commun Biol 5, 1125 (2022). |
| 1119 | *Physalis grisea* (*Solanaceae*) | 2022-10-21 | He J et al., Establishing Physalis as a Solanaceae model system enables genetic reevaluation of the inflated calyx syndrome. Plant Cell. 2022 Oct 21, koac305. Online ahead of print |
| 1118 | *Physalis pruinosa* (*Solanaceae*) | 2022-10-21 | He J et al., Establishing Physalis as a Solanaceae model system enables genetic reevaluation of the inflated calyx syndrome. Plant Cell. 2022 Oct 21, koac305. Online ahead of print |
| 1117 | *Aesculus chinensis* (*Sapindaceae*) | 2022-10-18 | Ye L et al., The Chromosome-level genome of Aesculus wilsonii provides new insights into terpenoid biosynthesis and Aesculus evolution. Front Plant Sci. 2022 Oct 18, eCollection 2022. |
| 1116 | *Angelica sinensis* (*Apiaceae*) | 2022-10-18 | Han X et al., The chromosome-level genome of female ginseng (Angelica sinensis) provides insights into molecular mechanisms and evolution of coumarin biosynthesis. Plant J. 2022 Oct 18. Online ahead of print. |
| 1115 | *Sesamum schinzianum* (*Pedaliaceae*) | 2022-10-18 | Wang X et al., The Wild Allotetraploid Sesame Genome Provides Novel Insights into Evolution and Lignan Biosynthesis. J Adv Res. 2022 Oct 17: S2090-1232(22)00233-8. Online ahead of print. |
| 1114 | *Cinnamomum burmannii* (*Lauraceae*) | 2022-10-15 | Li F et al., Genome assembly provided new insights into the Cinnamomum burmannii evolution and D-borneol biosynthesis differences between chemotypes. Ind Crops Prod. 2022 Oct, 186: 115181. Epub 2022 Oct 15. |
| 1113 | *Acacia pycnantha* (*Fabaceae*) | 2022-10-14 | McLay TGB et al., A genome resource for Acacia, Australia's largest plant genus. PLoS One. 2022 Oct 14, 17(10): e0274267. eCollection 2022. |
| 1112 | *Hirschfeldia incana* (*Brassicaceae*) | 2022-10-14 | Garassino F et al., The genome sequence of Hirschfeldia incana, a new Brassicaceae model to improve photosynthetic light-use efficiency. Plant J. 2022 Oct 14. Online ahead of print. |
| 1111 | *Cinchona pubescens* (*Rubiaceae*) | 2022-10-06 | Canales NA et al., A highly contiguous, scaffold-level nuclear genome assembly for the fever tree (Cinchona pubescens Vahl) as a novel resource for Rubiaceae research. Gigabyte. 2022: 71 |
| 1110 | *Vinca minor* (*Apocynaceae*) | 2022-10-06 | Stander EA et al., The Vinca minor genome highlights conserved evolutionary traits in monoterpene indole alkaloid synthesis. G3 (Bethesda). 2022 Oct 6, jkac268. Online ahead of print. |
| 1109 | *Gentiana macrophylla* (*Gentianaceae*) | 2022-10-05 | Zhou T et al., De novo genome assembly of the medicinal plant Gentiana macrophylla provides insights into the genomic evolution and biosynthesis of iridoids. DNA Res. 2022 Oct 5: dsac034. Online ahead of print. |
| 1108 | *Glebionis coronaria* (*Asteraceae*) | 2022-10-05 | Wang S et al., Chromosome-level genome of a leaf vegetable Glebionis coronaria provides insights into the biosynthesis of monoterpenoids contributing to its special aroma. DNA Res. 2022 Oct 5: dsac036. Online ahead of print. |
| 1107 | *Buxus austroyunnanensis* (*Buxaceae*) | 2022-10-04 | Wang Z et al., A high-quality Buxus austro-yunnanensis (Buxales) genome provides new insights into karyotype evolution in early eudicots. BMC Biol. 2022 Oct 4, 20(1): 216. |
| 1106 | *Galdieria partita* (*Galdieriaceae*) | 2022-10-04 | Hirooka S et al., Life cycle and functional genomics of the unicellular red alga Galdieria for elucidating algal and plant evolution and industrial use. Proc Natl Acad Sci U S A. 2022 Oct 11, 119(41): e2210665119. Epub 2022 Oct 4. |
| 1105 | *Vigna umbellata* (*Fabaceae*) | 2022-10-04 | Kaul T et al., De novo genome assembly of rice bean (Vigna umbellata) - A nominated nutritionally rich future crop reveals novel insights into flowering potential, habit, and palatability centric - traits for efficient domestication. Front Plant Sci. 2022 Oct 04, eCollection 2022. |
| 1104 | *Dracaena cochinchinensis* (*Asparagaceae*) | 2022-10-03 | Xu Y et al., A chromosome-level genome assembly for Dracaena cochinchinesis reveals molecular basis of its longevity and formation of dragons blood. Plant Commun. 2022 Oct 3: 100456. Online ahead of print. |
| 1103 | *Lantana camara* (*Verbenaceae*) | 2022-09-30 | Joshi AG et al., Draft genome sequence of an invasive plant Lantana camara L. Bioinformation. 2022 Sep 30, 18(9): 739-741. eCollection 2022. |
| 1102 | *Primulina eburnea* (*Gesneriaceae*) | 2022-09-30 | Yi H et al., Genomic insights into inter- and intraspecific mating system shifts in Primulina. Mol Ecol. 2022 Sep 30. Online ahead of print |
| 1101 | *Primulina tabacum* (*Gesneriaceae*) | 2022-09-30 | Yi H et al., Genomic insights into inter- and intraspecific mating system shifts in Primulina. Mol Ecol. 2022 Sep 30. Online ahead of print |
| 1100 | *Bursera bipinnata* (*Burseraceae*) | 2022-09-27 | Rico Y et al., Development of Genomic Resources in Mexican Bursera (Section: Bullockia: Burseraceae): Genome Assembly, Annotation, and Marker Discovery for Three Copal Species. Genes (Basel). 2022 Sep 27, 13(10): 1741. |
| 1099 | *Bursera cuneata* (*Burseraceae*) | 2022-09-27 | Rico Y et al., Development of Genomic Resources in Mexican Bursera (Section: Bullockia: Burseraceae): Genome Assembly, Annotation, and Marker Discovery for Three Copal Species. Genes (Basel). 2022 Sep 27, 13(10): 1741. |
| 1098 | *Bursera palmeri* (*Burseraceae*) | 2022-09-27 | Rico Y et al., Development of Genomic Resources in Mexican Bursera (Section: Bullockia: Burseraceae): Genome Assembly, Annotation, and Marker Discovery for Three Copal Species. Genes (Basel). 2022 Sep 27, 13(10): 1741. |
| 1097 | *Quercus gilva* (*Fagaceae*) | 2022-09-23 | Zhou X et al., A chromosome-scale genome assembly of Quercus gilva: Insights into the evolution of Quercus section Cyclobalanopsis (Fagaceae). Front Plant Sci. 2022 Sep 23, 13: 1012277. eCollection 2022. |
| 1096 | *Quercus variabilis* (*Fagaceae*) | 2022-09-23 | Han B et al., A chromosome-level genome assembly of the Chinese cork oak (Quercus variabilis). Front Plant Sci. 2022 Sep 23, 13: 1001583. eCollection 2022. |
| 1095 | *Salix arbutifolia* (*Salicaceae*) | 2022-09-23 | Wang D et al., Repeated turnovers keep sex chromosomes young in willows. Genome Biol. 2022 Sep 23, 23(1): 200. |
| 1094 | *Salix chaenomeloides* (*Salicaceae*) | 2022-09-23 | Wang D et al., Repeated turnovers keep sex chromosomes young in willows. Genome Biol. 2022 Sep 23, 23(1): 200. |
| 1093 | *Gossypium ekmanianum* (*Malvaceae*) | 2022-09-19 | Peng R et al., Evolutionary divergence of duplicated genomes in newly described allotetraploid cottons. Proc Natl Acad Sci U S A. 2022 Sep 27, 119(39): e2208496119. Epub 2022 Sep 19. |
| 1092 | *Gossypium stephensii* (*Malvaceae*) | 2022-09-19 | Peng R et al., Evolutionary divergence of duplicated genomes in newly described allotetraploid cottons. Proc Natl Acad Sci U S A. 2022 Sep 27, 119(39): e2208496119. Epub 2022 Sep 19. |
| 1091 | *Linanthus parryae* (*Polemoniaceae*) | 2022-09-15 | Anghel I et al., Reference genome of the color polymorphic desert annual plant Linanthus parryae. J Hered. 2022 Sep 15, esac052. Online ahead of print. |
| 1090 | *Papilionanthe hookeriana x teres* (*Orchidaceae*) | 2022-09-15 | Lim AH et al., Genome assembly and chemogenomic profiling of National Flower of Singapore Papilionanthe Miss Joaquim Agnes reveals metabolic pathways regulating floral traits. Commun Biol. 2022 Sep 15, 5(1): 967. |
| 1089 | *Thuja plicata* (*Cupressaceae*) | 2022-09-15 | Shalev TJ et al., The western redcedar genome reveals low genetic diversity in a self-compatible conifer. Genome Res. 2022 Sep 15: gr.276358.121. Online ahead of print. |
| 1088 | *Cissus rotundifolia* (*Vitaceae*) | 2022-09-13 | Xin H et al., A genome for Cissus illustrates features underlying the evolutionary success in dry savannas. Hortic Res. 2022 Sep 13, 9: uhac208. eCollection 2022. |
| 1087 | *Syzygium grande* (*Myrtaceae*) | 2022-09-12 | Low YW et al., Genomic insights into rapid speciation within the world's largest tree genus Syzygium. Nat Commun. 2022 Sep 12, 13(1): 5031. |
| 1086 | *Themeda triandra* (*Poaceae*) | 2022-09-12 | Dunning LT et al., Hybridisation and chloroplast capture between distinct Themeda triandra lineages in Australia. Mol Ecol. 2022 Sep 12. Online ahead of print. |
| 1085 | *Aubrieta canescens* (*Brassicaceae*) | 2022-09-10 | Kaya Y et al., Genome-wide characterization of two Aubrieta taxa: Aubrieta canescens subsp. canescens and Au. macrostyla (Brassicaceae). AoB Plants. 2022 Sep 10, 14(5): plac035. eCollection 2022 Oct. |

| | plant species | published | |
|---|---|---|---|
| 1084 | *Linum tenue*<br>*(Linaceae)* | 2022-09-09 | Gutierrez-Valencia J et al., Genomic analyses of the Linum distyly supergene reveal convergent evolution at the molecular level.<br>Curr Biol. 2022 Sep 4: S0960-9822(22)01364-1. Online ahead of print. |
| 1083 | *Ficus benghalensis*<br>*(Moraceae)* | 2022-09-08 | Chakraborty A et al., Genome sequencing and comparative analysis of Ficus benghalensis and Ficus religiosa species reveal evolutionary mechanisms of longevity.<br>iScience. 2022 Sep 8, 25(10): 105100. |
| 1082 | *Ficus religiosa*<br>*(Moraceae)* | 2022-09-08 | Chakraborty A et al., Genome sequencing and comparative analysis of Ficus benghalensis and Ficus religiosa species reveal evolutionary mechanisms of longevity.<br>iScience. 2022 Sep 8, 25(10): 105100. |
| 1081 | *Aechmea fasciata*<br>*(Bromeliaceae)* | 2022-09-07 | Li Z et al., The genome of Aechmea fasciata provides insights into the evolution of tank epiphytic habits and ethylene-induced flowering.<br>Commun Biol. 2022 Sep 7, 5(1): 920. |
| 1080 | *Orychophragmus violaceus*<br>*(Brassicaceae)* | 2022-09-07 | Zhang K et al., The genome of Orychophragmus violaceus provides genomic insights into the evolution of Brassicaceae polyploidization and its distinct traits.<br>Plant Commun. 2022 Sep 7: 100431. Online ahead of print. |
| 1079 | *Wurfbainia villosa*<br>*(Zingiberaceae)* | 2022-09-07 | Yang P et al., Chromosome-level genome assembly and functional characterization of terpene synthases provide new insights into the volatile terpenoid biosynthesis of Wurfbainia villosa.<br>Plant J. 2022 Sep 7. Online ahead of print. |
| 1078 | *Lathyrus tuberosus*<br>*(Fabaceae)* | 2022-09-04 | Flood PJ et al., Draft genome of the aardaker (Lathyrus tuberosus L.), a tuberous legume.<br>BMC Genom Data. 2022 Sep 4, 23(1): 70. |
| 1077 | *Adiantum capillus-veneris*<br>*(Pteridaceae)* | 2022-09-01 | Fang Y et al., The genome of homosporous maidenhair fern sheds light on the euphyllophyte evolution and defences.<br>Nat Plants. 2022 Sep 1. Online ahead of print. |
| 1076 | *Erianthus fulvus*<br>*(Poaceae)* | 2022-08-31 | Qian Z et al., EfGD: the Erianthus fulvus genome database.<br>Database (Oxford). 2022 Aug 31, 2022: baac076. |
| 1075 | *Hordeum spontaneum*<br>*(Poaceae)* | 2022-08-29 | Zhang W et al., Genome architecture and diverged selection shaping pattern of genomic differentiation in wild barley.<br>Plant Biotechnol J. 2022 Aug 29. Online ahead of print. |
| 1074 | *Gypsophila paniculata*<br>*(Caryophyllaceae)* | 2022-08-24 | Li F et al., The chromosome-level genome of Gypsophila paniculata reveals the molecular mechanism of floral development and ethylene insensitivity.<br>Hortic Res. 2022 Aug 24, 9: uhac176. eCollection 2022. |
| 1073 | *Musa troglodytarum*<br>*(Musaceae)* | 2022-08-24 | Li Z et al., The Musa troglodytarum L. genome provides insights into the mechanism of non-climacteric behaviour and enrichment of carotenoids.<br>BMC Biol. 2022 Aug 24, 20(1): 186. |
| 1072 | *Ilex latifolia*<br>*(Aquifoliaceae)* | 2022-08-22 | Xu KW et al., The chromosome-level holly (Ilex latifolia) genome reveals key enzymes in triterpenoid saponin biosynthesis and fruit color change.<br>Front Plant Sci. 2022 Aug 22, 13: 982323. eCollection 2022. |
| 1071 | *Mucuna pruriens*<br>*(Fabaceae)* | 2022-08-18 | Hao S et al., Chromosomal-level genome of velvet bean (Mucuna pruriens) provides resources for L-DOPA synthetic research and development.<br>DNA Res. 2022 Aug 18: dsac031. Online ahead of print. |
| 1070 | *Camelina hispida*<br>*(Brassicaceae)* | 2022-08-17 | Martin SL et al., Insights from the genomes of four diploid Camelina spp.<br>G3 (Bethesda). 2022 Aug 17: jkac182. Online ahead of print. |
| 1069 | *Camelina laxa*<br>*(Brassicaceae)* | 2022-08-17 | Martin SL et al., Insights from the genomes of four diploid Camelina spp.<br>G3 (Bethesda). 2022 Aug 17: jkac182. Online ahead of print. |
| 1068 | *Camelina neglecta*<br>*(Brassicaceae)* | 2022-08-17 | Martin SL et al., Insights from the genomes of four diploid Camelina spp.<br>G3 (Bethesda). 2022 Aug 17: jkac182. Online ahead of print. |
| 1067 | *Carex cristatella*<br>*(Cyperaceae)* | 2022-08-17 | Planta J et al., Chromosome scale genome assemblies and annotations for Poales species Carex cristatella, Carex scoparia, Juncus effusus and Juncus inflexus.<br>G3 (Bethesda). 2022 Aug 17: jkac211. Online ahead of print. |
| 1066 | *Carex scoparia*<br>*(Cyperaceae)* | 2022-08-17 | Planta J et al., Chromosome scale genome assemblies and annotations for Poales species Carex cristatella, Carex scoparia, Juncus effusus and Juncus inflexus.<br>G3 (Bethesda). 2022 Aug 17: jkac211. Online ahead of print. |
| 1065 | *Juncus inflexus*<br>*(Juncaceae)* | 2022-08-17 | Planta J et al., Chromosome scale genome assemblies and annotations for Poales species Carex cristatella, Carex scoparia, Juncus effusus and Juncus inflexus.<br>G3 (Bethesda). 2022 Aug 17: jkac211. Online ahead of print. |
| 1064 | *Salvia officinalis*<br>*(Lamiaceae)* | 2022-08-16 | Li CY et al., The sage genome provides insight into the evolutionary dynamics of diterpene biosynthesis gene cluster in plants.<br>Cell Rep. 2022 Aug 16, 40(7): 111236. Online ahead of print. |
| 1063 | *Vaccinium caesariense*<br>*(Ericaceae)* | 2022-08-13 | Mengist MF et al., Autopolyploid inheritance and a heterozygous reciprocal translocation shape chromosome genetic behavior in tetraploid blueberry (Vaccinium corymbosum).<br>New Phytol. 2022 Aug 13. Online ahead of print. |
| 1062 | *Adiantum nelumboides*<br>*(Pteridaceae)* | 2022-08-10 | Zhong Y et al., Genomic insights into genetic diploidization in the homosporous fern Adiantum nelumboides.<br>Genome Biol Evol. 2022 Aug 10: evac127. Online ahead of print. |
| 1061 | *Cercis chinensis*<br>*(Fabaceae)* | 2022-08-10 | Li J et al., The nearly complete assembly of Cercis chinensis genome and Fabaceae phylogenomic studies provide insights into new gene evolution.<br>Plant Commun. 2022 Aug 10: 100422. Online ahead of print. |
| 1060 | *Dendrobium hybrid cv. EM*<br>*(Orchidaceae)* | 2022-08-09 | Sherpa R et al., First De novo whole genome sequencing and assembly of mutant Dendrobium hybrid cultivar Emma White.<br>Gigabyte. 2022 Aug 09: gigabyte 066 |
| 1059 | *Gossypium bickii*<br>*(Malvaceae)* | 2022-08-09 | Sheng K et al., A reference-grade genome assembly for Gossypium bickii and insights into its genome evolution and formation of pigment gland and gossypol.<br>Plant Commun. 2022 Aug 9: 100421. Online ahead of print. |
| 1058 | *Larix sibirica*<br>*(Pinaceae)* | 2022-08-06 | Bondar EI et al., Annotation of Siberian Larch (Larix sibirica Ledeb.) nuclear genome - one of the most cold-resistant tree species in the only deciduous genus in Pinaceae.<br>Plants (Basel). 2022 Aug 06, 11(15), 2062 |
| 1057 | *Oxytropis ochrocephala*<br>*(Fabaceae)* | 2022-08-06 | Zhang L et al., Assembly of high-quality genomes of the locoweed Oxytropis ochrocephala and its endophyte Alternaria oxytropis provides new evidence for their symbiotic relationship and swainsonine biosynthesis.<br>Mol Ecol Resour. 2022 Aug 6. Online ahead of print. |
| 1056 | *Triadica sebifera*<br>*(Euphorbiaceae)* | 2022-08-04 | Luo J et al., The chromosome-scale genome sequence of Triadica sebifera provides insight into fatty acids and anthocyanin biosynthesis.<br>Commun Biol. 2022 Aug 4, 5(1): 786. |
| 1055 | *Juncus effusus*<br>*(Juncaceae)* | 2022-08-03 | Hofstatter PG et al., Repeat-based holocentromeres influence genome architecture and karyotype evolution.<br>Cell. 2022 Aug 3: S0092-8674(22)00797-8. Online ahead of print. |
| 1054 | *Rhynchospora breviuscula*<br>*(Cyperaceae)* | 2022-08-03 | Hofstatter PG et al., Repeat-based holocentromeres influence genome architecture and karyotype evolution.<br>Cell. 2022 Aug 3: S0092-8674(22)00797-8. Online ahead of print. |
| 1053 | *Rhynchospora pubera*<br>*(Cyperaceae)* | 2022-08-03 | Hofstatter PG et al., Repeat-based holocentromeres influence genome architecture and karyotype evolution.<br>Cell. 2022 Aug 3: S0092-8674(22)00797-8. Online ahead of print. |
| 1052 | *Rhynchospora tenuis*<br>*(Cyperaceae)* | 2022-08-03 | Hofstatter PG et al., Repeat-based holocentromeres influence genome architecture and karyotype evolution.<br>Cell. 2022 Aug 3: S0092-8674(22)00797-8. Online ahead of print. |
| 1051 | *Stellera chamaejasme*<br>*(Thymelaeaceae)* | 2022-08-03 | Hu H et al., Genomic divergence of Stellera chamaejasme through local selection across the Qinghai-Tibet plateau and northern China.<br>Mol Ecol. 2022 Sep, 31(18): 4782-4796. Epub 2022 Aug 3. |
| 1050 | *Monoraphidium minutum*<br>*(Selenastraceae)* | 2022-08-01 | Calhoun S et al., Multi-omics profiling of the cold tolerant Monoraphidium minutum 26B-AM in response to abiotic stress.<br>Algal Res. 2022 Jul 66: 102794 |
| 1049 | *Prosopis cineraria*<br>*(Fabaceae)* | 2022-07-31 | Sudalaimuthuasari N et al., The Genome of the Mimosoid Legume Prosopis cineraria, a Desert Tree.<br>Int J Mol Sci. 2022, 23, 8503. |

| | plant species | published | |
|---|---|---|---|
| 1048 | *Canna indica* *(Cannaceae)* | 2022-07-28 | Fu Y et al., High-quality reference genome sequences of two Cannaceae species provide insights into the evolution of Cannaceae. Front Plant Sci. 2022 Jul 28, 13: 955904. eCollection 2022. |
| 1047 | *Ipomoea cairica* *(Convolvulaceae)* | 2022-07-27 | Jiang F et al., A chromosome-level reference genome of a Convolvulaceae species Ipomoea cairica. G3 (Bethesda). 2022 Jul 27: jkac187. Online ahead of print. |
| 1046 | *Chenopodium giganteum* *(Amaranthaceae)* | 2022-07-26 | Jarvis DE et al., Chromosome-scale genome assembly of the hexaploid Taiwanese goosefoot djulis (Chenopodium formosanum). Genome Biol Evol. 2022 Jul 26: evac120. Online ahead of print. |
| 1045 | *Toona ciliata* *(Meliaceae)* | 2022-07-26 | Wang X et al., A Chromosome-Level Genome Assembly of Toona ciliata (Meliaceae). Genome Biol Evol. 2022 Jul 26: evac121. Online ahead of print. |
| 1044 | *Aeluropus littoralis* *(Poaceae)* | 2022-07-25 | Hashemi-Petroudi SH et al., Initial Description of the Genome of Aeluropus littoralis, a Halophile Grass. Front Plant Sci. 2022 Jul 11, 13: 906462. eCollection 2022. |
| 1043 | *Eustoma grandiflorum* *(Gentianaceae)* | 2022-07-25 | Liang Y et al., The genome of Eustoma grandiflorum reveals the whole-genome triplication event contributing to ornamental traits in cultivated lisianthus. Plant Biotechnol J. 2022 Jul 25. Online ahead of print. |
| 1042 | *Avena nuda* *(Poaceae)* | 2022-07-18 | Peng Y et al., Reference genome assemblies reveal the origin and evolution of allohexaploid oat. Nat Genet. 2022 Aug, 54(8): 1248-1258. Epub 2022 Jul 18. |
| 1041 | *Lindenbergia luchunensis* *(Orobanchaceae)* | 2022-07-18 | Xu Y et al., Comparative genomics of orobanchaceous species with different parasitic lifestyles reveals the origin and stepwise evolution of plant parasitism. Mol Plant. 2022 Jul 18, S1674-2052(22)00227-1. Online ahead of print. |
| 1040 | *Orobanche aegyptiaca* *(Orobanchaceae)* | 2022-07-18 | Xu Y et al., Comparative genomics of orobanchaceous species with different parasitic lifestyles reveals the origin and stepwise evolution of plant parasitism. Mol Plant. 2022 Jul 18, S1674-2052(22)00227-1. Online ahead of print. |
| 1039 | *Orobanche cumana* *(Orobanchaceae)* | 2022-07-18 | Xu Y et al., Comparative genomics of orobanchaceous species with different parasitic lifestyles reveals the origin and stepwise evolution of plant parasitism. Mol Plant. 2022 Jul 18, S1674-2052(22)00227-1. Online ahead of print. |
| 1038 | *Mesembryanthemum crystallinum* *(Aizoaceae)* | 2022-07-15 | Shen S et al., High-quality ice plant reference genome analysis provides insights into genome evolution and allows exploration of genes involved in the transition from C3 to CAM pathways. Plant Biotechnol J. 2022 Jul 15. Online ahead of print. |
| 1037 | *Rhododendron molle* *(Ericaceae)* | 2022-07-15 | Zhou GL et al., Chromosome-scale genome assembly of Rhododendron molle provides insights into its evolution and terpenoid biosynthesis. BMC Plant Biol. 2022 Jul 15, 22(1): 342. |
| 1036 | *Thymus quinquecostatus* *(Lamiaceae)* | 2022-07-15 | Sun M et al., Chromosome-level assembly and analysis of the Thymus genome provide insights on glandular secretory trichome formation and monoterpenoid biosynthesis in thyme. Plant Commun. 2022 Jul 15: 100413. Online ahead of print. |
| 1035 | *Acorus calamus* *(Acoraceae)* | 2022-07-14 | Shi T et al., The slow-evolving Acorus tatarinowii genome sheds light on ancestral monocot evolution. Nat Plants. 2022 Jul 14. Online ahead of print. |
| 1034 | *Lindera glauca* *(Lauraceae)* | 2022-07-14 | Xiong B et al., Genome of Lindera glauca provides insights into the evolution of biosynthesis genes for aromatic compounds. iScience. 2022 Jul 14, 25(8): 104761. eCollection 2022 Aug 19. |
| 1033 | *Eucalyptus albens* *(Myrtaceae)* | 2022-07-10 | Ferguson S et al., Interspecies genome divergence is predominantly due to frequent small scale rearrangements in Eucalyptus. Mol Ecol. 2023 Mar, 32(6): 1271-1287. Epub 2022 Jul 21. |
| 1032 | *Eucalyptus melliodora* *(Myrtaceae)* | 2022-07-10 | Ferguson S et al., Interspecies genome divergence is predominantly due to frequent small scale rearrangements in Eucalyptus. Mol Ecol. 2023 Mar, 32(6): 1271-1287. Epub 2022 Jul 21. |
| 1031 | *Eucalyptus sideroxylon* *(Myrtaceae)* | 2022-07-10 | Ferguson S et al., Interspecies genome divergence is predominantly due to frequent small scale rearrangements in Eucalyptus. Mol Ecol. 2023 Mar, 32(6): 1271-1287. Epub 2022 Jul 21. |
| 1030 | *Syzygium aromaticum* *(Myrtaceae)* | 2022-07-09 | Ouadi S et al., The clove (Syzygium aromaticum) genome provides insights into the eugenol biosynthesis pathway. Commun Biol. 2022 Jul 9, 5(1): 684. |
| 1029 | *Acer rubrum* *(Sapindaceae)* | 2022-07-08 | Lu X et al., The chromosome-scale genome provides insights into pigmentation in Acer rubrum. Plant Physiol Biochem. 2022 Jul 8, 186: 322-333. Online ahead of print. |
| 1028 | *Bletilla striata* *(Orchidaceae)* | 2022-07-04 | Jiang L et al., Haplotype-resolved genome assembly of Bletilla striata (Thunb.) Reichb.f. to elucidate medicinal values. Plant J. 2022 Jul 4. Online ahead of print. |
| 1027 | *Picea engelmannii* *(Pinaceae)* | 2022-07-04 | Gagalova KK et al., Spruce giga-genomes: structurally similar yet distinctive with differentially expanding gene families and rapidly evolving genes. Plant J. 2022 Jul 4. Online ahead of print. |
| 1026 | *Picea sitchensis* *(Pinaceae)* | 2022-07-04 | Gagalova KK et al., Spruce giga-genomes: structurally similar yet distinctive with differentially expanding gene families and rapidly evolving genes. Plant J. 2022 Jul 4. Online ahead of print. |
| 1025 | *Solanum verrucosum* *(Solanaceae)* | 2022-07-01 | Hosaka AJ et al., A de novo genome assembly of Solanum verrucosum Schlechtendal, a Mexican diploid species geographically isolated from other diploid A-genome species of potato relatives. G3 (Bethesda). 2022 Jul 1: jkac166. Online ahead of print. |
| 1024 | *Boesenbergia rotunda* *(Zingiberaceae)* | 2022-06-30 | Taheri S et al., Genome Assembly and Analysis of the Flavonoid and Phenylpropanoid Biosynthetic Pathways in Fingerroot Ginger (Boesenbergia rotunda). Int J Mol Sci. 2022, 23, 7269 |
| 1023 | *Populus qiongdaoensis* *(Salicaceae)* | 2022-06-30 | Li Y et al., A chromosome-level Populus qiongdaoensis genome assembly provides insights into tropical adaptation and a cryptic turnover of sex determination. Mol Ecol. 2023 Mar, 32(6): 1366-1380. Epub 2022 Jun 30. |
| 1022 | *Ulva compressa* *(Ulvaceae)* | 2022-06-30 | Osorio H et al., The Genome of the Marine Alga Ulva compressa (Chlorophyta) Reveals Protein-Coding Genes with Similarity to Plants and Green Microalgae, but also to Animal, Bacterial, and Fungal Genes. Int. J. Mol. Sci. 2022, 23(13), 7279 |
| 1021 | *Vaccinium bracteatum* *(Ericaceae)* | 2022-06-29 | Yang L et al., Chromosome-level genome assembly and annotation of the native Chinese wild blueberry Vaccinium bracteatum. Fruit Res. 2022 2: 8 |
| 1020 | *Scalesia atractyloides* *(Asteraceae)* | 2022-06-28 | Cerca J et al., The genomic basis of the plant island syndrome in Darwin's giant daisies. Nat Commun. 2022 Jun 28, 13(1): 3729. |
| 1019 | *Cichorium intybus* *(Asteraceae)* | 2022-06-25 | Fan W et al., The genomes of chicory, endive, great burdock and yacon provide insights into Asteraceae paleo-polyploidization history and plant inulin production. Mol Ecol Resour. 2022 Jun 25. Online ahead of print. |
| 1018 | *Nymphoides indica* *(Menyanthaceae)* | 2022-06-25 | Yang JS et al., Chromosome-level genome assembly of the aquatic plant Nymphoides indica reveals transposable element bursts and NBS-LRR gene family expansion shedding light on its invasiveness. DNA Res. 2022 Jun 25: dsac022. Online ahead of print. |
| 1017 | *Oryza australiensis* *(Poaceae)* | 2022-06-25 | Phillips AL et al., The first long-read nuclear genome assembly of Oryza australiensis, a wild rice from northern Australia. Sci Rep. 12, 10823 (2022). |
| 1016 | *Smallanthus sonchifolius* *(Asteraceae)* | 2022-06-25 | Fan W et al., The genomes of chicory, endive, great burdock and yacon provide insights into Asteraceae paleo-polyploidization history and plant inulin production. Mol Ecol Resour. 2022 Jun 25. Online ahead of print. |
| 1015 | *Crataegus pinnatifida* *(Rosaceae)* | 2022-06-24 | Zhang T et al., Cultivated hawthorn (Crataegus pinnatifida var. major) genome sheds light on the evolution of Maleae (apple tribe). J Integr Plant Biol. 2022 Jun 24. Online ahead of print. |
| 1014 | *Tanacetum coccineum* *(Asteraceae)* | 2022-06-24 | Yamashiro T et al., Draft Genome of Tanacetum Coccineum: Genomic Comparison of Closely Related Tanacetum-Family Plants. Int J Mol Sci. 2022, 23, 7039. |
| 1013 | *Abrus melanospermus* *(Fabaceae)* | 2022-06-22 | Xu S et al., Complete genome sequence and phylogenetic analysis of medicinal plant Abrus cantoniensis for evolutionary research and germplasm utilization. Plant Genome. 2022 Jun 24: e20236. Online ahead of print. |

| | plant species | published | |
|---|---|---|---|
| 1012 | *Solanum galapagense* *(Solanaceae)* | 2022-06-20 | Yu X et al., Chromosome-scale genome assemblies of wild tomato relatives Solanum habrochaites and S. galapagense reveal structural variants associated with stress tolerance and terpene biosynthesis. Hortic Res. 2022 Jun 20, uhac139 eCollection 2022. |
| 1011 | *Solanum habrochaites* *(Solanaceae)* | 2022-06-20 | Yu X et al., Chromosome-scale genome assemblies of wild tomato relatives Solanum habrochaites and S. galapagense reveal structural variants associated with stress tolerance and terpene biosynthesis. Hortic Res. 2022 Jun 20, uhac139 eCollection 2022. |
| 1010 | *Elaeagnus angustifolia* *(Elaeagnaceae)* | 2022-06-18 | Mao Y et al., De novo assembly provides new insights into the evolution of Elaeagnus angustifolia L.. Plant Methods 18, 84 (2022). |
| 1009 | *Pohlia nutans* *(Mniaceae)* | 2022-06-17 | Liu S et al., The Antarctic Moss Pohlia nutans Genome Provides Insights Into the Evolution of Bryophytes and the Adaptation to Extreme Terrestrial Habitats. Front Plant Sci., 17 June 2022 |
| 1008 | *Curcuma alismatifolia* *(Zingiberaceae)* | 2022-06-15 | Dong Q et al., The Chromosome-Scale Assembly of the Curcuma alismatifolia Genome Provides Insight Into Anthocyanin and Terpenoid Biosynthesis. Front Plant Sci. 2022 Jun 15, 13: 899588. eCollection 2022. |
| 1007 | *Artocarpus nanchuanensis* *(Moraceae)* | 2022-06-14 | He J et al., A chromosome-level genome assembly of Artocarpus nanchuanensis (Moraceae), an extremely endangered fruit tree. Gigascience. 2022 Jun 14, 11: giac042. |
| 1006 | *Fagopyrum cymosum* *(Polygonaceae)* | 2022-06-14 | He M et al., Comparison of buckwheat genomes reveals the genetic basis of metabolomic divergence and ecotype differentiation. New Phytol. 2022 Jun 14. Online ahead of print. |
| 1005 | *Striga hermonthica* *(Orobanchaceae)* | 2022-06-14 | Qiu S et al., Genome-enabled discovery of candidate virulence loci in Striga hermonthica, a devastating parasite of African cereal crops. New Phytol. 2022 Jun 14. Online ahead of print. |
| 1004 | *Prunus kansuensis* *(Rosaceae)* | 2022-06-13 | Cao K et al., Chromosome-level genome assemblies of four wild peach species provide insights into genome evolution and genetic basis of stress resistance. BMC Biol. 2022 Jun 13, 20(1): 139. |
| 1003 | *Artemisia argyi* *(Asteraceae)* | 2022-06-11 | Miao Y et al., Genome sequencing reveals chromosome fusion and extensive expansion of genes related to secondary metabolism in Artemisia argyi. Plant Biotechnol J. 2022 Jun 11. Online ahead of print. |
| 1002 | *Boswellia sacra* *(Burseraceae)* | 2022-06-10 | Khan AL et al., Genome structure and evolutionary history of frankincense producing Boswellia sacra. iScience. 2022 Jun 10, 25(7): 104574 |
| 1001 | *Cichorium endivia* *(Asteraceae)* | 2022-06-09 | Zhang B et al., The chromosome-scale assembly of endive (Cichorium endivia) genome provides insights into the sesquiterpenoid biosynthesis. Genomics. 2022 Jun 9:110400. Online ahead of print. |
| 1000 | *Solanum andreanum* *(Solanaceae)* | 2022-06-08 | Tang D et al., Genome evolution and diversity of wild and cultivated potatoes. Nature. 2022 Jun 8. Online ahead of print. |
| 999 | *Solanum boliviense* *(Solanaceae)* | 2022-06-08 | Tang D et al., Genome evolution and diversity of wild and cultivated potatoes. Nature. 2022 Jun 8. Online ahead of print. |
| 998 | *Solanum brevicaule* *(Solanaceae)* | 2022-06-08 | Tang D et al., Genome evolution and diversity of wild and cultivated potatoes. Nature. 2022 Jun 8. Online ahead of print. |
| 997 | *Solanum buesii* *(Solanaceae)* | 2022-06-08 | Tang D et al., Genome evolution and diversity of wild and cultivated potatoes. Nature. 2022 Jun 8. Online ahead of print. |
| 996 | *Solanum bulbocastanum* *(Solanaceae)* | 2022-06-08 | Tang D et al., Genome evolution and diversity of wild and cultivated potatoes. Nature. 2022 Jun 8. Online ahead of print. |
| 995 | *Solanum burkartii* *(Solanaceae)* | 2022-06-08 | Tang D et al., Genome evolution and diversity of wild and cultivated potatoes. Nature. 2022 Jun 8. Online ahead of print. |
| 994 | *Solanum cajamarquense* *(Solanaceae)* | 2022-06-08 | Tang D et al., Genome evolution and diversity of wild and cultivated potatoes. Nature. 2022 Jun 8. Online ahead of print. |
| 993 | *Solanum candolleanum* *(Solanaceae)* | 2022-06-08 | Tang D et al., Genome evolution and diversity of wild and cultivated potatoes. Nature. 2022 Jun 8. Online ahead of print. |
| 992 | *Solanum chomatophilum* *(Solanaceae)* | 2022-06-08 | Tang D et al., Genome evolution and diversity of wild and cultivated potatoes. Nature. 2022 Jun 8. Online ahead of print. |
| 991 | *Solanum etuberosum* *(Solanaceae)* | 2022-06-08 | Tang D et al., Genome evolution and diversity of wild and cultivated potatoes. Nature. 2022 Jun 8. Online ahead of print. |
| 990 | *Solanum jamesii* *(Solanaceae)* | 2022-06-08 | Tang D et al., Genome evolution and diversity of wild and cultivated potatoes. Nature. 2022 Jun 8. Online ahead of print. |
| 989 | *Solanum lignicaule* *(Solanaceae)* | 2022-06-08 | Tang D et al., Genome evolution and diversity of wild and cultivated potatoes. Nature. 2022 Jun 8. Online ahead of print. |
| 988 | *Solanum morelliforme* *(Solanaceae)* | 2022-06-08 | Tang D et al., Genome evolution and diversity of wild and cultivated potatoes. Nature. 2022 Jun 8. Online ahead of print. |
| 987 | *Solanum multiinterruptum* *(Solanaceae)* | 2022-06-08 | Tang D et al., Genome evolution and diversity of wild and cultivated potatoes. Nature. 2022 Jun 8. Online ahead of print. |
| 986 | *Solanum neorossii* *(Solanaceae)* | 2022-06-08 | Tang D et al., Genome evolution and diversity of wild and cultivated potatoes. Nature. 2022 Jun 8. Online ahead of print. |
| 985 | *Solanum palustre* *(Solanaceae)* | 2022-06-08 | Tang D et al., Genome evolution and diversity of wild and cultivated potatoes. Nature. 2022 Jun 8. Online ahead of print. |
| 984 | *Solanum paucissectum* *(Solanaceae)* | 2022-06-08 | Tang D et al., Genome evolution and diversity of wild and cultivated potatoes. Nature. 2022 Jun 8. Online ahead of print. |
| 983 | *Solanum pinnatisectum* *(Solanaceae)* | 2022-06-08 | Tang D et al., Genome evolution and diversity of wild and cultivated potatoes. Nature. 2022 Jun 8. Online ahead of print. |
| 982 | *Solanum piurae* *(Solanaceae)* | 2022-06-08 | Tang D et al., Genome evolution and diversity of wild and cultivated potatoes. Nature. 2022 Jun 8. Online ahead of print. |
| 981 | *Solanum sogarandinum* *(Solanaceae)* | 2022-06-08 | Tang D et al., Genome evolution and diversity of wild and cultivated potatoes. Nature. 2022 Jun 8. Online ahead of print. |
| 980 | *Solanum vernei* *(Solanaceae)* | 2022-06-08 | Tang D et al., Genome evolution and diversity of wild and cultivated potatoes. Nature. 2022 Jun 8. Online ahead of print. |
| 979 | *Syringa oblata* *(Oleaceae)* | 2022-06-08 | Wang Y et al., A chromosome-level genome of Syringa oblata provides new insights into chromosome formation in Oleaceae and evolutionary history of lilacs. Plant J. 2022 Jun 8. Online ahead of print. |
| 978 | *Papaver armeniacum* *(Papaveraceae)* | 2022-06-07 | Catania T et al., A functionally conserved STORR gene fusion in Papaver species that diverged 16.8 million years ago. Nat Commun. 2022 Jun 7, 13(1): 3150. |
| 977 | *Papaver atlanticum* *(Papaveraceae)* | 2022-06-07 | Catania T et al., A functionally conserved STORR gene fusion in Papaver species that diverged 16.8 million years ago. Nat Commun. 2022 Jun 7, 13(1): 3150. |

| | plant species | published | |
|---|---|---|---|
| 976 | *Papaver bracteatum* *(Papaveraceae)* | 2022-06-07 | Catania T et al., A functionally conserved STORR gene fusion in Papaver species that diverged 16.8 million years ago. Nat Commun. 2022 Jun 7, 13(1): 3150. |
| 975 | *Papaver californicum* *(Papaveraceae)* | 2022-06-07 | Catania T et al., A functionally conserved STORR gene fusion in Papaver species that diverged 16.8 million years ago. Nat Commun. 2022 Jun 7, 13(1): 3150. |
| 974 | *Papaver nudicaule* *(Papaveraceae)* | 2022-06-07 | Catania T et al., A functionally conserved STORR gene fusion in Papaver species that diverged 16.8 million years ago. Nat Commun. 2022 Jun 7, 13(1): 3150. |
| 973 | *Fragaria chiloensis* *(Rosaceae)* | 2022-06-06 | Cauret CMS et al., Chromosome-scale assembly with a phased sex-determining region resolves features of early Z and W chromosome differentiation in a wild octoploid strawberry. G3 (Bethesda). 2022 Jun 6: jkac139. Online ahead of print. |
| 972 | *Iochroma cyaneum* *(Solanaceae)* | 2022-06-06 | Powell AF et al., Genome sequence for the blue-flowered Andean shrub Iochroma cyaneum reveals extensive discordance across the berry clade of Solanaceae. Plant Genome. 2022 Jun 6: e20223. Online ahead of print. |
| 971 | *Entada phaseoloides* *(Fabaceae)* | 2022-06-04 | Lin M et al., Chromosome-level genome of Entada phaseoloides provides insights into genome evolution and triterpenoid saponins biosynthesis. Mol Ecol Resour. 2022 Jun 4. Online ahead of print. |
| 970 | *Lanxangia tsao-ko* *(Zingiberaceae)* | 2022-06-01 | Sun F et al., Genome Sequencing of Amomum tsao-ko Provides Novel Insight Into Its Volatile Component Biosynthesis. Front Plant Sci. 01 June 2022 |
| 969 | *Litsea coreana* *(Lauraceae)* | 2022-06-01 | Zhang B et al., High-quality chromosome-level genome assembly of Litsea coreana L. provides insights into Magnoliids evolution and flavonoid biosynthesis. Genomics. 2022 Jun 1: 110394. Online ahead of print. |
| 968 | *Toxicodendron vernicifluum* *(Anacardiaceae)* | 2022-06-01 | Bai G et al., The chromosome-level genome for Toxicodendron vernicifluum provides crucial insights into Anacardiaceae evolution and urushiol biosynthesis. iScience. 2022 Jun 2, 25(7): 104512. eCollection 2022 Jul 15. |
| 967 | *Pistia stratiotes* *(Araceae)* | 2022-05-27 | Qian Z et al., The chromosome-level genome of a free-floating aquatic weed Pistia stratiotes provides insights into its rapid invasion. Mol Ecol Resour. 2022 May 27. Online ahead of print. |
| 966 | *Rubus corchorifolius* *(Rosaceae)* | 2022-05-25 | Yang Y et al., Genome assembly and population resequencing reveal the geographical divergence of shanmei (Rubus corchorifolius). Genomics Proteomics Bioinformatics. 2022 May 25: S1672-0229(22)00067-5. Online ahead of print. |
| 965 | *Lepidium sativum* *(Brassicaceae)* | 2022-05-20 | Patat AS et al., Construction and characterization of a de novo draft genome of garden cress (Lepidium sativum L.). Funct Integr Genomics. 2022 May 20. Online ahead of print. |
| 964 | *Avena insularis* *(Poaceae)* | 2022-05-18 | Kamal N et al., The mosaic oat genome gives insights into a uniquely healthy cereal crop. Nature. 2022 Jun, 606(7912): 113-119. Epub 2022 May 18. |
| 963 | *Avena longiglumis* *(Poaceae)* | 2022-05-18 | Kamal N et al., The mosaic oat genome gives insights into a uniquely healthy cereal crop. Nature. 2022 Jun, 606(7912): 113-119. Epub 2022 May 18. |
| 962 | *Avena sativa* *(Poaceae)* | 2022-05-18 | Kamal N et al., The mosaic oat genome gives insights into a uniquely healthy cereal crop. Nature. 2022 Jun, 606(7912): 113-119. Epub 2022 May 18. |
| 961 | *Pedinomonas minor* *(Pedinomonadaceae)* | 2022-05-13 | Xu Y et al., Chromosome-level genome of Pedinomonas minor (Chlorophyta) unveils adaptations to abiotic stress in a rapidly fluctuating environment. New Phytol. 2022 May 13. Online ahead of print. |
| 960 | *Suaeda glauca* *(Amaranthaceae)* | 2022-05-12 | Yi L et al., Chromosome-Scale, Haplotype-Resolved Genome Assembly of Suaeda Glauca. Front. Genet. 12 May 2022 |
| 959 | *Alsophila spinulosa* *(Cyatheaceae)* | 2022-05-09 | Huang X et al., The flying spider-monkey tree fern genome provides insights into fern evolution and arborescence. Nat Plants. 2022 May 9. Online ahead of print. |
| 958 | *Bruguiera cylindrica* *(Rhizophoraceae)* | 2022-05-05 | Shearman JR et al., Assembly of a hybrid mangrove, Bruguiera hainesii, and its two ancestral contributors, Bruguiera cylindrica and Bruguiera gymnorhiza. Genomics. 2022 May 5: 110382. Online ahead of print. |
| 957 | *Bruguiera x hainesii* *(Rhizophoraceae)* | 2022-05-05 | Shearman JR et al., Assembly of a hybrid mangrove, Bruguiera hainesii, and its two ancestral contributors, Bruguiera cylindrica and Bruguiera gymnorhiza. Genomics. 2022 May 5: 110382. Online ahead of print. |
| 956 | *Hordeum marinum* *(Poaceae)* | 2022-05-05 | Kuang L et al., The genome and gene editing system of sea barleygrass provide a novel platform for cereal domestication and stress tolerance studies. Plant Commun. 2022 May 05: 100333 eCollection 2022 May 05. |
| 955 | *Leptochloa chinensis* *(Poaceae)* | 2022-05-05 | Wang L et al., Genomic insights into the origin, adaptive evolution a Genomic insights into the origin, adaptive evolution and herbicide resistance of Leptochloa chinensis, a devastating tetraploid weedy grass in rice fields.nd herbicide resistance of Leptochloa chinensis, a devastating tetraploid weedy grass in rice fields. Mol Plant. 2022 May 5: S1674-2052(22)00149-6. Online ahead of print. |
| 954 | *Limonium bicolor* *(Plumbaginaceae)* | 2022-05-05 | Yuan F et al., The genome of the recretohalophyte Limonium bicolor provides insights into salt gland development and salinity adaptation during terrestrial evolution. Mol Plant. 2022 May 5: S1674-2052(22)00148-4. Online ahead of print. |
| 953 | *Quercus acutissima* *(Fagaceae)* | 2022-05-05 | Fu R et al., Genome-wide analyses of introgression between two sympatric Asian oak species. Nat Ecol Evol. 2022 May 5. Online ahead of print. |
| 952 | *Ensete glaucum* *(Musaceae)* | 2022-04-30 | Wang Z et al., A chromosome-level reference genome of Ensete glaucum gives insight into diversity and chromosomal and repetitive sequence evolution in the Musaceae. Gigascience. 2022 Apr 30, 11: giac027. |
| 951 | *Annona glabra* *(Annonaceae)* | 2022-04-28 | He Z et al., Evolution of coastal forests based on a full set of mangrove genomes. Nat Ecol Evol. 2022 Apr 28. Online ahead of print. |
| 950 | *Avicennia alba* *(Acanthaceae)* | 2022-04-28 | He Z et al., Evolution of coastal forests based on a full set of mangrove genomes. Nat Ecol Evol. 2022 Apr 28. Online ahead of print. |
| 949 | *Avicennia germinans* *(Acanthaceae)* | 2022-04-28 | He Z et al., Evolution of coastal forests based on a full set of mangrove genomes. Nat Ecol Evol. 2022 Apr 28. Online ahead of print. |
| 948 | *Avicennia officinalis* *(Acanthaceae)* | 2022-04-28 | He Z et al., Evolution of coastal forests based on a full set of mangrove genomes. Nat Ecol Evol. 2022 Apr 28. Online ahead of print. |
| 947 | *Avicennia rumphiana* *(Acanthaceae)* | 2022-04-28 | He Z et al., Evolution of coastal forests based on a full set of mangrove genomes. Nat Ecol Evol. 2022 Apr 28. Online ahead of print. |
| 946 | *Barringtonia asiatica* *(Lecythidaceae)* | 2022-04-28 | He Z et al., Evolution of coastal forests based on a full set of mangrove genomes. Nat Ecol Evol. 2022 Apr 28. Online ahead of print. |
| 945 | *Barringtonia racemosa* *(Lecythidaceae)* | 2022-04-28 | He Z et al., Evolution of coastal forests based on a full set of mangrove genomes. Nat Ecol Evol. 2022 Apr 28. Online ahead of print. |
| 944 | *Carallia pectinifolia* *(Rhizophoraceae)* | 2022-04-28 | He Z et al., Evolution of coastal forests based on a full set of mangrove genomes. Nat Ecol Evol. 2022 Apr 28. Online ahead of print. |
| 943 | *Ceriops tagal* *(Rhizophoraceae)* | 2022-04-28 | He Z et al., Evolution of coastal forests based on a full set of mangrove genomes. Nat Ecol Evol. 2022 Apr 28. Online ahead of print. |
| 942 | *Clerodendrum inerme* *(Lamiaceae)* | 2022-04-28 | He Z et al., Evolution of coastal forests based on a full set of mangrove genomes. Nat Ecol Evol. 2022 Apr 28. Online ahead of print. |
| 941 | *Combretum micranthum* *(Combretaceae)* | 2022-04-28 | He Z et al., Evolution of coastal forests based on a full set of mangrove genomes. Nat Ecol Evol. 2022 Apr 28. Online ahead of print. |

| | plant species | published | |
|---|---|---|---|
| 940 | *Conocarpus erectus* *(Combretaceae)* | 2022-04-28 | He Z et al., Evolution of coastal forests based on a full set of mangrove genomes. Nat Ecol Evol. 2022 Apr 28. Online ahead of print. |
| 939 | *Duabanga grandiflora* *(Lythraceae)* | 2022-04-28 | He Z et al., Evolution of coastal forests based on a full set of mangrove genomes. Nat Ecol Evol. 2022 Apr 28. Online ahead of print. |
| 938 | *Heritiera littoralis* *(Malvaceae)* | 2022-04-28 | He Z et al., Evolution of coastal forests based on a full set of mangrove genomes. Nat Ecol Evol. 2022 Apr 28. Online ahead of print. |
| 937 | *Kandelia candel* *(Rhizophoraceae)* | 2022-04-28 | He Z et al., Evolution of coastal forests based on a full set of mangrove genomes. Nat Ecol Evol. 2022 Apr 28. Online ahead of print. |
| 936 | *Lagerstroemia speciosa* *(Lythraceae)* | 2022-04-28 | He Z et al., Evolution of coastal forests based on a full set of mangrove genomes. Nat Ecol Evol. 2022 Apr 28. Online ahead of print. |
| 935 | *Laguncularia racemosa* *(Combretaceae)* | 2022-04-28 | He Z et al., Evolution of coastal forests based on a full set of mangrove genomes. Nat Ecol Evol. 2022 Apr 28. Online ahead of print. |
| 934 | *Lumnitzera littorea* *(Combretaceae)* | 2022-04-28 | He Z et al., Evolution of coastal forests based on a full set of mangrove genomes. Nat Ecol Evol. 2022 Apr 28. Online ahead of print. |
| 933 | *Lumnitzera racemosa* *(Combretaceae)* | 2022-04-28 | He Z et al., Evolution of coastal forests based on a full set of mangrove genomes. Nat Ecol Evol. 2022 Apr 28. Online ahead of print. |
| 932 | *Morinda citrifolia* *(Rubiaceae)* | 2022-04-28 | He Z et al., Evolution of coastal forests based on a full set of mangrove genomes. Nat Ecol Evol. 2022 Apr 28. Online ahead of print. |
| 931 | *Nypa fruticans* *(Arecaceae)* | 2022-04-28 | He Z et al., Evolution of coastal forests based on a full set of mangrove genomes. Nat Ecol Evol. 2022 Apr 28. Online ahead of print. |
| 930 | *Pemphis acidula* *(Lythraceae)* | 2022-04-28 | He Z et al., Evolution of coastal forests based on a full set of mangrove genomes. Nat Ecol Evol. 2022 Apr 28. Online ahead of print. |
| 929 | *Pluchea indica* *(Asteraceae)* | 2022-04-28 | He Z et al., Evolution of coastal forests based on a full set of mangrove genomes. Nat Ecol Evol. 2022 Apr 28. Online ahead of print. |
| 928 | *Pongamia pinnata* *(Fabaceae)* | 2022-04-28 | He Z et al., Evolution of coastal forests based on a full set of mangrove genomes. Nat Ecol Evol. 2022 Apr 28. Online ahead of print. |
| 927 | *Premna obtusifolia* *(Lamiaceae)* | 2022-04-28 | He Z et al., Evolution of coastal forests based on a full set of mangrove genomes. Nat Ecol Evol. 2022 Apr 28. Online ahead of print. |
| 926 | *Rhizophora mangle* *(Rhizophoraceae)* | 2022-04-28 | He Z et al., Evolution of coastal forests based on a full set of mangrove genomes. Nat Ecol Evol. 2022 Apr 28. Online ahead of print. |
| 925 | *Rhizophora mucronata* *(Rhizophoraceae)* | 2022-04-28 | He Z et al., Evolution of coastal forests based on a full set of mangrove genomes. Nat Ecol Evol. 2022 Apr 28. Online ahead of print. |
| 924 | *Rhizophora stylosa* *(Rhizophoraceae)* | 2022-04-28 | He Z et al., Evolution of coastal forests based on a full set of mangrove genomes. Nat Ecol Evol. 2022 Apr 28. Online ahead of print. |
| 923 | *Scaevola hainanensis* *(Goodeniaceae)* | 2022-04-28 | He Z et al., Evolution of coastal forests based on a full set of mangrove genomes. Nat Ecol Evol. 2022 Apr 28. Online ahead of print. |
| 922 | *Scaevola sericea* *(Goodeniaceae)* | 2022-04-28 | He Z et al., Evolution of coastal forests based on a full set of mangrove genomes. Nat Ecol Evol. 2022 Apr 28. Online ahead of print. |
| 921 | *Scyphiphora hydrophyllacea* *(Rubiaceae)* | 2022-04-28 | He Z et al., Evolution of coastal forests based on a full set of mangrove genomes. Nat Ecol Evol. 2022 Apr 28. Online ahead of print. |
| 920 | *Sonneratia apetala* *(Lythraceae)* | 2022-04-28 | He Z et al., Evolution of coastal forests based on a full set of mangrove genomes. Nat Ecol Evol. 2022 Apr 28. Online ahead of print. |
| 919 | *Sonneratia ovata* *(Lythraceae)* | 2022-04-28 | He Z et al., Evolution of coastal forests based on a full set of mangrove genomes. Nat Ecol Evol. 2022 Apr 28. Online ahead of print. |
| 918 | *Swietenia macrophylla* *(Meliaceae)* | 2022-04-28 | He Z et al., Evolution of coastal forests based on a full set of mangrove genomes. Nat Ecol Evol. 2022 Apr 28. Online ahead of print. |
| 917 | *Taxillus chinensis* *(Loranthaceae)* | 2022-04-28 | Fu J et al., Chromosome-Level Genome Assembly of the Hemiparasitic Taxillus chinensis. Genome Biol Evol. 2022 Apr 28: evac060. Online ahead of print. |
| 916 | *Xylocarpus granatum* *(Meliaceae)* | 2022-04-28 | He Z et al., Evolution of coastal forests based on a full set of mangrove genomes. Nat Ecol Evol. 2022 Apr 28. Online ahead of print. |
| 915 | *Xylocarpus moluccensis* *(Meliaceae)* | 2022-04-28 | He Z et al., Evolution of coastal forests based on a full set of mangrove genomes. Nat Ecol Evol. 2022 Apr 28. Online ahead of print. |
| 914 | *Xylocarpus rumphii* *(Meliaceae)* | 2022-04-28 | He Z et al., Evolution of coastal forests based on a full set of mangrove genomes. Nat Ecol Evol. 2022 Apr 28. Online ahead of print. |
| 913 | *Corydalis tomentella* *(Papaveraceae)* | 2022-04-27 | Xu Z et al., The genome of Corydalis reveals the evolution of benzylisoquinoline alkaloid biosynthesis in Ranunculales. Plant J. 2022 Apr 27. Online ahead of print. |
| 912 | *Cynodon dactylon* *(Poaceae)* | 2022-04-25 | Zhang B et al., A High-Quality Haplotype-Resolved Genome of Common Bermudagrass (Cynodon dactylon L.) Provides Insights Into Polyploid Genome Stability and Prostrate Growth. Front Plant Sci. 2022 Apr 25, 13: 890980. eCollection 2022. |
| 911 | *Styphnolobium japonicum* *(Fabaceae)* | 2022-04-25 | Lei W et al., Chromosome-level genome assembly and characterization of Sophora japonica. DNA Res. 2022 Apr 25: dsac009. Online ahead of print. |
| 910 | *Platanthera guangdongensis* *(Orchidaceae)* | 2022-04-21 | Li MH et al., Genomes of leafy and leafless Platanthera orchids illuminate the evolution of mycoheterotrophy. Nat Plants (2022). |
| 909 | *Platanthera zijinensis* *(Orchidaceae)* | 2022-04-21 | Li MH et al., Genomes of leafy and leafless Platanthera orchids illuminate the evolution of mycoheterotrophy. Nat Plants (2022). |
| 908 | *Larix kaempferi* *(Pinaceae)* | 2022-04-20 | Sun C et al., The Larix kaempferi genome reveals new insights into wood properties. J Integr Plant Biol. 2022 Apr 20. Online ahead of print. |
| 907 | *Bauhinia variegata* *(Fabaceae)* | 2022-04-19 | Zhong Y et al., Chromosomal-level genome assembly of the orchid tree Bauhinia variegata (Leguminosae, Cercidoideae) supports the allotetraploid origin hypothesis of Bauhinia. DNA Res. 2022 Apr 19: dsac012. Online ahead of print. |
| 906 | *Cycas panzhihuaensis* *(Cycadaceae)* | 2022-04-18 | Liu Y et al., The Cycas genome and the early evolution of seed plants. Nat Plants. (2022). |
| 905 | *Dioscorea alata* *(Dioscoreaceae)* | 2022-04-14 | Bredeson JV et al., Chromosome evolution and the genetic basis of agronomically important traits in greater yam. Nat Commun. 2022 Apr 14, 13(1): 2001. |
| 904 | *Salvia hispanica* *(Lamiaceae)* | 2022-04-14 | Wang L et al., A chromosome-level genome assembly of chia provides insights into high omega-3 content and coat color variation of its seeds. Plant Commun. 2022 Apr 14: 100326. |

| | plant species | published | |
|---|---|---|---|
| 903 | *Carpinus viminea* *(Betulaceae)* | 2022-04-13 | Wang Z et al., Genomic evidence for homoploid hybrid speciation between ancestors of two different genera. Nat Commun. 13, 1987 (2022). |
| 902 | *Solanum lycopersicoides* *(Solanaceae)* | 2022-04-12 | Powell AF et al., A Solanum lycopersicoides reference genome facilitates insights into tomato specialized metabolism and immunity. Plant J. 2022 Apr 12. Online ahead of print. |
| 901 | *Paspalum notatum* *(Poaceae)* | 2022-04-11 | Yan Z et al., High-quality chromosome-scale de novo assembly of the Paspalum notatum Fluegge genome. BMC Genomics. 2022 Apr 11, 23(1): 293. |
| 900 | *Ilex asprella* *(Aquifoliaceae)* | 2022-04-09 | Kong BL et al., Chromosomal level genome of Ilex asprella and insight into antiviral triterpenoid pathway. Genomics. 2022 Apr 9: 110366. Online ahead of print. |
| 899 | *Gastrodia menghaiensis* *(Orchidaceae)* | 2022-04-07 | Jiang Y et al., The Gastrodia menghaiensis (Orchidaceae) genome provides new insights of orchid mycorrhizal interactions. BMC Plant Biol. 2022 Apr 7, 22(1): 179. |
| 898 | *Panax japonicus* *(Araliaceae)* | 2022-04-07 | Wang ZH et al., Reshuffling of the ancestral core-eudicot genome shaped chromatin topology and epigenetic modification in Panax. Nat Commun. 13, 1902 (2022) |
| 897 | *Panax quinquefolius* *(Araliaceae)* | 2022-04-07 | Wang ZH et al., Reshuffling of the ancestral core-eudicot genome shaped chromatin topology and epigenetic modification in Panax. Nat Commun. 13, 1902 (2022) |
| 896 | *Panax stipuleanatus* *(Araliaceae)* | 2022-04-07 | Wang ZH et al., Reshuffling of the ancestral core-eudicot genome shaped chromatin topology and epigenetic modification in Panax. Nat Commun. 13, 1902 (2022) |
| 895 | *Gentiana dahurica* *(Gentianaceae)* | 2022-04-05 | Li T et al., The chromosome-level genome assembly of Gentiana dahurica (Gentianaceae) provides insights into gentiopicroside biosynthesis. DNA Res. 2022 Apr 5: dsac008. Online ahead of print. |
| 894 | *Quercus mongolica* *(Fagaceae)* | 2022-04-04 | Ai W et al., A chromosome-scale genome assembly of the Mongolian oak (Quercus mongolica). Mol Ecol Resour. 2022 Apr 4. Online ahead of print. |
| 893 | *Streptocarpus rexii* *(Gesneriaceae)* | 2022-04-03 | Nishii K et al., The first genome for the Cape Primrose Streptocarpus rexii (Gesneriaceae), a model plant for studying meristem-driven shoot diversity. Plant Direct. 2022 Apr 03, eCollection 2022 Apr. |
| 892 | *Nelumbo lutea* *(Nelumbonaceae)* | 2022-04-01 | Zheng P et al., Comparative analyses of American and Asian lotus genomes reveal insights into petal color, carpel thermogenesis and domestication. Plant J. 2022 Apr 1. Online ahead of print. |
| 891 | *Bupleurum chinense* *(Apiaceae)* | 2022-03-31 | Zhang Q et al., Chromosome-Level Genome Assembly of Bupleurum chinense DC Provides Insights Into the Saikosaponin Biosynthesis. Front Genet. 31 March 2022 |
| 890 | *Malus prunifolia* *(Rosaceae)* | 2022-03-29 | Li Z et al., A Chromosome-Scale Reference Genome Provides Insights into the Genetic Origin and Grafting-Mediated Stress Tolerance of Malus prunifolia. Plant Biotechnol J. 2022 Mar 29. Online ahead of print. |
| 889 | *Carex kokanica* *(Cyperaceae)* | 2022-03-23 | Qu G et al., Draft genomes assembly and annotation of Carex parvula and Carex kokanica reveals stress-specific genes. Sci Rep. 2022 Mar 23, 12(1) |
| 888 | *Carex parvula* *(Cyperaceae)* | 2022-03-23 | Qu G et al., Draft genomes assembly and annotation of Carex parvula and Carex kokanica reveals stress-specific genes. Sci Rep. 2022 Mar 23, 12(1) |
| 887 | *Hibiscus hamabo* *(Malvaceae)* | 2022-03-23 | Wang Z et al., The genome of Hibiscus hamabo reveals its adaptation to saline and waterlogged habitat. Hortic Res. 2022 Mar 23: uhac067. Online ahead of print. |
| 886 | *Morus indica* *(Moraceae)* | 2022-03-21 | Jain M et al., Draft genome sequence of Indian mulberry (Morus indica) provides a resource for functional and translational genomics. Genomics. 2022 Mar 21: 110346. Online ahead of print. |
| 885 | *Camellia fluviatilis* *(Theaceae)* | 2022-03-20 | Gong W et al., Chromosome-level genome of Camellia lanceoleosa provides a valuable resource for understanding genome evolution and self-incompatibility. Plant J. 2022 Mar 20. Online ahead of print. |
| 884 | *Rubus idaeus* *(Rosaceae)* | 2022-03-16 | Davik J et al., A chromosome-level genome sequence assembly of the red raspberry (Rubus idaeus L.). PLoS One. 2022 Mar 16, 17(3): e0265096. eCollection 2022. |
| 883 | *Solanum muricatum* *(Solanaceae)* | 2022-03-16 | Song X et al., Chromosome-level pepino genome provides insights into genome evolution and anthocyanin biosynthesis in Solanaceae. Plant J. 2022 Mar 16. Online ahead of print. |
| 882 | *Portulaca amilis* *(Portulacaceae)* | 2022-03-14 | Gilman IS et al., Gene co-expression reveals the modularity and integration of C4 and CAM in Portulaca. Plant Physiol. 2022 Mar 14: kiac116. Online ahead of print. |
| 881 | *Haloxylon ammodendron* *(Amaranthaceae)* | 2022-03-10 | Wang M et al., Chromosome-level genome assembly of a xerophytic plant, Haloxylon ammodendron. DNA Res. 2022 Mar 10: dsac006. Online ahead of print. |
| 880 | *Nannochloris desiccata* *(Chlorellaceae)* | 2022-03-09 | Sanders CK et al., Phylogenetic analyses and reclassification of the oleaginous marine species Nannochloris sp. desiccata (Trebouxiophyceae, Chlorophyta), formerly Chlorella desiccata, supported by a high-quality genome assembly. J Phycol. 2022 Mar 9. Online ahead of print. |
| 879 | *Vaccinium oxycoccos* *(Ericaceae)* | 2022-03-07 | Kawash J et al., Contrasting a reference cranberry genome to a crop wild relative provides insights into adaptation, domestication, and breeding. PLoS One. 2022 Mar 7, 17(3): e0264966. eCollection 2022. |
| 878 | *Acer pseudosieboldianum* *(Sapindaceae)* | 2022-03-03 | Li X et al., Chromosome-Level genome assembly for Acer pseudosieboldianum and highlights to mechanisms for leaf color and shape change. Front. Plant Sci., 03 March 2022 \| https://doi.org/10.3389/fpls.2022.850054 |
| 877 | *Aralia elata* *(Araliaceae)* | 2022-03-01 | Liu W et al., A High-Quality Reference Genome Sequence and Genetic Transformation System of Aralia elata. Front Plant Sci. 01 March 2022 |
| 876 | *Dendrobium nobile* *(Orchidaceae)* | 2022-03-01 | Xu Q et al., Chromosome-Scale Assembly of the Dendrobium nobile Genome Provides Insights Into the Molecular Mechanism of the Biosynthesis of the Medicinal Active Ingredient of Dendrobium. Front Genet. 2022 Mar 01: 844622 eCollection 2022. |
| 875 | *Macadamia tetraphylla* *(Proteaceae)* | 2022-02-23 | Niu Y et al., The Chromosome-Scale Reference Genome of Macadamia tetraphylla Provides Insights Into Fatty Acid Biosynthesis. Front Genet. 23 February 2022 |
| 874 | *Sapindus mukorossi* *(Sapindaceae)* | 2022-02-17 | Xue T et al., Chromosome-scale assembly and population diversity analyses provide insights into the evolution of Sapindus mukorossi. Hortic Res. 2022, 9: uhac012 |
| 873 | *Vatica mangachapoi* *(Dipterocarpaceae)* | 2022-02-16 | Tang L et al., A chromosome-scale genome and transcriptomic analysis of the endangered tropical tree Vatica mangachapoi (Dipterocarpaceae). DNA Res. 2022 Feb 16: dsac005. Online ahead of print. |
| 872 | *Amorphophallus konjac* *(Araceae)* | 2022-02-15 | Gao Y et al., A chromosome-level genome assembly of Amorphophallus konjac provides insights into konjac glucomannan biosynthesis. Comput Struct Biotechnol J. 2022 Feb 15, 20: 1002-1011. eCollection 2022. |
| 871 | *Entodon seductrix* *(Entodontaceae)* | 2022-02-15 | Yu J et al., Chromosome-level genome assemblies of two Hypnales (mosses) reveal high intergeneric synteny. Genome Biol Evol. 2022 Feb 15: evac020. Online ahead of print. |
| 870 | *Hypnum curvifolium* *(Pylaisiaceae)* | 2022-02-15 | Yu J et al., Chromosome-level genome assemblies of two Hypnales (mosses) reveal high intergeneric synteny. Genome Biol Evol. 2022 Feb 15: evac020. Online ahead of print. |
| 869 | *Hibiscus mutabilis* *(Malvaceae)* | 2022-02-14 | Yang Y et al., A High-Quality, Chromosome-Level Genome Provides Insights Into Determinate Flowering Time and Color of Cotton Rose (Hibiscus mutabilis). Front Plant Sci. 14 February 2022 |
| 868 | *Heliosperma pusillum* *(Caryophyllaceae)* | 2022-02-13 | Szukala A et al., Polygenic routes lead to parallel altitudinal adaptation in Heliosperma pusillum (Caryophyllaceae). Mol Ecol. 2023 Apr, 32(8): 1832-1847. Epub 2022 Feb 13. |

| | plant species | published | |
|---|---|---|---|
| 867 | *Aristolochia contorta* (*Aristolochiaceae*) | 2022-02-11 | Cui X et al., Chromosome-level genome assembly of Aristolochia contorta provides insights into the biosynthesis of benzylisoquinoline alkaloids and aristolochic acids. Hortic Res. 2022 Feb 11: uhac005. Online ahead of print. |
| 866 | *Bruguiera sexangula* (*Rhizophoraceae*) | 2022-02-11 | Pootakham W et al., De novo reference assembly of the upriver orange mangrove (Bruguiera sexangula) genome. Genome Biol Evol. 2022 Feb 11: evac025. Online ahead of print. |
| 865 | *Hippophae rhamnoides* (*Elaeagnaceae*) | 2022-02-04 | Wu Z et al., The genome of Hippophae rhamnoides provides insights into a conserved molecular mechanism in actinorhizal and rhizobial symbiosis. New Phytol. 2022 Feb 4. Online ahead of print. |
| 864 | *Echinochloa colona* (*Poaceae*) | 2022-02-03 | Wu D et al., Genomic insights into the evolution of Echinochloa species as weed and orphan crop. Nat Commun. 13, 689 (2022). |
| 863 | *Buxus sinica* (*Buxaceae*) | 2022-02-02 | Chanderbali AS et al., Buxus and Tetracentron genomes help resolve eudicot genome history. Nat Commun. 13, 643, 2022, Epub ahead of print. |
| 862 | *Ceriops zippeliana* (*Rhizophoraceae*) | 2022-02-02 | Pootakham W et al., A de novo reference assembly of the yellow mangrove Ceriops zippeliana genome. G3 (Bethesda). 2022 Feb 2: jkac025. Online ahead of print. |
| 861 | *Chondria armata* (*Rhodomelaceae*) | 2022-02-02 | Steele TS et al., Domoic acid biosynthesis in the red alga Chondria armata suggests a complex evolutionary history for toxin production. Proc Natl Acad Sci U S A. 2022 Feb 8, 119(6): e2117407119. |
| 860 | *Gilia yorkii* (*Polemoniaceae*) | 2022-02-02 | Jarvis DE et al., Chromosome-scale genome assembly of Gilia yorkii enables genetic mapping of floral traits in an inter-species cross. Genome Biol Evol. 2022 Feb 2: evac017. Online ahead of print. |
| 859 | *Sporobolus pyramidalis* (*Poaceae*) | 2022-02-01 | Chavez Montes RA et al., A comparative genomics examination of desiccation tolerance and sensitivity in two sister grass species. Proc Natl Acad Sci U S A. 2022 Feb 1, 119(5): e2118886119. |
| 858 | *Sporobolus stapfianus* (*Poaceae*) | 2022-02-01 | Chavez Montes RA et al., A comparative genomics examination of desiccation tolerance and sensitivity in two sister grass species. Proc Natl Acad Sci U S A. 2022 Feb 1, 119(5): e2118886119. |
| 857 | *Crucihimalaya lasiocarpa* (*Brassicaceae*) | 2022-01-30 | Feng L et al., A chromosome-level genome assembly of an alpine plant Crucihimalaya lasiocarpa provides insights into high-altitude adaptation. DNA Res. 2022 Jan 30: dsac004. Online ahead of print. |
| 856 | *Bretschneidera sinensis* (*Akaniaceae*) | 2022-01-28 | Liu HL et al., The genome of the Paleogene relic tree Bretschneidera sinensis: insights into trade-offs in gene family evolution, demographic history, and adaptive SNPs. DNA Res. 2022 Jan 28, 29(1): dsac003. |
| 855 | *Gracilaria domingensis* (*Gracilariaceae*) | 2022-01-28 | Nakamura-Gouvea N et al., Insights into agar and secondary metabolite pathways from the genome of the red alga Gracilaria domingensis (Rhodophyta, Gracilariales). J Phycol. 2022 Jan 28. Online ahead of print. |
| 854 | *Speranskia yunnanensis* (*Euphorbiaceae*) | 2022-01-24 | Yuan G et al., Chromosome-Level Genome Assembly of the Rare and Endangered Tropical Plant Speranskia yunnanensis (Euphorbiaceae). Front Genet. 2022 Jan 24 |
| 853 | *Bruguiera parviflora* (*Rhizophoraceae*) | 2022-01-21 | Pootakham W et al., A chromosome-scale reference genome assembly of yellow mangrove (Bruguiera parviflora) reveals a whole genome duplication event associated with the Rhizophoraceae lineage. Mol Ecol Resour. 2022 Jan 21. Online ahead of print. |
| 852 | *Pueraria montana* (*Fabaceae*) | 2022-01-19 | Shang X et al., Chromosomal-level genome and multi-omics dataset of Pueraria lobata var. thomsonii provide new insights into legume family and the isoflavone and puerarin biosynthesis pathways. Hortic Res. 2022 Jan 19: uhab035. Online ahead of print. |
| 851 | *Asparagus kiusianus* (*Asparagaceae*) | 2022-01-18 | Shirasawa K et al., Chromosome-scale haplotype-phased genome assemblies of the male and female lines of wild asparagus (Asparagus kiusianus), a dioecious plant species. DNA Res. 2022 Jan 18: dsac002. Online ahead of print. |
| 850 | *Camellia chekiangoleosa* (*Theaceae*) | 2022-01-18 | Shen TF et al., The reference genome of Camellia chekiangoleosa provides insights into Camellia evolution and tea oil biosynthesis. Hortic Res. 2022 Jan 18, uhab083. Online ahead of print. |
| 849 | *Chrysanthemum lavandulifolium* (*Asteraceae*) | 2022-01-18 | Wen X et al., The Chrysanthemum lavandulifolium genome and the molecular mechanism underlying diverse capitulum types. Hortic Res. 2022 Jan 18: uhab022. Online ahead of print. |
| 848 | *Telopea speciosissima* (*Proteaceae*) | 2022-01-11 | Chen SH et al., Chromosome-level de novo genome assembly of Telopea speciosissima (New South Wales waratah) using long-reads, linked-reads and Hi-C. Mol Ecol Resour. 2022 Jan 11. Online ahead of print. |
| 847 | *Camellia oleifera* (*Theaceae*) | 2022-01-10 | Lin P et al., The genome of oil-Camellia and population genomics analysis provide insights into seed oil domestication. Genome Biol. 2022 Jan 10, 23(1): 14. |
| 846 | *Begonia darthvaderiana* (*Begoniaceae*) | 2022-01-08 | Li L et al., Genomes shed light on the evolution of Begonia, a mega-diverse genus. New Phytol. 2022 Jan 8. Online ahead of print. |
| 845 | *Begonia loranthoides* (*Begoniaceae*) | 2022-01-08 | Li L et al., Genomes shed light on the evolution of Begonia, a mega-diverse genus. New Phytol. 2022 Jan 8. Online ahead of print. |
| 844 | *Begonia masoniana* (*Begoniaceae*) | 2022-01-08 | Li L et al., Genomes shed light on the evolution of Begonia, a mega-diverse genus. New Phytol. 2022 Jan 8. Online ahead of print. |
| 843 | *Begonia peltatifolia* (*Begoniaceae*) | 2022-01-08 | Li L et al., Genomes shed light on the evolution of Begonia, a mega-diverse genus. New Phytol. 2022 Jan 8. Online ahead of print. |
| 842 | *Dendrocalamus latiflorus* (*Poaceae*) | 2022-01-06 | Zheng Y et al., Allele-aware chromosome-scale assembly of the allopolyploid genome of hexaploid Ma Bamboo (Dendrocalamus latiflorus Munro). J Integr Plant Biol. 2022 Jan 6. Online ahead of print. |
| 841 | *Ilex polyneura* (*Aquifoliaceae*) | 2022-01-05 | Yao X et al., A chromosome-scale genome assembly for the holly (Ilex polyneura) provides insights into genomic adaptations to elevation in Southwest China. Hortic Res. 2021 Dec 18, 9. |
| 840 | *Sorbus aucuparia* (*Rosaceae*) | 2022-01-05 | Zhao D et al., Genome sequence and transcriptome of Sorbus pohuashanensis provides insights into population evolution and leaf sunburn response. J Genet Genomics. 2022 Jan 05: Online ahead of print. |
| 839 | *Bidens hawaiensis* (*Asteraceae*) | 2022-01-04 | Bellinger MR et al., A genome for Bidens hawaiensis: a member of a hexaploid Hawaiian plant adaptive radiation. J Hered. 2022 Jan 4: esab077. Online ahead of print. |
| 838 | *Sindora glabra* (*Fabaceae*) | 2022-01-04 | Yu N et al., The Diesel Tree Sindora glabra Genome Provides Insights Into the Evolution of Oleoresin Biosynthesis. Front Plant Sci. 2022 Jan 4, 12: 794830. eCollection 2021. |
| 837 | *Litchi chinensis* (*Sapindaceae*) | 2022-01-03 | Hu G et al., Two divergent haplotypes from a highly heterozygous lychee genome suggest independent domestication events for early and late-maturing cultivars. Nat Genet. (2022) Epub ahead of print |
| 836 | *Synsepalum dulcificum* (*Sapotaceae*) | 2022-01-03 | Yang Z et al., The Chromosome-Level Genome of Miracle Fruit (Synsepalum dulcificum) Provides New Insights Into the Evolution and Function of Miraculin. Front Plant Sci. 2022 Jan 3, 12: 804662. eCollection 2021. |
| 835 | *Aegilops bicornis* (*Poaceae*) | 2022-01-01 | Li LF et al., Genome sequences of the five Sitopsis species of Aegilops and the origin of polyploid wheat B-subgenome. Mol Plant. 2022 Jan 01: Online ahead of print. |
| 834 | *Aegilops longissima* (*Poaceae*) | 2022-01-01 | Li LF et al., Genome sequences of the five Sitopsis species of Aegilops and the origin of polyploid wheat B-subgenome. Mol Plant. 2022 Jan 01: Online ahead of print. |
| 833 | *Aegilops searsii* (*Poaceae*) | 2022-01-01 | Li LF et al., Genome sequences of the five Sitopsis species of Aegilops and the origin of polyploid wheat B-subgenome. Mol Plant. 2022 Jan 01: Online ahead of print. |
| 832 | *Aegilops sharonensis* (*Poaceae*) | 2022-01-01 | Li LF et al., Genome sequences of the five Sitopsis species of Aegilops and the origin of polyploid wheat B-subgenome. Mol Plant. 2022 Jan 01: Online ahead of print. |

| | plant species | published | |
|---|---|---|---|
| 831 | *Aegilops speltoides* *(Poaceae)* | 2022-01-01 | Li LF et al., Genome sequences of the five Sitopsis species of Aegilops and the origin of polyploid wheat B-subgenome. Mol Plant. 2022 Jan 01: Online ahead of print. |
| 830 | *Acer negundo* *(Sapindaceae)* | 2021-12-29 | McEvoy SL et al., Strategies of tolerance reflected in two North American maple genomes. Plant J. 2021 Dec 29. Online ahead of print. |
| 829 | *Acer saccharum* *(Sapindaceae)* | 2021-12-29 | McEvoy SL et al., Strategies of tolerance reflected in two North American maple genomes. Plant J. 2021 Dec 29. Online ahead of print. |
| 828 | *Pinus tabuliformis* *(Pinaceae)* | 2021-12-22 | Niu S et al., The Chinese pine genome and methylome unveil key features of conifer evolution. Cell. 2021 Dec 22: S0092-8674(21)01428-8. Online ahead of print. |
| 827 | *P.incertae sedis sp.YPF701* *(P.incertae sedis)* | 2021-12-19 | Repetti SI et al., Nuclear genome of a pedinophyte pinpoints genomic innovation and streamlining in the green algae. New Phytol. 2022 Mar, 233(5): 2144-2154. |
| 826 | *Taraxacum mongolicum* *(Asteraceae)* | 2021-12-17 | Lin T et al., Extensive sequence divergence between the reference genomes of Taraxacum kok-saghyz and Taraxacum mongolicum. Sci China Life Sci. 2021 Dec 17. Online ahead of print. |
| 825 | *Argentina anserina* *(Rosaceae)* | 2021-12-15 | Gan X et al., Chromosome-Level Genome Assembly Provides New Insights into Genome Evolution and Tuberous Root Formation of Potentilla anserina. Genes. 2021, 12: 1993. |
| 824 | *Sequoia sempervirens* *(Cupressaceae)* | 2021-12-14 | Neale DB et al., Assembled and annotated 26.5 Gbp coast redwood genome: a resource for estimating evolutionary adaptive potential and investigating hexaploid origin. G3 (Bethesda). 2021 Dec 14: jkab380. Online ahead of print. |
| 823 | *Phragmites australis* *(Poaceae)* | 2021-12-11 | Oh DH et al., Novel genome characteristics contribute to the invasiveness of Phragmites australis (common reed). Mol Ecol. 2022 Feb, 31(4): 1142-1159. Epub 2021 Dec 11. |
| 822 | *Acer catalpifolium* *(Sapindaceae)* | 2021-12-08 | Yu T et al., Whole-genome sequencing of Acer catalpifolium reveals evolutionary history of endangered species. Genome Biol Evol. 2021 Dec 8: evab271. Online ahead of print. |
| 821 | *Lemna minuta* *(Araceae)* | 2021-12-06 | Abramson BW et al., The genome and preliminary single-nuclei transcriptome of Lemna minuta reveals mechanisms of invasiveness. Plant Physiol. 2021 Dec 6: kiab564 Online ahead of print. |
| 820 | *Trapa natans* *(Lythraceae)* | 2021-12-03 | Lu RS et al., Genome sequencing and transcriptome analyses provide insights into the origin and domestication of water caltrop (Trapa spp., Lythraceae). Plant Biotechnol J. 2021 Dec 3. Online ahead of print. |
| 819 | *Elaeagnus mollis* *(Elaeagnaceae)* | 2021-12-02 | Ren B et al., Genome sequence of Elaeagnus mollis, the first chromosome-level genome of the family Elaeagnaceae. Genome Biol Evol. 2021 Dec 2: evab266. Online ahead of print. |
| 818 | *Chrysanthemum makinoi* *(Asteraceae)* | 2021-12-01 | van Lieshout N et al., De novo whole-genome assembly of Chrysanthemum makinoi, a key wild chrysanthemum. G3 (Bethesda). 2021 Oct 13: jkab358. Online ahead of print. |
| 817 | *Cymbidium ensifolium* *(Orchidaceae)* | 2021-12-01 | Ai Y et al., The Cymbidium genome reveals the evolution of unique morphological traits. Hortic Res. 8, 255 (2021). |
| 816 | *Oenanthe javanica* *(Apiaceae)* | 2021-12-01 | Liu JX et al., Integrative genome, transcriptome, microRNA, and degradome analysis of water dropwort (Oenanthe javanica) in response to water stress. Hortic Res. 8, 262 (2021). |
| 815 | *Scutellaria indica* *(Lamiaceae)* | 2021-12-01 | Choi H et al., De Novo Assembly and Species-Specific Marker Development as a Useful Tool for the Identification of Scutellaria L. Species. Curr Issues Mol Biol. 2021 Dec 01, 43(3), 2177-2188 |
| 814 | *Chloranthus sessilifolius* *(Chloranthaceae)* | 2021-11-26 | Ma J et al., The Chloranthus sessilifolius genome provides insight into early diversification of angiosperms. Nat Commun. 2021 Nov 26, 12(1): 6929. |
| 813 | *Chloranthus spicatus* *(Chloranthaceae)* | 2021-11-26 | Guo X et al., Chloranthus genome provides insights into the early diversification of angiosperms. Nat Commun. 2021 Nov 26, 12(1): 6930. |
| 812 | *Arctostaphylos glauca* *(Ericaceae)* | 2021-11-24 | Huang Y et al., Reference genome assembly of the big berry Manzanita (Arctostaphylos glauca). J Hered. 2021 Nov 24, esab071. Online ahead of print. |
| 811 | *Astrephomene gubernaculifera* *(Goniaceae)* | 2021-11-22 | Yamashita S et al., Genome sequencing of the multicellular alga Astrephomene provides insights into convergent evolution of germ-soma differentiation. Sci Rep. 2021 Nov 22, 11(1): 22231. |
| 810 | *Neolamarckia cadamba* *(Rubiaceae)* | 2021-11-22 | Zhao X et al., Chromosome-level assembly of Neolamarckia cadamba genome provides insights into the evolution of cadambine biosynthesis. Plant J. 2021 Nov 22. Online ahead of print. |
| 809 | *Typha latifolia* *(Typhaceae)* | 2021-11-22 | Widanagama SD et al., Genome assembly, annotation, and comparative analysis of the cattail Typha latifolia. G3 (Bethesda). 2021 Nov 22: jkab401. Online ahead of print |
| 808 | *Physalis pubescens* *(Solanaceae)* | 2021-11-18 | Lu J et al., The Physalis floridana genome provides insights into the biochemical and morphological evolution of Physalis fruits. Hortic Res. 2021 Nov 18, 8(1): 244. |
| 807 | *Actinidia kolomikta* *(Actinidiaceae)* | 2021-11-17 | Wu H et al., A chromosome-level genome assembly for the wild kiwifruit Actinidia kolomikta provides insights into canker resistance and fruit development. Plant Biotechnol J. 2021 Nov 17. Online ahead of print. |
| 806 | *Cinnamomum camphora* *(Lauraceae)* | 2021-11-16 | Shen T et al., The chromosome-level genome sequence of the camphor tree provides insights into Lauraceae evolution and terpene biosynthesis. Plant Biotechnol J. 2021 Nov 16. Online ahead of print. |
| 805 | *Macadamia jansenii* *(Proteaceae)* | 2021-11-16 | Sharma P et al., Denovo chromosome level assembly of a plant genome from long read sequence data. Plant J. 2021 Nov 16. Online ahead of print. |
| 804 | *Arctium lappa* *(Asteraceae)* | 2021-11-10 | Yang Y et al., The first high-quality chromosomal genome assembly of a medicinal and edible plant Arctium lappa. Mol Ecol Resour. 2021 Nov 10. doi: 10.1111/1755-0998.13547. Online ahead of print. |
| 803 | *Prunus fruticosa* *(Rosaceae)* | 2021-11-10 | Wohner TW et al., The draft chromosome-level genome assembly of tetraploid ground cherry (Prunus fruticosa Pall.) from long reads. Genomics. 2021 Nov 10: S0888-7543 (21)00384-0. Online ahead of print. |
| 802 | *Astragalus sinicus* *(Fabaceae)* | 2021-11-08 | Chang D et al., The chromosome-level genome assembly of Astragalus sinicus and comparative genomic analyses provide new resources and insights for understanding legume-rhizobial interactions. Plant Commun. (2021) |
| 801 | *Fraxinus pennsylvanica* *(Oleaceae)* | 2021-11-08 | Huff M et al., A high quality reference genome for Fraxinus pennsylvanica for ash species restoration and research. Mol Ecol Resour. 2021 Nov 8. Online ahead of print. |
| 800 | *Isoetes taiwanensis* *(Isoetaceae)* | 2021-11-03 | Wickell D et al., Underwater CAM photosynthesis elucidated by Isoetes genome. Nat Commun. 2021 Nov 3, 12(1): 6348. |
| 799 | *Neoporphyra haitanensis* *(Bangiaceae)* | 2021-11-03 | Chen H et al., Insights into the Ancient Adaptation to Intertidal Environments by Red Algae Based on a Genomic and Multiomics Investigation of Neoporphyra haitanensis. Mol Biol Evol. 2022 Jan 7, 39(1): msab315. |
| 798 | *Vaccinium darrowii* *(Ericaceae)* | 2021-11-01 | Yu J et al., High-quality reference genome and annotation aids understanding of berry development for evergreen blueberry (Vaccinium darrowii) . Hortic Res. 2021 Nov 1, 8(1): 228. |
| 797 | *Melilotus albus* *(Fabaceae)* | 2021-10-30 | Wu F et al., Genome and systems biology of Melilotus albus provides insights into coumarins biosynthesis. Plant Biotechnol J. 2021 Oct 30. Online ahead of print. |
| 796 | *Melastoma dodecandrum* *(Melastomataceae)* | 2021-10-28 | Hao Y et al., The Melastoma dodecandrum genome and the evolution of Myrtales. J Genet Genomics. 2021 Oct 28: S1673-8527(21)00338-6. Online ahead of print. |

| | plant species | published | |
|---|---|---|---|
| 795 | *Microstegium vimineum* (*Poaceae*) | 2021-10-28 | Ramachandran D et al., Chromosome Level Genome Assembly and Annotation of Highly Invasive Japanese Stiltgrass (Microstegium vimineum). Genome Biol Evol. 2021 Nov 5, 13(11): evab238. |
| 794 | *Cyclocarya paliurus* (*Juglandaceae*) | 2021-10-25 | Zheng X et al., Insights into the evolution and hypoglycemic metabolite biosynthesis of autotetraploid Cyclocarya paliurus by combining genomic, transcriptomic and metabolomic analyses. Ind Crops Prod. 2021 Dec, 173: 114154 |
| 793 | *Fragaria daltoniana* (*Rosaceae*) | 2021-10-25 | Qiao Q et al., Evolutionary history and pan-genome dynamics of strawberry ( Fragaria spp.). Proc Natl Acad Sci USA. 2021 Nov 9, 118(45): e2105431118 |
| 792 | *Fragaria mandschurica* (*Rosaceae*) | 2021-10-25 | Qiao Q et al., Evolutionary history and pan-genome dynamics of strawberry ( Fragaria spp.). Proc Natl Acad Sci USA. 2021 Nov 9, 118(45): e2105431118 |
| 791 | *Fragaria pentaphylla* (*Rosaceae*) | 2021-10-25 | Qiao Q et al., Evolutionary history and pan-genome dynamics of strawberry ( Fragaria spp.). Proc Natl Acad Sci USA. 2021 Nov 9, 118(45): e2105431118 |
| 790 | *Castanopsis tibetana* (*Fagaceae*) | 2021-10-24 | Sun Y et al., The chromosome-scale genome assembly of Castanopsis tibetana provides a powerful comparative framework to study the evolution and adaptation of Fagaceae trees. Mol Ecol Resour. 2021 Oct 24. Online ahead of print. |
| 789 | *Cymbidium goeringii* (*Orchidaceae*) | 2021-10-23 | Chung O et al., A chromosome-scale genome assembly and annotation of the spring orchid (Cymbidium goeringii). Mol Ecol Resour. 2021 Oct 23. Online ahead of print. |
| 788 | *Dipterocarpus turbinatus* (*Dipterocarpaceae*) | 2021-10-22 | Wang S et al., The chromosome-scale genomes of Dipterocarpus turbinatus and Hopea hainanensis (Dipterocarpaceae) provide insights into fragrant oleoresin biosynthesis and hard wood formation. Plant Biotechnol J. 2021 Oct 22. Online ahead of print. |
| 787 | *Hopea hainanensis* (*Dipterocarpaceae*) | 2021-10-22 | Wang S et al., The chromosome-scale genomes of Dipterocarpus turbinatus and Hopea hainanensis (Dipterocarpaceae) provide insights into fragrant oleoresin biosynthesis and hard wood formation. Plant Biotechnol J. 2021 Oct 22. Online ahead of print. |
| 786 | *Artemisia tridentata* (*Asteraceae*) | 2021-10-19 | Melton AE et al., A draft genome provides hypotheses on drought tolerance in a keystone plant species in Western North America threatened by climate change. Ecol Evol. 2021 Oct 19, 11(21): 15417-15429. eCollection 2021 Nov. |
| 785 | *Euphorbia lathyris* (*Euphorbiaceae*) | 2021-10-19 | Wang M et al., High-quality genome assembly of an important biodiesel plant, Euphorbia lathyris L. DNA Res. 2021 Oct 19, dsab022. Online ahead of print. |
| 784 | *Pugionium cornutum* (*Brassicaceae*) | 2021-10-19 | Hu Q et al., Genome evolution of the psammophyte Pugionium for desert adaptation and further speciation. Proc Natl Acad Sci USA. 2021 Oct 19, 118(42): e2025711118. |
| 783 | *Pugionium dolabratum* (*Brassicaceae*) | 2021-10-19 | Hu Q et al., Genome evolution of the psammophyte Pugionium for desert adaptation and further speciation. Proc Natl Acad Sci USA. 2021 Oct 19, 118(42): e2025711118. |
| 782 | *Curcuma longa* (*Zingiberaceae*) | 2021-10-15 | Chakraborty A et al., Genome sequencing of turmeric provides evolutionary insights into its medicinal properties. Commun Biol. 2021 Oct 15, 4(1): 1193. |
| 781 | *Papaver rhoeas* (*Papaveraceae*) | 2021-10-15 | Yang X et al., Three chromosome-scale Papaver genomes reveal punctuated patchwork evolution of the morphinan and noscapine biosynthesis pathway. Nat Commun. 2021 Oct 15, 12(1): 6030. |
| 780 | *Papaver setigerum* (*Papaveraceae*) | 2021-10-15 | Yang X et al., Three chromosome-scale Papaver genomes reveal punctuated patchwork evolution of the morphinan and noscapine biosynthesis pathway. Nat Commun. 2021 Oct 15, 12(1): 6030. |
| 779 | *Rhododendron henanense* (*Ericaceae*) | 2021-10-15 | Zhou XJ et al., The chromosome-scale genome assembly, annotation and evolution of Rhododendron henanense subsp. lingbaoense Mol Ecol Resour. 2021 Oct 15. Online ahead of print. |
| 778 | *Panax vietnamensis* (*Araliaceae*) | 2021-10-11 | Tien NQD et al., De novo whole-genome assembly and discovery of genes involved in triterpenoid saponin biosynthesis of Vietnamese ginseng (Panax vietnamensis Ha et Grushv.). Physiol Mol Biol Plants. 2021 Oct, 27(10): 2215-2229. Epub 2021 Oct 11. |
| 777 | *Prunus x kanzakura* (*Rosaceae*) | 2021-10-11 | Shirasawa K et al., Genome sequencing and analysis of two early-flowering cherry (Cerasus x kanzakura) varieties, 'Kawazu-zakura' and 'Atami-zakura'. DNA Res. 2021 Oct 11, 28(6): dsab026. |
| 776 | *Macrotyloma uniflorum* (*Fabaceae*) | 2021-10-08 | Shirasawa K et al., A chromosome-scale draft genome sequence of horsegram (Macrotyloma uniflorum). Gigabyte. 1, 2021 |
| 775 | *Shorea leprosula* (*Dipterocarpaceae*) | 2021-10-07 | Ng KKS et al., The genome of Shorea leprosula (Dipterocarpaceae) highlights the ecological relevance of drought in aseasonal tropical rainforests. Commun Biol. 2021 Oct 7, 4(1): 1166. |
| 774 | *Vicia sativa* (*Fabaceae*) | 2021-10-07 | Shirasawa K et al., Genome features of common vetch (Vicia sativa) in natural habitats. Plant Direct. 2021 Oct 7, 5(10): e352. eCollection 2021 Oct. |
| 773 | *Streptochaeta angustifolia* (*Poaceae*) | 2021-10-04 | Seetharam AS et al., The Streptochaeta Genome and the Evolution of the Grasses. Front Plant Sci. 04 October 2021 |
| 772 | *Alliaria petiolata* (*Brassicaceae*) | 2021-10-02 | Alabi N et al., Genome Report: A draft genome of Alliaria petiolata (garlic mustard) as a model system for invasion genetics. G3 (Bethesda). 2021 Oct 2:jkab339. Online ahead of print. |
| 771 | *Prunus davidiana* (*Rosaceae*) | 2021-10-01 | Tan Q et al., Chromosome-level genome assemblies of five Prunus species and genome-wide association studies for key agronomic traits in peach. Hortic Res. 8, 213 (2021). |
| 770 | *Clausena lansium* (*Rutaceae*) | 2021-09-30 | Fan Y et al., The Clausena lansium (Wampee) genome reveal new insights into the carbazole alkaloids biosynthesis pathway. Genomics. 2021 Nov, 113(6):3696-3704. Epub 2021 Sep 11. |
| 769 | *Euscaphis japonica* (*Staphyleaceae*) | 2021-09-29 | Sun WH et al., The Euscaphis japonica genome and the evolution of malvids. Plant J. 2021 Sep 29. Online ahead of print. |
| 768 | *Populus tomentosa* (*Salicaceae*) | 2021-09-22 | An X et al., High quality haplotype-resolved genome assemblies of Populus tomentosa Carr., a stabilized interspecific hybrid species that is widespread in Asia. Mol Ecol Resour. 2021 Sep 22. Online ahead of print. |
| 767 | *Perilla citriodora* (*Lamiaceae*) | 2021-09-17 | Zhang Y et al., Incipient diploidization of the medicinal plant Perilla within 10,000 years. Nat Commun. 12, 5508 (2021). |
| 766 | *Perilla frutescens* (*Lamiaceae*) | 2021-09-17 | Zhang Y et al., Incipient diploidization of the medicinal plant Perilla within 10,000 years. Nat Commun. 12, 5508 (2021). |
| 765 | *Bruguiera gymnorhiza* (*Rhizophoraceae*) | 2021-09-16 | Miryeganeh M et al., De novo genome assembly and in natura epigenomics reveal salinity-induced DNA methylation in the mangrove tree Bruguiera gymnorhiza. New Phytol. 2021 Sep 16. Online ahead of print. |
| 764 | *Castanea crenata* (*Fagaceae*) | 2021-09-13 | Shirasawa K et al., Chromosome-level genome assembly of Japanese chestnut (Castanea crenata Sieb. et Zucc.) reveals conserved chromosomal segments in woody rosids. DNA Res. 2021 Sep 13, 28(5): dsab016. |
| 763 | *Nephelium lappaceum* (*Sapindaceae*) | 2021-09-13 | Zhang W et al., Rambutan genome revealed gene networks for spine formation and aril development. Plant J. 2021 Sep 13. Online ahead of print. |
| 762 | *Gynostemma pentaphyllum* (*Cucurbitaceae*) | 2021-09-09 | Huang D et al., Chromosome-level genome assembly of Gynostemma pentaphyllum provides insights into gypenoside biosynthesis. DNA Res. 2021 Sep 9: dsab018. Online ahead of print. |
| 761 | *Vitellaria paradoxa* (*Sapotaceae*) | 2021-09-09 | Hale I et al., Genomic Resources to Guide Improvement of the Shea Tree. Front Plant Sci. 09 September 2021 |
| 760 | *Achnatherum splendens* (*Poaceae*) | 2021-09-06 | Ren G et al., The genome sequence provides insights into salt tolerance of Achnatherum splendens (Gramineae), a constructive species of alkaline grassland. Plant Biotechnol J. 2021 Sep 6. Online ahead of print. |

| | plant species | published | |
|---|---|---|---|
| 759 | *Gossypium anomalum* (*Malvaceae*) | 2021-09-06 | Grover CE et al., The Gossypium anomalum genome as a resource for cotton improvement and evolutionary analysis of hybrid incompatibility. G3 (Bethesda). 2021 Sep 06 |
| 758 | *Zanthoxylum bungeanum* (*Rutaceae*) | 2021-09-03 | Feng S et al., Zanthoxylum-specific whole genome duplication and recent activity of transposable elements in the highly repetitive paleotetraploid Z. bungeanum genome. Hortic Res. 2021 Sep 3, 8(1): 205. |
| 757 | *Aristolochia fimbriata* (*Aristolochiaceae*) | 2021-09-02 | Qin L et al., Insights into angiosperm evolution, floral development and chemical biosynthesis from the Aristolochia fimbriata genome. Nat Plants. 2021 Sep 2. Online ahead of print. |
| 756 | *Dendrobium chrysotoxum* (*Orchidaceae*) | 2021-09-01 | Zhang Y et al., Chromosome-scale assembly of the Dendrobium chrysotoxum genome enhances the understanding of orchid evolution. Hortic Res. 2021 Sep 1, 8(1): 183. |
| 755 | *Eremochloa ophiuroides* (*Poaceae*) | 2021-09-01 | Wang J et al., A high-quality chromosome-scale assembly of the centipedegrass Eremochloa ophiuroides (Munro) Hack. genome provides insights into chromosomal structural evolution and prostrate growth habit. Hortic Res. 2021 Sep 1, 8(1): 201. |
| 754 | *Jasminum sambac* (*Oleaceae*) | 2021-08-30 | Xu S et al., A high-quality genome assembly of Jasminum sambac provides insight into floral trait formation and Oleaceae genome evolution. Mol Ecol Resour. 2021 Aug 30. Online ahead of print. |
| 753 | *Alpinia nigra* (*Zingiberaceae*) | 2021-08-24 | Ranavat S et al., A Draft Genome of the Ginger Species Alpinia nigra and New Insights into the Genetic Basis of Flexistyly. Genes. 2021, 12, 1297. |
| 752 | *Ipomoea aquatica* (*Convolvulaceae*) | 2021-08-24 | Hao Y et al., The chromosome-based genome provides insights into the evolution in water spinach. Sci Hortic. 2021 Nov, 289: 110501 |
| 751 | *Gossypium rotundifolium* (*Malvaceae*) | 2021-08-23 | Wang M et al., Comparative Genome Analyses Highlight Transposon-Mediated Genome Expansion and the Evolutionary Architecture of 3D Genomic Folding in Cotton. Mol Biol Evol. 2021 Aug 23, 38(9): 3621-3636. |
| 750 | *Lunularia cruciata* (*Lunulariaceae*) | 2021-08-23 | Linde AM et al., Rates and patterns of molecular evolution in bryophyte genomes, with focus on complex thalloid liverworts, Marchantiopsida. Mol Phylogenet Evol. 2021 Dec, 165: 107295. Epub 2021 Aug 23. |
| 749 | *Arabidopsis arenosa* (*Brassicaceae*) | 2021-08-19 | Burns R et al., Gradual evolution of allopolyploidy in Arabidopsis suecica. Nat Ecol Evol. (2021) |
| 748 | *Arabidopsis suecica* (*Brassicaceae*) | 2021-08-19 | Jiang X et al., Concerted genomic and epigenomic changes accompany stabilization of Arabidopsis allopolyploids. Nat Ecol Evol. (2021) |
| 747 | *Magnolia officinalis* (*Magnoliaceae*) | 2021-08-17 | Yin Y et al., The chromosome-scale genome of Magnolia officinalis provides insight into the evolutionary position of magnoliids. iScience. 2021 Aug 17 |
| 746 | *Cardamine enshiensis* (*Brassicaceae*) | 2021-08-10 | Huang C et al., The Cardamine enshiensis genome reveals whole genome duplication and insight into selenium hyperaccumulation and tolerance. Cell Discov. 7, 62 (2021) |
| 745 | *Rhododendron ovatum* (*Ericaceae*) | 2021-08-10 | Wang X et al., High-quality evergreen azalea genome reveals tandem duplication-facilitated low-altitude adaptability and floral scent evolution. Plant Biotechnol J. 2021 Aug 10. Epub ahead of print |
| 744 | *Melaleuca alternifolia* (*Myrtaceae*) | 2021-08-09 | Voelker J et al., A high-quality draft genome for Melaleuca alternifolia (tea tree): a new platform for evolutionary genomics of myrtaceous terpene-rich species. Gigabyte. 1, 2021 |
| 743 | *Zingiber officinale* (*Zingiberaceae*) | 2021-08-05 | Li HL et al., Haplotype-resolved genome of diploid ginger (Zingiber officinale) and its unique gingerol biosynthetic pathway. Hortic Res. 8, 189 (2021). |
| 742 | *Cymbidium sinense* (*Orchidaceae*) | 2021-08-03 | Yang FX et al., The genome of Cymbidium sinense revealed the evolution of orchid traits. Plant Biotechnol J. 2021 Aug 3. Epub ahead of print |
| 741 | *Musa textilis* (*Musaceae*) | 2021-08-02 | Galvez LC et al., Sequencing and de Novo Assembly of Abaca (Musa textilis Nee) var. Abuab Genome Genes. 2021, 12, 1202. |
| 740 | *Solanum stenotomum* (*Solanaceae*) | 2021-07-29 | Yan L et al., Genome assembly of primitive cultivated potato Solanum stenotomum provides insights into potato evolution. G3 (Bethesda). 2021 Jul 29. Online ahead of print. |
| 739 | *Stipa capillata* (*Poaceae*) | 2021-07-28 | Baiakhmetov E et al., The first draft genome of feather grasses using SMRT sequencing and its implications in molecular studies of Stipa. Sci Rep. 11, 15345 (2021). |
| 738 | *Allium cepa* (*Amaryllidaceae*) | 2021-07-27 | Finkers R et al., Insights from the first genome assembly of Onion (Allium cepa). G3 (Bethesda). 2021 Jul 27 |
| 737 | *Zizania palustris* (*Poaceae*) | 2021-07-26 | Haas M et al., Whole Genome Assembly and Annotation of Northern Wild Rice, Zizania palustris L., Supports a Whole Genome Duplication in the Zizania Genus. Plant J. 2021 Jul 26. Epub ahead of print |
| 736 | *Passiflora organensis* (*Passifloraceae*) | 2021-07-23 | Costa ZP et al., A genome sequence resource for the genus Passiflora, the genome of the wild diploid species Passiflora organensis. Plant Genome. 2021 Jul 23: e20117. Epub ahead of print |
| 735 | *Buddleja alternifolia* (*Scrophulariaceae*) | 2021-07-22 | Ma YP et al., Genome-wide analysis of butterfly bush (Buddleja alternifolia) in three uplands provides insights into biogeography, demography and speciation. New Phytol. 2021 Jul 22. Epub ahead of print |
| 734 | *Picrorhiza kurrooa* (*Plantaginaceae*) | 2021-07-22 | Sharma T et al., The first draft genome of Picrorhiza kurrooa, an endangered medicinal herb from Himalayas. Sci Rep. 11, 14944 (2021). |
| 733 | *Prunus mira* (*Rosaceae*) | 2021-07-22 | Wang X et al., Genomic basis of high-altitude adaptation in Tibetan Prunus fruit trees. Curr Biol. 2021 Jul 22: S0960-9822(21)00891-5. Epub ahead of print |
| 732 | *Rhododendron kiyosumense* (*Ericaceae*) | 2021-07-20 | Shirasawa K et al., Whole-genome sequencing and analysis of two azaleas, Rhododendron ripense and Rhododendron kiyosumense. DNA Res. 2021 Jul 20: dsab010. Epub ahead of print |
| 731 | *Rhododendron ripense* (*Ericaceae*) | 2021-07-20 | Shirasawa K et al., Whole-genome sequencing and analysis of two azaleas, Rhododendron ripense and Rhododendron kiyosumense. DNA Res. 2021 Jul 20: dsab010. Epub ahead of print |
| 730 | *Zanthoxylum armatum* (*Rutaceae*) | 2021-07-18 | Wang M et al., Chromosome-level genome assembly of Sichuan pepper provides insights into apomixis, drought tolerance, and alkaloid biosynthesis. Mol Ecol Resour. 2021 Oct, 21(7): 2533-2545. Epub 2021 Jul 18. |
| 729 | *Taxus chinensis* (*Taxaceae*) | 2021-07-15 | Xiong X et al., The Taxus genome provides insights into paclitaxel biosynthesis. Nat Plants. (2021). Epub ahead of print |
| 728 | *Vaccinium myrtillus* (*Ericaceae*) | 2021-07-14 | Wu C et al., A chromosome-scale assembly of the bilberry genome identifies a complex locus controlling berry anthocyanin composition. Mol Ecol Resour. 2021 Jul 14. Epub ahead of print |
| 727 | *Apostasia ramifera* (*Orchidaceae*) | 2021-07-13 | Zhang W et al., Genome sequence of Apostasia ramifera provides insights into the adaptive evolution in orchids. BMC Genomics. 2021 Jul 13, 22(1): 536. |
| 726 | *Welwitschia mirabilis* (*Welwitschiaceae*) | 2021-07-12 | Wan T et al., The Welwitschia genome reveals a unique biology underpinning extreme longevity in deserts. Nat Commun. 12, 4247 (2021). |
| 725 | *Leptodermis oblonga* (*Rubiaceae*) | 2021-07-08 | Guo XM et al., Chromosomal-level assembly of the Leptodermis oblonga (Rubiaceae) genome and its phylogenetic implications. Genomics. 2021 Jul 9 113(5): 3072-3082. Epub ahead of print |
| 724 | *Rehmannia glutinosa* (*Orobanchaceae*) | 2021-07-08 | Ma L et al., De novo genome assembly of the potent medicinal plant Rehmannia glutinosa using nanopore technology. Comput Struct Biotechnol J. 2021 Jul 8 |

| | plant species | published | |
|---|---|---|---|
| 723 | *Rhododendron griersonianum* *(Ericaceae)* | 2021-06-30 | Ma H et al., Chromosome-level genome assembly and population genetic analysis of a critically endangered rhododendron provide insights into its conservation. Plant J. 2021 Jun 30. Epub ahead of print |
| 722 | *Paulownia fortunei* *(Paulowniaceae)* | 2021-06-29 | Cao Y et al., Genomic insights into the fast growth of paulownias and the formation of Paulownia witches broom. Mol Plant. 2021 Jun 29: S1674-2052(21)00233-1. Epub ahead of print |
| 721 | *Rubus chingii* *(Rosaceae)* | 2021-06-26 | Wang L et al., The chromosome-scale reference genome of Rubus chingii Hu provides insight into the biosynthetic pathway of hydrolysable tannins. Plant J. 2021 Jun 26. Epub ahead of print |
| 720 | *Prunus armeniaca* *(Rosaceae)* | 2021-06-25 | Groppi A et al., Population genomics of apricots unravels domestication history and adaptive events. Nat Commun. 12, 3956 (2021). |
| 719 | *Prunus mandshurica* *(Rosaceae)* | 2021-06-25 | Groppi A et al., Population genomics of apricots unravels domestication history and adaptive events. Nat Commun. 12, 3956 (2021). |
| 718 | *Prunus sibirica* *(Rosaceae)* | 2021-06-25 | Groppi A et al., Population genomics of apricots unravels domestication history and adaptive events. Nat Commun. 12, 3956 (2021). |
| 717 | *Acanthochlamys bracteata* *(Velloziaceae)* | 2021-06-24 | Gao ZY et al., Chromosome-scale genome assembly of the resurrection plant Acanthochlamys bracteata (Velloziaceae). Genome Biol Evol. 2021 Aug 3 13(8): evab147. |
| 716 | *Knightia excelsa* *(Proteaceae)* | 2021-06-19 | McCartney AM et al., An exploration of assembly strategies and quality metrics on the accuracy of the rewarewa (Knightia excelsa) genome. Mol Ecol Resour. 2021 Aug, 21(6): 2125-2144. Epub 2021 Jun 19. |
| 715 | *Rosa rugosa* *(Rosaceae)* | 2021-06-18 | Chen F et al., A chromosome-level genome assembly of rugged rose (Rosa rugosa) provides insights into its evolution, ecology, and floral characteristics. Hortic Res. 8, 141 (2021). |
| 714 | *Acetabularia acetabulum* *(Polyphysaceae)* | 2021-06-17 | Andresen IJ et al., Genome sequencing and de novo assembly of the giant unicellular alga Acetabularia acetabulum using droplet MDA. Sci Rep. 2021 Jun 17, 11(1): 12820. |
| 713 | *Areca catechu* *(Arecaceae)* | 2021-06-16 | Yang Y et al., Chromosome-scale genome assembly of areca palm (Areca catechu). Mol Ecol Resour. 2021 Jun 16. Epub ahead of print |
| 712 | *Lycium barbarum* *(Solanaceae)* | 2021-06-03 | Cao YL et al., Wolfberry genomes and the evolution of Lycium (Solanaceae). Commun Biol. 4, 671 (2021). |
| 711 | *Lycium ruthenicum* *(Solanaceae)* | 2021-06-03 | Cao YL et al., Wolfberry genomes and the evolution of Lycium (Solanaceae). Commun Biol. 4, 671 (2021). |
| 710 | *Puya raimondii* *(Bromeliaceae)* | 2021-06-02 | Liu L et al., Draft genome of Puya raimondii (Bromeliaceae), the Queen of the Andes. Genomics. 2021 Jul 113(4): 2537-2546. Epub 2021 Jun 2. |
| 709 | *Coptis chinensis* *(Ranunculaceae)* | 2021-06-01 | Chen D et al., The chromosome-level reference genome of Coptis chinensis provides insights into genomic evolution and berberine biosynthesis. Hortic Res. 8, 121 (2021). |
| 708 | *Morinda officinalis* *(Rubiaceae)* | 2021-06-01 | Wang J et al., A high-quality genome assembly of Morinda officinalis, a famous native southern herb in the Lingnan region of southern China. Hortic Res. 8, 135 (2021). |
| 707 | *Stevia rebaudiana* *(Asteraceae)* | 2021-06-01 | Xu X et al., The chromosome-level Stevia genome provides insights into steviol glycoside biosynthesis. Hortic Res. 8, 129 (2021). |
| 706 | *Volvox africanus* *(Volvocaceae)* | 2021-05-25 | Yamamoto K et al., Three genomes in the algal genus Volvox reveal the fate of a haploid sex-determining region after a transition to homothallism. Proc Natl Acad Sci USA. 2021 May 25, 118(21): e2100712118. |
| 705 | *Volvox reticuliferus* *(Volvocaceae)* | 2021-05-25 | Yamamoto K et al., Three genomes in the algal genus Volvox reveal the fate of a haploid sex-determining region after a transition to homothallism. Proc Natl Acad Sci USA. 2021 May 25, 118(21): e2100712118. |
| 704 | *Populus deltoides* *(Salicaceae)* | 2021-05-24 | Bai S et al., Genome Assembly of Salicaceae Populus deltoides (Eastern Cottonwood) I-69 Based on Nanopore Sequencing and Hi-C Technologies. J Hered. 2021 May 24, 112(3): 303-310. |
| 703 | *Rosa lucieae* *(Rosaceae)* | 2021-05-22 | Zhong MC et al., Rose without prickle: genomic insights linked to moisture adaptation. Natl Sci Rev. 2021 May 22, 8(12): nwab092. eCollection 2021 Dec. |
| 702 | *Collinsia rattanii* *(Plantaginaceae)* | 2021-05-20 | Frazee LJ et al., New genomic resources and comparative analyses reveal differences in floral gene expression in selfing and outcrossing Collinsia sister species. G3 (Bethesda). 2021 May 20: jkab177. Epub ahead of print |
| 701 | *Centella asiatica* *(Apiaceae)* | 2021-05-19 | Pootakham W et al., De novo chromosome-level assembly of Centella asiatica genome. Genomics. 2021 Jul 113(4): 2221-2228. Epub 2021 May 19. |
| 700 | *Gillenia trifoliata* *(Rosaceae)* | 2021-05-18 | Su W et al., Polyploidy underlies co-option and diversification of biosynthetic triterpene pathways in the apple tribe. Proc Natl Acad Sci U S A. 2021 May 18, 118(20): e2101767118. |
| 699 | *Phaseolus acutifolius* *(Fabaceae)* | 2021-05-11 | Moghaddam SM et al., The tepary bean genome provides insight into evolution and domestication under heat stress. Nat Commun. 2021 May 11, 12(1): 2638. |
| 698 | *Corymbia citriodora* *(Myrtaceae)* | 2021-05-10 | Healey AI et al., Pests, diseases, and aridity have shaped the genome of Corymbia citriodora. Commun Biol. 4, 537 (2021). |
| 697 | *Amaranthus cruentus* *(Amaranthaceae)* | 2021-05-07 | Ma X et al., A chromosome-level Amaranthus cruentus genome assembly highlights gene family evolution and biosynthetic gene clusters that may underpin the nutritional value of this traditional crop. Plant J. 2021 Jul 107(2): 613-628. Epub 2021 Jun 24. |
| 696 | *Avena strigosa* *(Poaceae)* | 2021-05-07 | Li Y et al., Subtelomeric assembly of a multi-gene pathway for antimicrobial defense compounds in cereals. Nat Commun. 2021 May 7, 12(1): 2563. |
| 695 | *Medicago ruthenica* *(Fabaceae)* | 2021-05-06 | Wang T et al., The genome of a wild Medicago species provides insights into the tolerant mechanisms of legume forage to environmental stress. BMC Biol. 2021 May 6, 19(1): 96. |
| 694 | *Miscanthus floridulus* *(Poaceae)* | 2021-05-06 | Zhang G et al., The reference genome of Miscanthus floridulus illuminates the evolution of Saccharinae. Nat Plants. 2021 May 7(5): 608-618. Epub 2021 May 6. |
| 693 | *Jacaranda mimosifolia* *(Bignoniaceae)* | 2021-05-03 | Wang M et al., Chromosomal-level reference genome of the neotropical tree Jacaranda mimosifolia D. Don. Genome Biol Evol. 2021 Jun 8 13(6): evab094. |
| 692 | *Taxus wallichiana* *(Taxaceae)* | 2021-05-02 | Cheng J et al., Chromosome-level genome of Himalayan yew provides insights into the origin and evolution of the paclitaxel biosynthetic pathway. Mol Plant. 2021 Jul 5 14(7): 1199-1209. Epub 2021 May 2. |
| 691 | *Cynodon transvaalensis* *(Poaceae)* | 2021-05-01 | Cui F et al., The genome of the warm-season turfgrass African bermudagrass (Cynodon transvaalensis). Hortic Res. 2021 May 1, 8(1): 93 |
| 690 | *Miscanthus lutarioriparius* *(Poaceae)* | 2021-04-28 | Miao J et al., Chromosome-scale assembly and analysis of biomass crop Miscanthus lutarioriparius genome. Nat Commun. 2021 Apr 28, 12(1): 2458 |
| 689 | *Korthalsia laciniosa* *(Arecaceae)* | 2021-04-20 | Ghosh Dasgupta M et al., Draft genome of Korthalsia laciniosa (Griff.) Mart., a climbing rattan elucidates its phylogenetic position. Genomics. 2021 Jul 113(4): 2010-2022. Epub 2021 Apr 20. |
| 688 | *Corylus heterophylla* *(Betulaceae)* | 2021-04-19 | Zhao T et al., A chromosome-level reference genome of the hazelnut, Corylus heterophylla Fisch. Gigascience. 2021 Apr 19, 10(4): giab027 |

| | plant species | published | |
|---|---|---|---|
| 687 | *Gossypium stocksii* (*Malvaceae*) | 2021-04-19 | Grover CE et al., The Gossypium stocksii genome as a novel resource for cotton improvement.<br>G3 (Bethesda). 2021 Apr 19, 11(7): jkab125. Online ahead of print. |
| 686 | *Cucumis metuliferus* (*Cucurbitaceae*) | 2021-04-18 | Ling J et al., High-quality chromosome-level genomes of Cucumis metuliferus and Cucumis melo provide insight into Cucumis genome evolution.<br>Plant J. 2021 Jul 107(1): 136-148. Epub 2021 May 19. |
| 685 | *Coffea humblotiana* (*Rubiaceae*) | 2021-04-14 | Raharimalala N et al., The absence of the caffeine synthase gene is involved in the naturally decaffeinated status of Coffea humblotiana, a wild species from Comoro archipelago.<br>Sci Rep. 2021 Apr 14, 11(1): 8119. |
| 684 | *Hemerocallis citrina* (*Asphodelaceae*) | 2021-04-07 | Qing Z et al., The chromosome-level Hemerocallis citrina Borani genome provides new insights into the rutin biosynthesis and the lack of colchicine.<br>Hortic Res. 2021 Apr 7, 8(1): 89. |
| 683 | *Juglans mandshurica* (*Juglandaceae*) | 2021-04-04 | Yan F et al., Improved de novo chromosome-level genome assembly of the vulnerable walnut tree Juglans mandshurica reveals gene family evolution and possible genome basis of resistance to lesion nematode.<br>Mol Ecol Resour. 2021 Aug 21(6): 2063-2076. Epub 2021 Apr 17. |
| 682 | *Solanum appendiculatum* (*Solanaceae*) | 2021-03-26 | Wu M et al., Inferring the genetic basis of sex determination from the genome of a dioecious nightshade.<br>Mol Biol Evol. 2021 Jun 25 38(7): 2946-2957. |
| 681 | *Carthamus tinctorius* (*Asteraceae*) | 2021-03-25 | Wu Z et al., The chromosome-scale reference genome of safflower (Carthamus tinctorius) provides insights into linoleic acid and flavonoid biosynthesis.<br>Plant Biotechnol J. 2021 Mar 25. Epub ahead of print |
| 680 | *Salix dunnii* (*Salicaceae*) | 2021-03-16 | He L et al., Chromosome-scale assembly of the genome of Salix dunnii reveals a male-heterogametic sex determination system on chromosome 7.<br>Mol Ecol Resour. 2021 Aug, 21(6): 1966-1982. Epub 2021 Mar 16. |
| 679 | *Selenicereus undatus* (*Cactaceae*) | 2021-03-10 | Zheng J et al., The chromosome-level genome of dragon fruit reveals whole-genome duplication and chromosomal co-localization of betacyanin biosynthetic genes.<br>Hortic Res. 8, 63 (2021). |
| 678 | *Prototheca wickerhamii* (*Chlorellaceae*) | 2021-03-09 | Bakula Z et al., A first insight into the genome of Prototheca wickerhamii, a major causative agent of human protothecosis.<br>BMC Genomics. 2021 Mar 9, 22(1): 168. |
| 677 | *Hamamelis virginiana* (*Hamamelidaceae*) | 2021-03-02 | Korgaonkar A et al., A novel family of secreted insect proteins linked to plant gall development.<br>Curr Biol. 2021 May 10, 31(9): 1836-1849.e12. Epub 2021 Mar 2. |
| 676 | *Mitragyna speciosa* (*Rubiaceae*) | 2021-03-02 | Brose J et al., The Mitragyna speciosa (Kratom) Genome: A resource for data-mining potent pharmaceuticals that impact human health.<br>G3 (Bethesda). 2021 Apr 15 11(4): jkab058. |
| 675 | *Corylus mandshurica* (*Betulaceae*) | 2021-03-01 | Li Y et al., The Corylus mandshurica genome provides insights into the evolution of Betulaceae genomes and hazelnut breeding.<br>Hortic Res. 8, 54 (2021). |
| 674 | *Cucumis hystrix* (*Cucurbitaceae*) | 2021-03-01 | Qin X et al., Chromosome-scale genome assembly of Cucumis hystrix - a wild species interspecifically cross-compatible with cultivated cucumber.<br>Hortic Res. 8, 40 (2021). |
| 673 | *Magnolia biondii* (*Magnoliaceae*) | 2021-03-01 | Dong S et al., The genome of Magnolia biondii Pamp. provides insights into the evolution of Magnoliales and biosynthesis of terpenoids.<br>Hortic Res. 8, 38 (2021). |
| 672 | *Medicago polymorpha* (*Fabaceae*) | 2021-03-01 | Cui J et al., The genome of Medicago polymorpha provides insights into its edibility and nutritional value as a vegetable and forage legume.<br>Hortic Res. 8, 47 (2021). |
| 671 | *Moricandia arvensis* (*Brassicaceae*) | 2021-03-01 | Lin MY et al., Whole genome sequencing and assembly data of Moricandia moricandioides and M. arvensis.<br>Data Brief. 2021 Mar 1, 35: 106922. eCollection 2021 Apr |
| 670 | *Moricandia moricandioides* (*Brassicaceae*) | 2021-03-01 | Lin MY et al., Whole genome sequencing and assembly data of Moricandia moricandioides and M. arvensis.<br>Data Brief. 2021 Mar 1, 35: 106922. eCollection 2021 Apr |
| 669 | *Pyrus pyrifolia* (*Rosaceae*) | 2021-02-26 | Shirasawa K et al., Chromosome-scale genome assembly of Japanese pear (Pyrus pyrifolia) variety Nijisseiki.<br>DNA Res. 2021 May 2 28(2): dsab001. |
| 668 | *Salvia bowleyana* (*Lamiaceae*) | 2021-02-26 | Zheng X et al., Insights into salvianolic acid B biosynthesis from chromosome-scale assembly of the Salvia bowleyana genome.<br>J Integr Plant Biol. 2021 Jul 63(7): 1309-1323. Epub 2021 Apr 13. |
| 667 | *Agarophyton vermiculophyllum* (*Gracilariaceae*) | 2021-02-19 | Flanagan BA et al., Founder effects shape linkage disequilibrium and genomic diversity of a partially clonal invader.<br>Mol Ecol. 2021 May, 30(9): 1962-1978. |
| 666 | *Aloe vera* (*Asphodelaceae*) | 2021-02-19 | Jaiswal SK et al., The genome sequence of Aloe vera reveals adaptive evolution of drought tolerance mechanisms.<br>iScience. 2021 Feb 19, 24(2): 102079 |
| 665 | *Cucumis x hytivus* (*Cucurbitaceae*) | 2021-02-15 | Yu X et al., Whole-Genome Sequence of Synthesized Allopolyploids in Cucumis Reveals Insights into the Genome Evolution of Allopolyploidization.<br>Adv Sci (Weinh). 2021 Feb. |
| 664 | *Cydonia oblonga* (*Rosaceae*) | 2021-02-15 | Soyturk A et al., De novo assembly and characterization of the first draft genome of quince (Cydonia oblonga Mill.).<br>Sci Rep. 11, 3818 (2021). |
| 663 | *Annona muricata* (*Annonaceae*) | 2021-02-11 | Strijk JS et al., Chromosome-level reference genome of the soursop (Annona muricata): a new resource for Magnoliid research and tropical pomology.<br>Mol Ecol Resour. 2021 Jul 21(5): 1608-1619. Epub 2021 Mar 10. |
| 662 | *Betula platyphylla* (*Betulaceae*) | 2021-02-11 | Chen S et al., Genome sequence and evolution of Betula platyphylla.<br>Hortic Res. 2021 Feb 11, 8(1): 37. |
| 661 | *Vaccinium microcarpum* (*Ericaceae*) | 2021-02-10 | Diaz-Garcia L et al., Chromosome-Level Genome Assembly of the American Cranberry (Vaccinium macrocarpon Ait.) and Its Wild Relative Vaccinium microcarpum.<br>Front Plant Sci. 2021 Feb 10, eCollection 2021. |
| 660 | *Handroanthus guayacan* (*Bignoniaceae*) | 2021-02-09 | Burley JT et al., Genome assemblies for two Neotropical trees: Jacaranda copaia and Handroanthus guayacan.<br>G3 (Bethesda). 2021 Feb 9, 11(2): jkab010. |
| 659 | *Jacaranda copaia* (*Bignoniaceae*) | 2021-02-09 | Burley JT et al., Genome assemblies for two Neotropical trees: Jacaranda copaia and Handroanthus guayacan.<br>G3 (Bethesda). 2021 Feb 9, 11(2): jkab010. |
| 658 | *Aegiceras corniculatum* (*Primulaceae*) | 2021-02-07 | Ma D et al., Chromosome-level assembly of the mangrove plant Aegiceras corniculatum genome generated through Illumina, PacBio and Hi-C sequencing technologies.<br>Mol Ecol Resour. 2021 Jul 21(5): 1593-1607. Epub 2021 Mar 8. |
| 657 | *Aeschynomene evenia* (*Fabaceae*) | 2021-02-05 | Quilbe J et al., Genetics of nodulation in Aeschynomene evenia uncovers mechanisms of the rhizobium-legume symbiosis.<br>Nat Commun. 2021 Feb 5, 12(1): 829. |
| 656 | *Ostreobium quekettii* (*Ostreobiaceae*) | 2021-02-05 | Iha C et al., Genomic adaptations to an endolithic lifestyle in the coral-associated alga Ostreobium.<br>Curr Biol. 2021 Apr 12, 31(7): 1393-1402.e5. Epub 2021 Feb 5. |
| 655 | *Vitis rotundifolia* (*Vitaceae*) | 2021-02-05 | Cochetel N et al., Diploid chromosome-scale assembly of the Muscadinia rotundifolia genome supports chromosome fusion and disease resistance gene expansion during Vitis and Muscadinia divergence.<br>G3 (Bethesda). 2021 Feb 5. |
| 654 | *Brassica carinata* (*Brassicaceae*) | 2021-02-04 | Song X et al., Brassica carinata genome characterization clarifies Us triangle model of evolution and polyploidy in Brassica<br>Plant Physiol. 2021 Feb 4 186(1): 388-406. Epub ahead of print |
| 653 | *Oryza alta* (*Poaceae*) | 2021-02-03 | Yu H et al., A route to de novo domestication of wild allotetraploid rice.<br>Cell. 2021 Mar 4 184(5): 1156-1170.e14. Epub 2021 Feb 3. |
| 652 | *Chlamydomonas incerta* (*Chlamydomonadaceae*) | 2021-02-02 | Craig RJ et al., Comparative genomics of Chlamydomonas.<br>Plant Cell. 2021, koab026, Epub ahead of print |

| | plant species | published | |
|---|---|---|---|
| 651 | *Chlamydomonas schloesseri* *(Chlamydomonadaceae)* | 2021-02-02 | Craig RJ et al., Comparative genomics of Chlamydomonas. Plant Cell. 2021, koab026, Epub ahead of print |
| 650 | *Edaphochlamys debaryana* *(C.incertae sedis)* | 2021-02-02 | Craig RJ et al., Comparative genomics of Chlamydomonas. Plant Cell. 2021, koab026, Epub ahead of print |
| 649 | *Sechium edule* *(Cucurbitaceae)* | 2021-01-31 | Fu A et al., Combined genomic, transcriptomic, and metabolomic analyses provide insights into chayote (Sechium edule) evolution and fruit development. Hortic Res. 8, 35 (2021) |
| 648 | *Solanum sitiens* *(Solanaceae)* | 2021-01-30 | Molitor C et al., De Novo Genome Assembly Of Solanum sitiens Reveals Structural Variation Associated With Drought And Salinity Tolerance. Bioinformatics. 2021 Jan 30: btab048. Epub ahead of print |
| 647 | *Phaseolus lunatus* *(Fabaceae)* | 2021-01-29 | Garcia T et al., Comprehensive genomic resources related to domestication and crop improvement traits in Lima bean. Nat Commun. 12, 702 (2021) |
| 646 | *Thalictrum thalictroides* *(Ranunculaceae)* | 2021-01-29 | Arias T et al., Genomic and transcriptomic resources for candidate gene discovery in the Ranunculids. Appl Plant Sci. 2021 Jan 29, 9(1): e11407. eCollection 2021 Jan. |
| 645 | *Panicum virgatum* *(Poaceae)* | 2021-01-27 | Lovell JT et al., Genomic mechanisms of climate adaptation in polyploid bioenergy switchgrass. Nature. 2021 Feb 590(7846): 438-444. Epub 2021 Jan 27. |
| 644 | *Pharus latifolius* *(Poaceae)* | 2021-01-25 | Ma PF et al., The Pharus latifolius genome bridges the gap of early grass evolution. Plant Cell. 2021, koab015 |
| 643 | *Sapria himalayana* *(Rafflesiaceae)* | 2021-01-22 | Cai L et al., Deeply Altered Genome Architecture in the Endoparasitic Flowering Plant Sapria himalayana Griff. (Rafflesiaceae). Curr Biol. 2021 Mar 8, Epub ahead of print |
| 642 | *Chlamydomonas sp. UWO241* *(Chlamydomonadaceae)* | 2021-01-20 | Zhang X et al., Draft genome sequence of the Antarctic green alga Chlamydomonas sp. UWO241. iScience. 2021 Jan 20, 24(2): 102084. |
| 641 | *Ophiorrhiza pumila* *(Rubiaceae)* | 2021-01-15 | Rai A et al., Chromosome-level genome assembly of Ophiorrhiza pumila reveals the evolution of camptothecin biosynthesis. Nat Commun. 2021 Jan 15, 12(1): 405. |
| 640 | *Datura stramonium* *(Solanaceae)* | 2021-01-13 | De-la-Cruz IM et al., Genomic signatures of the evolution of defence against its natural enemies in the poisonous and medicinal plant Datura stramonium (Solanaceae). Sci Rep. 2021 Jan 13, 11(1): 882. |
| 639 | *Dioscorea zingiberensis* *(Dioscoreaceae)* | 2021-01-11 | Cheng J et al., The origin and evolution of the diosgenin biosynthetic pathway in yam. Plant Commun. 2020 Jun 2, 2(1): 100079. eCollection 2021 Jan 11. |
| 638 | *Toona sinensis* *(Meliaceae)* | 2021-01-09 | Ji YT et al., Long read sequencing of Toona sinensis (A. Juss) Roem: a chromosome-level reference genome for the Meliaceae family. Mol Ecol Resour. 2021 May 21(4): 1243-1255. Epub 2021 Jan 27. |
| 637 | *Raddia distichophylla* *(Poaceae)* | 2021-01-06 | Li W et al., Draft genome of the herbaceous bamboo Raddia distichophylla. G3 (Bethesda). 2021 Jan 6 |
| 636 | *Akebia trifoliata* *(Lardizabalaceae)* | 2021-01-05 | Zhang Z et al., Genome survey sequencing and genetic diversity of cultivated Akebia trifoliata assessed via phenotypes and SSR markers. Mol Biol Rep. 2021 Jan 48(1): 241-250. Epub 2021 Jan 5. |
| 635 | *Prunus domestica* *(Rosaceae)* | 2021-01-01 | Callahan AM et al., Defining the HoneySweet insertion event utilizing NextGen sequencing and a de novo genome assembly of plum (Prunus domestica). Hortic Res. 2021 Jan 1, 8(1): 8. |
| 634 | *Wolffia australiana* *(Araceae)* | 2020-12-23 | Michael TP et al., Genome and time-of-day transcriptome of Wolffia australiana link morphological minimization with gene loss and less growth control. Genome Res. 2020 Dec 23 31(2): 225-38. Epub ahead of print |
| 633 | *Passiflora edulis* *(Passifloraceae)* | 2020-12-16 | Ma D et al., Chromosome-level reference genome assembly provides insights into aroma biosynthesis in passion fruit (Passiflora edulis). Mol Ecol Resour. 2021 Apr 21(3): 955-968. Epub 2020 Dec 28. |
| 632 | *Hypericum perforatum* *(Hypericaceae)* | 2020-12-14 | Zhou W et al., Whole genome sequence data of Hypericum perforatum and functional characterization of melatonin biosynthesis by N-acetylserotonin O-methyltransferase. J Pineal Res. 2021 Mar 70(2): e12709. Epub 2020 Dec 25. |
| 631 | *Vanilla planifolia* *(Orchidaceae)* | 2020-12-11 | Hasing T et al., A phased Vanilla planifolia genome enables genetic improvement of flavour and production. Nat Food. 1, 811-819 (2020). |
| 630 | *Prunus salicina* *(Rosaceae)* | 2020-12-10 | Liu C et al., Chromosome-level draft genome of a diploid plum (Prunus salicina). Gigascience. 2020 Dec 10, 9(12): giaa130. |
| 629 | *Vitis amurensis* *(Vitaceae)* | 2020-12-10 | Wang Y et al., The genome of Shanputao (Vitis amurensis) provides a new insight into cold tolerance of grapevine. Plant J. 2021 Mar 105(6): 1495-1506. Epub 2021 Jan 21. |
| 628 | *Dendrobium huoshanense* *(Orchidaceae)* | 2020-12-06 | Han B et al., A Chromosome-Level Genome Assembly of Dendrobium Huoshanense Using Long Reads and Hi-C Data. Genome Biol Evol. 2020 Dec 6, 12(12): 2486-2490. |
| 627 | *Syntrichia caninervis* *(Pottiaceae)* | 2020-12-04 | Silva AT et al., To dry perchance to live: Insights from the genome of the desiccation-tolerant biocrust moss Syntrichia caninervis. Plant J. 2021 Mar 105(5): 1339-1356. Epub 2020 Dec 20. |
| 626 | *Tetracentron sinense* *(Trochodendraceae)* | 2020-12-02 | Liu PL et al., The Tetracentron genome provides insight into the early evolution of eudicots and the formation of vessel elements. Genome Biol. 2020 Dec 2, 21(1): 291. |
| 625 | *Lobularia maritima* *(Brassicaceae)* | 2020-12-01 | Huang L et al., A chromosome-scale reference genome of Lobularia maritima, an ornamental plant with high stress tolerance. Hortic Res. 7, 197 (2020). |
| 624 | *Salix matsudana* *(Salicaceae)* | 2020-12-01 | Zhang J et al., Genome sequencing and phylogenetic analysis of allotetraploid Salix matsudana Koidz. Hortic Res. 7, 201 (2020). |
| 623 | *Tapiscia sinensis* *(Tapisciaceae)* | 2020-12-01 | Zhao P et al., The de novo genome assembly of Tapiscia sinensis and the transcriptomic and developmental bases of androdioecy. Hortic Res. 7, 191 (2020). |
| 622 | *Trichosanthes anguina* *(Cucurbitaceae)* | 2020-12-01 | Ma L et al., The genome and transcriptome analysis of snake gourd provide insights into its evolution and fruit development and ripening. Hortic Res. 7, 199 (2020). |
| 621 | *Corylus avellana* *(Betulaceae)* | 2020-11-29 | Lucas SJ et al., A chromosome-scale genome assembly of European Hazel (Corylus avellana L.) reveals targets for crop improvement. Plant J. 2021 Mar 105(5): 1413-1430. Epub 2021 Jan 6. |
| 620 | *Amphicarpaea edgeworthii* *(Fabaceae)* | 2020-11-25 | Liu Y et al., Insights into amphicarpy from the compact genome of the legume Amphicarpaea edgeworthii. Plant Biotechnol J. 2021 May 19(5): 952-965. Epub 2020 Dec 11. |
| 619 | *Senna tora* *(Fabaceae)* | 2020-11-18 | Kang SH et al., Genome-enabled discovery of anthraquinone biosynthesis in Senna tora. Nat Commun. 2020 Nov 18, 11(1): 5875. |
| 618 | *Fontinalis antipyretica* *(Fontinalaceae)* | 2020-11-16 | Yu J et al., Draft genome of the aquatic moss Fontinalis antipyretica (Fontinalaceae, Bryophyta). Gigabyte, 1, 2020 |
| 617 | *Hydrangea macrophylla* *(Hydrangeaceae)* | 2020-11-11 | Nashima K et al., Genome sequence of Hydrangea macrophylla and its application in analysis of the double flower phenotype. DNA Res. 2021 Jan 19 28(1): dsaa026. |
| 616 | *Megadenia pygmaea* *(Brassicaceae)* | 2020-11-05 | Yang W et al., The chromosome-level genome sequence and karyotypic evolution of Megadenia pygmaea (Brassicaceae). Mol Ecol Resour. 2021 Apr 21(3): 871-879. Epub 2020 Dec 1. |
| 615 | *Spirodela intermedia* *(Araceae)* | 2020-11-05 | Hoang PTN et al., Chromosome-scale genome assembly for the duckweed Spirodela intermedia, integrating cytogenetic maps, PacBio and Oxford Nanopore libraries. Sci Rep. 10, 19230 (2020). |

| | plant species | published | |
|---|---|---|---|
| 614 | *Scutellaria barbata* (Lamiaceae) | 2020-11-04 | Xu Z et al., Comparative Genome Analysis of Scutellaria baicalensis and Scutellaria barbata Reveals the Evolution of Active Flavonoid Biosynthesis.<br>Genomics Proteomics Bioinformatics. 2020 Jun 18(3): 230-240. Epub 2020 Nov 4. |
| 613 | *Malus sylvestris* (Rosaceae) | 2020-11-02 | Sun X et al., Phased diploid genome assemblies and pan-genomes provide insights into the genetic history of apple domestication.<br>Nat Genet. 2020 Dec 52(12): 1423-1432. Epub 2020 Nov 2. |
| 612 | *Miscanthus sinensis* (Poaceae) | 2020-10-28 | Mitros T et al., Genome biology of the paleotetraploid perennial biomass crop Miscanthus.<br>Nat Commun. 2020 Oct 28, 11(1): 5442 |
| 611 | *Phtheirospermum japonicum* (Orobanchaceae) | 2020-10-28 | Cui S et al., Ethylene signaling mediates host invasion by parasitic plants.<br>Sci Adv. 2020 Oct 28, 6(44): eabc2385. Print 2020 Oct. |
| 610 | *Strobilanthes cusia* (Acanthaceae) | 2020-10-28 | Xu W et al., Blue genome: chromosome-scale genome reveals the evolutionary and molecular basis of indigo biosynthesis in Strobilanthes cusia.<br>Plant J. 2020 Nov, 104(4): 864-879. Epub 2020 Oct 28. |
| 609 | *Eruca sativa* (Brassicaceae) | 2020-10-27 | Bell L et al., The Eruca sativa Genome and Transcriptome: A Targeted Analysis of Sulfur Metabolism and Glucosinolate Biosynthesis Pre and Postharvest.<br>Front Plant Sci. 11: 525102 |
| 608 | *Psidium guajava* (Myrtaceae) | 2020-10-24 | Feng C et al., A chromosome-level genome assembly provides insights into ascorbic acid accumulation and fruit softening in guava (Psidium guajava).<br>Plant Biotechnol J. 2021 Apr 19(4): 717-730. Epub 2020 Nov 12. |
| 607 | *Rumex hastatulus* (Polygonaceae) | 2020-10-23 | Rifkin JL et al., Widespread Recombination Suppression Facilitates Plant Sex Chromosome Evolution.<br>Mol Biol Evol. 2021 Mar 9 38(3): 1018-1030. |
| 606 | *Rhododendron simsii* (Ericaceae) | 2020-10-19 | Yang F et al., Chromosome-level genome assembly of a parent species of widely cultivated azaleas.<br>Nat Commun. 11, 5269 (2020) |
| 605 | *Draba nivalis* (Brassicaceae) | 2020-10-15 | Nowak MD et al., The genome of Draba nivalis shows signatures of adaptation to the extreme environmental stresses of the Arctic.<br>Mol Ecol Resour. 2021 Apr 21(3): 661-676. Epub 2020 Nov 12. |
| 604 | *Cenchrus purpureus* (Poaceae) | 2020-10-11 | Yan Q et al., The elephant grass (Cenchrus purpureus) genome provides insights into anthocyanidin accumulation and fast growth.<br>Mol Ecol Resour. 2021 Feb 21(2): 526-542. Epub 2020 Oct 28. |
| 603 | *Ficus hispida* (Moraceae) | 2020-10-08 | Zhang X et al., Genomes of the Banyan Tree and Pollinator Wasp Provide Insights into Fig-Wasp Coevolution.<br>Cell. 2020 Nov 12 183(4): 875-889.e17. Epub 2020 Oct 8. |
| 602 | *Ficus microcarpa* (Moraceae) | 2020-10-08 | Zhang X et al., Genomes of the Banyan Tree and Pollinator Wasp Provide Insights into Fig-Wasp Coevolution.<br>Cell. 2020 Nov 12 183(4): 875-889.e17. Epub 2020 Oct 8. |
| 601 | *Mesua ferrea* (Calophyllaceae) | 2020-10-08 | Patil AB et al., The genome sequence of Mesua ferrea and comparative demographic histories of forest trees.<br>Gene. 2021 Feb 15 769: 145214. Epub 2020 Oct 8. |
| 600 | *Prunus humilis* (Rosaceae) | 2020-10-06 | Wang P et al., Chromosome-Level Genome Assembly of Cerasus humilis Using PacBio and Hi-C Technologies.<br>Front Genet. 2020 Oct 6, 11: 956. eCollection 2020. |
| 599 | *Colocasia esculenta* (Araceae) | 2020-10-04 | Yin J et al., A high-quality genome of taro (Colocasia esculenta (L.) Schott), one of the world's oldest crops.<br>Mol Ecol Resour. 2021 Jan, 21(1): 68-77. Epub 2020 Oct 4. |
| 598 | *Prunus serrulata* (Rosaceae) | 2020-10-01 | Yi X et al., The genome of Chinese flowering cherry (Cerasus serrulata) provides new insights into Cerasus species.<br>Hortic Res. 7, 165 (2020) |
| 597 | *Citrus trifoliata* (Rutaceae) | 2020-09-27 | Peng Z et al., A Chromosome-scale Reference Genome of Trifoliate Orange (Poncirus trifoliata) Provides Insights into Disease Resistance, Cold Tolerance and Genome Evolution in Citrus.<br>Plant J. 2020 Dec 104(5): 1215-1232. Epub 2020 Oct 18. |
| 596 | *Cardamine resedifolia* (Brassicaceae) | 2020-09-24 | Rellstab C et al., Genomic signatures of convergent adaptation to Alpine environments in three Brassicaceae species.<br>Mol Ecol. 2020 Nov 29(22): 4350-4365. Epub 2020 Oct 14. |
| 595 | *Cleistogenes songorica* (Poaceae) | 2020-09-22 | Zhang J et al., The genome of Cleistogenes songorica provides a blueprint for functional dissection of dimorphic flower differentiation and drought adaptability.<br>Plant Biotechnol J. 2021 Mar 19(3): 532-547. Epub 2020 Oct 28. |
| 594 | *Sequoiadendron giganteum* (Cupressaceae) | 2020-09-18 | Scott AD et al., A Reference Genome Sequence for Giant Sequoia.<br>G3 (Bethesda). 2020 Nov 5 10(11): 3907-3919. |
| 593 | *Fragaria nubicola* (Rosaceae) | 2020-09-17 | Feng C et al., Tracing the Diploid Ancestry of the Cultivated Octoploid Strawberry.<br>Mol Biol Evol. 2021 Jan 23, 38(2): 478-485. |
| 592 | *Fragaria viridis* (Rosaceae) | 2020-09-17 | Feng C et al., Tracing the Diploid Ancestry of the Cultivated Octoploid Strawberry.<br>Mol Biol Evol. 2021 Jan 23, 38(2): 478-485. |
| 591 | *Clethra arborea* (Clethraceae) | 2020-09-10 | Hartmann S et al., Annotated genome sequences of the carnivorous plant Roridula gorgonias and a non-carnivorous relative, Clethra arborea.<br>BMC Res Notes. 2020 Sep 10, 13(1): 426 |
| 590 | *Roridula gorgonias* (Roridulaceae) | 2020-09-10 | Hartmann S et al., Annotated genome sequences of the carnivorous plant Roridula gorgonias and a non-carnivorous relative, Clethra arborea.<br>BMC Res Notes. 2020 Sep 10, 13(1): 426 |
| 589 | *Digitaria exilis* (Poaceae) | 2020-09-08 | Abrouk M et al., Fonio millet genome unlocks African orphan crop diversity for agriculture in a changing climate.<br>Nat Commun. 2020 Sep 8, 11(1): 4488 |
| 588 | *Callicarpa americana* (Lamiaceae) | 2020-09-07 | Hamilton JP et al., Generation of a chromosome-scale genome assembly of the insect-repellent terpenoid-producing Lamiaceae species, Callicarpa americana.<br>Gigascience. 2020 Sep 1, 9(9): giaa093 |
| 587 | *Nyssa yunnanensis* (Nyssaceae) | 2020-09-01 | Mu W et al., The draft genome assembly of the critically endangered Nyssa yunnanensis, a plant species with extremely small populations endemic to Yunnan Province, China.<br>Gigabyte, 1, 2020 |
| 586 | *Phoebe bournei* (Lauraceae) | 2020-09-01 | Chen SP et al., The Phoebe genome sheds light on the evolution of magnoliids.<br>Hortic Res. 2020 Sep 1, 7(1): 146. |
| 585 | *Amaranthus hybridus* (Amaranthaceae) | 2020-08-24 | Montgomery JS et al., Draft genomes of Amaranthus tuberculatus, Amaranthus hybridus and Amaranthus palmeri.<br>Genome Biol Evol. 2020 Nov 3 12(11): 1988-1993. |
| 584 | *Amaranthus palmeri* (Amaranthaceae) | 2020-08-24 | Montgomery JS et al., Draft genomes of Amaranthus tuberculatus, Amaranthus hybridus and Amaranthus palmeri.<br>Genome Biol Evol. 2020 Nov 3 12(11): 1988-1993. |
| 583 | *Dalbergia odorifera* (Fabaceae) | 2020-08-18 | Hong Z et al., The chromosome-level draft genome of Dalbergia odorifera.<br>Gigascience. 2020 Aug 18, 9(8) |
| 582 | *Cercidiphyllum japonicum* (Cercidiphyllaceae) | 2020-08-17 | Zhu S et al., Genomic insights on the contribution of balancing selection and local adaptation to the long-term survival of a widespread living fossil tree, Cercidiphyllum japonicum.<br>New Phytol. 2020 Dec, 228(5): 1674-1689. Epub 2020 Aug 17. |
| 581 | *Coelastrum proboscideum* (Scenedesmaceae ) | 2020-08-17 | Liang H et al., The Draft Genome of Coelastrum proboscideum (Sphaeropleales, Chlorophyta).<br>Protist. 2020 Nov 171(5): 125758. Epub 2020 Aug 17. |
| 580 | *Vigna mungo* (Fabaceae) | 2020-08-13 | Pootakham W et al., A chromosome-scale assembly of the black gram (Vigna mungo) genome.<br>Mol Ecol Resour. 2021 Jan 21(1): 238-250. Epub 2020 Sep 8. |
| 579 | *Chimonanthus praecox* (Calycanthaceae) | 2020-08-10 | Shang J et al., The chromosome-level wintersweet (Chimonanthus praecox) genome provides insights into floral scent biosynthesis and flowering in winter.<br>Genome Biol. 2020 Aug 10, 21(1): 200 |
| 578 | *Acer truncatum* (Sapindaceae) | 2020-08-09 | Ma Q et al., The Acer truncatum genome provides insights into the nervonic acid biosynthesis.<br>Plant J. 2020 Nov 104(3): 662-678. Epub 2020 Oct 2. |

| | plant species | published | |
|---|---|---|---|
| 577 | *Luffa acutangula* (*Cucurbitaceae*) | 2020-08-09 | Pootakham W et al., De novo assemblies of Luffa acutangula and Luffa cylindrica genomes reveal an expansion associated with substantial accumulation of transposable elements.<br>Mol Ecol Resour. 2021 Jan 21(1): 212-225. Epub 2020 Aug 25. |
| 576 | *Megacarpaea delavayi* (*Brassicaceae*) | 2020-08-03 | Yang Q et al., The Genome Sequence of Alpine Megacarpaea delavayi Identifies Species-Specific Whole-Genome Duplication.<br>Front Genet. 11: 812 (Aug 2020) |
| 575 | *Forsythia suspensa* (*Oleaceae*) | 2020-08-01 | Li L et al., Genome sequencing and population genomics modeling provide insights into the local adaptation of weeping forsythia.<br>Hortic Res. 7, 130 (2020) |
| 574 | *Ocimum basilicum* (*Lamiaceae*) | 2020-07-31 | Bornowski N et al., Genome sequencing of four culinary herbs reveals terpenoid genes underlying chemodiversity in the Nepetoideae.<br>DNA Res. 2020 Jun 1 27(3): dsaa016. |
| 573 | *Origanum majorana* (*Lamiaceae*) | 2020-07-31 | Bornowski N et al., Genome sequencing of four culinary herbs reveals terpenoid genes underlying chemodiversity in the Nepetoideae.<br>DNA Res. 2020 Jun 1 27(3): dsaa016. |
| 572 | *Origanum vulgare* (*Lamiaceae*) | 2020-07-31 | Bornowski N et al., Genome sequencing of four culinary herbs reveals terpenoid genes underlying chemodiversity in the Nepetoideae.<br>DNA Res. 2020 Jun 1 27(3): dsaa016. |
| 571 | *Salvia rosmarinus* (*Lamiaceae*) | 2020-07-31 | Bornowski N et al., Genome sequencing of four culinary herbs reveals terpenoid genes underlying chemodiversity in the Nepetoideae.<br>DNA Res. 2020 Jun 1 27(3): dsaa016. |
| 570 | *Brachypodium hybridum* (*Poaceae*) | 2020-07-29 | Gordon SP et al., Gradual polyploid genome evolution revealed by pan-genomic analysis of Brachypodium hybridum and its diploid progenitors.<br>Nat Commun. 2020 Jul 29, 11(1): 3670 |
| 569 | *Brachypodium stacei* (*Poaceae*) | 2020-07-29 | Gordon SP et al., Gradual polyploid genome evolution revealed by pan-genomic analysis of Brachypodium hybridum and its diploid progenitors.<br>Nat Commun. 2020 Jul 29, 11(1): 3670 |
| 568 | *Cyanidiococcus yangmingshanensis* (*Cyanidioschyzonaceae*) | 2020-07-29 | Liu SL et al., Comparative genome analysis reveals Cyanidiococcus gen. nov., a new extremophilic red algal genus sister to Cyanidioschyzon (Cyanidioschyzonaceae, Rhodophyta).<br>J Phycol. 2020 Jul 29. Online ahead of print. |
| 567 | *Allium sativum* (*Amaryllidaceae*) | 2020-07-27 | Sun X et al., A chromosome-level genome assembly of garlic (Allium sativum L.) provides insights into genome evolution and allicin biosynthesis.<br>Mol Plant. 2020 Sep 7 13(9): 1328-1339. Epub 2020 Jul 28. |
| 566 | *Setaria viridis* (*Poaceae*) | 2020-07-21 | Thielen PM et al., Reference Genome for the Highly Transformable Setaria viridis ME034V.<br>G3 (Bethesda). 2020 Oct 5 10(10): 3467-3478. |
| 565 | *Mychonastes homosphaera* (*Mychonastaceae*) | 2020-07-11 | Liu C et al., Genome analyses provide insights into the evolution and adaptation of the eukaryotic Picophytoplankton Mychonastes homosphaera.<br>BMC Genomics. 2020 Jul 11, 21(1): 477 |
| 564 | *Chiococca alba* (*Rubiaceae*) | 2020-07-08 | Lau KH et al., Genome assembly of Chiococca alba uncovers key enzymes involved in the biosynthesis of unusual terpenoids.<br>DNA Res. 2020 Jun 1 27(3): dsaa013. |
| 563 | *Microthlaspi erraticum* (*Brassicaceae*) | 2020-07-03 | Mishra B et al., The Genome of Microthlaspi erraticum (Brassicaceae) Provides Insights Into the Adaptation to Highly Calcareous Soils.<br>Front Plant Sci. 03 July 2020 |
| 562 | *Chlamydomonas sp. ICE-L* (*Chlamydomonadaceae*) | 2020-07-02 | Zhang Z et al., Adaptation to Extreme Antarctic Environments Revealed by the Genome of a Sea Ice Green Alga.<br>Curr Biol. 2020 Sep 7 30(17): 3330-3341.e7. Epub 2020 Jul 2. |
| 561 | *Echinochloa haploclada* (*Poaceae*) | 2020-07-02 | Ye CY et al., The Genomes of the Allohexaploid Echinochloa crus-galli and Its Progenitors Provide Insights into Polyploidization-Driven Adaptation.<br>Mol Plant. 2020 Sep 7, 13(9): 1298-1310. Epub 2020 Jul 2. |
| 560 | *Echinochloa oryzoides* (*Poaceae*) | 2020-07-02 | Ye CY et al., The Genomes of the Allohexaploid Echinochloa crus-galli and Its Progenitors Provide Insights into Polyploidization-Driven Adaptation.<br>Mol Plant. 2020 Sep 7, 13(9): 1298-1310. Epub 2020 Jul 2. |
| 559 | *Aquilegia oxysepala* (*Ranunculaceae*) | 2020-07-01 | Xie J et al., A chromosome-scale reference genome of Aquilegia oxysepala var. kansuensis.<br>Hortic Res. 7, 113 (2020) |
| 558 | *Platycodon grandiflorus* (*Campanulaceae*) | 2020-07-01 | Kim J et al., Whole-genome, transcriptome, and methylome analyses provide insights into the evolution of platycoside biosynthesis in Platycodon grandiflorus, a medicinal plant.<br>Hortic Res. 7, 112 (2020) |
| 557 | *Salix viminalis* (*Salicaceae*) | 2020-06-30 | Almeida P et al., Genome assembly of the basket willow, Salix viminalis, reveals earliest stages of sex chromosome expansion.<br>BMC Biol. 2020 Jun 30, 18(1): 78 |
| 556 | *Prasinoderma coloniale* (*Prasinodermataceae*) | 2020-06-22 | Li L et al., The Genome of Prasinoderma Coloniale Unveils the Existence of a Third Phylum Within Green Plants.<br>Nat Ecol Evol. 2020 Sep 4(9): 1220-1231. Epub 2020 Jun 22. |
| 555 | *Gardenia jasminoides* (*Rubiaceae*) | 2020-06-18 | Xu Z et al., Tandem gene duplications drive divergent evolution of caffeine and crocin biosynthetic pathways in plants.<br>BMC Biol. 18, 63 (2020) |
| 554 | *Carex littledalei* (*Cyperaceae*) | 2020-06-11 | Can M et al., Genome Sequence of Kobresia Littledalei, the First Chromosome-Level Genome in the Family Cyperaceae.<br>Sci Data. 2020 Jun 11, 7(1): 175 |
| 553 | *Chimonanthus salicifolius* (*Calycanthaceae*) | 2020-06-10 | Lv Q et al., The Chimonanthus Salicifolius Genome Provides Insight Into Magnoliid Evolution and Flavonoid Biosynthesis.<br>Plant J. 2020 Aug 103(5): 1910-1923. Epub 2020 Jul 10. |
| 552 | *Vitis arizonica* (*Vitaceae*) | 2020-06-09 | Massonnet M et al., The genetic basis of sex determination in grapes.<br>Nat Commun. 2020 Jun 9, 11(1): 2902. |
| 551 | *Averrhoa carambola* (*Oxalidaceae*) | 2020-06-01 | Fan Y et al., Dissecting the genome of star fruit (Averrhoa carambola L.).<br>Hortic Res. 2020 Jun 1 7: 94. eCollection 2020. |
| 550 | *Lithospermum erythrorhizon* (*Boraginaceae*) | 2020-06-01 | Auber RP et al., Hybrid de novo genome assembly of red gromwell (Lithospermum erythrorhizon) reveals evolutionary insight into shikonin biosynthesis.<br>Hortic Res. 2020 Jun 1 7: 82. eCollection 2020. |
| 549 | *Atriplex hortensis* (*Amaranthaceae*) | 2020-05-25 | Hunt SP et al., A Chromosome-Scale Assembly of the Garden Orach (Atriplex hortensis L.) Genome Using Oxford Nanopore Sequencing.<br>Front Plant Sci. 25 May 2020 |
| 548 | *Kingdonia uniflora* (*Circaeasteraceae*) | 2020-05-22 | Sun Y et al., Genome sequencing of the endangered Kingdonia uniflora (Circaeasteraceae, Ranunculales) reveals potential mechanisms of evolutionary specialization.<br>iScience. 2020 May 22 23(5): 101124. Epub 2020 May 1. |
| 547 | *Penium margaritaceum* (*Peniaceae*) | 2020-05-21 | Jiao C et al., The Penium margaritaceum Genome: Hallmarks of the Origins of Land Plants.<br>Cell. 2020 May 28 181(5): 1097-1111.e12. Epub 2020 May 21. |
| 546 | *Argania spinosa* (*Sapotaceae*) | 2020-05-19 | Khayi S et al., First draft genome assembly of the Argane tree (Argania spinosa).<br>F1000Res. 2020 May 19, 7: 1310 |
| 545 | *Medicago sativa* (*Fabaceae*) | 2020-05-19 | Chen H et al., Allele-aware Chromosome-Level Genome Assembly and Efficient Transgene-Free Genome Editing for the Autotetraploid Cultivated Alfalfa.<br>Nat Commun. 2020 May 19, 11(1): 2494 |
| 544 | *Coix lacryma-jobi* (*Poaceae*) | 2020-05-18 | Kang SH et al., Genome Assembly and Annotation of Soft-Shelled Adlay (Coix lacryma-jobi Variety ma-yuen), a Cereal and Medicinal Crop in the Poaceae Family.<br>Front Plant Sci. 2020 May 18 |
| 543 | *Morus alba* (*Moraceae*) | 2020-05-16 | Jiao F et al., Chromosome-Level Reference Genome and Population Genomic Analysis Provide Insights into the Evolution and Improvement of Domesticated Mulberry (Morus alba).<br>Mol Plant. 2020 Jul 6, 13(7): 1001-1012. Epub 2020 May 16. |
| 542 | *Calohypnum plumiforme* (*Hypnaceae*) | 2020-05-14 | Mao L et al., Genomic evidence for convergent evolution of gene clusters for momilactone biosynthesis in land plants.<br>Proc Natl Acad Sci U S A. 2020 Jun 2 117(22): 12472-12480. Epub 2020 May 14. |
| 541 | *Primulina huaijiensis* (*Gesneriaceae*) | 2020-05-14 | Feng C et al., The genome of a cave plant, Primulina huaijiensis, provides insights into adaptation to limestone karst habitats.<br>New Phytol. 2020 Aug, 227(4): 1249-1263 |

| | plant species | published | |
|---|---|---|---|
| 540 | *Hyssopus officinalis* *(Lamiaceae)* | 2020-05-13 | Lichman BR et al., The evolutionary origins of the cat attractant nepetalactone in catnip. Sci Adv. 2020 May 13, 6(20): eaba0721. eCollection 2020 May. |
| 539 | *Nepeta cataria* *(Lamiaceae)* | 2020-05-13 | Lichman BR et al., The evolutionary origins of the cat attractant nepetalactone in catnip. Sci Adv. 2020 May 13, 6(20): eaba0721. eCollection 2020 May. |
| 538 | *Nepeta racemosa* *(Lamiaceae)* | 2020-05-13 | Lichman BR et al., The evolutionary origins of the cat attractant nepetalactone in catnip. Sci Adv. 2020 May 13, 6(20): eaba0721. eCollection 2020 May. |
| 537 | *Paeonia suffruticosa* *(Paeoniaceae)* | 2020-05-12 | Lv S et al., Draft genome of the famous ornamental plant Paeonia suffruticosa. Ecol Evol. 2020, 00: 1-13 |
| 536 | *Aldrovanda vesiculosa* *(Droseraceae)* | 2020-05-11 | Palfalvi G et al., Genomes of the Venus Flytrap and Close Relatives Unveil the Roots of Plant Carnivory. Curr Biol. 2020 Jun 22 30(12): 2312-2320.e5. Epub 2020 May 14. |
| 535 | *Dionaea muscipula* *(Droseraceae)* | 2020-05-11 | Palfalvi G et al., Genomes of the Venus Flytrap and Close Relatives Unveil the Roots of Plant Carnivory. Curr Biol. 2020 Jun 22 30(12): 2312-2320.e5. Epub 2020 May 14. |
| 534 | *Drosera spatulata* *(Droseraceae)* | 2020-05-11 | Palfalvi G et al., Genomes of the Venus Flytrap and Close Relatives Unveil the Roots of Plant Carnivory. Curr Biol. 2020 Jun 22 30(12): 2312-2320.e5. Epub 2020 May 14. |
| 533 | *Kandelia obovata* *(Rhizophoraceae)* | 2020-05-02 | Hu MJ et al., Chromosome-scale assembly of the Kandelia obovata genome. Hortic Res. 2020 May 2, 7: 75 |
| 532 | *Eragrostis nindensis* *(Poaceae)* | 2020-04-23 | Pardo J et al., Intertwined signatures of desiccation and drought tolerance in grasses. Proc Natl Acad Sci U S A. 2020 May 5 117(18): 10079-10088. Epub 2020 Apr 23. |
| 531 | *Gossypium darwinii* *(Malvaceae)* | 2020-04-20 | Chen ZJ et al., Genomic diversifications of five Gossypium allopolyploid species and their impact on cotton improvement. Nat Genet. 2020 May 52(5): 525-533. Epub 2020 Apr 20. |
| 530 | *Gossypium mustelinum* *(Malvaceae)* | 2020-04-20 | Chen ZJ et al., Genomic diversifications of five Gossypium allopolyploid species and their impact on cotton improvement. Nat Genet. 2020 May 52(5): 525-533. Epub 2020 Apr 20. |
| 529 | *Gossypium tomentosum* *(Malvaceae)* | 2020-04-20 | Chen ZJ et al., Genomic diversifications of five Gossypium allopolyploid species and their impact on cotton improvement. Nat Genet. 2020 May 52(5): 525-533. Epub 2020 Apr 20. |
| 528 | *Gossypium herbaceum* *(Malvaceae)* | 2020-04-13 | Huang G et al., Genome sequence of Gossypium herbaceum and genome updates of Gossypium arboreum and Gossypium hirsutum provide insights into cotton A-genome evolution. Nat Genet. 2020 May 52(5): 516-524. Epub 2020 Apr 13. |
| 527 | *Sinapis alba* *(Brassicaceae)* | 2020-04-09 | Kumari P et al., Draft genome of multiple resistance donor plant Sinapis alba: An insight into SSRs, annotations and phylogenetics. PLoS One. 2020 Apr 9, 15(4): e0231002 |
| 526 | *Thinopyrum elongatum* *(Poaceae)* | 2020-04-09 | Wang H et al., Horizontal gene transfer of Fhb7 from fungus underlies Fusarium head blight resistance in wheat. Science. 2020 May 22, 368(6493): eaba5435. Epub 2020 Apr 9. |
| 525 | *Erysimum cheiranthoides* *(Brassicaceae)* | 2020-04-07 | Zust T et al., Independent evolution of ancestral and novel defenses in a genus of toxic plants (Erysimum, Brassicaceae). Elife. 2020 Apr 7, 9: e51712 |
| 524 | *Litsea cubeba* *(Lauraceae)* | 2020-04-03 | Chen YC et al., The Litsea genome and the evolution of the laurel family. Nat Commun. 2020 Apr 3, 11(1): 1675 |
| 523 | *Asparagus setaceus* *(Asparagaceae)* | 2020-04-01 | Li SF et al., Chromosome-level genome assembly, annotation and evolutionary analysis of the ornamental plant Asparagus setaceus. Hortic Res. 2020 Apr 1 7: 48. eCollection 2020. |
| 522 | *Cladopus chinensis* *(Podostemaceae)* | 2020-04-01 | Xue T et al., A high-quality genome provides insights into the new taxonomic status and genomic characteristics of Cladopus chinensis (Podostemaceae). Hortic Res. 2020 Apr 1 7: 46. eCollection 2020. |
| 521 | *Nymphaea thermarum* *(Nymphaeaceae)* | 2020-03-31 | Povilus RA et al., Water lily (Nymphaea thermarum) genome reveals variable genomic signatures of ancient vascular cambium losses. Proc Natl Acad Sci U S A. 2020 Apr 14 117(15): 8649-8656. Epub 2020 Mar 31. |
| 520 | *Lonicera japonica* *(Caprifoliaceae)* | 2020-03-18 | Pu X et al., The honeysuckle genome provides insight into the molecular mechanism of carotenoid metabolism underlying dynamic flower coloration. New Phytol. 2020 Aug 227(3): 930-943. Epub 2020 Apr 18. |
| 519 | *Anthoceros agrestis* *(Anthocerotaceae)* | 2020-03-13 | Li FW et al., Anthoceros genomes illuminate the origin of land plants and the unique biology of hornworts. Nat Plants. 2020 Mar, 6(3): 259-272 |
| 518 | *Anthoceros punctatus* *(Anthocerotaceae)* | 2020-03-13 | Li FW et al., Anthoceros genomes illuminate the origin of land plants and the unique biology of hornworts. Nat Plants. 2020 Mar, 6(3): 259-272 |
| 517 | *Populus ilicifolia* *(Salicaceae)* | 2020-03-13 | Chen Z et al., Survival in the Tropics despite isolation, inbreeding and asexual reproduction: insights from the genome of the world's southernmost poplar (Populus ilicifolia). Plant J. 2020 Jul 103(1): 430-442. Epub 2020 Apr 3. |
| 516 | *Puccinellia tenuiflora* *(Poaceae)* | 2020-03-12 | Zhang W et al., A high-quality genome sequence of alkaligrass provides insights into halophyte stress tolerance. Sci China Life Sci. 2020 Sep 63(9): 1269-1282. Epub 2020 Mar 12. |
| 515 | *Simmondsia chinensis* *(Simmondsiaceae)* | 2020-03-11 | Sturtevant D et al., The genome of jojoba (Simmondsia chinensis): A taxonomically isolated species that directs wax ester accumulation in its seeds. Sci Adv. 2020 Mar 11, 6(11) |
| 514 | *Eriobotrya japonica* *(Rosaceae)* | 2020-03-06 | Jiang S et al., Chromosome-level genome assembly and annotation of the loquat (Eriobotrya japonica) genome. Gigascience. 2020 Mar 1, 9(3) |
| 513 | *Linum bienne* *(Linaceae)* | 2020-03-06 | Zhang J et al., Genomic Comparison and Population Diversity Analysis Provide Insights into the Domestication and Improvement of Flax. iScience. 2020 Apr 24, 23(4): 100967. Epub 2020 Mar 6. |
| 512 | *Mangifera indica* *(Anacardiaceae)* | 2020-03-06 | Wang P et al., The genome evolution and domestication of tropical fruit mango. Genome Biol. 2020 Mar 6, 21(1): 60 |
| 511 | *Dioscorea dumetorum* *(Dioscoreaceae)* | 2020-03-04 | Siadjeu C et al., High Contiguity De Novo Genome Sequence Assembly of Trifoliate Yam (Dioscorea dumetorum) Using Long Read Sequencing. Genes (Basel). 2020 Mar 4, 11(3) |
| 510 | *Oryza eichingeri* *(Poaceae)* | 2020-03-03 | Shenton M et al., Evolution and diversity of the wild rice Oryza officinalis complex, across continents genome types, and ploidy levels. Genome Biol Evol. 2020 Apr 1 12(4): 413-428. |
| 509 | *Oryza officinalis* *(Poaceae)* | 2020-03-03 | Shenton M et al., Evolution and diversity of the wild rice Oryza officinalis complex, across continents genome types, and ploidy levels. Genome Biol Evol. 2020 Apr 1 12(4): 413-428. |
| 508 | *Oryza rhizomatis* *(Poaceae)* | 2020-03-03 | Shenton M et al., Evolution and diversity of the wild rice Oryza officinalis complex, across continents genome types, and ploidy levels. Genome Biol Evol. 2020 Apr 1 12(4): 413-428. |
| 507 | *Aquilaria sinensis* *(Thymelaeaceae)* | 2020-03-02 | Ding X et al., Genome sequence of the agarwood tree Aquilaria sinensis (Lour.) Spreng: the first chromosome-level draft genome in the Thymelaeceae family. Gigascience. 2020 Mar 1, 9(3) |
| 506 | *Gossypium longicalyx* *(Malvaceae)* | 2020-03-02 | Grover CE et al., The Gossypium longicalyx Genome as a Resource for Cotton Breeding and Evolution. G3 (Bethesda). 2020 May 4, 10(5): 1457-1467 |
| 505 | *Ceratophyllum demersum* *(Ceratophyllaceae)* | 2020-02-24 | Yang Y et al., Prickly waterlily and rigid hornwort genomes shed light on early angiosperm evolution. Nat Plants. 2020 Mar 6(3): 215-222. Epub 2020 Feb 24. |
| 504 | *Euryale ferox* *(Nymphaeaceae)* | 2020-02-24 | Yang Y et al., Prickly waterlily and rigid hornwort genomes shed light on early angiosperm evolution. Nat Plants. 2020 Mar 6(3): 215-222. Epub 2020 Feb 24. |

| | plant species | published | |
|---|---|---|---|
| 503 | *Avicennia marina* *(Acanthaceae)* | 2020-02-20 | He Z et al., Convergent adaptation of the genomes of woody plants at the land-sea interface. Natl Sci Rev. 2020 Jun, 7(6): 978-993. Epub 2020 Feb 20. |
| 502 | *Sonneratia alba* *(Lythraceae)* | 2020-02-20 | He Z et al., Convergent adaptation of the genomes of woody plants at the land-sea interface. Natl Sci Rev. 2020 Jun, 7(6): 978-993. Epub 2020 Feb 20. |
| 501 | *Sonneratia caseolaris* *(Lythraceae)* | 2020-02-20 | He Z et al., Convergent adaptation of the genomes of woody plants at the land-sea interface. Natl Sci Rev. 2020 Jun, 7(6): 978-993. Epub 2020 Feb 20. |
| 500 | *Tripterygium wilfordii* *(Celastraceae)* | 2020-02-20 | Tu L et al., Genome of Tripterygium wilfordii and identification of cytochrome P450 involved in triptolide biosynthesis. Nat Commun. 2020 Feb 20, 11(1): 971 |
| 499 | *Diospyros lotus* *(Ebenaceae)* | 2020-02-18 | Akagi T et al., The persimmon genome reveals clues to the evolution of a lineage-specific sex determination system in plants. PLoS Genet. 2020 Feb 18 16(2): e1008566. eCollection 2020 Feb. |
| 498 | *Davidia involucrata* *(Nyssaceae)* | 2020-02-11 | Chen Y et al., Genomic analyses of a living fossil: The endangered dove-tree. Mol Ecol Resour. 2020 May, 20(3), Epub 2020 Feb 11 |
| 497 | *Anthoceros angustus* *(Anthocerotaceae)* | 2020-02-10 | Zhang J et al., The hornwort genome and early land plant evolution. Nat Plants. 2020 Feb 6(2): 107-118. Epub 2020 Feb 10. |
| 496 | *Populus simonii* *(Salicaceae)* | 2020-02-06 | Wu H et al., De Novo Genome Assembly of Populus simonii Further Supports That Populus simonii and Populus trichocarpa Belong to Different Sections. G3 (Bethesda). 2020 Feb 6, 10(2): 455-466. |
| 495 | *Isatis indigotica* *(Brassicaceae)* | 2020-02-01 | Kang M et al., A chromosome-scale genome assembly of Isatis indigotica, an important medicinal plant used in traditional Chinese medicine: An Isatis genome. Hortic Res. 2020 Feb 1, 7: 18 |
| 494 | *Fragaria nilgerrensis* *(Rosaceae)* | 2020-01-31 | Zhang J et al., The high-quality genome of diploid strawberry (Fragaria nilgerrensis) provides new insights into anthocyanin accumulation. Plant Biotechnol J. 2020 Sep 18(9): 1908-1924. Epub 2020 Feb 15. |
| 493 | *Ficus erecta* *(Moraceae)* | 2020-01-24 | Shirasawa K et al., The Ficus erecta genome towards Ceratocystis canker resistance breeding in common fig (F. carica). Plant J. 2020 Jun 102(6): 1313-1322. Epub 2020 Feb 24. |
| 492 | *Lupinus albus* *(Fabaceae)* | 2020-01-24 | Hufnagel B et al., High-quality genome sequence of white lupin provides insight into soil exploration and seed quality. Nat Commun. 2020 Jan 24 11(1): 492. |
| 491 | *Hibiscus cannabinus* *(Malvaceae)* | 2020-01-23 | Zhang L et al., The genome of kenaf (Hibiscus cannabinus L.) provides insights into bast fiber and leaf shape biogenesis. Plant Biotechnol J. 2020 Aug 18(8): 1796-1809. Epub 2020 Jan 30. |
| 490 | *Carpinus fangiana* *(Betulaceae)* | 2020-01-21 | Yang X et al., A Chromosome-Level Reference Genome of the Hornbeam, Carpinus Fangiana. Sci Data. 2020 Jan 21, 7(1): 24 |
| 489 | *Mikania micrantha* *(Asteraceae)* | 2020-01-17 | Liu B et al., Mikania micrantha genome provides insights into the molecular mechanism of rapid growth. Nat Commun. 2020 Jan 17, 11(1): 340 |
| 488 | *Apium graveolens* *(Apiaceae)* | 2020-01-06 | Li MY et al., The genome sequence of celery (Apium graveolens L.), an important leaf vegetable crop rich in apigenin in the Apiaceae family. Hortic Res. 2020 Jan 6 7: 9. eCollection 2020. |
| 487 | *Eucalyptus pauciflora* *(Myrtaceae)* | 2020-01-02 | Wang W et al., The draft nuclear genome assembly of Eucalyptus pauciflora: a pipeline for comparing de novo assemblies. Gigascience. 2020 Jan 1 9(1): giz160. |
| 486 | *Castanea dentata* *(Fagaceae)* | 2019-12-29 | Westbrook JW et al., Optimizing genomic selection for blight resistance in American chestnut backcross populations: A trade-off with American chestnut ancestry implies resistance is polygenic. Evol Appl. 2019 Dec 29, 13(1): 31-47. eCollection 2020 Jan. |
| 485 | *Artocarpus altilis* *(Moraceae)* | 2019-12-24 | Sahu SK et al., Draft Genomes of Two Artocarpus Plants, Jackfruit (A. heterophyllus) and Breadfruit (A. altilis). Genes (Basel). 2019 Dec 24, 11(1) |
| 484 | *Artocarpus heterophyllus* *(Moraceae)* | 2019-12-24 | Sahu SK et al., Draft Genomes of Two Artocarpus Plants, Jackfruit (A. heterophyllus) and Breadfruit (A. altilis). Genes (Basel). 2019 Dec 24, 11(1) |
| 483 | *Luffa cylindrica* *(Cucurbitaceae)* | 2019-12-23 | Zhang T et al., Long-read sequencing and de novo assembly of the Luffa cylindrica (L.) Roem. Genome. Mol Ecol Resour. 2020 Mar 20(2): 511-519. Epub 2020 Jan 16. |
| 482 | *Diospyros oleifera* *(Ebenaceae)* | 2019-12-18 | Zhu QG et al., The persimmon (Diospyros oleifera Cheng) genome provides new insights into the inheritance of astringency and ancestral evolution. Hortic Res. 2019 Dec 18, 6: 138 |
| 481 | *Nymphaea colorata* *(Nymphaeaceae)* | 2019-12-18 | Zhang L et al., The water lily genome and the early evolution of flowering plants. Nature. 2020 Jan 577(7788): 79-84. Epub 2019 Dec 18. |
| 480 | *Utricularia reniformis* *(Lentibulariaceae)* | 2019-12-18 | Silva SR et al., The Terrestrial Carnivorous Plant Utricularia reniformis Sheds Light on Environmental and Life-Form Genome Plasticity. Int J Mol Sci. 2019 Dec 18, 21(1) |
| 479 | *Chlorokybus atmophyticus* *(Chlorokybaceae)* | 2019-12-16 | Wang S et al., Genomes of early-diverging streptophyte algae shed light on plant terrestrialization. Nat Plants. 2020 Feb 6(2): 95-106. Epub 2019 Dec 16. |
| 478 | *Fragaria iinumae* *(Rosaceae)* | 2019-12-16 | Edger PP et al., Reply to: Revisiting the origin of octoploid strawberry. Nat Genet. 2020 Jan, 52(1): 5-7. |
| 477 | *Mesostigma viride* *(Mesostigmataceae)* | 2019-12-16 | Wang S et al., Genomes of early-diverging streptophyte algae shed light on plant terrestrialization. Nat Plants. 2020 Feb 6(2): 95-106. Epub 2019 Dec 16. |
| 476 | *Coriandrum sativum* *(Apiaceae)* | 2019-12-04 | Song X et al., Deciphering the High Quality Genome Sequence of Coriander that Causes Controversial Feelings. Plant Biotechnol J. 2020 Jun 18(6): 1444-1456. Epub 2020 Feb 5. |
| 475 | *Ceratopteris richardii* *(Pteridaceae)* | 2019-12-03 | Marchant DB et al., The C-Fern (Ceratopteris richardii) genome: insights into plant genome evolution with the first partial homosporous fern genome assembly. Sci Rep. 2019 Dec 3, 9(1): 18181 |
| 474 | *Tanacetum cinerariifolium* *(Asteraceae)* | 2019-12-03 | Yamashiro T et al., Draft genome of Tanacetum cinerariifolium, the natural source of mosquito coil. Sci Rep. 2019 Dec 3, 9(1): 18249 |
| 473 | *Gossypioides kirkii* *(Malvaceae)* | 2019-11-27 | Udall JA et al., The Genome Sequence of Gossypioides kirkii Illustrates a Descending Dysploidy in Plants. Front Plant Sci. 2019 Nov 27 10: 1541. eCollection 2019. |
| 472 | *Abrus precatorius* *(Fabaceae)* | 2019-11-25 | Hovde BT et al., Detection of Abrin-Like and Prepropulchellin-Like Toxin Genes and Transcripts Using Whole Genome Sequencing and Full-Length Transcript Sequencing of Abrus precatorius. Toxins (Basel). 2019 Nov 25, 11(12) |
| 471 | *Coix aquatica* *(Poaceae)* | 2019-11-25 | Guo C et al., The Coix genome provides insights into Panicoideae evolution and papery hull domestication. Mol Plant. 2020 Feb 3 13(2): 309-320. Epub 2019 Nov 25. |
| 470 | *Leptospermum scoparium* *(Myrtaceae)* | 2019-11-25 | Thrimawithana AH et al., A whole genome assembly of Leptospermum scoparium (Myrtaceae) for manuka research. New Zealand J Crop Hort Sci. 2019 Nov 25, Epub ahead of print |
| 469 | *Nyssa sinensis* *(Nyssaceae)* | 2019-11-25 | Yang X et al., A chromosome-level genome assembly of the Chinese tupelo Nyssa sinensis. Sci Data. 2019 Nov 25, 6(1): 282 |
| 468 | *Pyrus ussuriensis x communis* *(Rosaceae)* | 2019-11-25 | Ou C et al., A de novo genome assembly of the dwarfing pear rootstock Zhongai 1. Sci Data. 2019 Nov 25, 6(1): 281 |
| 467 | *Avena atlantica* *(Poaceae)* | 2019-11-22 | Maughan PJ et al., Genomic insights from the first chromosome-scale assemblies of oat (Avena spp.) diploid species. BMC Biol. 2019 Nov 22, 17(1): 92. |

| | plant species | published | |
|---|---|---|---|
| 466 | *Avena eriantha* *(Poaceae)* | 2019-11-22 | Maughan PJ et al., Genomic insights from the first chromosome-scale assemblies of oat (Avena spp.) diploid species. BMC Biol. 2019 Nov 22, 17(1): 92. |
| 465 | *Salix brachista* *(Salicaceae)* | 2019-11-19 | Chen JH et al., Genome-wide analysis of Cushion willow provides insights into alpine plant divergence in a biodiversity hotspot. Nat Commun. 2019 Nov 19, 10(1): 5230 |
| 464 | *Trochodendron aralioides* *(Trochodendraceae)* | 2019-11-18 | Strijk JS et al., Trochodendron aralioides, the first chromosome-level draft genome in Trochodendrales and a valuable resource for basal eudicot research. Gigascience. 2019 Nov 1, 8(11) |
| 463 | *Benincasa hispida* *(Cucurbitaceae)* | 2019-11-14 | Xie D et al., The wax gourd genomes offer insights into the genetic diversity and ancestral cucurbit karyotype. Nat Commun. 2019 Nov 14, 10(1): 5158 |
| 462 | *Mesotaenium endlicherianum* *(Mesotaeniaceae)* | 2019-11-14 | Cheng S et al., Genomes of Subaerial Zygnematophyceae Provide Insights into Land Plant Evolution. Cell. 2019 Nov 14, 179(5): 1057-1067.e14 |
| 461 | *Spirogloea muscicola* *(Spirogloeaceae)* | 2019-11-14 | Cheng S et al., Genomes of Subaerial Zygnematophyceae Provide Insights into Land Plant Evolution. Cell. 2019 Nov 14, 179(5): 1057-1067.e14 |
| 460 | *Pachycladon cheesemanii* *(Brassicaceae)* | 2019-11-12 | Dong Y et al., Genome draft of the Arabidopsis relative Pachycladon cheesemanii reveals novel strategies to tolerate New Zealand's high ultraviolet B radiation environment BMC Genomics. 2019 Nov 12, 20: 838 |
| 459 | *Chenopodium pallidicaule* *(Amaranthaceae)* | 2019-11-08 | Mangelson H et al., The genome of Chenopodium pallidicaule: An emerging Andean super grain. Appl Plant Sci. 2019 Nov 8 7(11): e11300. eCollection 2019 Nov. |
| 458 | *Rhododendron williamsianum* *(Ericaceae)* | 2019-11-08 | Soza VL et al., The Rhododendron genome and chromosomal organization provide insight into shared whole genome duplications across the heath family (Ericaceae). Genome Biol Evol. 2019 Dec 1 11(12): 3353-3371. |
| 457 | *Tetraselmis striata* *(Chlorodendraceae)* | 2019-10-24 | Steadman-Tyler CR et al., High-Quality Draft Genome Sequence of the Green Alga Tetraselmis striata (Chlorophyta) Generated from PacBio Sequencing. Microbiol Resour Announc. 2019 Oct 24, 8(43) |
| 456 | *Picochlorum renovo* *(C.incertae sedis)* | 2019-10-23 | Dahlin LR et al., Development of a high-productivity, halophilic, thermotolerant microalga Picochlorum renovo. Commun Biol. 2019, 2(): 388. |
| 455 | *Echium plantagineum* *(Boraginaceae)* | 2019-10-18 | Tang CY et al., Comparative genome/transcriptome analysis probes Boraginales phylogenetic position, WGDs in Boraginales, and key enzyme genes in the alkannin/shikonin core pathway. Mol Ecol Resour. 2020 Jan, 20(1): 228-241. |
| 454 | *Messastrum gracile* *(Selenastraceae)* | 2019-10-18 | Teh KY et al., De novo whole genome sequencing data of two mangrove-isolated microalgae from Terengganu coastal waters. Data Brief. 2019 Dec, 27: 104680 |
| 453 | *Polygonum cuspidatum* *(Polygonaceae)* | 2019-10-18 | Zhang Y et al., Assembly and Annotation of a Draft Genome of the Medicinal Plant Polygonum cuspidatum. Front Plant Sci. 2019 Oct 18 10: 1274. eCollection 2019. |
| 452 | *Piper nigrum* *(Piperaceae)* | 2019-10-16 | Hu L et al., The chromosome-scale reference genome of black pepper provides insight into piperine biosynthesis Nat Commun. 2019 Oct 16, 10(1): 4702 |
| 451 | *Solanum chilense* *(Solanaceae)* | 2019-10-11 | Stam R et al., The de Novo Reference Genome and Transcriptome Assemblies of the Wild Tomato Species Solanum chilense Highlights Birth and Death of NLR Genes Between Tomato Species. G3 (Bethesda). 2019 Dec 3 9(12): 3933-3941. |
| 450 | *Solanum aethiopicum* *(Solanaceae)* | 2019-10-01 | Song B et al., Draft genome sequence of Solanum aethiopicum provides insights into disease resistance, drought tolerance, and the evolution of the genome. Gigascience. 2019 Oct 1, 8(10) |
| 449 | *Amaranthus tuberculatus* *(Amaranthaceae)* | 2019-09-30 | Kreiner JM et al., Multiple modes of convergent adaptation in the spread of glyphosate-resistant Amaranthus tuberculatus. Proc Natl Acad Sci U S A. 2019 Oct 15, 116(42): 21076-21084 |
| 448 | *Asclepias syriaca* *(Apocynaceae)* | 2019-09-20 | Weitemier K et al., A draft genome and transcriptome of common milkweed (Asclepias syriaca) as resources for evolutionary, ecological, and molecular studies in milkweeds and Apocynaceae. PeerJ 7: e7649 |
| 447 | *Bassia scoparia* *(Amaranthaceae)* | 2019-09-13 | Patterson EL et al., The Draft Genome of Kochia scoparia and the Mechanism of Glyphosate Resistance via Transposon-Mediated EPSPS Tandem Gene Duplication. Genome Biol Evol. 2019 Oct 1 11(10): 2927-2940. |
| 446 | *Gossypium australe* *(Malvaceae)* | 2019-09-13 | Cai Y et al., Genome sequencing of the Australian wild diploid species Gossypium australe highlights disease resistance and delayed gland morphogenesis. Plant Biotechnol J. 2020 Mar, 18(3): 814-828. Epub 2019 Sep 13. |
| 445 | *Castanea mollissima* *(Fagaceae)* | 2019-09-12 | Xing Y et al., Hybrid de novo genome assembly of Chinese chestnut (Castanea mollissima). Gigascience. 2019 Sep 1, 8(9) |
| 444 | *Suaeda aralocaspica* *(Amaranthaceae)* | 2019-09-12 | Wang L et al., A draft genome assembly of halophyte Suaeda aralocaspica, a plant that performs C4 photosynthesis within individual cells. Gigascience. 2019 Sep 1, 8(9) |
| 443 | *Arabis nemorensis* *(Brassicaceae)* | 2019-09-11 | Dittberner H et al., Strengths and potential pitfalls of hay transfer for ecological restoration revealed by RAD-seq analysis in floodplain Arabis species. Mol Ecol. 2019 Sep, 28(17): 3887-3901. |
| 442 | *Striga asiatica* *(Orobanchaceae)* | 2019-09-10 | Yoshida S et al., Genome Sequence of Striga asiatica Provides Insight into the Evolution of Plant Parasitism. Curr Biol. 2019 Sep 23 29(18):3041-3052.e4. Epub 2019 Sep 12. |
| 441 | *Chloropicon primus* *(Chloropicaceae)* | 2019-09-06 | Lemieux C et al., A streamlined and predominantly diploid genome in the tiny marine green alga Chloropicon primus. Nat Commun. 2019 Sep 6, 10(1): 4061 |
| 440 | *Pisum sativum* *(Fabaceae)* | 2019-09-02 | Kreplak J et al., A reference genome for pea provides insight into legume genome evolution. Nat Genet. 2019 Sep, 51(9):1411-1422 Epub 2019 Sep 2. |
| 439 | *Gossypium turneri* *(Malvaceae)* | 2019-08-28 | Udall JA et al., De novo genome sequence assemblies of Gossypium raimondii and G. turneri. G3 (Bethesda). 2019 Oct 7 9(10):3079-3085. |
| 438 | *Gelsemium elegans* *(Gelsemiaceae)* | 2019-08-16 | Liu Y et al., Whole-genome sequencing and analysis of the Chinese herbal plant Gelsemium elegans. Acta Pharm Sin B. 2020 Feb, 10(2): 374-382 |
| 437 | *Phytolacca americana* *(Phytolaccaceae)* | 2019-08-08 | Neller KCM et al., De novo Assembly of the Pokeweed Genome Provides Insight Into Pokeweed Antiviral Protein (PAP) Gene Expression. Front Plant Sci. 2019 Aug 6, 10: 1002. eCollection 2019. |
| 436 | *Persea americana* *(Lauraceae)* | 2019-08-06 | Rendon-Anaya M et al., The avocado genome informs deep angiosperm phylogeny, highlights introgressive hybridization, and reveals pathogen-influenced gene space adaptation. Proc Natl Acad Sci U S A. 2019 Aug 20 116(34):17081-17089. Epub 2019 Aug 6. |
| 435 | *Pyrus betulifolia* *(Rosaceae)* | 2019-08-01 | Dong X et al., De novo assembly of a wild pear (Pyrus betuleafolia) genome. Plant Biotechnol J. 2020 Feb 18(2): 581-595. Epub 2019 Aug 12. |
| 434 | *Asterochloris glomerata* *(Trebouxiaceae)* | 2019-07-23 | Armaleo D et al., The lichen symbiosis re-viewed through the genomes of Cladonia grayi and its algal partner Asterochloris glomerata. BMC Genomics. 2019 Jul 23, 20(1): 605 |
| 433 | *Vitis riparia* *(Vitaceae)* | 2019-07-19 | Girollet N et al., De Novo Phased Assembly of the Vitis Riparia Grape Genome. Sci Data. 2019 Jul 19, 6(1): 127 |
| 432 | *Vigna marina* *(Fabaceae)* | 2019-07-18 | Singh AK et al., Draft genome sequence of a less-known wild Vigna: Beach pea (V. marina cv. ANBp-14-03) Crop Journal (2019), epub ahead of print |
| 431 | *Rhodelphis limnetica* *(Rhodelphidaceae)* | 2019-07-17 | Gawryluk RMR et al., Non-photosynthetic predators are sister to red algae. Nature. 2019 Aug, 572(7768): 240-243. Epub 2019 Jul 17. |
| 430 | *Acer yangbiense* *(Sapindaceae)* | 2019-07-15 | Yang J et al., De novo genome assembly of the endangered Acer yangbiense, a plant species with extremely small populations endemic to Yunnan Province, China. Gigascience. 2019 Jul 1, 8(7) |

| | plant species | published | |
|---|---|---|---|
| 429 | *Eragrostis curvula* (*Poaceae*) | 2019-07-15 | Carballo J et al., A high-quality genome of Eragrostis curvula grass provides insights into Poaceae evolution and supports new strategies to enhance forage quality. Sci Rep. 2019 Jul 15, 9(1): 10250 |
| 428 | *Pleurozium schreberi* (*Hylocomiaceae*) | 2019-07-08 | Pederson ERA et al., Genome Sequencing of Pleurozium schreberi: The Assembled and Annotated Draft Genome of a Pleurocarpous Feather Moss. G3 (Bethesda). 2019 Sep 4, 9(9): 2791-2797 |
| 427 | *Dactylis glomerata* (*Poaceae*) | 2019-07-05 | Huang L et al., Genome assembly provides insights into the genome evolution and flowering regulation of orchardgrass. Plant Biotechnol J. 2020 Feb 18(2): 373-388. Epub 2019 Jul 30. |
| 426 | *Spatholobus suberectus* (*Fabaceae*) | 2019-07-04 | Qin S et al., A draft genome for Spatholobus suberectus. Sci Data. 2019 Jul 4, 6(1): 113 |
| 425 | *Xanthoceras sorbifolium* (*Sapindaceae*) | 2019-06-26 | Liang Q et al., The genome assembly and annotation of yellowhorn (Xanthoceras sorbifolium Bunge). Gigascience. 2019 Jun, 8(6): giz071 |
| 424 | *Abies alba* (*Pinaceae*) | 2019-06-20 | Mosca E et al., A Reference Genome Sequence for the European Silver Fir (Abies alba Mill.): A Community-Generated Genomic Resource. G3 (Bethesda). 2019 Jul 9 9(7): 2039-2049. |
| 423 | *Marchantia inflexa* (*Marchantiaceae*) | 2019-06-19 | Marks RA et al., Genome of the tropical plant Marchantia inflexa: implications for sex chromosome evolution and dehydration tolerance. Sci Rep. 2019 Jun 19, 9(1): 8722 |
| 422 | *Prunus dulcis* (*Rosaceae*) | 2019-06-14 | Sanchez-Perez R et al., Mutation of a bHLH transcription factor allowed almond domestication. Science. 2019 Jun 14, 364(6445): 1095-1098. |
| 421 | *Sedum album* (*Crassulaceae*) | 2019-06-14 | Wai CM et al., Time of day and network reprogramming during drought induced CAM photosynthesis in Sedum album. PLoS Genet. 2019 Jun 14 15(6):e1008209. eCollection 2019 Jun. |
| 420 | *Trichopus zeylanicus* (*Dioscoreaceae*) | 2019-06-12 | Vadakkemukadiyil-Chellappan B et al., High Quality Draft Genome of Arogyapacha (Trichopus zeylanicus), an Important Medicinal Plant Endemic to Western Ghats of India. G3 (Bethesda). 2019 Aug 8 9(8):2395-2404. |
| 419 | *Bonia amplexicaulis* (*Poaceae*) | 2019-05-27 | Guo ZH et al., Genome sequences provide insights into the reticulate origin and unique traits of woody bamboos. Mol Plant. 2019 Oct 7 12(10):1353-1365. Epub 2019 May 27. |
| 418 | *Guadua angustifolia* (*Poaceae*) | 2019-05-27 | Guo ZH et al., Genome sequences provide insights into the reticulate origin and unique traits of woody bamboos. Mol Plant. 2019 Oct 7 12(10):1353-1365. Epub 2019 May 27. |
| 417 | *Olyra latifolia* (*Poaceae*) | 2019-05-27 | Guo ZH et al., Genome sequences provide insights into the reticulate origin and unique traits of woody bamboos. Mol Plant. 2019 Oct 7 12(10):1353-1365. Epub 2019 May 27. |
| 416 | *Raddia guianensis* (*Poaceae*) | 2019-05-27 | Guo ZH et al., Genome sequences provide insights into the reticulate origin and unique traits of woody bamboos. Mol Plant. 2019 Oct 7 12(10):1353-1365. Epub 2019 May 27. |
| 415 | *Malus baccata* (*Rosaceae*) | 2019-05-24 | Chen X et al., Sequencing of a Wild Apple (Malus baccata) Genome Unravels the Differences Between Cultivated and Wild Apple Species Regarding Disease Resistance and Cold Tolerance. G3 (Bethesda). 2019 Jul 9 9(7):2051-2060. |
| 414 | *Mercurialis annua* (*Euphorbiaceae*) | 2019-05-24 | Veltsos P et al., Early Sex-chromosome Evolution in the Diploid Dioecious Plant Mercurialis annua. Genetics. 2019 Jul 212(3):815-835. Epub 2019 May 21. |
| 413 | *Citrus japonica* (*Rutaceae*) | 2019-05-21 | Zhu C et al., Genome sequencing and CRISPR/Cas9 gene editing of an early flowering Mini-Citrus (Fortunella hindsii). Plant Biotechnol J. 2019 Nov, 17(11): 2199-2210. Epub 2019 May 21. |
| 412 | *Beta patula* (*Amaranthaceae*) | 2019-05-18 | Del Rio AR et al., Genomes of the wild beets Beta patula and Beta vulgaris ssp. maritima. Plant J. 2019 Sep 99(6):1242-1253. Epub 2019 Jul 15. |
| 411 | *Carya cathayensis* (*Juglandaceae*) | 2019-05-02 | Huang Y et al., The genomes of pecan and Chinese hickory provide insights into Carya evolution and nut nutrition. GigaScience 2019 May 02 8(5): giz036 |
| 410 | *Carya illinoinensis* (*Juglandaceae*) | 2019-05-02 | Huang Y et al., The genomes of pecan and Chinese hickory provide insights into Carya evolution and nut nutrition. GigaScience 2019 May 02 8(5): giz036 |
| 409 | *Trifolium occidentale* (*Fabaceae*) | 2019-04-25 | Griffiths AG et al., Breaking Free: The Genomics of Allopolyploidy-Facilitated Niche Expansion in White Clover. Plant Cell. 2019 Jul, 31(7): 1466-1487. Epub 2019 Apr 25. |
| 408 | *Trifolium pallescens* (*Fabaceae*) | 2019-04-25 | Griffiths AG et al., Breaking Free: The Genomics of Allopolyploidy-Facilitated Niche Expansion in White Clover. Plant Cell. 2019 Jul, 31(7): 1466-1487. Epub 2019 Apr 25. |
| 407 | *Trifolium repens* (*Fabaceae*) | 2019-04-25 | Griffiths AG et al., Breaking Free: The Genomics of Allopolyploidy-Facilitated Niche Expansion in White Clover. Plant Cell. 2019 Jul, 31(7): 1466-1487. Epub 2019 Apr 25. |
| 406 | *Vigna unguiculata* (*Fabaceae*) | 2019-04-24 | Lonardi S et al., The genome of cowpea (Vigna unguiculata (L.) Walp.). Plant J. 2019 Jun 98(5):767-782. |
| 405 | *Pistacia vera* (*Anacardiaceae*) | 2019-04-18 | Zeng L et al., Whole genomes and transcriptomes reveal adaptation and domestication of pistachio. Genome Biol. 2019 Apr 18, 20(1):79 |
| 404 | *Scutellaria baicalensis* (*Lamiaceae*) | 2019-04-15 | Zhao Q et al., The Reference Genome Sequence of Scutellaria baicalensis Provides Insights into the Evolution of Wogonin Biosynthesis. Mol Plant. 2019 Jul 1 12(7):935-950. Epub 2019 Apr 15. |
| 403 | *Actinidia eriantha* (*Actinidiaceae*) | 2019-04-03 | Tang W et al., Chromosome-scale genome assembly of kiwifruit Actinidia eriantha with single-molecule sequencing and chromatin interaction mapping. Gigascience. 2019 Apr 1,8(4): giz027 |
| 402 | *Caulerpa lentillifera* (*Caulerpaceae*) | 2019-03-28 | Arimoto A et al., A siphonous macroalgal genome suggests convergent functions of homeobox genes in algae and land plants. DNA Res. 2019 Apr 1 26(2): 183-192. |
| 401 | *Crucihimalaya himalaica* (*Brassicaceae*) | 2019-03-20 | Zhang T et al., Genome of Crucihimalaya himalaica, a close relative of Arabidopsis, shows ecological adaptation to high altitude. Proc Natl Acad Sci U S A. 2019 Apr 2 116(14):7137-7146. Epub 2019 Mar 20. |
| 400 | *Arachis hypogaea* (*Fabaceae*) | 2019-03-19 | Chen X et al., Sequencing of Cultivated Peanut, Arachis hypogaea, Yields Insights into Genome Evolution and Oil Improvement. Mol Plant. 2019 Jul 1 12(7):920-934. Epub 2019 Mar 19. |
| 399 | *Eleusine indica* (*Poaceae*) | 2019-03-08 | Zhang H et al., Development of a goosegrass (Eleusine indica) draft genome and application to weed science research. Pest Manag Sci. 2019 Oct 75(10):2776-2784. Epub 2019 Apr 10. |
| 398 | *Fragaria x ananassa* (*Rosaceae*) | 2019-02-25 | Edger PP et al., Origin and evolution of the octoploid strawberry genome. Nat Genet. 2019 Mar 51(3):541-547. Epub 2019 Feb 25. |
| 397 | *Alloteropsis semialata* (*Poaceae*) | 2019-02-20 | Dunning LT et al., Lateral transfers of large DNA fragments spread functional genes among grasses. Proc Natl Acad Sci U S A. 2019 Mar 5, 116(10): 4416-4425. Epub 2019 Feb 20. |
| 396 | *Broussonetia papyrifera* (*Moraceae*) | 2019-01-31 | Peng X et al., A Chromosome-Scale Genome Assembly of Paper Mulberry (Broussonetia papyrifera) Provides New Insights into Its Forage and Papermaking Usage. Mol Plant. 2019 May 6 12(5):661-677. Epub 2019 Feb 26. |
| 395 | *Antirrhinum majus* (*Plantaginaceae*) | 2019-01-28 | Li M et al., Genome structure and evolution of Antirrhinum majus L. Nat Plants. 2019 Feb 5(2):174-183. Epub 2019 Jan 28. |
| 394 | *Chrysanthemum seticuspe* (*Asteraceae*) | 2019-01-27 | Hirakawa H et al., De novo whole-genome assembly in Chrysanthemum seticuspe, a model species of Chrysanthemums, and its application to genetic and gene discovery analysis DNA Res. 2019 Jun 1, 26(3): 195-203 |
| 393 | *Panicum miliaceum* (*Poaceae*) | 2019-01-25 | Zou C et al., The genome of broomcorn millet. Nat Commun. 2019 Jan 25, 10(1): 436 |

| | plant species | published | |
|---|---|---|---|
| 392 | *Malania oleifera* (*Olacaceae*) | 2019-01-24 | Xu CQ et al., Genome sequence of Malania oleifera, a tree with great value for nervonic acid production.<br>GigaScience 2019 Jan 24, giy164 |
| 391 | *Picocystis sp.* (*Picocystaceae*) | 2019-01-24 | Junkins EN et al., Draft Genome Sequence of Picocystis sp. Strain ML, Cultivated from Mono Lake, California.<br>Microbiol Resour Announc. 2019 Jan, 8(4)): e01353-18. |
| 390 | *Cinnamomum kanehirae* (*Lauraceae*) | 2019-01-09 | Chaw SM et al., Stout camphor tree genome fills gaps in understanding of flowering plant genome evolution.<br>Nat Plants. 2019 Jan, 5(1): 63-73 |
| 389 | *Cucurbita argyrosperma* (*Cucurbitaceae*) | 2019-01-07 | Barrera-Redondo J et al., The genome of Cucurbita argyrosperma (silver-seed gourd) reveals faster rates of protein-coding gene and long noncoding RNA turnover and neofunctionalization within Cucurbita.<br>Mol Plant. 2019 Apr 1 12(4):506-520. Epub 2019 Jan 8. |
| 388 | *Jaltomata sinuosa* (*Solanaceae*) | 2019-01-04 | Wu M et al., Genome sequence of Jaltomata addresses rapid reproductive trait evolution and enhances comparative genomics in the hyper-diverse Solanaceae.<br>Genome Biol Evol. 2019 Feb 1 11(2):335-349. |
| 387 | *Ostrya chinensis* (*Betulaceae*) | 2018-12-21 | Yang Y et al., Genomic effects of population collapse in a critically endangered ironwood tree Ostrya rehderiana.<br>Nat Commun. 2018 Dec 21, 9(1): 5449 |
| 386 | *Ostrya rehderiana* (*Betulaceae*) | 2018-12-21 | Yang Y et al., Genomic effects of population collapse in a critically endangered ironwood tree Ostrya rehderiana.<br>Nat Commun. 2018 Dec 21, 9(1): 5449 |
| 385 | *Primula vulgaris* (*Primulaceae*) | 2018-12-18 | Cocker JM et al., Primula vulgaris (primrose) genome assembly, annotation and gene expression, with comparative genomics on the heterostyly supergene.<br>Sci Rep. 2018 Dec 18, 8(1): 17942 |
| 384 | *Liriodendron chinense* (*Magnoliaceae*) | 2018-12-17 | Chen J et al., Liriodendron genome sheds light on angiosperm phylogeny and species-pair differentiation.<br>Nat Plants. 2019 Jan 5(1):18-25. Epub 2018 Dec 17. |
| 383 | *Dracaena cambodiana* (*Asparagaceae*) | 2018-12-14 | Ding X et al., Genome survey sequencing for the characterization of genetic background of Dracaena cambodiana and its defense response during dragon's blood formation.<br>PLoS One. 2018 Dec 14, 13(12): e0209258. |
| 382 | *Faidherbia albida* (*Fabaceae*) | 2018-12-07 | Chang Y et al., The draft genomes of five agriculturally important African orphan crops.<br>GigaScience 2018, giy152 |
| 381 | *Lablab purpureus* (*Fabaceae*) | 2018-12-07 | Chang Y et al., The draft genomes of five agriculturally important African orphan crops.<br>GigaScience 2018, giy152 |
| 380 | *Sclerocarya birrea* (*Anacardiaceae*) | 2018-12-07 | Chang Y et al., The draft genomes of five agriculturally important African orphan crops.<br>GigaScience 2018, giy152 |
| 379 | *Vigna subterranea* (*Fabaceae*) | 2018-12-07 | Chang Y et al., The draft genomes of five agriculturally important African orphan crops.<br>GigaScience 2018, giy152 |
| 378 | *Panicum hallii* (*Poaceae*) | 2018-12-06 | Lovell JT et al., The genomic landscape of molecular responses to natural drought stress in Panicum hallii.<br>Nat Commun. 2018 Dec 6, 9(1): 5213 |
| 377 | *Haematococcus pluvialis* (*Haematococcaceae*) | 2018-11-29 | Luo Q et al., Genome and Transcriptome Sequencing of the Astaxanthin-Producing Green Microalga, Haematococcus pluvialis.<br>Genome Biol Evol. 2019 Jan 1 11(1): 166-173. |
| 376 | *Nicotiana knightiana* (*Solanaceae*) | 2018-11-29 | Sierro N et al., The impact of genome evolution on the allotetraploid Nicotiana rustica - an intriguing story of enhanced alkaloid production.<br>BMC Genomics 2018 Nov 29, 19(1): 855 |
| 375 | *Nicotiana paniculata* (*Solanaceae*) | 2018-11-29 | Sierro N et al., The impact of genome evolution on the allotetraploid Nicotiana rustica - an intriguing story of enhanced alkaloid production.<br>BMC Genomics 2018 Nov 29, 19(1): 855 |
| 374 | *Nicotiana rustica* (*Solanaceae*) | 2018-11-29 | Sierro N et al., The impact of genome evolution on the allotetraploid Nicotiana rustica - an intriguing story of enhanced alkaloid production.<br>BMC Genomics 2018 Nov 29, 19(1): 855 |
| 373 | *Nicotiana undulata* (*Solanaceae*) | 2018-11-29 | Sierro N et al., The impact of genome evolution on the allotetraploid Nicotiana rustica - an intriguing story of enhanced alkaloid production.<br>BMC Genomics 2018 Nov 29, 19(1): 855 |
| 372 | *Gossypium aridum* (*Malvaceae*) | 2018-11-23 | Grover CE et al., Insights into the Evolution of the New World Diploid Cottons (Gossypium, Subgenus Houzingenia) Based on Genome Sequencing.<br>Genome Biol Evol. 2019 Jan 1, 11(1): 53-71. |
| 371 | *Gossypium gossypioides* (*Malvaceae*) | 2018-11-23 | Grover CE et al., Insights into the Evolution of the New World Diploid Cottons (Gossypium, Subgenus Houzingenia) Based on Genome Sequencing.<br>Genome Biol Evol. 2019 Jan 1, 11(1): 53-71. |
| 370 | *Gossypium harknessii* (*Malvaceae*) | 2018-11-23 | Grover CE et al., Insights into the Evolution of the New World Diploid Cottons (Gossypium, Subgenus Houzingenia) Based on Genome Sequencing.<br>Genome Biol Evol. 2019 Jan 1, 11(1): 53-71. |
| 369 | *Gossypium klotzschianum* (*Malvaceae*) | 2018-11-23 | Grover CE et al., Insights into the Evolution of the New World Diploid Cottons (Gossypium, Subgenus Houzingenia) Based on Genome Sequencing.<br>Genome Biol Evol. 2019 Jan 1, 11(1): 53-71. |
| 368 | *Gossypium laxum* (*Malvaceae*) | 2018-11-23 | Grover CE et al., Insights into the Evolution of the New World Diploid Cottons (Gossypium, Subgenus Houzingenia) Based on Genome Sequencing.<br>Genome Biol Evol. 2019 Jan 1, 11(1): 53-71. |
| 367 | *Gossypium lobatum* (*Malvaceae*) | 2018-11-23 | Grover CE et al., Insights into the Evolution of the New World Diploid Cottons (Gossypium, Subgenus Houzingenia) Based on Genome Sequencing.<br>Genome Biol Evol. 2019 Jan 1, 11(1): 53-71. |
| 366 | *Gossypium schwendimanii* (*Malvaceae*) | 2018-11-23 | Grover CE et al., Insights into the Evolution of the New World Diploid Cottons (Gossypium, Subgenus Houzingenia) Based on Genome Sequencing.<br>Genome Biol Evol. 2019 Jan 1, 11(1): 53-71. |
| 365 | *Gossypium thurberi* (*Malvaceae*) | 2018-11-23 | Grover CE et al., Insights into the Evolution of the New World Diploid Cottons (Gossypium, Subgenus Houzingenia) Based on Genome Sequencing.<br>Genome Biol Evol. 2019 Jan 1, 11(1): 53-71. |
| 364 | *Gossypium trilobum* (*Malvaceae*) | 2018-11-23 | Grover CE et al., Insights into the Evolution of the New World Diploid Cottons (Gossypium, Subgenus Houzingenia) Based on Genome Sequencing.<br>Genome Biol Evol. 2019 Jan 1, 11(1): 53-71. |
| 363 | *Osmanthus fragrans* (*Oleaceae*) | 2018-11-20 | Yang X et al., The chromosome-level quality genome provides insights into the evolution of the biosynthesis genes for aroma compounds of Osmanthus fragrans.<br>Hortic Res. 2018 Nov 20, 5: 72 |
| 362 | *Andrographis paniculata* (*Acanthaceae*) | 2018-11-16 | Sun W et al., The medicinal plant Andrographis paniculata genome provides insight into biosynthesis of the bioactive diterpenoid neoandrographolide.<br>Plant J. 2019 Mar 97(5):841-857. Epub 2019 Jan 2. |
| 361 | *Casuarina equisetifolia* (*Casuarinaceae*) | 2018-11-14 | Ye G et al., De novo genome assembly of the stress tolerant forest species Casuarina equisetifolia provides insight into secondary growth.<br>Plant J. 2019 Feb 97(4):779-794. Epub 2019 Jan 22. |
| 360 | *Scenedesmus quadricauda* (*Scenedesmaceae* ) | 2018-11-09 | Nag Dasgupta C et al., Draft genome sequence and detailed characterization of biofuel production by oleaginous microalga Scenedesmus quadricauda LWG002611.<br>Biotechnol Biofuels. 2018 Nov 9, 11: 308 |
| 359 | *Ipomoea triloba* (*Convolvulaceae*) | 2018-11-02 | Wu S et al., Genome sequences of two diploid wild relatives of cultivated sweetpotato reveal targets for genetic improvement.<br>Nat Commun. 2018 Nov 2, 9(1): 4580 |
| 358 | *Musa schizocarpa* (*Musaceae*) | 2018-11-02 | Belser C et al., Chromosome-scale assemblies of plant genomes using nanopore long reads and optical maps.<br>Nat Plants. 2018 Nov, 4(11):879-887 |
| 357 | *Populus tremula* (*Salicaceae*) | 2018-10-29 | Lin YC et al., Functional and evolutionary genomic inferences in Populus through genome and population sequencing of American and European aspen<br>Proc Natl Acad Sci U S A. 2018 Nov 13 115(46):E10970-E10978. Epub 2018 Oct 29. |

| | plant species | published | |
|---|---|---|---|
| 356 | *Populus tremuloides* *(Salicaceae)* | 2018-10-29 | Lin YC et al., Functional and evolutionary genomic inferences in Populus through genome and population sequencing of American and European aspen Proc Natl Acad Sci U S A. 2018 Nov 13 115(46):E10970-E10978. Epub 2018 Oct 29. |
| 355 | *Linderniella brevidens* *(Linderniaceae)* | 2018-10-25 | VanBuren R et al., Desiccation tolerance evolved through gene duplication and network rewiring in Lindernia. Plant Cell. 2018 Dec 30(12):2943-2958. Epub 2018 Oct 25. |
| 354 | *Vandellia subracemosa* *(Linderniaceae)* | 2018-10-25 | VanBuren R et al., Desiccation tolerance evolved through gene duplication and network rewiring in Lindernia. Plant Cell. 2018 Dec 30(12):2943-2958. Epub 2018 Oct 25. |
| 353 | *Chrysanthemum nankingense* *(Asteraceae)* | 2018-10-18 | Song C et al., The Chrysanthemum nankingense Genome Provides Insights into the Evolution and Diversification of Chrysanthemum Flowers and Medicinal Traits. Mol Plant. 2018 Dec 3 |
| 352 | *Aquilegia coerulea* *(Ranunculaceae)* | 2018-10-16 | Filiault DL et al., The Aquilegia genome provides insight into adaptive radiation and reveals an extraordinarily polymorphic chromosome with a unique history. Elife. 2018 Oct 16,7 |
| 351 | *Gelsemium sempervirens* *(Gelsemiaceae)* | 2018-10-09 | Franke J et al., Gene Discovery in Gelsemium Highlights Conserved Gene Clusters in Monoterpene Indole Alkaloid Biosynthesis. Chembiochem. 2019 Jan 2 20(1):83-87. Epub 2018 Nov 13. |
| 350 | *Saccharum spontaneum* *(Poaceae)* | 2018-10-08 | Zhang J et al., Allele-defined genome of the autopolyploid sugarcane Saccharum spontaneum L. Nat Genet. 2018 Nov 50(11):1565-1573. Epub 2018 Oct 8. |
| 349 | *Prototheca zopfii* *(Chlorellaceae)* | 2018-10-02 | Severgnini M et al., Genome sequencing of Prototheca zopfii genotypes 1 and 2 provides evidence of a severe reduction in organellar genomes. Sci Rep. 2018 Oct 02, 8: 14637 |
| 348 | *Lavandula angustifolia* *(Lamiaceae)* | 2018-09-29 | Malli RPN et al., De novo sequencing of the Lavandula angustifolia genome reveals highly duplicated and optimized features for essential oil production. Planta. 2019 Jan 249(1):251-256. Epub 2018 Sep 29. |
| 347 | *Ulva mutabilis* *(Ulvaceae)* | 2018-09-24 | De Clerck O et al., Insights into the Evolution of Multicellularity from the Sea Lettuce Genome. Curr Biol. 2018 Sep 24, 28(18): 2921-2933 |
| 346 | *Prunus x yedoensis* *(Rosaceae)* | 2018-09-04 | Baek S et al., Draft genome sequence of wild Prunus yedoensis reveals massive inter-specific hybridization between sympatric flowering cherries. Genome Biol. 2018 Sep 4 |
| 345 | *Picochlorum costavermella* *(C.incertae sedis)* | 2018-09-01 | Krasovec M et al., Genome Analyses of the Microalga Picochlorum Provide Insights into the Evolution of Thermotolerance in the Green Lineage. Genome Biol. Evol. 10(9): 2347-2365 |
| 344 | *Papaver somniferum* *(Papaveraceae)* | 2018-08-30 | Guo L et al., The opium poppy genome and morphinan production. Science. 2018 Oct 19 362(6412):343-347. Epub 2018 Aug 30. |
| 343 | *Vernicia fordii* *(Euphorbiaceae)* | 2018-08-21 | Cui P et al., Tung tree (Vernicia fordii, Hemsl.) genome and transcriptome sequencing reveals coordinate upregulation of fatty acid beta-oxidation and triacylglycerol biosynthesis pathways during eleostearic acid accumulation in seeds. Plant Cell Physiol. 2018 Oct 1 59(10):1990-2003. |
| 342 | *Calamus simplicifolius* *(Arecaceae)* | 2018-08-07 | Zhao H et al., The chromosome-level genome assemblies of two rattans (Calamus simplicifolius and Daemonorops jenkinsiana) GigaScience 2018: giy097 |
| 341 | *Daemonorops jenkinsiana* *(Arecaceae)* | 2018-08-07 | Zhao H et al., The chromosome-level genome assemblies of two rattans (Calamus simplicifolius and Daemonorops jenkinsiana) GigaScience 2018: giy097 |
| 340 | *Populus alba* *(Salicaceae)* | 2018-07-25 | Ma J et al., Genome sequence and genetic transformation of a widely distributed and cultivated poplar Plant Biotechnol J. 2019 Feb 17(2):451-460. Epub 2018 Oct 24. |
| 339 | *Alnus glutinosa* *(Betulaceae)* | 2018-07-13 | Griesmann M et al., Phylogenomics reveals multiple losses of nitrogen-fixing root nodule symbiosis. Science. 2018 Jul 13, 361(6398) |
| 338 | *Begonia fuchsioides* *(Begoniaceae)* | 2018-07-13 | Griesmann M et al., Phylogenomics reveals multiple losses of nitrogen-fixing root nodule symbiosis. Science. 2018 Jul 13, 361(6398) |
| 337 | *Casuarina glauca* *(Casuarinaceae)* | 2018-07-13 | Griesmann M et al., Phylogenomics reveals multiple losses of nitrogen-fixing root nodule symbiosis. Science. 2018 Jul 13, 361(6398) |
| 336 | *Cercis canadensis* *(Fabaceae)* | 2018-07-13 | Griesmann M et al., Phylogenomics reveals multiple losses of nitrogen-fixing root nodule symbiosis. Science. 2018 Jul 13, 361(6398) |
| 335 | *Chamaecrista fasciculata* *(Fabaceae)* | 2018-07-13 | Griesmann M et al., Phylogenomics reveals multiple losses of nitrogen-fixing root nodule symbiosis. Science. 2018 Jul 13, 361(6398) |
| 334 | *Datisca glomerata* *(Datiscaceae)* | 2018-07-13 | Griesmann M et al., Phylogenomics reveals multiple losses of nitrogen-fixing root nodule symbiosis. Science. 2018 Jul 13, 361(6398) |
| 333 | *Dryas drummondii* *(Rosaceae)* | 2018-07-13 | Griesmann M et al., Phylogenomics reveals multiple losses of nitrogen-fixing root nodule symbiosis. Science. 2018 Jul 13, 361(6398) |
| 332 | *Mimosa pudica* *(Fabaceae)* | 2018-07-13 | Griesmann M et al., Phylogenomics reveals multiple losses of nitrogen-fixing root nodule symbiosis. Science. 2018 Jul 13, 361(6398) |
| 331 | *Nissolia schottii* *(Fabaceae)* | 2018-07-13 | Griesmann M et al., Phylogenomics reveals multiple losses of nitrogen-fixing root nodule symbiosis. Science. 2018 Jul 13, 361(6398) |
| 330 | *Ochetophila trinervis* *(Rhamnaceae)* | 2018-07-13 | Griesmann M et al., Phylogenomics reveals multiple losses of nitrogen-fixing root nodule symbiosis. Science. 2018 Jul 13, 361(6398) |
| 329 | *Chara braunii* *(Characeae)* | 2018-07-12 | Nishiyama T et al., The Chara Genome: Secondary Complexity and Implications for Plant Terrestrialization. Cell. 2018 Jul 12, 174(2) |
| 328 | *Cuscuta australis* *(Convolvulaceae)* | 2018-07-11 | Sun G et al., Large-scale gene losses underlie the genome evolution of parasitic plant Cuscuta australis. Nat Commun. 2018 Jul 11, 9(1):2683 |
| 327 | *Myrica rubra* *(Myricaceae)* | 2018-07-10 | Jia HM et al., The red bayberry genome and genetic basis of sex determination Plant Biotechnol J. 2018 Jul 10 |
| 326 | *Saccharum x sp.* *(Poaceae)* | 2018-07-06 | Garsmeur O et al., A mosaic monoploid reference sequence for the highly complex genome of sugarcane. Nat Commun. 2018 Jul 6, 9(1): 2638 |
| 325 | *Azolla filiculoides* *(Salviniaceae)* | 2018-07-02 | Li FW et al., Fern genomes elucidate land plant evolution and cyanobacterial symbioses. Nat Plants. 2018 Jul 4(7): 460-472. Epub 2018 Jul 2. |
| 324 | *Salvinia cucullata* *(Salviniaceae)* | 2018-07-02 | Li FW et al., Fern genomes elucidate land plant evolution and cyanobacterial symbioses. Nat Plants. 2018 Jul 4(7): 460-472. Epub 2018 Jul 2. |
| 323 | *Oryza granulata* *(Poaceae)* | 2018-06-29 | Wu Z et al., De novo genome assembly of Oryza granulata reveals rapid genome expansion and adaptive evolution. Commun Biol. 2018 Jun 29, 1:84 |
| 322 | *Cuscuta campestris* *(Convolvulaceae)* | 2018-06-28 | Vogel A et al., Footprints of parasitism in the genome of the parasitic flowering plant Cuscuta campestris. Nat Commun. 2018 Jun 28, 9(1): 2515 |
| 321 | *Salvia splendens* *(Lamiaceae)* | 2018-06-19 | Dong AX et al., High quality assembly of the reference genome for scarlet sage, Salvia splendens, an economically important ornamental plant. Gigascience. 2018 Jul 1 7(7). |

| | plant species | published | |
|---|---|---|---|
| 320 | *Ammopiptanthus nanus* *(Fabaceae)* | 2018-06-18 | Gao F et al., Long-read sequencing and de novo genome assembly of Ammopiptanthus nanus, a desert shrub. GigaScience 2018, giy074 |
| 319 | *Quercus robur* *(Fagaceae)* | 2018-06-18 | Plomion C et al., Oak genome reveals facets of long lifespan. Nat Plants. 2018 Jul 4(7):440-452. Epub 2018 Jun 18. |
| 318 | *Citrus reticulata* *(Rutaceae)* | 2018-06-06 | Wang L et al., Genome of wild mandarin and domestication history of mandarin Mol Plant. 2018 Aug 6 11(8):1024-1037. Epub 2018 Jun 6. |
| 317 | *Arachis monticola* *(Fabaceae)* | 2018-06-01 | Yin D et al., Genome of an allotetraploid wild peanut Arachis monticola: a de novo assembly. Gigascience. 2018 Jun 1, 7(6) |
| 316 | *Fagus sylvatica* *(Fagaceae)* | 2018-05-28 | Mishra B et al., A reference genome of the European Beech (Fagus sylvatica L.) GigaScience 2018, giy063 |
| 315 | *Tectona grandis* *(Lamiaceae)* | 2018-05-24 | Yasodha R et al., Draft genome of a high value tropical timber tree, Teak (Tectona grandis L. f): insights into SSR diversity, phylogeny and conservation. DNA Res. 2018 Aug 1 25(4):409-419. |
| 314 | *Juglans cathayensis* *(Juglandaceae)* | 2018-05-23 | Stevens KA et al., Genomic Variation Among and Within Six Juglans Species. G3 (Bethesda). 2018 Jul 2 8(7):2153-2165. |
| 313 | *Juglans hindsii* *(Juglandaceae)* | 2018-05-23 | Stevens KA et al., Genomic Variation Among and Within Six Juglans Species. G3 (Bethesda). 2018 Jul 2 8(7):2153-2165. |
| 312 | *Juglans microcarpa* *(Juglandaceae)* | 2018-05-23 | Stevens KA et al., Genomic Variation Among and Within Six Juglans Species. G3 (Bethesda). 2018 Jul 2 8(7):2153-2165. |
| 311 | *Juglans nigra* *(Juglandaceae)* | 2018-05-23 | Stevens KA et al., Genomic Variation Among and Within Six Juglans Species. G3 (Bethesda). 2018 Jul 2 8(7):2153-2165. |
| 310 | *Juglans sigillata* *(Juglandaceae)* | 2018-05-23 | Stevens KA et al., Genomic Variation Among and Within Six Juglans Species. G3 (Bethesda). 2018 Jul 2 8(7):2153-2165. |
| 309 | *Pterocarya stenoptera* *(Juglandaceae)* | 2018-05-23 | Stevens KA et al., Genomic Variation Among and Within Six Juglans Species. G3 (Bethesda). 2018 Jul 2 8(7):2153-2165. |
| 308 | *Raphidocelis subcapitata* *(Selenastraceae)* | 2018-05-23 | Suzuki S et al., Raphidocelis subcapitata (=Pseudokirchneriella subcapitata) provides an insight into genome evolution and environmental adaptations in the Sphaeropleales. Sci Rep. 2018 May 23, 8(1): 8058 |
| 307 | *Quercus suber* *(Fagaceae)* | 2018-05-22 | Ramos AM et al., The draft genome sequence of cork oak. Sci Data. 2018 May 22, 5:180069 |
| 306 | *Selaginella tamariscina* *(Selaginellaceae)* | 2018-05-16 | Xu Z et al., Genome analysis of the ancient tracheophyte Selaginella tamariscina reveals evolutionary features relevant to the acquisition of desiccation tolerance. Mol Plant. 2018 Jul 2 11(7): 983-994. Epub 2018 May 17. |
| 305 | *Bombax ceiba* *(Malvaceae)* | 2018-05-10 | Gao Y et al., De novo genome assembly of the red silk cotton tree (Bombax ceiba) Gigascience. 2018 May 1 7(5). |
| 304 | *Parasponia andersonii* *(Cannabaceae)* | 2018-05-01 | van Velzen R et al., Comparative genomics of the nonlegume Parasponia reveals insights into evolution of nitrogen-fixing rhizobium symbioses. Proc Natl Acad Sci U S A. 2018 May 15 115(20):E4700-E4709. Epub 2018 May 1. |
| 303 | *Trema orientalis* *(Cannabaceae)* | 2018-05-01 | van Velzen R et al., Comparative genomics of the nonlegume Parasponia reveals insights into evolution of nitrogen-fixing rhizobium symbioses. Proc Natl Acad Sci U S A. 2018 May 15 115(20):E4700-E4709. Epub 2018 May 1. |
| 302 | *Rosa chinensis* *(Rosaceae)* | 2018-04-30 | Raymond O et al., The Rosa genome provides new insights into the domestication of modern roses. Nat Genet. 2018 Jun 50(6):772-777. Epub 2018 Apr 30. |
| 301 | *Phalaenopsis aphrodite* *(Orchidaceae)* | 2018-04-28 | Chao YT et al., Chromosome-level assembly, genetic and physical mapping of Phalaenopsis aphrodite genome provides new insights into species adaptation and resources for orchid breeding. Plant Biotechnol J. 2018 Dec 16(12):2027-2041. Epub 2018 May 23. |
| 300 | *Gastrodia elata* *(Orchidaceae)* | 2018-04-24 | Yuan Y et al., The Gastrodia elata genome provides insights into plant adaptation to heterotrophy. Nat Commun. 2018 Apr 24, 9(1):1615. |
| 299 | *Gracilariopsis chorda* *(Gracilariaceae)* | 2018-04-23 | Lee J et al., Analysis of the draft genome of the red seaweed Gracilariopsis chorda provides insights into genome size evolution in Rhodophyta. Mol Biol Evol. 2018 Aug 1 35(8): 1869-1886. |
| 298 | *Artemisia annua* *(Asteraceae)* | 2018-04-20 | Shen Q et al., The Genome of Artemisia annua Provides Insight into the Evolution of Asteraceae Family and Artemisinin Biosynthesis. Mol Plant. 2018 Jun 4 11(6):776-788. Epub 2018 Apr 24. |
| 297 | *Glycine latifolia* *(Fabaceae)* | 2018-04-19 | Liu Q et al., Assembly and annotation of a draft genome sequence for Glycine latifolia, a perennial wild relative of soybean Plant J. 2018 Jul 95(1):71-85. Epub 2018 May 23. |
| 296 | *Chlorella vulgaris* *(Chlorellaceae)* | 2018-04-05 | Guarnieri MT et al., Genome Sequence of the Oleaginous Green Alga, Chlorella vulgaris UTEX 395. Front Bioeng Biotechnol. 2018 Apr 5, 6: 37 |
| 295 | *Boechera retrofracta* *(Brassicaceae)* | 2018-03-28 | Kliver S et al., Assembly of the Boechera retrofracta Genome and Evolutionary Analysis of Apomixis-Associated Genes. Genes (Basel). 2018 Mar 28, 9(4) |
| 294 | *Eudorina sp.* *(Volvocaceae)* | 2018-03-08 | Hamaji T et al., Anisogamy evolved with a reduced sex-determining region in volvocine green algae. Commun Biol. 2018 Mar 8, 1: 17 eCollection 2018. |
| 293 | *Yamagishiella unicocca* *(Volvocaceae)* | 2018-03-08 | Hamaji T et al., Anisogamy evolved with a reduced sex-determining region in volvocine green algae. Commun Biol. 2018 Mar 8, 1: 17 eCollection 2018. |
| 292 | *Coffea arabica* *(Rubiaceae)* | 2018-03-06 | Tran HTM et al., Use of a draft genome of coffee (Coffea arabica) to identify SNPs associated with caffeine content. Plant Biotechnol J. 2018 Oct 16(10):1756-1766. Epub 2018 Apr 13. |
| 291 | *Potentilla micrantha* *(Rosaceae)* | 2018-02-15 | Buti M et al., The genome sequence and transcriptome of Potentilla micrantha and their comparison to Fragaria vesca (the woodland strawberry). GigaScience 2018: giy010 |
| 290 | *Santalum album* *(Santalaceae)* | 2018-02-12 | Mahesh HB et al., Multi-omics driven assembly and annotation of the sandalwood (Santalum album) genome. Plant Physiol. 2018 Apr 176(4):2772-2788. Epub 2018 Feb 12. |
| 289 | *Solanum chacoense* *(Solanaceae)* | 2018-02-05 | Leisner CP et al., Genome sequence of M6, a diploid inbred clone of the high glycoalkaloid-producing tuber-bearing potato species Solanum chacoense, reveals residual heterozygosity. Plant J. 2018 May 94(3):562-570. Epub 2018 Mar 22. |
| 288 | *Eutrema heterophyllum* *(Brassicaceae)* | 2018-01-30 | Guo X et al., The genomes of two Eutrema species provide insight into plant adaptation to high altitudes. DNA Res. 2018 Jan 30 25(3): 307-15. Epub ahead of print |
| 287 | *Eutrema yunnanense* *(Brassicaceae)* | 2018-01-30 | Guo X et al., The genomes of two Eutrema species provide insight into plant adaptation to high altitudes. DNA Res. 2018 Jan 30 25(3): 307-15. Epub ahead of print |
| 286 | *Gnetum montanum* *(Gnetaceae)* | 2018-01-29 | Wan T et al., A genome for gnetophytes and early evolution of seed plants Nat Plants. 2018 Feb 4(2): 82-89. Epub 2018 Jan 29. |
| 285 | *Picochlorum soloecismus* *(C.incertae sedis)* | 2018-01-25 | Gonzalez-Esquer CR et al., Nuclear, Chloroplast, and Mitochondrial Genome Sequences of the Prospective Microalgal Biofuel Strain Picochlorum soloecismus. Genome Announc. 2018 Jan 25, 6(4). |
| 284 | *Leersia perrieri* *(Poaceae)* | 2018-01-22 | Stein JC et al., Genomes of 13 domesticated and wild rice relatives highlight genetic conservation, turnover and innovation across the genus Oryza Nat Genet. 2018 Feb 50(2):285-296. Epub 2018 Jan 22. |

| | plant species | published | |
|---|---|---|---|
| 283 | *Oryza punctata* *(Poaceae)* | 2018-01-22 | Stein JC et al., Genomes of 13 domesticated and wild rice relatives highlight genetic conservation, turnover and innovation across the genus Oryza<br>Nat Genet. 2018 Feb 50(2):285-296. Epub 2018 Jan 22. |
| 282 | *Oryza rufipogon* *(Poaceae)* | 2018-01-22 | Stein JC et al., Genomes of 13 domesticated and wild rice relatives highlight genetic conservation, turnover and innovation across the genus Oryza<br>Nat Genet. 2018 Feb 50(2):285-296. Epub 2018 Jan 22. |
| 281 | *Selaginella lepidophylla* *(Selaginellaceae)* | 2018-01-02 | VanBuren R et al., Extreme haplotype variation in the desiccation-tolerant clubmoss Selaginella lepidophylla.<br>Nat Commun. 2018 Jan 2, 9(1):13 |
| 280 | *Eschscholzia californica* *(Papaveraceae)* | 2017-12-29 | Hori K et al., Mining of the uncharacterized cytochrome P450 genes involved in alkaloid biosynthesis in California poppy using a draft genome sequence<br>Plant Cell Physiol. 2018 Feb 1 59(2):222-233. |
| 279 | *Tetrabaena socialis* *(Tetrabaenaceae)* | 2017-12-26 | Featherston J et al., The 4-celled Tetrabaena socialis nuclear genome reveals the essential components for genetic control of cell number at the origin of multicellularity in the volvocine lineage.<br>Mol Biol Evol. 2018 Apr 1 35(4): 855-870. |
| 278 | *Oryza coarctata* *(Poaceae)* | 2017-12-15 | Mondal TK et al., First de novo draft genome sequence of Oryza coarctata, the only halophytic species in the genus Oryza.<br>F1000Res. 2017 Dec 15, 6: 1750 |
| 277 | *Handroanthus impetiginosus* *(Bignoniaceae)* | 2017-12-13 | da Silva-Junior OB et al., Genome assembly of the Pink Ipe (Handroanthus impetiginosus, Bignoniaceae), a highly-valued ecologically keystone Neotropical timber forest tree<br>Gigascience. 2018 Jan 1 7(1):1-16. |
| 276 | *Calotropis gigantea* *(Apocynaceae)* | 2017-12-12 | Hoopes GM et al., Genome Assembly and Annotation of the Medicinal Plant Calotropis gigantea, a Producer of Anti-Cancer and Anti-Malarial Cardenolides<br>G3 (Bethesda). 2018 Feb 2 8(2):385-391. |
| 275 | *Eucommia ulmoides* *(Eucommiaceae)* | 2017-12-08 | Wuyun TN et al., The hardy rubber tree genome provides insights into the evolution of polyisoprene biosynthesis.<br>Mol Plant. 2018 Mar 5 11(3):429-442. Epub 2017 Dec 9. |
| 274 | *Citrus unshiu* *(Rutaceae)* | 2017-12-05 | Shimizu T1 et al., Draft Sequencing of the Heterozygous Diploid Genome of Satsuma (Citrus unshiu Marc.) Using a Hybrid Assembly Approach.<br>Front Genet. 2017 Dec 5 |
| 273 | *Kalanchoe fedtschenkoi* *(Crassulaceae)* | 2017-12-01 | Yang X et al., The Kalanchoe genome provides insights into convergent evolution and building blocks of crassulacean acid metabolism<br>Nat Commun. 2017 Dec 1, 8(1): 1899 |
| 272 | *Chlorella sorokiniana* *(Chlorellaceae)* | 2017-11-27 | Arriola MB et al., Genome Sequences of Chlorella sorokiniana UTEX 1602 and Micractinium conductrix SAG 241.80: Implications to Maltose Excretion by a Green Alga<br>Plant J. 2018 Feb 93(3): 566-586. Epub 2018 Jan 10. |
| 271 | *Micractinium conductrix* *(Chlorellaceae)* | 2017-11-27 | Arriola MB et al., Genome Sequences of Chlorella sorokiniana UTEX 1602 and Micractinium conductrix SAG 241.80: Implications to Maltose Excretion by a Green Alga<br>Plant J. 2018 Feb 93(3): 566-586. Epub 2018 Jan 10. |
| 270 | *Boehmeria nivea* *(Urticaceae)* | 2017-11-15 | Liu C et al., Draft genome analysis provides insights into the fiber yield, crude protein biosynthesis, and vegetative growth of domesticated ramie (Boehmeria nivea L. Gaud)<br>DNA Res. 2018 Apr 1 25(2):173-181. |
| 269 | *Cucurbita pepo* *(Cucurbitaceae)* | 2017-11-07 | Montero-Pau J et al., De novo assembly of the zucchini genome reveals a whole genome duplication associated with the origin of the Cucurbita genus.<br>Plant Biotechnol J. 2018 Jun 16(6):1161-1171. Epub 2017 Dec 4. |
| 268 | *Asparagus officinalis* *(Asparagaceae)* | 2017-11-02 | Harkess A et al., The asparagus genome sheds light on the origin and evolution of a young Y chromosome.<br>Nat Commun. 2017 Nov 2, 8(1):1279 |
| 267 | *Capsicum baccatum* *(Solanaceae)* | 2017-11-01 | Kim S et al., New reference genome sequences of hot pepper reveal the massive evolution of plant disease-resistance genes by retroduplication<br>Genome Biol. 2017 Nov 1, 18(1):210 |
| 266 | *Capsicum chinense* *(Solanaceae)* | 2017-11-01 | Kim S et al., New reference genome sequences of hot pepper reveal the massive evolution of plant disease-resistance genes by retroduplication<br>Genome Biol. 2017 Nov 1, 18(1):210 |
| 265 | *Dunaliella salina* *(Dunaliellaceae)* | 2017-10-26 | Polle JEW et al., Draft Nuclear Genome Sequence of the Halophilic and Beta-Carotene-Accumulating Green Alga Dunaliella salina Strain CCAP19/18.<br>Genome Announc. 2017 Oct 26, 5(43): e01105-17 |
| 264 | *Carnegiea gigantea* *(Cactaceae)* | 2017-10-23 | Copetti D et al., Extensive gene tree discordance and hemiplasy shaped the genomes of North American columnar cacti<br>Proc Natl Acad Sci U S A. 2017 Nov 7 114(45):12003-12008. Epub 2017 Oct 23. |
| 263 | *Echinochloa crus-galli* *(Poaceae)* | 2017-10-18 | Guo L et al., Echinochloa crus-galli genome analysis provides insight into its adaptation and invasiveness as a weed.<br>Nat Commun. 2017 Oct 18 8(1):1031. |
| 262 | *Rosa multiflora* *(Rosaceae)* | 2017-10-16 | Nakamura N et al., Genome structure of Rosa multiflora, a wild ancestor of cultivated roses<br>DNA Res. 2018 Apr 1 25(2):113-121. |
| 261 | *Durio zibethinus* *(Malvaceae)* | 2017-10-09 | Teh BT et al., The draft genome of tropical fruit durian (Durio zibethinus)<br>Nat Genet. 2017 Nov 49(11):1633-1641. Epub 2017 Oct 9. |
| 260 | *Cocos nucifera* *(Arecaceae)* | 2017-10-05 | Xiao Y et al., The genome draft of Coconut (Cocos nucifera)<br>Gigascience. 2017 Nov 1, 6(11): 1-11 |
| 259 | *Marchantia polymorpha* *(Marchantiaceae)* | 2017-10-05 | Bowman JL et al., Insights into Land Plant Evolution Garnered from the Marchantia polymorpha Genome.<br>Cell. 2017 Oct 5, 171(2): 287-304 |
| 258 | *Panax ginseng* *(Araliaceae)* | 2017-10-05 | Jiang X et al., Panax ginseng genome examination for ginsenoside biosynthesis<br>Gigascience. 2017 Nov 1, 6(11): 1-15 |
| 257 | *Lagenaria siceraria* *(Cucurbitaceae)* | 2017-09-23 | Wu S et al., The bottle gourd genome provides insights in to Cucurbitaceae evolution and facilitates mapping of a Papaya ringspot virus resistance locus<br>Plant J. 2017 Dec 92(5):963-975. Epub 2017 Nov 12. |
| 256 | *Dioscorea cayenensis* *(Dioscoreaceae)* | 2017-09-19 | Tamiru M1 et al., Genome sequencing of the staple food crop white Guinea yam enables the development of a molecular marker for sex determination.<br>BMC Biol. 2017 Sep 19, 15(1):86 |
| 255 | *Cenchrus americanus* *(Poaceae)* | 2017-09-18 | Varshney RK et al., Pearl millet genome sequence provides a resource to improve agronomic traits in arid environments.<br>Nat Biotechnol. 2017 Oct 35(10):969-976. Epub 2017 Sep 18. |
| 254 | *Apostasia shenzhenica* *(Orchidaceae)* | 2017-09-13 | Zhang GQ et al., The Apostasia genome and the evolution of orchids<br>Nature. 2017 Sep 21 549(7672):379-383. Epub 2017 Sep 13. |
| 253 | *Cucurbita maxima* *(Cucurbitaceae)* | 2017-09-13 | Sun H et al., Karyotype stability and unbiased fractionation in the paleo-allotetraploid Cucurbita genomes.<br>Mol Plant. 2017 Oct 9 10(10):1293-1306. Epub 2017 Sep 14. |
| 252 | *Cucurbita moschata* *(Cucurbitaceae)* | 2017-09-13 | Sun H et al., Karyotype stability and unbiased fractionation in the paleo-allotetraploid Cucurbita genomes.<br>Mol Plant. 2017 Oct 9 10(10):1293-1306. Epub 2017 Sep 14. |
| 251 | *Chlamydomonas eustigma* *(Chlamydomonadaceae)* | 2017-09-11 | Hirooka S et al., Acidophilic green algal genome provides insights into adaptation to an acidic environment.<br>Proc Natl Acad Sci U S A. 2017 Sep 26 114(39): E8304-E8313. Epub 2017 Sep 11. |
| 250 | *Gracilaria changii* *(Gracilariaceae)* | 2017-09-07 | Ho CL et al., Unraveling the nuclear and chloroplast genomes of an agar producing red macroalga, Gracilaria changii (Rhodophyta, Gracilariales).<br>Genomics. 2018 Mar, 110(2): 124-133. Epub 2017 Sep 7. |
| 249 | *Pseudotsuga menziesii* *(Pinaceae)* | 2017-09-07 | Neale DB et al., The Douglas-Fir Genome Sequence Reveals Specialization of the Photosynthetic Apparatus in Pinaceae.<br>G3 (Bethesda). 2017 Sep 7, 7(9): 3157-3167 |
| 248 | *Fagopyrum tataricum* *(Polygonaceae)* | 2017-08-30 | Zhang L et al., The Tartary Buckwheat Genome Provides Insights into Rutin Biosynthesis and Abiotic Stress Tolerance.<br>Mol Plant. 2017 Sep 12 10(9):1224-1237. Epub 2017 Sep 1. |
| 247 | *Rhododendron delavayi* *(Ericaceae)* | 2017-08-26 | Zhang L et al., The draft genome assembly of Rhododendron delavayi Franch. var. delavayi<br>Gigascience. 2017 Oct 1, 6(10):1-11 epub: gix076 |

| | plant species | published | |
|---|---|---|---|
| 246 | *Taraxacum kok-saghyz*<br>*(Asteraceae)* | 2017-08-25 | Lin T et al., Genome analysis of Taraxacum kok-saghyz Rodin provides new insights into rubber biosynthesis<br>National Science Review, nwx101 |
| 245 | *Ipomoea batatas*<br>*(Convolvulaceae)* | 2017-08-21 | Yang J et al., Haplotype-resolved sweet potato genome traces back its hexaploidization history.<br>Nat Plants. 2017 Sep 3(9):696-703. Epub 2017 Aug 21. |
| 244 | *Erythranthe luteus*<br>*(Phrymaceae)* | 2017-08-16 | Edger PP et al., Subgenome Dominance in an Interspecific Hybrid, Synthetic Allopolyploid, and a 140-Year-Old Naturally Established Neo-Allopolyploid Monkeyflower.<br>Plant Cell. 2017 Sep, 29(9): 2150-2167 |
| 243 | *Populus pruinosa*<br>*(Salicaceae)* | 2017-08-08 | Yang W et al., The draft genome sequence of a desert tree Populus pruinosa<br>Gigascience. 2017 Sep 1, 6(9): 1-7 |
| 242 | *Camptotheca acuminata*<br>*(Nyssaceae)* | 2017-07-24 | Zhao D et al., De novo genome assembly of Camptotheca acuminata, a natural source of the anti-cancer compound camptothecin<br>Gigascience. 2017 Sep 1 6(9):1-7. |
| 241 | *Porphyra umbilicalis*<br>*(Bangiaceae)* | 2017-07-17 | Brawley SH et al., Insights into the red algae and eukaryotic evolution from the genome of Porphyra umbilicalis (Bangiophyceae, Rhodophyta).<br>Proc Natl Acad Sci U S A. 2017 Aug 1 114(31):E6361-E6370. Epub 2017 Jul 17. |
| 240 | *Triticum turgidum*<br>*(Poaceae)* | 2017-07-07 | Avni R et al., Wild emmer genome architecture and diversity elucidate wheat evolution and domestication.<br>Science. 2017 Jul 7, 357(6346): 93-97 |
| 239 | *Punica granatum*<br>*(Lythraceae)* | 2017-06-27 | Qin G et al., The pomegranate (Punica granatum L.) genome and the genomics of punicalagin biosynthesis<br>Plant J. 2017 Sep 91(6):1108-1128. Epub 2017 Aug 3. |
| 238 | *Chloroidium sp.*<br>*(Watanabeaceae)* | 2017-06-17 | Nelson DR et al., The genome and phenome of the green alga Chloroidium sp. UTEX 3007 reveal adaptive traits for desert acclimatization.<br>Elife. 2017 Jun 17 |
| 237 | *Eleusine coracana*<br>*(Poaceae)* | 2017-06-15 | Hittalmani S et al., Genome and Transcriptome sequence of Finger millet (Eleusine coracana (L.) Gaertn.) provides insights into drought tolerance and nutraceutical properties.<br>BMC Genomics. 2017 Jun 15 18(1):465. |
| 236 | *Rhizophora apiculata*<br>*(Rhizophoraceae)* | 2017-06-05 | Xu S et al., The origin, diversification and adaptation of a major mangrove clade (Rhizophoreae) revealed by whole-genome sequencing<br>National Science Review, 2017 Jun 5, nwx065 |
| 235 | *Macleaya cordata*<br>*(Papaveraceae)* | 2017-05-25 | Liu X et al., The genome of the medicinal plant Macleaya cordata provides new insights into benzylisoquinoline alkaloids metabolism.<br>Mol Plant. 2017 Jul 5 10(7):975-989. Epub 2017 May 26. |
| 234 | *Prunus avium*<br>*(Rosaceae)* | 2017-05-25 | Shirasawa K et al., The genome sequence of sweet cherry (Prunus avium) for use in genomics-assisted breeding.<br>DNA Res. 2017 Oct 1 24(5):499-508. |
| 233 | *Spinacia oleracea*<br>*(Amaranthaceae)* | 2017-05-24 | Xu C et al., Draft genome of spinach and transcriptome diversity of 120 Spinacia accessions.<br>Nat Commun. 2017 May 24 8:15275. |
| 232 | *Nicotiana attenuata*<br>*(Solanaceae)* | 2017-05-23 | Xu S et al., Wild tobacco genomes reveal the evolution of nicotine biosynthesis<br>Proc Natl Acad Sci U S A. 2017 Jun 6 114(23):6133-6138. Epub 2017 May 23. |
| 231 | *Nicotiana obtusifolia*<br>*(Solanaceae)* | 2017-05-23 | Xu S et al., Wild tobacco genomes reveal the evolution of nicotine biosynthesis<br>Proc Natl Acad Sci U S A. 2017 Jun 6 114(23):6133-6138. Epub 2017 May 23. |
| 230 | *Helianthus annuus*<br>*(Asteraceae)* | 2017-05-22 | Badouin H et al., The sunflower genome provides insights into oil metabolism, flowering and Asterid evolution<br>Nature. 2017 Jun 1 546(7656):148-152. Epub 2017 May 22. |
| 229 | *Betula pendula*<br>*(Betulaceae)* | 2017-05-08 | Salojaervi J et al., Genome sequencing and population genomic analyses provide insights into the adaptive landscape of silver birch.<br>Nat Genet. 2017 Jun 49(6):904-912. Epub 2017 May 8. |
| 228 | *Chromochloris zofingiensis*<br>*(Chromochloridaceae)* | 2017-05-08 | Roth MS et al., Chromosome-level genome assembly and transcriptome of the green alga Chromochloris zofingiensis illuminates astaxanthin production.<br>Proc Natl Acad Sci U S A. 2017 May 23 114(21):E4296-E4305. Epub 2017 May 8. |
| 227 | *Rhodiola crenulata*<br>*(Crassulaceae)* | 2017-05-05 | Fu Y et al., Draft genome sequence of the Tibetan medicinal herb, Rhodiola crenulata.<br>Gigascience. 2017 Jun 1 6(6):1-5. |
| 226 | *Camellia sinensis*<br>*(Theaceae)* | 2017-05-01 | Xia EH et al., The tea tree genome provides insights into tea flavor and independent evolution of caffeine biosynthesis<br>Mol Plant. 2017 Jun 5 10(6):866-877. Epub 2017 May 2. |
| 225 | *Botryococcus braunii*<br>*(Botryococcaceae)* | 2017-04-20 | Browne DR et al., Draft Nuclear Genome Sequence of the Liquid Hydrocarbon–Accumulating Green Microalga Botryococcus braunii Race B (Showa)<br>Genome Announc. 2017 Apr 20 5(16). |
| 224 | *Erigeron breviscapus*<br>*(Asteraceae)* | 2017-04-18 | Yang J et al., Hybrid de novo genome assembly of the Chinese herbal fleabane Erigeron breviscapus<br>Gigascience. 2017 Jun 1 6(6):1-7. |
| 223 | *Lactuca sativa*<br>*(Asteraceae)* | 2017-04-12 | Reyes-Chin-Wo S et al., Genome assembly with in vitro proximity ligation data and whole-genome triplication in lettuce<br>Nat Commun. 2017 Apr 12 8:14953. |
| 222 | *Atalantia buxifolia*<br>*(Rutaceae)* | 2017-04-10 | Wang X et al., Genomic analyses of primitive, wild and cultivated citrus provide insights into asexual reproduction<br>Nat Genet. 2017 May 49(5):765-772. Epub 2017 Apr 10. |
| 221 | *Citrus cavaleriei*<br>*(Rutaceae)* | 2017-04-10 | Wang X et al., Genomic analyses of primitive, wild and cultivated citrus provide insights into asexual reproduction<br>Nat Genet. 2017 May 49(5):765-772. Epub 2017 Apr 10. |
| 220 | *Citrus maxima*<br>*(Rutaceae)* | 2017-04-10 | Wang X et al., Genomic analyses of primitive, wild and cultivated citrus provide insights into asexual reproduction<br>Nat Genet. 2017 May 49(5):765-772. Epub 2017 Apr 10. |
| 219 | *Citrus medica*<br>*(Rutaceae)* | 2017-04-10 | Wang X et al., Genomic analyses of primitive, wild and cultivated citrus provide insights into asexual reproduction<br>Nat Genet. 2017 May 49(5):765-772. Epub 2017 Apr 10. |
| 218 | *Capsella bursa-pastoris*<br>*(Brassicaceae)* | 2017-04-07 | Kasianov AS et al., High quality genome assembly of Capsella bursa-pastoris reveals asymmetry of regulatory elements at early stages of polyploid genome evolution<br>Plant J. 2017 Jul 91(2):278-291. Epub 2017 Jun 12. |
| 217 | *Boechera stricta*<br>*(Brassicaceae)* | 2017-04-03 | Lee CR et al., Young inversion with multiple linked QTLs under selection in a hybrid zone.<br>Nat Ecol Evol. 2017 Apr 3, 1(5): 119. |
| 216 | *Dimocarpus longan*<br>*(Sapindaceae)* | 2017-03-28 | Lin Y et al., Genome-wide sequencing of longan (Dimocarpus longan Lour.) provides insights into molecular basis of its polyphenol-rich characteristics<br>Gigascience. 2017 May 1 6(5):1-14. |
| 215 | *Xerophyta viscosa*<br>*(Velloziaceae)* | 2017-03-27 | Costa MD et al., A footprint of desiccation tolerance in the genome of Xerophyta viscosa<br>Nat Plants. 2017 Mar 27 3:17038. |
| 214 | *Panax notoginseng*<br>*(Araliaceae)* | 2017-03-15 | Chen W et al., Whole-genome Sequencing and Analysis of the Chinese herbal Plant Panax Notoginseng<br>Mol Plant. 2017 Jun 5 10(6):899-902. Epub 2017 Mar 15. |
| 213 | *Chenopodium suecicum*<br>*(Amaranthaceae)* | 2017-02-16 | Jarvis DE et al., The genome of Chenopodium quinoa.<br>Nature. 2017 Feb 16 542(7641):307-312. Epub 2017 Feb 8. |
| 212 | *Secale cereale*<br>*(Poaceae)* | 2017-02-08 | Bauer E et al., Towards a whole-genome sequence for rye (Secale cereale L.).<br>Plant J. 2017 Mar 89(5):853-869. Epub 2017 Feb 8. |
| 211 | *Cephalotus follicularis*<br>*(Cephalotaceae)* | 2017-02-06 | Fukushima K et al., Genome of the pitcher plant Cephalotus reveals genetic changes associated with carnivory<br>Nat Ecol Evol. 2017 Feb 6, 1(3):59 |
| 210 | *Corchorus capsularis*<br>*(Malvaceae)* | 2017-01-30 | Islam MS et al., Comparative genomics of two jute species and insight into fibre biogenesis.<br>Nat Plants. 2017 Jan 30 3:16223. |

| | plant species | published | |
|---|---|---|---|
| 209 | *Corchorus olitorius* *(Malvaceae)* | 2017-01-30 | Islam MS et al., Comparative genomics of two jute species and insight into fibre biogenesis. Nat Plants. 2017 Jan 30 3:16223. |
| 208 | *Ruellia speciosa* *(Acanthaceae)* | 2017-01-27 | Zhuang Y et al., The draft genome of Ruellia speciosa (Beautiful Wild Petunia: Acanthaceae). DNA Res. 2017 Apr 1, 24(2): 179-192. |
| 207 | *Ficus carica* *(Moraceae)* | 2017-01-25 | Mori K et al., Identification of RAN1 orthologue associated with sex determination through whole genome sequencing analysis in fig (Ficus carica L.) Sci Rep. 2017 Jan 25 7:41124. |
| 206 | *Tetradesmus obliquus* *(Scenedesmaceae )* | 2017-01-19 | Carreres BM et al., Draft Genome Sequence of the Oleaginous Green Alga Tetradesmus obliquus UTEX 393 Genome Announc. 2017 Jan 19 5(3). |
| 205 | *Barbarea vulgaris* *(Brassicaceae)* | 2017-01-17 | Byrne SL et al., The genome sequence of Barbarea vulgaris facilitates the study of ecological biochemistry. Sci Rep. 2017 Jan 17 7:40728. |
| 204 | *Momordica charantia* *(Cucurbitaceae)* | 2016-12-27 | Urasaki N et al., Draft genome sequence of bitter gourd (Momordica charantia), a vegetable and medicinal plant in tropical and subtropical regions. DNA Res. 2017 Feb 1 24(1):51-58. |
| 203 | *Fraxinus excelsior* *(Oleaceae)* | 2016-12-26 | Sollars ES et al., Genome sequence and genetic diversity of European ash trees. Nature. 2017 Jan 12 541(7636):212-216. Epub 2016 Dec 26. |
| 202 | *Hibiscus syriacus* *(Malvaceae)* | 2016-12-22 | Kim YM et al., Genome analysis of Hibiscus syriacus provides insights of polyploidization and indeterminate flowering in woody plants DNA Res. 2017 Feb 1 24(1):71-80. |
| 201 | *Pinus lambertiana* *(Pinaceae)* | 2016-12-01 | Stevens KA et al., Sequence of the Sugar Pine Megagenome Genetics. 2016 Dec 204(4):1613-1626. Epub 2016 Oct 28. |
| 200 | *Ginkgo biloba* *(Ginkgoaceae)* | 2016-11-21 | Guan R et al., Draft genome of the living fossil Ginkgo biloba. Gigascience. 2016 Nov 21 5(1):49. |
| 199 | *Macadamia integrifolia* *(Proteaceae)* | 2016-11-17 | Nock CJ et al., Genome and transcriptome sequencing characterises the gene space of Macadamia integrifolia (Proteaceae). BMC Genomics. 2016 Nov 17 17(1):937. |
| 198 | *Mentha longifolia* *(Lamiaceae)* | 2016-11-17 | Vining KJ et al., Draft Genome Sequence of Mentha longifolia (L.) and Development of Resources for Mint Cultivar Improvement. Mol Plant. 2017 Feb 13 10(2):323-339. Epub 2016 Nov 17. |
| 197 | *Ipomoea nil* *(Convolvulaceae)* | 2016-11-08 | Hoshino A et al., Genome sequence and analysis of the Japanese morning glory Ipomoea nil Nat Commun. 2016 Nov 8 7:13295. |
| 196 | *Siraitia grosvenorii* *(Cucurbitaceae)* | 2016-11-07 | Itkin M et al., The biosynthetic pathway of the nonsugar, high-intensity sweetener mogroside V from Siraitia grosvenorii. Proc Natl Acad Sci U S A. 2016 Nov 22 113(47):E7619-E7628. Epub 2016 Nov 7. |
| 195 | *Cardamine hirsuta* *(Brassicaceae)* | 2016-10-31 | Gan X et al., The Cardamine hirsuta genome offers insight into the evolution of morphological diversity. Nat Plants. 2016 Oct 31 2(11):16167. |
| 194 | *Dichanthelium oligosanthes* *(Poaceae)* | 2016-10-28 | Studer AJ et al., The draft genome of the C3 panicoid grass species Dichanthelium oligosanthes. Genome Biol. 2016 Oct 28 17(1):223. |
| 193 | *Glycyrrhiza uralensis* *(Fabaceae)* | 2016-10-24 | Mochida K et al., Draft genome assembly and annotation of Glycyrrhiza uralensis, a medicinal legume. Plant J. 2017 Jan 89(2):181-194. Epub 2016 Oct 24. |
| 192 | *Rhazya stricta* *(Apocynaceae)* | 2016-09-22 | Sabir JS et al., The nuclear genome of Rhazya stricta and the evolution of alkaloid diversity in a medically relevant clade of Apocynaceae. Sci Rep. 2016 Sep 22 6:33782. |
| 191 | *Quercus lobata* *(Fagaceae)* | 2016-09-12 | Sork VL et al., First Draft Assembly and Annotation of the Genome of a California Endemic Oak Quercus lobata Née (Fagaceae). G3 (Bethesda). 2016 Nov 8 6(11):3485-3495. |
| 190 | *Brassica juncea* *(Brassicaceae)* | 2016-09-05 | Yang J et al., The genome sequence of allopolyploid Brassica juncea and analysis of differential homoeolog gene expression influencing selection. Nat Genet. 2016 Oct 48(10):1225-32. Epub 2016 Sep 5. |
| 189 | *Brassica nigra* *(Brassicaceae)* | 2016-09-05 | Yang J et al., The genome sequence of allopolyploid Brassica juncea and analysis of differential homoeolog gene expression influencing selection. Nat Genet. 2016 Oct 48(10):1225-32. Epub 2016 Sep 5. |
| 188 | *Cicer reticulatum* *(Fabaceae)* | 2016-08-26 | Gupta S et al., Draft genome sequence of Cicer reticulatum L., the wild progenitor of chickpea provides a resource for agronomic trait improvement. DNA Res. 2017 Feb 1 24(1):1-10. |
| 187 | *Trifolium subterranum* *(Fabaceae)* | 2016-08-22 | Hirakawa H et al., Draft genome sequence of subterranean clover, a reference for genus Trifolium. Sci Rep. 2016 Aug 22 6:30358. |
| 186 | *Musa itinerans* *(Musaceae)* | 2016-08-17 | Wu W et al., Whole genome sequencing of a banana wild relative Musa itinerans provides insights into lineage-specific diversification of the Musa genus Sci Rep. 2016 Aug 17 6:31586. |
| 185 | *Chenopodium quinoa* *(Amaranthaceae)* | 2016-07-25 | Yasui Y et al., Draft genome sequence of an inbred line of Chenopodium quinoa, an allotetraploid crop with great environmental adaptability and outstanding nutritional properties DNA Res. 2016 Dec 23(6):535-546. Epub 2016 Jul 25. |
| 184 | *Artocarpus camansi* *(Moraceae)* | 2016-07-13 | Gardner EM et al., Low-coverage, whole-genome sequencing of Artocarpus camansi (Moraceae) for phylogenetic marker development and gene discovery Appl Plant Sci. 2016 Jul 13 4(7). eCollection 2016 Jul. |
| 183 | *Citrus x aurantium* *(Rutaceae)* | 2016-07-08 | Zhang Y et al., Transcriptome analysis of root response to citrus blight based on the newly assembled Swingle citrumelo draft genome. BMC Genomics. 2016 Jul 8 17:485. |
| 182 | *Zostera muelleri* *(Zosteraceae)* | 2016-07-03 | Lee HT et al., The genome of a southern hemisphere seagrass species (Zostera muelleri) Plant Physiol. 2016 Sep 172(1):272-83. Epub 2016 Jul 3. |
| 181 | *Drosera capensis* *(Droseraceae)* | 2016-06-29 | Butts CT et al., Novel proteases from the genome of the carnivorous plant Drosera capensis: structural prediction and comparative analysis Proteins. 2016 Oct 84(10):1517-33. Epub 2016 Jul 13. |
| 180 | *Olea europaea* *(Oleaceae)* | 2016-06-27 | Cruz F et al., Genome sequence of the olive tree, Olea europaea. Gigascience. 2016 Jun 27 5:29. |
| 179 | *Petunia axillaris* *(Solanaceae)* | 2016-05-27 | Bombarely A et al., Insight into the evolution of the Solanaceae from the parental genomes of Petunia hybrida Nature Plants 2: 16074 (2016) |
| 178 | *Petunia inflata* *(Solanaceae)* | 2016-05-27 | Bombarely A et al., Insight into the evolution of the Solanaceae from the parental genomes of Petunia hybrida Nature Plants 2: 16074 (2016) |
| 177 | *Rubus occidentalis* *(Rosaceae)* | 2016-05-26 | VanBuren R et al., The genome of black raspberry (Rubus occidentalis). Plant J. 2016 Sep 87(6):535-47. Epub 2016 Jul 20. |
| 176 | *Pogostemon cablin* *(Lamiaceae)* | 2016-05-20 | He Y et al., Survey of the genome of Pogostemon cablin provides insights into its evolutionary history and sesquiterpenoid biosynthesis. Sci Rep. 2016 May 20 6:26405. |
| 175 | *Phalaenopsis hybrid cv. KHM190* *(Orchidaceae)* | 2016-05-12 | Huang JZ et al., The genome and transcriptome of Phalaenopsis yield insights into floral organ development and flowering regulation PeerJ. 2016 May 12 4:e2017. eCollection 2016. |
| 174 | *Lepidium meyenii* *(Brassicaceae)* | 2016-05-10 | Zhang J et al., Genome of plant maca (Lepidium meyenii) illuminates genomic basis for high altitude adaptation in the central Andes. Mol Plant. 2016 Jul 6 9(7):1066-77. Epub 2016 May 10. |
| 173 | *Daucus carota* *(Apiaceae)* | 2016-05-09 | Iorizzo M et al., A high-quality carrot genome assembly provides new insights into carotenoid accumulation and asterid genome evolution. Nat Genet. 2016 Jun 48(6):657-66. Epub 2016 May 9. |

| | plant species | published | |
|---|---|---|---|
| 172 | *Juglans regia* (Juglandaceae) | 2016-05-03 | Martinez-Garcia PJ et al., The walnut (Juglans regia) genome sequence reveals diversity in genes coding for the biosynthesis of nonstructural polyphenols. Plant J. 2016 Sep 87(5):507-32. Epub 2016 Jul 18. |
| 171 | *Gonium pectorale* (Goniaceae) | 2016-04-22 | Hanschen ER et al., The Gonium pectorale genome demonstrates co-option of cell cycle regulation during the evolution of multicellularity. Nat Commun. 2016 Apr 22 7:11370. |
| 170 | *Fagopyrum esculentum* (Polygonaceae) | 2016-04-02 | Yasui Y et al., Assembly of the draft genome of buckwheat and its applications in identifying agronomically useful genes. DNA Res. 2016 Jun 23(3):215-24. Epub 2016 Apr 2. |
| 169 | *Zoysia japonica* (Poaceae) | 2016-03-14 | Tanaka et al., Sequencing and comparative analyses of the genomes of Zoysiagrasses DNA Res. 2016 Apr 23(2):171-80. Epub 2016 Mar 14. |
| 168 | *Zoysia matrella* (Poaceae) | 2016-03-14 | Tanaka et al., Sequencing and comparative analyses of the genomes of Zoysiagrasses DNA Res. 2016 Apr 23(2):171-80. Epub 2016 Mar 14. |
| 167 | *Zoysia pacifica* (Poaceae) | 2016-03-14 | Tanaka et al., Sequencing and comparative analyses of the genomes of Zoysiagrasses DNA Res. 2016 Apr 23(2):171-80. Epub 2016 Mar 14. |
| 166 | *Arachis duranensis* (Fabaceae) | 2016-02-22 | Bertioli DJ et al., The genome sequences of Arachis duranensis and Arachis ipaensis, the diploid ancestors of cultivated peanut. Nat Genet. 2016 Apr 48(4):438-46. Epub 2016 Feb 22. |
| 165 | *Arachis ipaensis* (Fabaceae) | 2016-02-22 | Bertioli DJ et al., The genome sequences of Arachis duranensis and Arachis ipaensis, the diploid ancestors of cultivated peanut. Nat Genet. 2016 Apr 48(4):438-46. Epub 2016 Feb 22. |
| 164 | *Rosa roxburghii* (Rosaceae) | 2016-02-05 | Lu M et al., Genome Survey Sequencing for the Characterization of the Genetic Background of Rosa roxburghii Tratt and Leaf Ascorbate Metabolism Genes. PLoS One. 2016 Feb 5 11(2):e0147530. eCollection 2016. |
| 163 | *Zostera marina* (Zosteraceae) | 2016-01-27 | Olsen JL et al., The genome of the seagrass Zostera marina reveals angiosperm adaptation to the sea. Nature. 2016 Feb 18 530(7590):331-5. Epub 2016 Jan 27. |
| 162 | *Parachlorella kessleri* (Chlorellaceae) | 2016-01-25 | Ota S et al., Highly efficient lipid production in the green alga Parachlorella kessleri : draft genome and transcriptome endorsed by whole - cell 3D ultrastructure Biotechnol Biofuels. 2016 Jan 25 9:13. eCollection 2016. |
| 161 | *Cynara cardunculus* (Asteraceae) | 2016-01-20 | Scaglione D et al., The genome sequence of the outbreeding globe artichoke constructed de novo incorporating a phase-aware low-pass sequencing strategy of F1 progeny Sci Rep. 2016 Jan 20 6:19427. |
| 160 | *Dendrobium catenatum* (Orchidaceae) | 2016-01-12 | Zhang GQ et al., The Dendrobium catenatum Lindl. genome sequence provides insights into polysaccharide synthase, floral development and adaptive evolution Sci Rep. 2016 Jan 12 6:19029. |
| 159 | *Salvia miltiorrhiza* (Lamiaceae) | 2015-12-14 | Zhang G et al., Hybrid de novo genome assembly of the Chinese herbal plant danshen (Salvia miltiorrhiza Bunge) Gigascience. 2015 Dec 14 4:62. eCollection 2015. |
| 158 | *Lemna minor* (Araceae) | 2015-11-25 | Van Hoeck A et al., The first draft genome of the aquatic model plant Lemna minor opens the route for future stress physiology research and biotechnological applications. Biotechnol Biofuels. 2015 Nov 25 8:188. eCollection 2015. |
| 157 | *Oropetium thomaeum* (Poaceae) | 2015-11-11 | VanBuren R et al., Single-molecule sequencing of the desiccation-tolerant grass Oropetium thomaeum. Nature. 2015 Nov 26 527(7579):508-11. Epub 2015 Nov 11. |
| 156 | *Ananas bracteatus* (Bromeliaceae) | 2015-11-02 | Ming R et al., The pineapple genome and the evolution of CAM photosynthesis. Nat Genet. 2015 Dec 47(12):1435-42. Epub 2015 Nov 2. |
| 155 | *Ananas comosus* (Bromeliaceae) | 2015-11-02 | Ming R et al., The pineapple genome and the evolution of CAM photosynthesis. Nat Genet. 2015 Dec 47(12):1435-42. Epub 2015 Nov 2. |
| 154 | *Oryza longistaminata* (Poaceae) | 2015-11-02 | Zhang Y et al., Genome and Comparative Transcriptomics of African Wild Rice Oryza longistaminata Provide Insights into Molecular Mechanism of Rhizomatousness and Self-Incompatibility. Mol Plant. 2015 Nov 2, 8(11): 1683-6 |
| 153 | *Auxenochlorella pyrenoidosa* (Chlorellaceae) | 2015-10-20 | Fan J et al., Genomic Foundation of Starch to Lipid Switch in Oleaginous Chlorella. Plant Physiol. 2015 Dec 169(4):2444-61. Epub 2015 Oct 20. |
| 152 | *Gossypium barbadense* (Malvaceae) | 2015-09-30 | Liu X et al., Gossypium barbadense genome sequence provides insight into the evolution of extra-long staple fiber and specialized metabolites. Sci Rep. 2015 Sep 30 5:14139. |
| 151 | *Lolium perenne* (Poaceae) | 2015-09-26 | Byrne SL et al., A synteny-based draft genome sequence of the forage grass Lolium perenne. Plant J. 2015 Nov 84(4):816-26. |
| 150 | *Cymbomonas tetramitiformis* (Pyramimonadaceae) | 2015-07-29 | Burns JA et al., Comparative Genomics of a Bacterivorous Green Alga Reveals Evolutionary Causalities and Consequences of Phago-Mixotrophic Mode of Nutrition. Genome Biol Evol. 2015 Jul 29 7(11):3047-61. |
| 149 | *Zizania latifolia* (Poaceae) | 2015-06-13 | Guo L et al., A host plant genome (Zizania latifolia) after a century-long endophyte infection Plant J. 2015 Aug 83(4):600-9. Epub 2015 Jul 7. |
| 148 | *Moringa oleifera* (Moringaceae) | 2015-06-01 | Tian Y et al., High quality reference genome of drumstick tree (Moringa oleifera Lam.), a potential perennial crop. Sci China Life Sci. 2015 Jul 58(7):627-38. Epub 2015 Jun 1. |
| 147 | *Ocimum sanctum* (Lamiaceae) | 2015-05-28 | Rastogi S et al., Unravelling the genome of Holy basil: an "incomparable" "elixir of life" of traditional Indian medicine. BMC Genomics. 2015 May 28 16:413. |
| 146 | *Picea glauca* (Pinaceae) | 2015-05-28 | Warren RL et al., Improved white spruce (Picea glauca) genome assemblies and annotation of large gene families of conifer terpenoid and phenolic defense metabolism. Plant J. 2015 Jul, 83(2): 189-212 |
| 145 | *Dorcoceras hygrometrica* (Gesneriaceae) | 2015-04-20 | Xiao L et al., The resurrection genome of Boea hygrometrica: A blueprint for survival of dehydration Proc Natl Acad Sci U S A. 2015 May 5 112(18):5833-7. Epub 2015 Apr 20. |
| 144 | *Gossypium hirsutum* (Malvaceae) | 2015-04-20 | Li F et al., Genome sequence of cultivated Upland cotton (Gossypium hirsutum TM-1) provides insights into genome evolution. Nat Biotechnol. 2015 May 33(5):524-30. Epub 2015 Apr 20. |
| 143 | *Solanum commersonii* (Solanaceae) | 2015-04-14 | Aversano R et al., The Solanum commersonii genome sequence provides insights into adaptation to stress conditions and genome evolution of wild potato relatives. Plant Cell. 2015 Apr 27(4):954-68. Epub 2015 Apr 14. |
| 142 | *Catharanthus roseus* (Apocynaceae) | 2015-04-11 | Kellner F et al., Genome-guided investigation of plant natural product biosynthesis. Plant J. 2015 May 82(4):680-92. Epub 2015 Apr 11. |
| 141 | *Ipomoea trifida* (Convolvulaceae) | 2015-03-24 | Hirakawa H et al., Survey of genome sequences in a wild sweet potato, Ipomoea trifida (H. B. K.) G. Don. DNA Res. 2015 Apr 22(2):171-9. Epub 2015 Mar 24. |
| 140 | *Vaccinium corymbosum* (Ericaceae) | 2015-02-13 | Gupta V et al., RNA-Seq analysis and annotation of a draft blueberry genome assembly identifies candidate genes involved in fruit ripening, biosynthesis of bioactive compounds, and stage-specific alternative splicing Gigascience. 2015 Feb 13, 4: 5 |
| 139 | *Arabis alpina* (Brassicaceae) | 2015-02-02 | Willing EM et al., Genome expansion of Arabis alpina linked with retrotransposition and reduced symmetric DNA methylation. Nat Plants. 2015 Feb 2, 1:14023 |
| 138 | *Vigna angularis* (Fabaceae) | 2015-01-28 | Kang YJ et al., Draft genome sequence of adzuki bean, Vigna angularis. Sci Rep. 2015 Jan 28 5:8069. |
| 137 | *Thlaspi arvense* (Brassicaceae) | 2015-01-27 | Dorn KM et al., A draft genome of field pennycress (Thlaspi arvense) provides tools for the domestication of a new winter biofuel crop. DNA Res. 2015 Apr 22(2):121-31. Epub 2015 Jan 27. |

| | plant species | published | |
|---|---|---|---|
| 136 | *Primula veris* (*Primulaceae*) | 2015-01-24 | Nowak MD et al., The draft genome of Primula veris yields insights into the molecular basis of heterostyly<br>Genome Biol. 2015 Jan 24 16:12. |
| 135 | *Dendrobium officinale* (*Orchidaceae*) | 2014-12-23 | Yan L et al., The genome of Dendrobium officinale illuminates the biology of the important traditional Chinese orchid herb.<br>Mol Plant. 2015 Jun 8(6):922-34. Epub 2014 Dec 24. |
| 134 | *Phalaenopsis equestris* (*Orchidaceae*) | 2014-11-24 | Cai J et al., The genome sequence of the orchid Phalaenopsis equestris<br>Nat Genet. 2015 Jan 47(1):65-72. Epub 2014 Nov 24. |
| 133 | *Humulus lupulus* (*Cannabaceae*) | 2014-11-20 | Natsume S et al., The Draft Genome of Hop (Humulus lupulus), an Essence for Brewing.<br>Plant Cell Physiol. 2015 Mar 56(3):428-41. Epub 2014 Nov 20. |
| 132 | *Vigna glabrescens* (*Fabaceae*) | 2014-11-11 | Kang YJ et al., Genome sequence of mungbean and insights into evolution within Vigna species<br>Nat Commun. 2014 Nov 11 5:5443. |
| 131 | *Vigna radiata* (*Fabaceae*) | 2014-11-11 | Kang YJ et al., Genome sequence of mungbean and insights into evolution within Vigna species<br>Nat Commun. 2014 Nov 11 5:5443. |
| 130 | *Oryza barthii* (*Poaceae*) | 2014-11-03 | Zhang QJ et al., Rapid diversification of five Oryza AA genomes associated with rice adaptation<br>Proc Natl Acad Sci U S A. 2014 Nov 18 111(46):E4954-62. Epub 2014 Nov 3. |
| 129 | *Oryza glumaepatula* (*Poaceae*) | 2014-11-03 | Zhang QJ et al., Rapid diversification of five Oryza AA genomes associated with rice adaptation<br>Proc Natl Acad Sci U S A. 2014 Nov 18 111(46):E4954-62. Epub 2014 Nov 3. |
| 128 | *Oryza meridionalis* (*Poaceae*) | 2014-11-03 | Zhang QJ et al., Rapid diversification of five Oryza AA genomes associated with rice adaptation<br>Proc Natl Acad Sci U S A. 2014 Nov 18 111(46):E4954-62. Epub 2014 Nov 3. |
| 127 | *Oryza nivara* (*Poaceae*) | 2014-11-03 | Zhang QJ et al., Rapid diversification of five Oryza AA genomes associated with rice adaptation<br>Proc Natl Acad Sci U S A. 2014 Nov 18 111(46):E4954-62. Epub 2014 Nov 3. |
| 126 | *Ziziphus jujuba* (*Rhamnaceae*) | 2014-10-28 | Liu MJ et al., The complex jujube genome provides insights into fruit tree biology.<br>Nat Commun. 2014 Oct 28 5:5315. |
| 125 | *Solanum melongena* (*Solanaceae*) | 2014-09-18 | Hirakawa H et al., Draft Genome Sequence of Eggplant (Solanum melongena L.): the Representative Solanum Species Indigenous to the Old World<br>DNA Res. 2014 Dec 21(6):649-60. Epub 2014 Sep 18. |
| 124 | *Glycine soja* (*Fabaceae*) | 2014-09-14 | Li YH et al., De novo assembly of soybean wild relatives for pan-genome analysis of diversity and agronomic traits.<br>Nat Biotechnol. 2014 Oct 32(10):1045-52. Epub 2014 Sep 14. |
| 123 | *Conyza canadensis* (*Asteraceae*) | 2014-09-10 | Peng Y et al., De novo genome assembly of the economically-important weed Conyza canadensis using integrated data from multiple sequencing platforms.<br>Plant Physiol. 2014 Nov 166(3):1241-54. Epub 2014 Sep 10. |
| 122 | *Coffea canephora* (*Rubiaceae*) | 2014-09-04 | Denoeud F et al., The coffee genome provides insight into the convergent evolution of caffeine biosynthesis.<br>Science. 2014 Sep 5 345(6201):1181-4. Epub 2014 Sep 4. |
| 121 | *Brassica napus* (*Brassicaceae*) | 2014-08-22 | Chalhoub B et al., Early allopolyploid evolution in the post-Neolithic Brassica napus oilseed genome.<br>Science. 2014 Aug 22 345(6199):950-3. Epub 2014 Aug 21. |
| 120 | *Amaranthus hypochondriacus* (*Amaranthaceae*) | 2014-07-28 | Sunil M et al., The Draft Genome and Transcriptome of Amaranthus hypochondriacus: A C4 Dicot Producing High-Lysine Edible Pseudo-Cereal.<br>DNA Res. 2014 Dec 21(6):585-602. Epub 2014 Jul 28. |
| 119 | *Oryza glaberrima* (*Poaceae*) | 2014-07-27 | Wang M et al., The genome sequence of African rice (Oryza glaberrima) and evidence for independent domestication.<br>Nat Genet. 2014 Sep 46(9):982-8. Epub 2014 Jul 27. |
| 118 | *Solanum pennellii* (*Solanaceae*) | 2014-07-27 | Bolger A et al., The genome of the stress-tolerant wild tomato species Solanum pennellii.<br>Nat Genet. 2014 Sep 46(9):1034-8. Epub 2014 Jul 27. |
| 117 | *Auxenochlorella protothecoides* (*Chlorellaceae*) | 2014-07-10 | Gao C et al., Oil accumulation mechanisms of the oleaginous microalga Chlorella protothecoides revealed through its genome, transcriptomes, and proteomes.<br>BMC Genomics. 2014 Jul 10 15:582. |
| 116 | *Aquilaria agallocha* (*Thymelaeaceae*) | 2014-07-09 | Chen CH et al., Identification of cucurbitacins and assembly of a draft genome for Aquilaria agallocha.<br>BMC Genomics. 2014 Jul 9 15:578. |
| 115 | *Eragrostis tef* (*Poaceae*) | 2014-07-09 | Cannarozzi G et al., Genome and transcriptome sequencing identifies breeding targets in the orphan crop tef (Eragrostis tef).<br>BMC Genomics. 2014 Jul 9 15:581. |
| 114 | *Salix suchowensis* (*Salicaceae*) | 2014-07-01 | Dai X et al., The willow genome and divergent evolution from poplar after the common genome duplication.<br>Cell Res. 2014 Oct 24(10):1274-7. Epub 2014 Jul 1. |
| 113 | *Picochlorum sp. SENEW3* (*C.incertae sedis*) | 2014-06-25 | Foflonker F et al., Genome of the halotolerant green alga Picochlorum SENEW3 reveals strategies for thriving under fluctuating environmental conditions.<br>Environ Microbiol. 2015 Feb 17(2):412-26. Epub 2014 Jul 24. |
| 112 | *Vaccinium macrocarpon* (*Ericaceae*) | 2014-06-13 | Polashock J et al., The American cranberry: first insights into the whole genome of a species adapted to bog habitat<br>BMC Plant Biol. 2014 Jun 13 14:165. |
| 111 | *Eucalyptus grandis* (*Myrtaceae*) | 2014-06-11 | Myburg AA et al., The genome of Eucalyptus grandis<br>Nature. 2014 Jun 19 510(7505):356-62. Epub 2014 Jun 11. |
| 110 | *Citrus clementina* (*Rutaceae*) | 2014-06-08 | Wu GA et al., Sequencing of diverse mandarin, pummelo and orange genomes reveals complex history of admixture during citrus domestication<br>Nat Biotechnol. 2014 Jul 32(7):656-62. Epub 2014 Jun 8. |
| 109 | *Phaseolus vulgaris* (*Fabaceae*) | 2014-06-08 | Schmutz J et al., A reference genome for common bean and genome-wide analysis of dual domestications<br>Nat Genet. 2014 Jul 46(7):707-13. Epub 2014 Jun 8. |
| 108 | *Klebsormidium flaccidum* (*Klebsormidiaceae*) | 2014-05-28 | Hori K et al., Klebsormidium flaccidum genome reveals primary factors for plant terrestrial adaptation<br>Nat Commun. 2014 May 28 5:3978. |
| 107 | *Brassica oleracea* (*Brassicaceae*) | 2014-05-23 | Liu S et al., The Brassica oleracea genome reveals the asymmetrical evolution of polyploid genomes<br>Nat Commun. 2014 May 23 5:3930. |
| 106 | *Gossypium arboreum* (*Malvaceae*) | 2014-05-18 | Li F et al., Genome sequence of the cultivated cotton Gossypium arboreum<br>Nat Genet. 2014 Jun 46(6):567-72. Epub 2014 May 18. |
| 105 | *Raphanus raphanistrum* (*Brassicaceae*) | 2014-05-16 | Kitashiba H et al., Draft Sequences of the Radish (Raphanus sativus L.) Genome<br>DNA Res. 2014 Oct 21(5):481-90. Epub 2014 May 16. |
| 104 | *Helicosporidium sp.* (*Chlorellaceae*) | 2014-05-08 | Pombert JF et al., A lack of parasitic reduction in the obligate parasitic green alga Helicosporidium.<br>PLoS Genet. 2014 May 8 10(5):e1004355. eCollection 2014 May. |
| 103 | *Nicotiana otophora* (*Solanaceae*) | 2014-05-08 | Sierro N et al., The tobacco genome sequence and its comparison with those of tomato and potato.<br>Nat Commun. 2014 May 8 5:3833. |
| 102 | *Nicotiana tabacum* (*Solanaceae*) | 2014-05-08 | Sierro N et al., The tobacco genome sequence and its comparison with those of tomato and potato.<br>Nat Commun. 2014 May 8 5:3833. |
| 101 | *Camelina sativa* (*Brassicaceae*) | 2014-04-23 | Kagale S et al., The emerging biofuel crop Camelina sativa retains a highly undifferentiated hexaploid genome structure<br>Nat Commun. 2014 Apr 23 5:3706. |
| 100 | *Pyrus communis* (*Rosaceae*) | 2014-04-03 | Chagne D et al., The Draft Genome Sequence of European Pear (Pyrus communis L. Bartlett).<br>PLoS One. 2014 Apr 3 9(4):e92644. eCollection 2014. |

| | plant species | published | |
|---|---|---|---|
| 99 | *Pinus taeda* (*Pinaceae*) | 2014-03-20 | Neale DB et al., Decoding the massive genome of loblolly pine using haploid DNA and novel assembly strategies. Genome Biol. 2014 Mar 4 15(3):R59. |
| 98 | *Sesamum indicum* (*Pedaliaceae*) | 2014-02-27 | Wang L et al., Genome sequencing of the high oil crop sesame provides insight into oil biosynthesis Genome Biol. 2014 Feb 27 15(2):R39. |
| 97 | *Spirodela polyrhiza* (*Araceae*) | 2014-02-19 | Wang W et al., The Spirodela polyrhiza genome reveals insights into its neotenous reduction fast growth and aquatic lifestyle. Nat Commun. 2014 5:3311. |
| 96 | *Trifolium pratense* (*Fabaceae*) | 2014-02-05 | Istvanek J1 et al., Genome assembly and annotation for red clover (Trifolium pratense Am J Bot. 2014 Feb, 101(2): 327-37 Epub 2014 Feb 5. |
| 95 | *Capsicum annuum* (*Solanaceae*) | 2014-01-19 | Kim S et al., Genome sequence of the hot pepper provides insights into the evolution of pungency in Capsicum species. Nat Genet. 2014 Mar 46(3):270-8. Epub 2014 Jan 19. |
| 94 | *Ensete ventricosum* (*Musaceae*) | 2014-01-17 | Harrison J et al., A Draft Genome Sequence for Ensete ventricosum, the Drought-Tolerant 'Tree Against Hunger'. Agronomy 2014, 4, 13-33 |
| 93 | *Arabidopsis halleri* (*Brassicaceae*) | 2014-01-13 | Akama S et al., Genome-wide quantification of homeolog expression ratio revealed nonstochastic gene regulation in synthetic allopolyploid Arabidopsis. Nucleic Acids Res. 2014 Apr, 42(6): e46, Epub 2014 Jan 13. |
| 92 | *Monoraphidium neglectum* (*Selenastraceae*) | 2013-12-28 | Bogen C et al., Reconstruction of the lipid metabolism for the microalga Monoraphidium neglectum from its genome sequence reveals characteristics suitable for biofuel production. BMC Genomics. 2013 Dec 28 14:926. |
| 91 | *Amborella trichopoda* (*Amborellaceae*) | 2013-12-20 | Amborella Genome Project et al., The Amborella genome and the evolution of flowering plants. Science. 2013 Dec 20 342(6165):1241089. |
| 90 | *Beta vulgaris* (*Amaranthaceae*) | 2013-12-18 | Dohm JC et al., The genome of the recently domesticated crop plant sugar beet (Beta vulgaris). Nature. 2014 Jan 23 505(7484):546-9. Epub 2013 Dec 18. |
| 89 | *Dianthus caryophyllus* (*Caryophyllaceae*) | 2013-12-17 | Yagi M et al., Sequence Analysis of the Genome of Carnation (Dianthus caryophyllus L.). DNA Res. 2014 Jun 21(3):231-41. Epub 2013 Dec 17. |
| 88 | *Erythranthe guttatus* (*Phrymaceae*) | 2013-11-26 | Hellsten U et al., Fine-scale variation in meiotic recombination in Mimulus inferred from population shotgun sequencing. Proc Natl Acad Sci U S A. 2013 Nov 26, 110(48): 19478-82 |
| 87 | *Populus euphratica* (*Salicaceae*) | 2013-11-21 | Ma T et al., Genomic insights into salt adaptation in a desert poplar. Nat Commun. 2013, 4: 2797 |
| 86 | *Actinidia chinensis* (*Actinidiaceae*) | 2013-10-18 | Huang S et al., Draft genome of the kiwifruit Actinidia chinensis. Nat Commun. 2013 4:2640. |
| 85 | *Galdieria phlegrea* (*Galdieriaceae*) | 2013-10-07 | Qiu H et al., Adaptation through horizontal gene transfer in the cryptoendolithic red alga Galdieria phlegrea. Curr Biol. 2013 Oct 7 23(19):R865-6. |
| 84 | *Musa balbisiana* (*Musaceae*) | 2013-10-05 | Davey MW et al., A draft Musa balbisiana genome sequence for molecular genetics in polyploid, inter- and intra-specific Musa hybrids. BMC Genomics. 2013 Oct 5 14:683. |
| 83 | *Morus notabilis* (*Moraceae*) | 2013-09-19 | He N et al., Draft genome sequence of the mulberry tree Morus notabilis. Nat Commun. 2013 4:2445. |
| 82 | *Tarenaya hassleriana* (*Cleomaceae*) | 2013-08-27 | Cheng S et al., The Tarenaya hassleriana genome provides insight into reproductive trait and genome evolution of crucifers. Plant Cell. 2013 Aug 25(8):2813-30. Epub 2013 Aug 27. |
| 81 | *Elaeis guineensis* (*Arecaceae*) | 2013-07-24 | Singh R et al., Oil palm genome sequence reveals divergence of interfertile species in Old and New worlds. Nature. 2013 Aug 15 500(7462):335-9. Epub 2013 Jul 24. |
| 80 | *Gracilariopsis lemaneiformis* (*Gracilariaceae*) | 2013-07-16 | Zhou W et al., Genome survey sequencing and genetic background characterization of Gracilariopsis lemaneiformis (Rhodophyta) based on next-generation sequencing. PLoS One. 2013 Jul 16 8(7):e69909. Print 2013. |
| 79 | *Genlisea aurea* (*Lentibulariaceae*) | 2013-07-15 | Leushkin EV et al., The miniature genome of a carnivorous plant Genlisea aurea contains a low number of genes and short non-coding sequences. BMC Genomics. 2013 Jul 15 14:476. |
| 78 | *Aethionema arabicum* (*Brassicaceae*) | 2013-06-30 | Haudry A et al., An atlas of over 90,000 conserved noncoding sequences provides insight into crucifer regulatory regions. Nat Genet. 2013 Aug 45(8):891-8. Epub 2013 Jun 30. |
| 77 | *Leavenworthia alabamica* (*Brassicaceae*) | 2013-06-30 | Haudry A et al., An atlas of over 90,000 conserved noncoding sequences provides insight into crucifer regulatory regions. Nat Genet. 2013 Aug 45(8):891-8. Epub 2013 Jun 30. |
| 76 | *Sisymbrium irio* (*Brassicaceae*) | 2013-06-30 | Haudry A et al., An atlas of over 90,000 conserved noncoding sequences provides insight into crucifer regulatory regions. Nat Genet. 2013 Aug 45(8):891-8. Epub 2013 Jun 30. |
| 75 | *Nicotiana sylvestris* (*Solanaceae*) | 2013-06-17 | Sierro N et al., Reference genomes and transcriptomes of Nicotiana sylvestris and Nicotiana tomentosiformis Genome Biol. 2013 Jun 17 14(6):R60. |
| 74 | *Nicotiana tomentosiformis* (*Solanaceae*) | 2013-06-17 | Sierro N et al., Reference genomes and transcriptomes of Nicotiana sylvestris and Nicotiana tomentosiformis Genome Biol. 2013 Jun 17 14(6):R60. |
| 73 | *Porphyridium purpureum* (*Porphyridiaceae*) | 2013-06-17 | Bhattacharya D et al., Genome of the red alga Porphyridium purpureum Nat Commun. 2013 4:1941. |
| 72 | *Capsella grandiflora* (*Brassicaceae*) | 2013-06-09 | Slotte T et al., The Capsella rubella genome and the genomic consequences of rapid mating system evolution. Nat Genet. 2013 Jul 45(7):831-5. Epub 2013 Jun 9. |
| 71 | *Capsella rubella* (*Brassicaceae*) | 2013-06-09 | Slotte T et al., The Capsella rubella genome and the genomic consequences of rapid mating system evolution. Nat Genet. 2013 Jul 45(7):831-5. Epub 2013 Jun 9. |
| 70 | *Lupinus angustifolius* (*Fabaceae*) | 2013-05-29 | Yang H et al., Draft genome sequence, and a sequence-defined genetic linkage map of the legume crop species Lupinus angustifolius L. PLoS One. 2013 May 29 8(5):e64799. Print 2013. |
| 69 | *Picea abies* (*Pinaceae*) | 2013-05-22 | Nystedt B et al., The Norway spruce genome sequence and conifer genome evolution. Nature. 2013 May 30 497(7451):579-84. Epub 2013 May 22. |
| 68 | *Utricularia gibba* (*Lentibulariaceae*) | 2013-05-12 | Ibarra-Laclette E et al., Architecture and evolution of a minute plant genome. Nature. 2013 Jun 6 498(7452):94-8. Epub 2013 May 12. |
| 67 | *Nelumbo nucifera* (*Nelumbonaceae*) | 2013-05-10 | Ming R et al., Genome of the long-living sacred lotus (Nelumbo nucifera Gaertn.). Genome Biol. 2013 May 10 14(5):R41. |
| 66 | *Aegilops tauschii* (*Poaceae*) | 2013-03-24 | Jia J et al., Aegilops tauschii draft genome sequence reveals a gene repertoire for wheat adaptation. Nature. 2013 Apr 4 496(7443):91-5. Epub 2013 Mar 24. |
| 65 | *Prunus persica* (*Rosaceae*) | 2013-03-24 | Verde I et al., The high-quality draft genome of peach (Prunus persica) identifies unique patterns of genetic diversity, domestication and genome evolution. Nat Genet. 2013 May 45(5):487-94. Epub 2013 Mar 24. |
| 64 | *Triticum urartu* (*Poaceae*) | 2013-03-24 | Ling HQ et al., Draft genome of the wheat A-genome progenitor Triticum urartu. Nature. 2013 Apr 4 496(7443):87-90. Epub 2013 Mar 24. |
| 63 | *Chondrus crispus* (*Gigartinaceae*) | 2013-03-15 | Collen J et al., Genome structure and metabolic features in the red seaweed Chondrus crispus shed light on evolution of the Archaeplastida. Proc Natl Acad Sci U S A. 2013 Mar 26 110(13):5247-52. Epub 2013 Mar 15. |

| | plant species | published | |
|---|---|---|---|
| 62 | *Oryza brachyantha* *(Poaceae)* | 2013-03-12 | Chen J et al., Whole-genome sequencing of Oryza brachyantha reveals mechanisms underlying Oryza genome evolution. Nat Commun. 2013 4:1595. |
| 61 | *Neopyropia yezoensis* *(Bangiaceae)* | 2013-03-11 | Nakamura Y et al., The first symbiont-free genome sequence of marine red alga, Susabi-nori (Pyropia yezoensis). PLoS One. 2013 8(3):e57122. Epub 2013 Mar 11. |
| 60 | *Galdieria sulphuraria* *(Galdieriaceae)* | 2013-03-08 | Schonknecht G et al., Gene transfer from bacteria and archaea facilitated evolution of an extremophilic eukaryote. Science. 2013 Mar 8 339(6124):1207-10. |
| 59 | *Phyllostachys edulis* *(Poaceae)* | 2013-02-24 | Peng Z et al., The draft genome of the fast-growing non-timber forest species moso bamboo (Phyllostachys heterocycla). Nat Genet. 2013 Apr 45(4):456-61, 461e1-2. Epub 2013 Feb 24. |
| 58 | *Hevea brasiliensis* *(Euphorbiaceae)* | 2013-02-02 | Rahman AY et al., Draft genome sequence of the rubber tree Hevea brasiliensis BMC Genomics. 2013 Feb 2 14:75. |
| 57 | *Cicer arietinum* *(Fabaceae)* | 2013-01-27 | Varshney RK et al., Draft genome sequence of chickpea (Cicer arietinum) provides a resource for trait improvement. Nat Biotechnol. 2013 Mar 31(3):240-6. Epub 2013 Jan 27. |
| 56 | *Prunus mume* *(Rosaceae)* | 2012-12-27 | Zhang Q et al., The genome of Prunus mume. Nat Commun. 2012 3:1318. |
| 55 | *Nicotiana benthamiana* *(Solanaceae)* | 2012-12-01 | Bombarely A et al., A draft genome sequence of Nicotiana benthamiana to enhance molecular plant-microbe biology research. Mol Plant Microbe Interact. 2012 Dec 25(12):1523-30. |
| 54 | *Triticum aestivum* *(Poaceae)* | 2012-11-29 | Brenchley R et al., Analysis of the bread wheat genome using whole-genome shotgun sequencing. Nature. 2012 Nov 29 491(7426):705-10. |
| 53 | *Citrullus lanatus* *(Cucurbitaceae)* | 2012-11-25 | Guo S et al., The draft genome of watermelon (Citrullus lanatus) and resequencing of 20 diverse accessions. Nat Genet. 2013 Jan 45(1):51-8. Epub 2012 Nov 25. |
| 52 | *Citrus sinensis* *(Rutaceae)* | 2012-11-25 | Xu Q et al., The draft genome of sweet orange (Citrus sinensis). Nat Genet. 2013 Jan 45(1):59-66. Epub 2012 Nov 25. |
| 51 | *Betula nana* *(Betulaceae)* | 2012-11-21 | Wang N et al., Genome sequence of dwarf birch (Betula nana) and cross-species RAD markers. Mol Ecol. 2013 Jun 22(11):3098-111. Epub 2012 Nov 21. |
| 50 | *Pyrus bretschneideri* *(Rosaceae)* | 2012-11-13 | Wu J et al., The genome of the pear (Pyrus bretschneideri Rehd.). Genome Res. 2013 Feb 23(2):396-408. Epub 2012 Nov 13. |
| 49 | *Hordeum vulgare* *(Poaceae)* | 2012-10-17 | Mayer KF et al., A physical, genetic and functional sequence assembly of the barley genome. Nature. 2012 Nov 29 491(7426):711-6. Epub 2012 Oct 17. |
| 48 | *Azadirachta indica* *(Meliaceae)* | 2012-09-09 | Krishnan NM et al., A draft of the genome and four transcriptomes of a medicinal and pesticidal angiosperm Azadirachta indica. BMC Genomics. 2012 Sep 9 13:464. |
| 47 | *Gossypium raimondii* *(Malvaceae)* | 2012-08-26 | Wang K et al., The draft genome of a diploid cotton Gossypium raimondii. Nat Genet. 2012 Oct 44(10):1098-103. Epub 2012 Aug 26. |
| 46 | *Bathycoccus prasinos* *(Bathycoccaceae)* | 2012-08-24 | Moreau H et al., Gene functionalities and genome structure in Bathycoccus prasinos reflect cellular specializations at the base of the green lineage. Genome Biol. 2012 Aug 24 13(8):R74. |
| 45 | *Linum usitatissimum* *(Linaceae)* | 2012-08-14 | Wang Z et al., The genome of flax (Linum usitatissimum) assembled de novo from short shotgun sequence reads. Plant J. 2012 Nov 72(3):461-73. Epub 2012 Aug 14. |
| 44 | *Musa acuminata* *(Musaceae)* | 2012-08-09 | D'Hont A et al., The banana (Musa acuminata) genome and the evolution of monocotyledonous plants. Nature. 2012 Aug 9 488(7410):213-7. |
| 43 | *Eutrema salsugineum* *(Brassicaceae)* | 2012-07-24 | Wu HJ et al., Insights into salt tolerance from the genome of Thellungiella salsuginea. Proc Natl Acad Sci U S A. 2012 Jul 24 109(30):12219-24. |
| 42 | *Cucumis melo* *(Cucurbitaceae)* | 2012-07-02 | Garcia-Mas J et al., The genome of melon (Cucumis melo L.). Proc Natl Acad Sci U S A. 2012 Jul 17 109(29):11872-7. Epub 2012 Jul 2. |
| 41 | *Solanum lycopersicum* *(Solanaceae)* | 2012-05-30 | Tomato Genome Consortium et al., The tomato genome sequence provides insights into fleshy fruit evolution. Nature. 2012 May 30 485(7400):635-41. |
| 40 | *Solanum pimpinellifolium* *(Solanaceae)* | 2012-05-30 | Tomato Genome Consortium et al., The tomato genome sequence provides insights into fleshy fruit evolution. Nature. 2012 May 30 485(7400):635-41. |
| 39 | *Coccomyxa subellipsoidea* *(Coccomyxaceae)* | 2012-05-25 | Blanc G et al., The genome of the polar eukaryotic microalga Coccomyxa subellipsoidea reveals traits of cold adaptation. Genome Biol. 2012 May 25 13(5):R39. |
| 38 | *Setaria italica* *(Poaceae)* | 2012-05-13 | Zhang G et al., Genome sequence of foxtail millet (Setaria italica) provides insights into grass evolution and biofuel potential. Nat Biotechnol. 2012 May 13 30(6):549-54. |
| 37 | *Cyanophora paradoxa* *(Glaucocystaceae)* | 2012-02-17 | Price DC et al., Cyanophora paradoxa genome elucidates origin of photosynthesis in algae and plants. Science. 2012 Feb 17 335(6070):843-7. |
| 36 | *Manihot esculenta* *(Euphorbiaceae)* | 2012-01-05 | Prochnik S et al., The Cassava Genome: Current Progress, Future Directions. Trop Plant Biol. 2012 Mar 5(1):88-94. Epub 2012 Jan 5. |
| 35 | *Eucalyptus camaldulensis* *(Myrtaceae)* | 2011-12-25 | Hirakawa H et al., Survey of the genetic information carried in the genome of Eucalyptus camaldulensis. Plant Biotechnol. 2011 Dec 25, 28(5): 471-480 |
| 34 | *Medicago truncatula* *(Fabaceae)* | 2011-11-16 | Young ND et al., The Medicago genome provides insight into the evolution of rhizobial symbioses. Nature. 2011 Nov 16 480(7378):520-4. |
| 33 | *Cajanus cajan* *(Fabaceae)* | 2011-11-06 | Varshney RK et al., Draft genome sequence of pigeonpea (Cajanus cajan), an orphan legume crop of resource-poor farmers. Nat Biotechnol. 2011 Nov 6 30(1):83-9. |
| 32 | *Cannabis sativa* *(Cannabaceae)* | 2011-10-20 | van Bakel H et al., The draft genome and transcriptome of Cannabis sativa. Genome Biol. 2011 Oct 20 12(10):R102. |
| 31 | *Brassica rapa* *(Brassicaceae)* | 2011-08-28 | Wang X et al., The genome of the mesopolyploid crop species Brassica rapa. Nat Genet. 2011 Aug 28 43(10):1035-9. |
| 30 | *Thellungiella parvula* *(Brassicaceae)* | 2011-08-07 | Dassanayake M et al., The genome of the extremophile crucifer Thellungiella parvula. Nat Genet. 2011 Aug 7 43(9):913-8. |
| 29 | *Solanum tuberosum* *(Solanaceae)* | 2011-07-10 | Xu X et al., Genome sequence and analysis of the tuber crop potato. Nature. 2011 Jul 10 475(7355):189-95. |
| 28 | *Phoenix dactylifera* *(Arecaceae)* | 2011-05-29 | Al-Dous EK et al., De novo genome sequencing and comparative genomics of date palm (Phoenix dactylifera). Nat Biotechnol. 2011 May 29 29(6):521-7. |
| 27 | *Selaginella moellendorffii* *(Selaginellaceae)* | 2011-05-05 | Banks JA et al., The Selaginella genome identifies genetic changes associated with the evolution of vascular plants. Science. 2011 May 20 332(6032):960-3. Epub 2011 May 5. |
| 26 | *Arabidopsis lyrata* *(Brassicaceae)* | 2011-04-10 | Hu TT et al., The Arabidopsis lyrata genome sequence and the basis of rapid genome size change. Nat Genet. 2011 May 43(5):476-81. Epub 2011 Apr 10. |

| | plant species | published | |
|---|---|---|---|
| 25 | *Fragaria vesca*<br>*(Rosaceae)* | 2010-12-26 | Shulaev V et al., The genome of woodland strawberry (Fragaria vesca).<br>Nat Genet. 2011 Feb 43(2):109-16. Epub 2010 Dec 26. |
| 24 | *Theobroma cacao*<br>*(Malvaceae)* | 2010-12-26 | Argout X et al., The genome of Theobroma cacao.<br>Nat Genet. 2011 Feb 43(2):101-8. Epub 2010 Dec 26. |
| 23 | *Jatropha curcas*<br>*(Euphorbiaceae)* | 2010-12-13 | Sato S et al., Sequence analysis of the genome of an oil-bearing tree, Jatropha curcas L.<br>DNA Res. 2011 Feb 18(1):65-76. Epub 2010 Dec 13. |
| 22 | *Chlorella variabilis*<br>*(Chlorellaceae)* | 2010-09-17 | Blanc G et al., The Chlorella variabilis NC64A genome reveals adaptation to photosymbiosis, coevolution with viruses, and cryptic sex.<br>Plant Cell. 2010 Sep 22(9):2943-55. Epub 2010 Sep 17. |
| 21 | *Malus domestica*<br>*(Rosaceae)* | 2010-08-29 | Velasco R et al., The genome of the domesticated apple (Malus x domestica Borkh.).<br>Nat Genet. 2010 Oct 42(10):833-9. Epub 2010 Aug 29. |
| 20 | *Ricinus communis*<br>*(Euphorbiaceae)* | 2010-08-22 | Chan AP et al., Draft genome sequence of the oilseed species Ricinus communis.<br>Nat Biotechnol. 2010 Sep 28(9):951-6. Epub 2010 Aug 22. |
| 19 | *Volvox carteri*<br>*(Volvocaceae)* | 2010-07-09 | Prochnik SE et al., Genomic analysis of organismal complexity in the multicellular green alga Volvox carteri.<br>Science. 2010 Jul 9 329(5988):223-6. |
| 18 | *Brachypodium distachyon*<br>*(Poaceae)* | 2010-02-11 | International Brachypodium Initiative et al., Genome sequencing and analysis of the model grass Brachypodium distachyon.<br>Nature. 2010 Feb 11 463(7282):763-8. |
| 17 | *Glycine max*<br>*(Fabaceae)* | 2010-01-14 | Schmutz J et al., Genome sequence of the palaeopolyploid soybean.<br>Nature. 2010 Jan 14 463(7278):178-83. |
| 16 | *Zea mays*<br>*(Poaceae)* | 2009-11-20 | Schnable PS et al., The B73 maize genome: complexity, diversity, and dynamics.<br>Science. 2009 Nov 20 326(5956):1112-5. |
| 15 | *Cucumis sativus*<br>*(Cucurbitaceae)* | 2009-11-01 | Huang S et al., The genome of the cucumber, Cucumis sativus L.<br>Nat Genet. 2009 Dec 41(12):1275-81. Epub 2009 Nov 1. |
| 14 | *Micromonas commoda*<br>*(Mamiellaceae)* | 2009-04-10 | Worden AZ et al., Green evolution and dynamic adaptations revealed by genomes of the marine picoeukaryotes Micromonas.<br>Science. 2009 Apr 10 324(5924):268-72. |
| 13 | *Micromonas pusilla*<br>*(Mamiellaceae)* | 2009-04-10 | Worden AZ et al., Green evolution and dynamic adaptations revealed by genomes of the marine picoeukaryotes Micromonas.<br>Science. 2009 Apr 10 324(5924):268-72. |
| 12 | *Sorghum bicolor*<br>*(Poaceae)* | 2009-01-29 | Paterson AH et al., The Sorghum bicolor genome and the diversification of grasses.<br>Nature. 2009 Jan 29 457(7229):551-6. |
| 11 | *Lotus japonicus*<br>*(Fabaceae)* | 2008-05-28 | Sato S et al., Genome structure of the legume, Lotus japonicus.<br>DNA Res. 2008 Aug 15(4):227-39. Epub 2008 May 28. |
| 10 | *Carica papaya*<br>*(Caricaceae)* | 2008-04-24 | Ming R et al., The draft genome of the transgenic tropical fruit tree papaya (Carica papaya Linnaeus).<br>Nature. 2008 Apr 24 452(7190):991-6. |
| 9 | *Physcomitrium patens*<br>*(Funariaceae)* | 2007-12-13 | Rensing SA et al., The Physcomitrella genome reveals evolutionary insights into the conquest of land by plants.<br>Science. 2008 Jan 4 319(5859):64-9. Epub 2007 Dec 13. |
| 8 | *Chlamydomonas reinhardtii*<br>*(Chlamydomonadaceae)* | 2007-10-12 | Merchant SS et al., The Chlamydomonas genome reveals the evolution of key animal and plant functions.<br>Science. 2007 Oct 12 318(5848):245-50. |
| 7 | *Vitis vinifera*<br>*(Vitaceae)* | 2007-08-26 | Jaillon O et al., The grapevine genome sequence suggests ancestral hexaploidization in major angiosperm phyla.<br>Nature. 2007 Sep 27 449(7161):463-7. Epub 2007 Aug 26. |
| 6 | *Ostreococcus lucimarinus*<br>*(Bathycoccaceae)* | 2007-04-25 | Palenik B et al., The tiny eukaryote Ostreococcus provides genomic insights into the paradox of plankton speciation.<br>Proc Natl Acad Sci U S A. 2007 May 1 104(18):7705-10. Epub 2007 Apr 25. |
| 5 | *Populus trichocarpa*<br>*(Salicaceae)* | 2006-09-15 | Tuskan GA et al., The genome of black cottonwood, Populus trichocarpa (Torr. & Gray).<br>Science. 2006 Sep 15 313(5793):1596-604. |
| 4 | *Ostreococcus tauri*<br>*(Bathycoccaceae)* | 2006-07-25 | Derelle E et al., Genome analysis of the smallest free-living eukaryote Ostreococcus tauri unveils many unique features.<br>Proc Natl Acad Sci U S A. 2006 Aug 1 103(31):11647-52. Epub 2006 Jul 25. |
| 3 | *Cyanidioschyzon merolae*<br>*(Cyanidioschyzonaceae)* | 2004-04-08 | Matsuzaki M et al., Genome sequence of the ultrasmall unicellular red alga Cyanidioschyzon merolae 10D.<br>Nature. 2004 Apr 8 428(6983):653-7. |
| 2 | *Oryza sativa*<br>*(Poaceae)* | 2002-04-05 | Goff SA et al., A draft sequence of the rice genome (Oryza sativa L. ssp. japonica).<br>Science. 2002 Apr 5 296(5565):92-100. |
| 1 | *Arabidopsis thaliana*<br>*(Brassicaceae)* | 2000-12-14 | Arabidopsis Genome Initiative et al., Analysis of the genome sequence of the flowering plant Arabidopsis thaliana.<br>Nature. 2000 Dec 14 408(6814):796-815. |
